# A Versatile Marker-Free Genome Engineering Platform to Overcome Homologous Recombination Bias in Microbes: A Case Study in *Synechococcus elongatus* UTEX 2973

**DOI:** 10.1101/2025.04.27.650839

**Authors:** Shubin Li, Tao Sun, Dailin Liu, Tong Zhang, Chen Lei, Weiwen Zhang

## Abstract

The single crossover occurring via homologous recombination is a common phenomenon exited among most microbes like *Escherichia, Clostridium, Streptococcus, Lactobacillus*, and cyanobacteria, threatening the stability of engineered strains. Among them, fast-growing cyanobacterium *Synechococcus elongatus* UTEX 2973 (Syn2973) is an attractive photosynthetic chassis for CO_2_ bioconversion. To address the challenge of homologous recombination, we constructed two endogenous plasmid-based shuttle vectors, pSES and pSEL, enabling stable DNA delivery without reliance on homologous recombination. In parallel, two robust counter-selection systems were developed based on *sepT_2_* and *rpsL*, which, in combination with positive selection markers, significantly improved the screening efficiency of double-crossover mutants. Building upon these tools, we established three marker-free platforms: (i) T4CROSS, which employs two plasmids and four rounds of single crossover; (ii) TRIPLEARM, which uses a single plasmid containing three homologous arms for three rounds of single crossover; and (iii) CRISPRARM, which integrates CRISPR/Cpf1-mediated genome editing with homologous recombination. All three methods successfully repaired mutations in the *pilMNOQ* pilus gene cluster in Syn2973, restoring natural competence with high efficiency and positive selection rates. As proof of concept, we employed the CRISPRARM platform for a three-step sequential engineering of the sucrose biosynthetic pathway. The final engineered strain produced 7.12 g·L^-1^ of sucrose within four days.

## Introduction

Homologous recombination is a widely used strategy for precise genome modification in microbes, enabling accurate and site-specific exchange of genetic material between homologous DNA sequences (1). Typically, a suicide vector carrying the desired cargo cassette flanked by homologous arms is introduced into the target microbe via electroporation or conjugation. The recombination process ideally proceeds through two sequential crossover events: an initial single crossover, followed by a second recombination event that results in a double crossover and stable genomic integration. However, in many cases including *Lactococcus lactis* (2), *Clostridium acetobutylicum* (3), *Anabaena* (*Nostoc*) (4) and etc. the second crossover is inefficient, leading to the persistence of single-crossover intermediates. This partial integration is problematic, as it compromises the genetic stability of engineered strains and hinders applications such as clean gene knockouts or iterative genome modifications. Moreover, in polyploid microbes, residual plasmid backbone sequences retained after single crossover events may serve as unintended homologous targets in subsequent integrations, further complicating genetic engineering efforts. Among them, cyanobacteria are the only prokaryotes capable of oxygenic photosynthesis, contributing 20% to 30% of global CO_2_ fixation (5). This unique capability makes them attractive chassis for developing photosynthetic cell factories, especially in the pursuit of carbon neutrality. Notably, various high-value chemicals have been successfully produced using several model cyanobacteria such as *Synechocystis* sp. PCC 6803 (hereafter Syn6803) and *Synechococcus elongatus* PCC 7942 (hereafter Syn7942), including biofuels including biofuels (e.g., ethanol (6)), terpene compounds (e.g., farnesene (7)), carotenoids (e.g., astaxanthin (8)), and long-chain hydrocarbons (e.g., fatty acids (9)). However, the relatively slow growth and limited biomass accumulation of these strains have hindered their industrial scalability (10). *Synechococcus elongatus* UTEX 2973 (hereafter Syn2973) is a recently identified, fast-growing cyanobacterial strain (11). Although its genome shares 99.8% sequence identity with Syn7942 (12), it exhibits a significantly shorter doubling time of just 1.5 hours under optimal conditions (13). Moreover, Syn2973 shows enhanced tolerance to high light intensities (up to 1500 μmol photons·m^-2^·s^-1^) and elevated temperatures (up to 42 °C). Its potential as a photosynthetic production platform has been demonstrated, with engineered strains achieving sucrose titers of up to 8 g·L^-1^ (1.9 g·L^-1^·day^-1^) under the optimized conditions (14)—more than double the yields obtained with traditional strains. Additionally, Syn2973 can reach a dry cell weight of 23.41 g·L^-^¹ under semi-continuous cultivation, with a productivity of 2.4 g·L^-^¹·day^-^¹ (15). Despite these advantages, iterative genetic engineering of Syn2973 remains challenging especially considering its genome polyploid and limited efficiency of double cross over after single cross-over recombination (16,17). This underscores the urgent need for efficient, marker-free genome editing strategies in this promising organism.

Marker-free genetic manipulation in cyanobacteria generally follows two main strategies: *i*) two-step homologous recombination using a dual-selection cassette (typically combining an antibiotic resistance gene and a counter-selection marker), and *ii*) CRISPR/Cas-mediated genome editing. For the first approach, *sacB*, encoding *Bacillus subtilis* levansucrase, is commonly used as a counter-selection marker. It converts sucrose into levan, which is toxic to cyanobacterial cells such as Syn6803. Using a *sacB-Km^R^* cassette, Liu et al. achieved six rounds of iterative genome editing in Syn6803 for fatty acid biosynthesis (18). The *sacB-Km^R^* cassette is first integrated at the target locus via antibiotic selection, followed by its replacement with a gene of interest (GOI) in a second round using counter-selection. Viola et al. later simplified this process to a single transformation by designing constructs with dual homologous arms, enabling two recombination events (single and double crossover) in Syn6803 (19). However, this method is not directly transferrable to Syn2973 because, first, *sacB* has been shown to be ineffective in Syn2973 (17), and second, homologous recombination in Syn2973 predominantly results in single-crossover events, significantly reducing the frequency and stability of double-crossover recombinants (17). To overcome these limitations, Chen et al. have screened new counter-selection markers compatible with Syn2973 and achieved marker-free genome editing via homologous recombination (17). Nevertheless, this strategy remains underdeveloped, and efficient, broadly applicable methods for iterative, marker-free engineering in Syn2973 are still lacking. The second strategy employs CRISPR-based systems, including Cas9, Cpf1 (Cas12a), and Cas3, which have been applied to Syn2973 genome editing (20). For example, Wendt et al. achieved marker-free gene knockouts by transient expression of Cas9 (21), while Racharaks et al. developed CRISPR/Cas9-based knock-in tools for free fatty acid production by optimizing transformation protocols and sgRNA design (22). However, Cas9 exhibits high cytotoxicity in Syn2973, limiting its utility—especially considering the polyploid nature of cyanobacteria. Transient or low-level Cas9 expression often results in incomplete editing and low rates of homozygous transformants. Using CRISPR/Cpf1, Ungerer conducted a genome-wide mutational analysis in Syn2973 and identified three key genes (*atpA, ppnK*, and *rpaA*) with SNPs that enhance growth rates (23). However, constructing CRISPR plasmids in Syn2973 remains technically challenging. For instance, plasmids based on the RSF1010 origin exhibit low transformation efficiency in Syn2973 (24), as also confirmed in this study.

To address the limitations, we systematically optimized marker-free genetic manipulation strategies in Syn2973 from the following aspects: *i*) Optimization of stable shuttle plasmids (**Fig. 1A**): We constructed two shuttle vectors, pSES and pSEL, based on the endogenous plasmids pANS and pANL of Syn2973. In addition, a re-engineered plasmid (pRSF), derived from the RSF1010 origin of replication, was developed to enhance transformation efficiency. *ii*) Screening of effective counter-selection markers (**Fig. 1B**): We evaluated four commonly used counter-selection markers—*sacB* (25), *rpsl_12_* (26), *sepT_2_* (27), and *tetA* (28)—in Syn2973. Among them, *rpsl_12_* and *sepT_2_*demonstrated reliable performance in enabling counter-selection. *iii*) Development of marker-free genetic manipulation tools (**Figs. 1C-1E**): Three novel strategies were established for efficient, marker-free genome editing in fast-growing cyanobacterium Syn2973: T4CROSS, which employs two plasmids for four rounds of single-crossover recombination; TRIPLEARM, which uses a single plasmid containing three homologous arms to facilitate iterative recombination; CRISPRARM, which combines CRISPR/Cpf1 with homologous recombination to enable precise genome modifications. *iv*) Iterative marker-free gene editing in Syn2973 (**Fig. 1F**): Using sucrose biosynthesis as a proof-concept, we applied the CRISPRARM system in a three-step iterative editing workflow. The resulting strain achieved a sucrose titer of 7.12 g·L^-1^ within four days, demonstrating both the feasibility and high efficiency of the system for sequential genome engineering in fast-growing cyanobacterium Syn973.

**Fig. 1.**
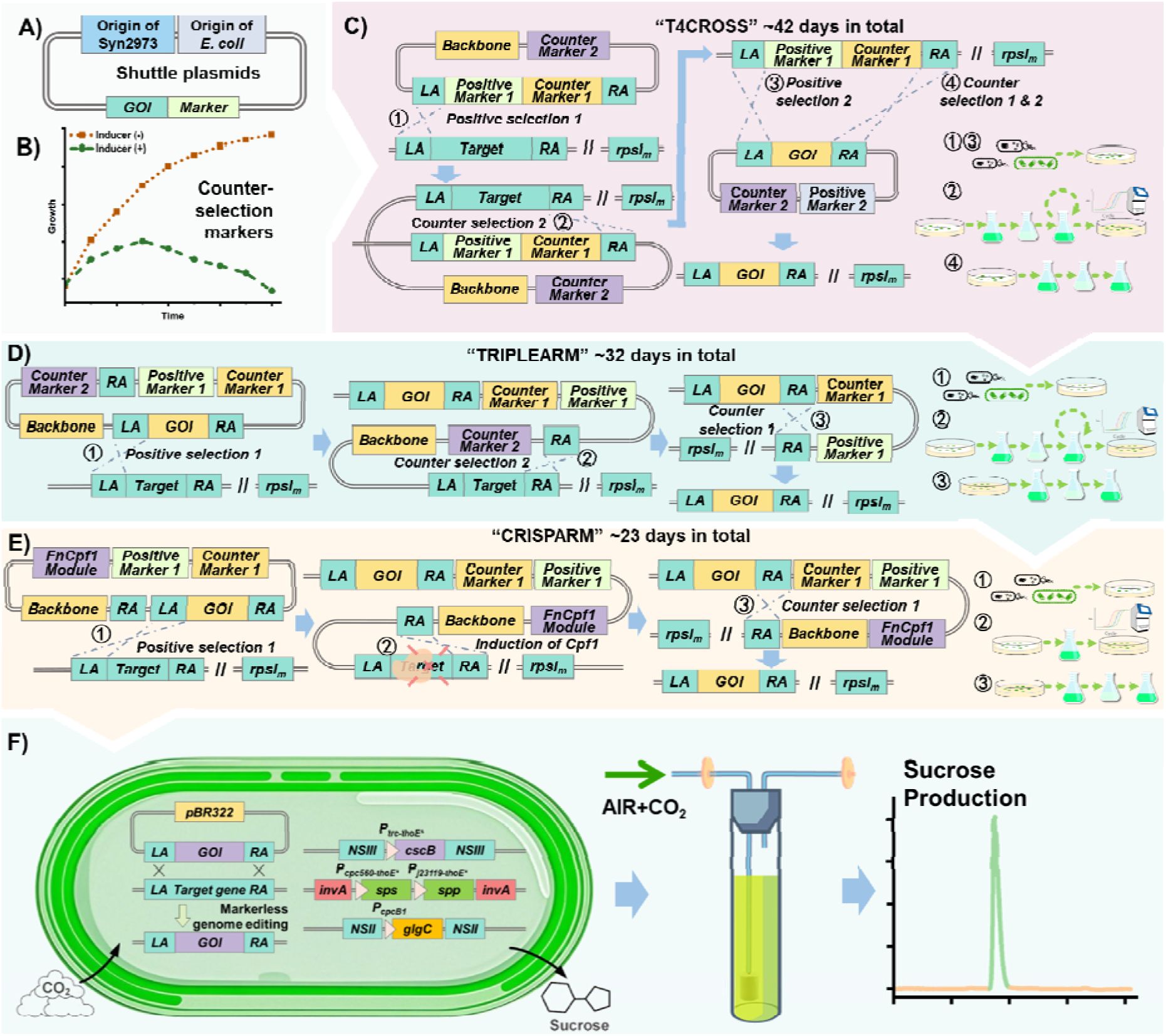
Schematic overview of this study. **A)** Diagram of the shuttle vector structure. **B)** Illustration of the growth effects of the counter-selectable marker before and after induction. **C)** Schematic representation of the recombination strategy and experimental procedure of the “T4CROSS” method. Circled numbers indicate the sequence of the four single crossover events. **D)** Recombination strategy and experimental procedure of the “TRIPLEARM” method. Circled numbers indicate the sequence of the three single crossover events. **E)** Recombination strategy and experimental procedure of the “CRISPARM” method. Circled numbers indicate the sequence of the three single crossover events. **F)** Iterative scarless genome editing using the developed methods to construct a sucrose-producing *Synechococcus elongatus* UTEX 2973 strain, and a schematic diagram of sucrose production in a photobioreactor.

## Methods

### Bacterial strains and growth conditions

The wild-type Syn2973 (hereafter referred to as WT) and engineered strains were cultivated in standard BG11 medium (pH 7.5) (29). For liquid cultures, cells were grown in shaking flasks using an illuminated incubator (HNYC-202T, Honour, Tianjin, China) at 37 °C with agitation at 200 rpm and continuous illumination of ∼500 μmol photons·m^-2^·s^-1^. For solid cultures, plates were incubated at 37 °C under a light intensity of ∼300 μmol photons·m^-2^·s^-1^ in a growth chamber (SPX-250B-G, Boxun, Shanghai, China). *Escherichia coli* strains were cultured under similar temperature and shaking conditions but without light exposure. When appropriate, antibiotics were added to the media at the following final concentrations: for Syn2973, 20 μg·mL^-1^ kanamycin, 20 μg·mL^-1^ chloramphenicol, 50 μg·mL^-1^ spectinomycin, 50 μg·mL^-1^ erythromycin, and/or 50 μg·mL^-1^ streptomycin; for *E. coli*, 100 μg·mL^-1^ ampicillin, 50 μg·mL^-1^ kanamycin, 50 μg·mL^-1^, 100 μg·mL^-1^ spectinomycin, 200 μg·mL^-1^ erythromycin, and/or 50 μg·mL^-1^ streptomycin. When required, 50 μg·mL^-1^streptomycin, 1 mM IPTG (isopropyl β-D-1-thiogalactopyranoside), or 2 mM theophylline was used as an inducer. Optical density was measured at 750 nm (OD_750_) for Syn2973 and at 600 nm (OD_600_) for *E. coli* using a UV-1750 spectrophotometer (Shimadzu, Kyoto, Japan).

### Plasmids and strains construction

*E. coli* DH5α was used for plasmids construction and amplification. All the primers and DNA fragments were chemically synthesized by GENEWIZ Inc. (Suzhou, China). In this study, plasmid construction was mainly carried out using ClonExpress MultiS One Step Cloning Kit (Vazyme Biotech Co., Ltd, Nanjing, China) and Golden Gate assembly technology. The constructed plasmids, their construction methods, and the corresponding primers used were listed in **Table S1**, among which the synthetic DNA sequences used for plasmid construction are listed in **Table S2**. The primers used for PCR or qPCR verification are listed in **Table S3** and some of the key plasmid sequences are listed in **Table S4**. The transformation of Syn2973 was carried out through conjugation according to studies reported previously (30). When testing transformation efficiency, the initial Syn2973 input was controlled at an appropriate level, where volume * OD_750 nm_ = 0.01, 0.1 or 0.5 (approximately equals to 1.45 x 10^6^ , 1.45 x 10^7^ or 7.25 x 10^6^ cells). The strains used in this study were listed in **Table S5**.

### Natural transformation

For natural transformation (31), 1 mL of exponentially growing Syn2973 culture (OD_750_ = 1.0) was collected by centrifugation and resuspended in 50 μL of BG11 medium containing 500 ng of linear DNA. The mixture was then incubated statically at 37°C under a light intensity of 50 μmol photons m^-2^·s^-1^ for 5 hours, with gentle shaking every 2.5 h. Finally, the mixture was evenly spread on sterile filters (0.45 µm pore size) coated on BG11 agar plates containing the appropriate antibiotics and cultured under a light intensity of approximately 200 μmol photons m^-2^·s^-1^.

### Plasmid stability test

Plasmid stability was tested according to previously described methods (32). The strain was centrifuged at 4000 rpm and 4 °C then resuspended in BG11 medium. This process was repeated twice. Subsequently, continuous cultivation was carried out in BG11 medium without antibiotics starting from an initial OD_750_ of 0.1. Each time the OD_750_ of the culture reached 1, the above process was repeated. For each round, strains with an OD_750 nm_ of 1.0 (approximately equals to 1.45 x 10^8^ cells) were diluted by 1000-fold and spread onto antibiotic-free BG11 plates. Fifty single colonies obtained were then subjected to resistance testing.

### β-galactosidase activity measurement

The method for measuring β-galactosidase activity is based on a previous study (33). In simple terms, cells from the exponential growth phase of Syn2973 (volume × OD_750 nm_ =1, approximately equals to 1.45 x 10^8^ cells) are centrifuged and resuspended in 1 mL of Z buffer (containing 60 mM Na_2_HPO_4_, 40 mM NaH_2_PO_4_, 10 mM KCl, 1 mM MgSO_4_, and 40 mM β-mercaptoethanol). Then, 50 μL of 0.1% SDS and 50 μL of chloroform are added to lyse the cells. Subsequently, 200 μL of ortho-nitrophenyl-beta-d-galactopyranoside (ONPG; 4 g·L^-1^) is added to initiate the reaction. The reaction mixture is incubated for a certain period at 30 °C and 750 rpm in a constant temperature shaker for 3 minutes. Finally, 500 μL of 1 M Na_2_CO_3_ is added to stop the reaction. The Miller value is calculated using the OD_420_ of the supernatant (Miller = 1000 × OD_420_ / 3).

### QRT-PCR analysis

The determination of copy numbers and the assessment of genome homozygosity are performed using qRT-PCR (34). After growing for 48 h, the culture (volume*OD_750_ = 2, approximately equals to 2.9 x 10^8^ cells) was collected and the DNA was extracted through a TIANamp Bacteria DNA Kit (TIANGE, Beijing, China). QRT-PCR reactions were performed using PowerUp SYBR Green Master Mix (Thermo Fisher Scientific, MA, USA) in a 10 μL reaction volume, consisting of 5 μL of the mixture, 3 μL of ddH_2_O, 1 μL of template (diluted to 10 ng·μL^-1^ of gDNA), and 0.5 μL of each primer. The reactions were carried out using the StepOnePlus^TM^ Real-Time PCR System (Applied Biosystems, CA, USA). Data analysis was conducted using StepOnePlus Analysis Software (Applied Biosystems, CA, USA). The relative gene copy numbers were calculated using the 2^-ΔΔCT^ method (35), with the 16S rDNA gene (Two copies per genome) serving as an internal reference. Three biological replicates were performed for each condition.

### Sucrose measurement

The concentrate of sucrose was quantified from culture supernatants using high-performance liquid chromatography (HPLC) on an Agilent 1260 instrument equipped with an Aminex HPX-87C column (300 mm × 7.8 mm; Bio-Rad, CA, USA) (36). Ultra-pure water was used as the mobile phase at 80 °C and a flow rate of 0.6 mL·min^-1^ in isocratic mode. Compounds were detected and quantified from 10 μL sample injections using a refractive index detector. Reported metabolite concentrations represent the average of three biological replicates. Standard products were purchased from Aladdin Scientific. The sucrose standard curve is shown in **Fig. S1**.

### Cultivation of Syn2973 using bubble column bioreactor

The high-density cultivation of Syn2973 is conducted in 100 mL flat-bottom glass tubes with a diameter of 30 millimeters, in a volume of 60 milliliters of 5×BG optimized medium (37). Air and carbon dioxide flow rates and ratios (1%, 3% and 5% CO_2_ in air) were controlled using gas mass flow controllers. The total gas flow rate in the tube was maintained at a ratio of 1:1 to the culture volume per minute. Light intensity (500 or 2000 μmol photons m^-2^ s^-1^) and temperature were controlled externally using LEDs and circulating water, respectively.

## Results

### Construction and evaluation of new shuttle plasmids

Syn2973 is not naturally competent due to the incomplete *pilMNOQ* gene cluster involved in pilus biogenesis (33), and its transformation typically relies on *E. coli*-mediated conjugation. Therefore, shuttle plasmids capable of replication in both *E. coli* and Syn2973 are essential for genetic manipulation. In this study, we focused on two endogenous plasmids from Syn2973—designated pANS and pANL, following their nomenclature in the closely related strain Syn7942. The replication origins of pANS and pANL have been well-characterized in Syn7942 (38,39). We constructed shuttle plasmids by combining these origins with the *E. coli*-compatible pBR322 backbone, resulting in four variants: pSES-ori and pSEL-ori (containing the full reported replication regions), and pSES and pSEL (with *Bsm*BI/*Bsa*I restriction sites removed to support Golden Gate Assembly, along with non-essential sequence deletions) (**Fig. 2A**, **Table S4**). As controls, we included pRSF-ori (based on the RSF1010 broad-host-range replicon (24)) and pNSI (a pBR322-derived plasmid containing a single homologous arm to support single crossover integration in Syn2973). To evaluate transformation efficiency, 0.01 or 0.1 OD of Syn2973 cells (∼1.45 or 14.5 × 10^6^ cells) were used for conjugation. As shown in **Fig. 2B** and **Fig. S2**, pSES-ori and pSEL-ori yielded approximately 1,540 and 6 colonies, respectively, at 0.01 OD, while 78,513 (the colonies could not be clearly distinguished used 90 mm plates thus 200 mm plates were used) and 541 colonies were obtained at OD 0.1. Notably, the modified versions (pSES and pSEL) displayed comparable transformation efficiencies, indicating that the modifications did not impair performance. All four pANS/pANL-derived plasmids outperformed pNSI in transformation efficiency. In contrast, pRSF-ori yielded no colonies at OD 0.01 and only 5 colonies at OD 0.1, suggesting poor compatibility. To investigate the cause, we performed qRT-PCR to quantify plasmid copy number relative to the genome. pRSF-ori exhibited significantly lower copy numbers than pSES and pSEL, and even lower than the genomic baseline. Given that the RSF1010-encoded *repF* gene represses the expression of replication proteins RepA and RepC (40), we deleted *repF* to create the pRSF variant. This modification significantly improved transformation efficiency, yielding 1 and 167 colonies at OD 0.01 and 0.1, respectively, along with increased plasmid copy number (**Fig. 2B**). The results indicated that copy number is only partial to the reasons altering the conjugation efficiency of RSF1010-based vector in Syn2973.

**Fig. 2.**
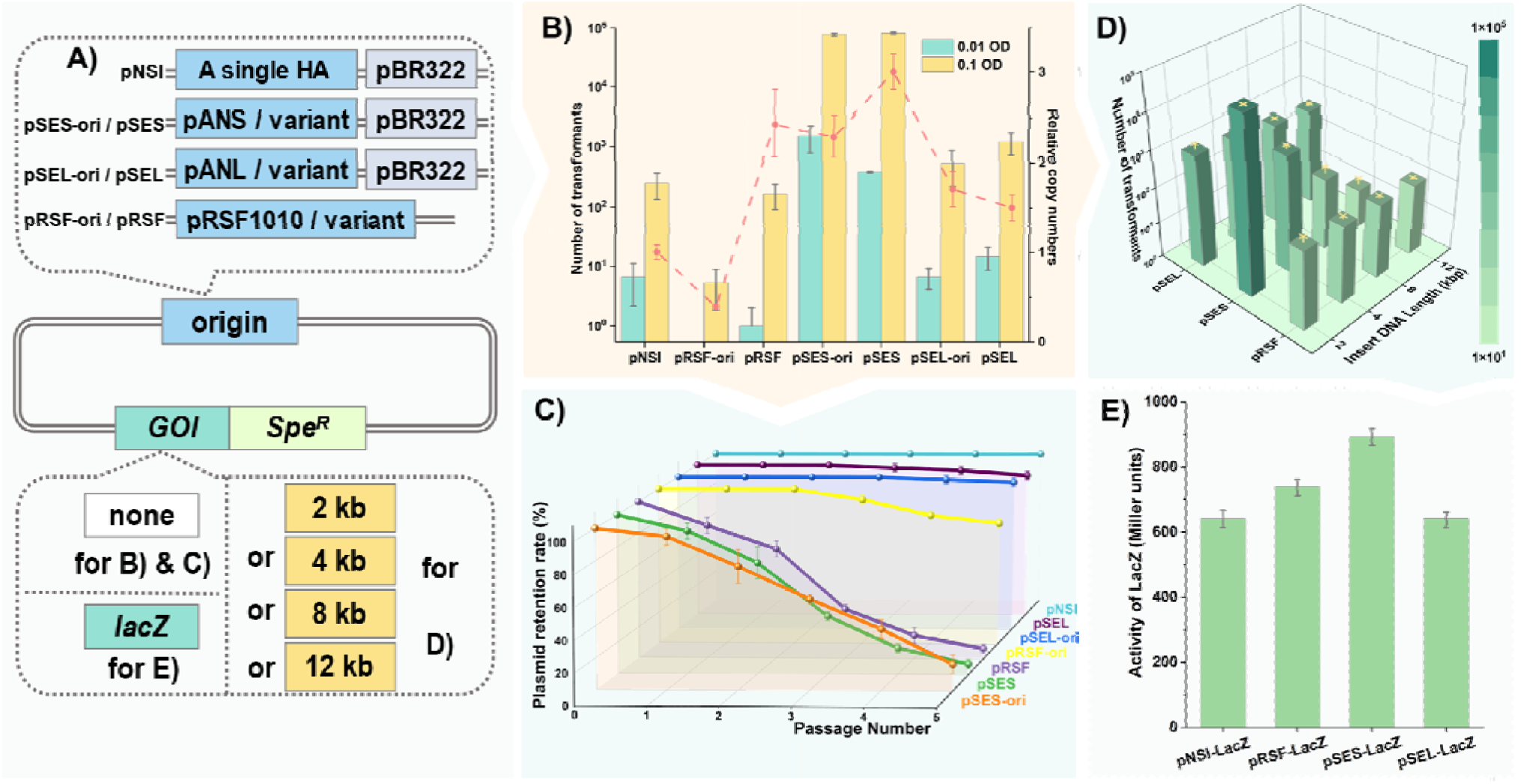
Construction and characterization of shuttle plasmids. **A)** Schematic overview of shuttle plasmid design and testing. The dashed box at the top shows the replication/integration elements being tested, while the lower dashed box indicates the insertion of the gene of interest (GOI) in panels **B–E**. **B)** Transformation efficiency of different synthetic shuttle plasmids at initial OD of 0.01 and 0.1, along with their relative copy numbers compared to the genome in Syn2973. **C)** Relative plasmid retention rates of various synthetic shuttle plasmids during serial passaging without antibiotic pressure. **D)** Transformation efficiencies of pSES, pSEL, and pRSF plasmids carrying gene fragments of varying lengths (2 kb, 4 kb, 8 kb, and 12 kb), with an initial OD of 0.1. **E)** Activity levels of β-galactosidase encoded by pSES, pSEL, and pRSF, and comparison to activity from integration at the NSI locus. Error bars represent standard deviations from three independent replicates.

We next assessed plasmid stability, insert capacity, and gene expression performance. After five passages without antibiotic selection, retention rates of pSEL, pSEL-ori, and pRSF-ori were 93.3%, 96.7%, and 78.3%, respectively, indicating good plasmid stability. In contrast, retention rates for pSES, pSES-ori, and pRSF were only 6.7%, 16.7%, and 6.7%, respectively (**Fig. 2C**). To assess fragment tolerance, we inserted 2 kb, 4 kb, 8 kb, and 12 kb sequences into pSES, pSEL, and pRSF. All three vectors successfully carried up to 12 kb inserts (**Fig. 2D**). However, the transformation efficiency of pSES declined sharply with increasing insert size, whereas pSEL and pRSF maintained relatively stable performance across the tested range. Finally, we evaluated expression strength using a *lacZ* reporter gene under the control of the Ptrc promoter. As expected, plasmids with higher copy numbers (pSES and pRSF) exhibited stronger β-galactosidase activity than pSEL (**Fig. 2E**). Taken together, pSES and pSEL were selected as the primary shuttle plasmids for subsequent genetic manipulations in Syn2973 due to their high efficiency, stability, and performance.

### Development of counter-selection systems and marker-free genetic tools

As previously reported (17) and confirmed in this study (**Fig. 3A** and **S3A**), even with the use of two homologous arms, most transformants (n = 20, three groups) retained a single crossover state after five passages. To overcome this limitation, we tested a range of potential counter-selection markers, including: *i*) *rpsl_12_*, which encodes ribosomal protein S12 (its wild-type form is toxic in the presence of streptomycin when introduced into a streptomycin-resistant background) (26); *ii*) *sepT_2_*, a VapC-like toxin encoding gene known to be lethal to Syn2973 (27); *iii*) *tetA*, which confers tetracycline resistance but increases Ni^2+^ sensitivity (28). The commonly used *sacB* gene in Syn6803, which confers sucrose sensitivity, served as a control (25). The *tetA* and *sacB* cassettes were inserted into the stable plasmid pSEL under the control of the *P_tetA_* and *P_sacB_* promoters, respectively. *sepT_2_* was placed under a theophylline-inducible *P_trc-tho_* switch, resulting in the strains WT-pSEL-TetA, WT-pSEL-SacB, and WT-pSEL-SepT_2_ (**Table S5**). A streptomycin-resistant strain, WT-RPSLm, was generated by mutating the native *rpsl_12_* gene (**Fig. S4**). The wild-type *rpsl_12_* gene from Syn6803 was then introduced into WT-RPSLm, generating the strain WTR-pSEL-68Rpsl. As shown in **Fig. 3B** and **3C**, neither Ni^2+^-induced expression of *tetA* nor sucrose-induced *sacB* expression was lethal to Syn2973. In contrast, theophylline-induced expression of *sepT_2_* significantly inhibited the growth of WT-pSEL-SepT_2_ after 48 h (**Fig. 3D**), and streptomycin addition suppressed the growth of WTR-pSEL-68Rpsl (**Fig. 3E**). Both systems also successfully eliminated plasmids during two passages post-induction (**Fig. 3F, S3B**), providing new applications in plasmid cure. We then employed these counter-selection markers in a two-step single-crossover recombination strategy to knock out the *pilMNOQ* gene cluster (**Fig. 3G**). Following a single passage with theophylline induction, ∼93.3% of colonies (n = 20, three groups) exhibited successful double crossovers, generating a strain WT-PIL-KM (**Fig. 3H** and **3I**, with PCR validation shown in **Fig. S3C**).

**Fig. 3.**
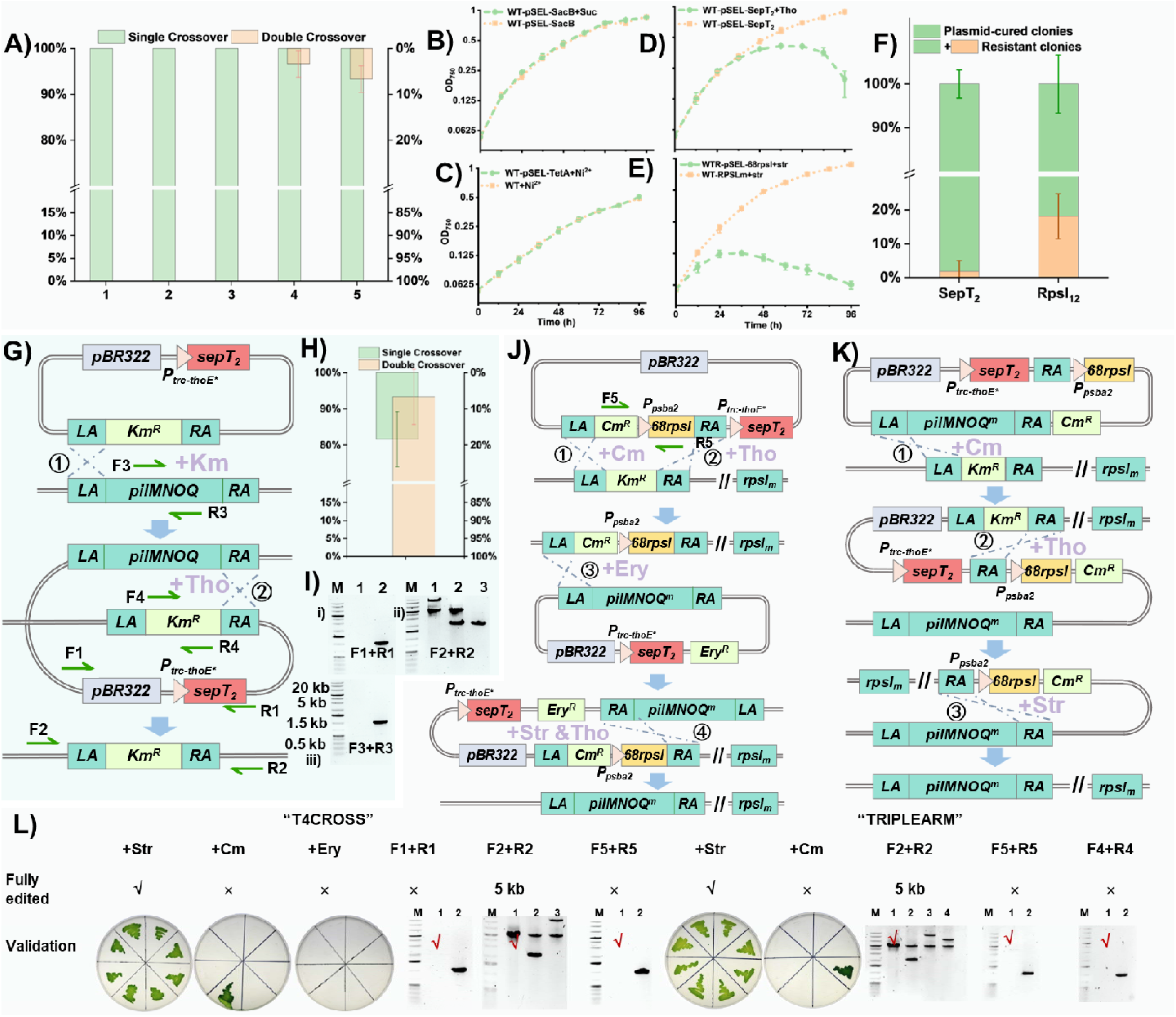
Identification and application of negative selection markers. **A)** Proportion of Syn2973 transformants undergoing single-crossover and double-crossover recombination across five consecutive passages. **B–E)** Effects of different negative selection systems on Syn2973 growth, with or without inducers: B) SacB system (50 g·L^-1^ sucrose), C) TetA system (2.5 μM Ni^2+^), D) SepT_2_ system (2 mM theophylline), and E) RpsL_12_ system (50 μg·mL^-1^ streptomycin). **F)** Application of SepT_2_ and RpsL_12_ for plasmid curing. The bar graph shows the proportions of plasmid-free versus plasmid-retaining colonies (20 colonies per group, 3 groups total) after inducer treatment. Error bars represent standard deviations from three independent groups. **G)** Schematic diagram of gene knockout using negative selection markers. Validation primer locations are also indicated. **H)** Proportions of single- and double-crossover events in transformants (20 colonies per group, 3 groups total) after induction of negative selection markers. Error bars represent standard deviations from three independent groups. I) Representative PCR gel electrophoresis results using three primer sets corresponding to panel G. Lane annotations: F1–R1: Lane 1, no single crossover; Lane 2, presence of single crossover; F2–R2: Lane 1, coexistence of wild-type and single-crossover; Lane 2, coexistence of wild-type and double-crossover; Lane 3, double-crossover homozygote; F3–R3: Lane 1, absence of wild-type genome; Lane 2, presence of wild-type genome. **J)** Schematic diagram of replacing the Km^R^ cassette with *pilMNOQ^m^* in the WTR-PIL-KM strain using the T4CROSS method. Validation primers F5–R5 are indicated. **K)** Schematic diagram of the same gene replacement using the TRIPLEARM method. **L)** Phenotypic or molecular validation results of correctly modified strains under corresponding conditions for both T4CROSS and TRIPLEARM strategies.

Next, we aimed to construct marker-free genetic tools based on the validated counter-selection markers. As a proof of concept, we replaced the native *pilMNOQ* cluster with a mutant version (*pilMNOQ^m^*) in the WT-PIL-KM background (the original *rpsl_12_* gene was first mutated to obtain WTR-PIL-KM). The first strategy, named “T4CROSS” (**Fig. 3J**), used two plasmids and four rounds of single-crossover recombination. In the first step, a cassette containing both a positive selection marker and the *rpsl_12_* counter-selection marker replaced the kanamycin-resistance cassette in WTR-PIL-KM. Another counter-selection marker *sepT_2_* was introduced to facilitate the second single crossover at the same time. The resulting strain was chloramphenicol-resistant and streptomycin-sensitive. In the second round, a plasmid carrying *pilMNOQ^m^*, an erythromycin resistance marker, and *sepT_2_* was introduced. After selecting transformants on erythromycin, marker-free replacement of the target gene was achieved via combined induction of streptomycin and theophylline, activating both counter-selection systems. The second strategy, “TRIPLEARM” (**Fig. 3K**), used a single plasmid and three single-crossover steps. Here, the cassette containing a positive selection marker and *rpsl_12_* was flanked by two identical homologous arms, enabling excision of the marker in the final step. As with T4CROSS, *sepT_2_* was used to facilitate the intermediate recombination event. As shown in **Fig. 3L** (with PCR validation in **Fig. S3D**, **S3E**), both approaches successfully enabled marker-free genome manipulation. Notably, the resulting strain, WTR-pilNm, regained natural competence and could be transformed directly using DNA fragments (**Fig. S3F**).

### Development of marker-free gene knock-in tool based on modified CRISPR/Cpf1

Genome editing using CRISPR/Cpf1 avoids the cytotoxicity associated with Cas9 (41) by directly cleaving target sequences, thereby forcing cells to survive through either homology-directed repair (HDR) or non-homologous end joining (NHEJ). A CRISPR/Cpf1-based system has previously been reported in Syn2973 (23). Initially, we constructed an all-in-one shuttle plasmid based on pRSF-ori, harboring *cpf1*, crRNA, and repair templates, targeting the *pilMNOQ* cluster (**Fig. 4A**). Three promoters of varying strength—*P_lac_, P_trc_*, and *P_tho_* (specifically P_trc-thoE*_)—were used to drive *cpf1* expression. However, using our standard transformation conditions (0.5 OD, ∼7.25 x 10^7^ cells), no transformants were obtained, suggesting low editing efficiency. Switching the backbone from pRSF-ori to our modified vectors (pSES, pSEL, and pRSF, **Fig. 4B**) did not yield any edited colonies either (**Fig. 4C**). To overcome this limitation, we explored a stepwise transformation strategy by separating *cpf1* and crRNA expression into two different plasmids (**Fig. 4D**). All six possible plasmid combinations using pSES, pSEL, and pRSF were evaluated. As shown in **Fig. 4E**, the stepwise strategy significantly improved transformation efficiency, with pSES-based plasmids generating the highest number of colonies. Notably, over 100 transformants were obtained with combinations of pSES (carrying crRNA) and pSEL or pRSF (carrying *cpf1*). However, most transformants failed to survive subsequent subculturing (**Fig. 4E**), and only a small fraction of the survivors was confirmed to be successfully edited (PCR verification in **Fig. S5A**). We hypothesized that the mismatch between the high DNA cleavage activity of Cpf1 and the low HDR efficiency in Syn2973 was a key bottleneck. Unlike *E. coli* or Syn6803, Syn2973 lacks key recombination-related genes such as *lexA* and *recD*, which may also contribute to the reduced resistance to DNA damage observed in its close relative Syn7942 (42).

**Fig. 4.**
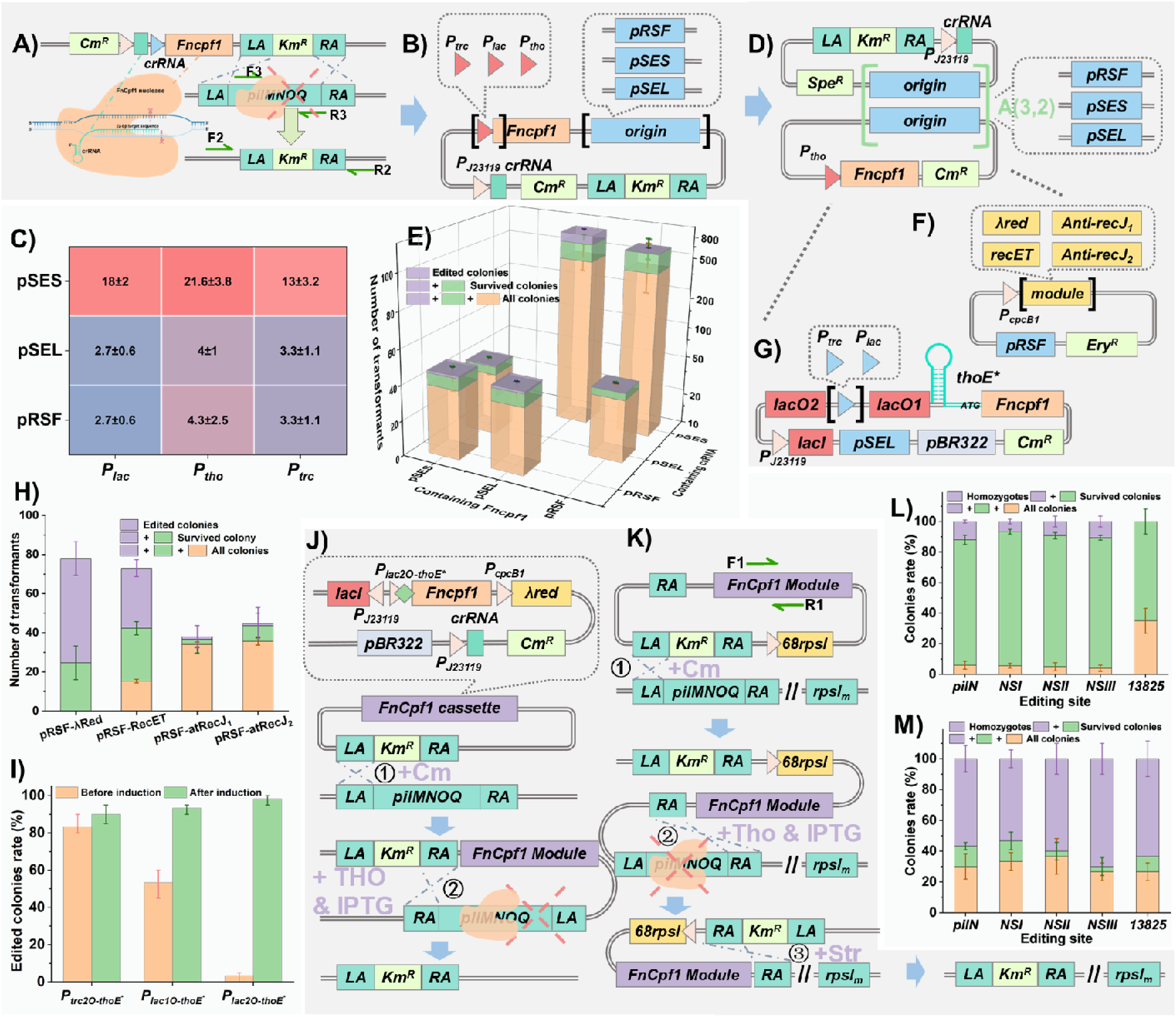
Testing and optimization of the CRISPR/Cpf1 system. Error bars representstandard deviations from three independent experiments. **A)** Schematic diagram showing the evaluation of CRISPR/Cpf1 efficiency via knockout of the *pilMNOQ* operon. **B)** Diagram of shuttle plasmids carrying the crRNA cassette, homologous insert fragments, and the cpf1 gene driven by different promoters. **C)** Transformation efficiency of plasmids described in panel B using an initial culture with OD= 0.5. **D)** Schematic of a two-plasmid CRISPR system in which the functional modules are separated. **E)** Number of transformants, survival rate, and editing outcomes after sequential introduction of the FnCpf1 expression plasmid followed by the crRNA plasmid. The transformation efficiency of the first-step introduction of the plasmid carrying **Cpf1** is shown in **Fig. S3G**. **F)** Schematic of plasmid constructs based on panel D, incorporating either a recombination-enhancing module or a plasmid designed to repress endogenous recJ expression. **G)** Modified system from panel D using a more tightly regulated inducible promoter to control FnCpf1 expression. **H)** Number of transformants, survival rate, and editing success after introduction of homologous recombination-enhancing elements (20 colonies per group, 3 groups total). **I)** Editing efficiency before and after induction of FnCpf1 expression under different inducible promoters (20 colonies per group, 3 groups total). **J)** Schematic of a one-plasmid CRISPR strategy in which the CRISPR and recombination modules are placed on a recombination plasmid, which integrates into the target site via a single crossover and is subsequently excised along with the target gene in a second single crossover event. **K)** Schematic illustration of the “CRISPARM” strategy. **L)** Editing efficiency of the strategy shown in panel J at different genomic loci (20 colonies per group, 3 groups total). **M)** Editing efficiency of the “CRISPARM” method across various gene loci.

To balance DNA cleavage with recombination, we utilized a two-pronged approach: *i*) Enhancing recombination via introduction of additional systems such as λ-red (1) and RecET (43), and inhibition of endogenous *recJ*, which was previously shown to increase HDR efficiency in Syn2973 (22) (**Fig. 4F**); and *ii*) Implementing tighter control of *cpf1* expression using a tandem inducible system responsive to IPTG and theophylline (**Fig. 4G**). As shown in **Fig. 4H**, both λ-Red and RecET improved editing efficiency, with λ-Red being more effective (PCR results in **Fig. S5B**).

Moreover, combining λ-Red with *cpf1* under the dual-inducible promoter *P_lac2O-thoE*_* allowed precise control: no editing occurred without induction, whereas clear editing was observed following induction with 2 mM theophylline and 1 mM IPTG (**Fig. 4I, S6A**). However, even with this strategy, homozygous edited colonies were not obtained (**Fig. S5C**), likely due to instability of the pSES-based plasmid or insufficient recombination efficiency. To further improve outcomes, we employed an integrative plasmid approach (**Fig. 4J**), wherein the CRISPR/Cpf1 cassette was integrated into the genome via an initial single crossover. Induction of CRISPR/Cpf1 then promoted a second crossover, achieving marker-free editing by excising the target gene and editing cassette itself together. This approach was validated by targeting *pilMNOQ*, three neutral sites (NSI, NSII, NSIII), and gene *M744_13825*, which affects growth upon deletion (44) (**Fig .4L S6B**). Homozygous transformants were obtained for all non-essential targets (∼10% efficiency; **Fig. S6D**). However, for *M744_13825*, no homozygotes were recovered, potentially due to the loss of Cpf1 function before complete homozygosity was achieved. To further optimize this approach, we combined the enhanced CRISPR/Cpf1 tool with our previously established “TRIPLEARM” method, creating a hybrid strategy named “CRISPARM” Here (**Fig. 4K**), both the Cpf1 cassette and the Rpsl_12_ counter-selection marker were flanked by identical homologous arms. Dual induction with IPTG and theophylline, combined with positive selection for resistance, enabled the isolation of edited strains. The Cpf1 cassette was subsequently removed through streptomycin-based counter-selection. Using an 8 kb GOI to evaluate the system, we achieved homozygous editing in all target loci with >50% efficiency (**Fig. 4M** and **S6C, E**). Approximately 30% of transformants exhibited unedited outcomes, mainly due to survival failure during the sub-induction process. These were likely caused by unintended single crossover events due to the presence of three homologous arms, which is discussed further in the subsequent sections.

### Standardization and comparison of scarless operation methods

To facilitate broader application of the genome editing tools developed above, we standardized the entire manipulation workflow and compared the editing efficiency of each method. As a model case, we introduced the *cscB* cassette—encoding sucrose permease—into the WT-RPSLm strain to enable sucrose secretion (**Fig. 5A**).

**Fig. 5.**
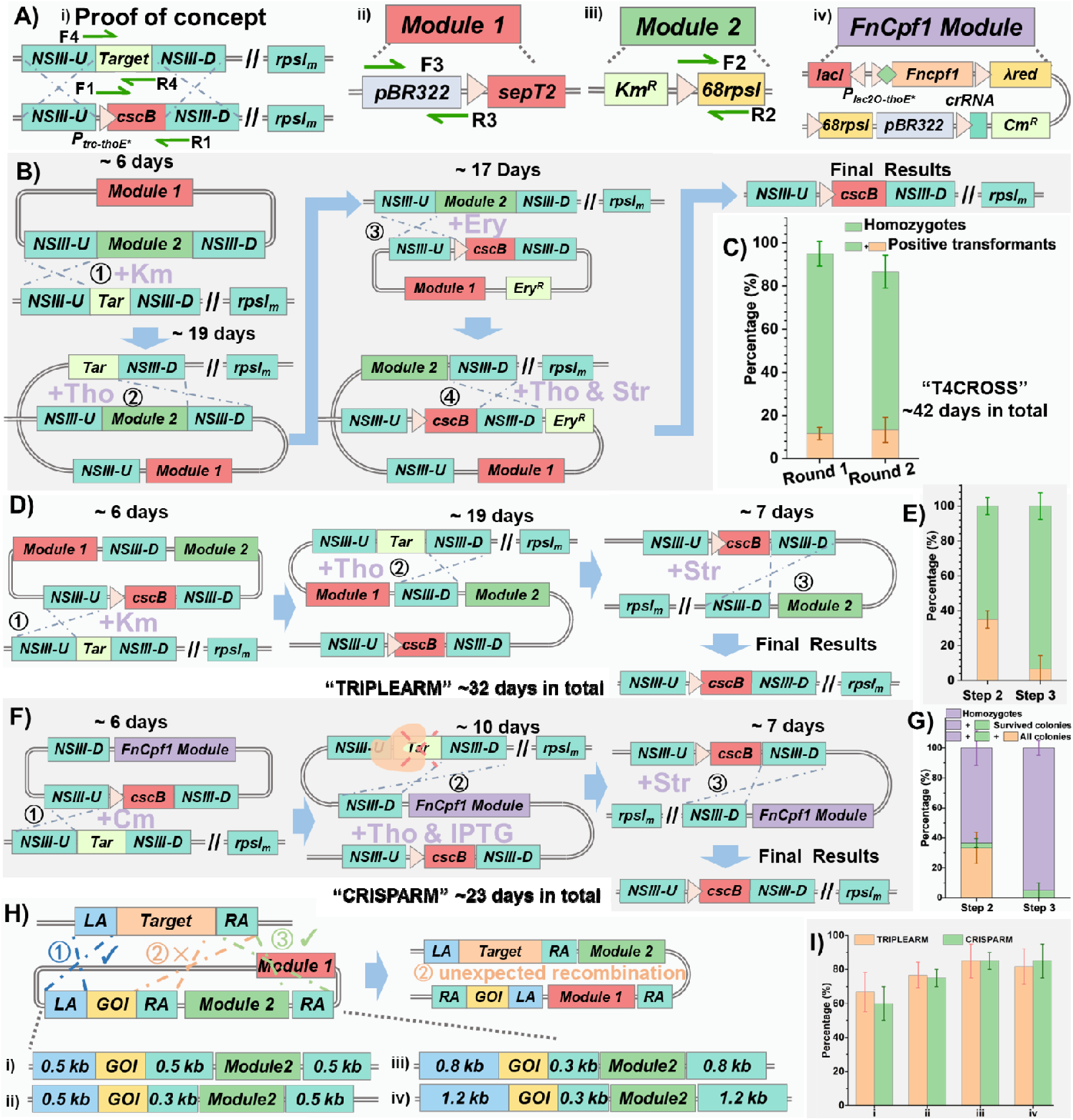
Integration of the *cscB* expression cassette into Syn2973 using three marker-free genome editing strategies for sucrose secretion. Error bars represent standard deviations from three independent groups. A) Schematic overview of introducing *cscB* into Syn2973 to construct the WTR-B3 strain, along with simplified diagram legends used in other subpanels. **B)** Construction of WTR-B3 using the T4CROSS method. **C)** Proportions of positive colonies and homozygotes obtained in the first and second rounds of transformation using the T4CROSS strategy (20 colonies per group, 3 groups total). **D)** Construction of WTR-B3 using the TRIPLEARM method. **E)** Proportions of positive colonies and homozygotes in the second and third recombination steps using TRIPLEARM (20 colonies per group, 3 groups total). **F)** Construction of WTR-B3 using the CRISPARM method. **G)** Survival, positivity, and homozygosity rates in the second and third recombination steps using CRISPARM (20 colonies per group, 3 groups total). **H)** Schematic representation of possible homologous recombination pathways during the first step of TRIPLEARM and CRISPARM, along with the optimized strategy to improve recombination specificity. **I)** Effect of different homologous arm lengths on the survival rate of transformants after induction.

For the first method, “T4CROSS” (**Fig. 5B**), a plasmid designed to replace the NSIII site with both a positive and a counter-selection cassette was first introduced. Following approximately six days of selection, individual colonies were transferred to shake flasks for expansion. Inducers and antibiotics were then added to trigger the second single-crossover recombination event. After approximately three passages, homozygous mutants were verified by qRT-PCR. This step took around 19 days in total, with 83.3% of the colonies identified as homozygous (**Fig. 5C**). Subsequently, a second plasmid carrying the *cscB* cassette was introduced into the verified homozygotes. After two further rounds of single-crossover recombination, induced by streptomycin and theophylline, the final sucrose-producing strain WTR-B3—free of any selection markers—was obtained. This second editing phase required approximately 17 days, yielding a final positive colony rate of 73.3% (n = 20, three groups) (**Fig. 5C**). Overall, the T4CROSS method required about 42 days to complete the full editing cycle.

For the “TRIPLEARM” (**Fig. 5D**) and “CRISPARM” (**Fig. 5F**) methods, a single plasmid targeting the NSIII site was constructed for each approach. Selection for the first integration event was performed using kanamycin or chloramphenicol, respectively, and both methods required approximately 6 days for this step. In the TRIPLEARM workflow, the second single-crossover recombination was induced using theophylline after colonies growing to mid-log phase in shaking flasks, followed by three serial passages to enrich for homozygous mutants, which were again confirmed by qRT-PCR. In contrast, for the CRISPARM method, IPTG and theophylline were added directly after transferring the selected colonies into flasks, enabling faster induction. The TRIPLEARM approach required approximately 19 days, with a final homozygous positive rate of 65.0% (**Fig. 5E**). The CRISPARM method took only 10 days, yielding a positive rate of 62.7% (**Fig. 5G**). In both approaches, streptomycin was used in the final step to trigger a third single-crossover recombination event, leading to the removal of editing cassettes and recovery of marker-free strains. The entire workflows required approximately 32 days for TRIPLEARM and 23 days for CRISPARM. It is worth noting that both TRIPLEARM and CRISPARM require the simultaneous presence of three homologous arms, which introduces the potential for unintended recombination events due to three possible crossover sites (**Fig. 5H**). To optimize recombination directionality and efficiency, we systematically adjusted the lengths of the three homologous arms. The best performance was observed when the two terminal arms were extended to 800 bp and the central arm was shortened to 300 bp. Under these conditions, the editing efficiency reached approximately 85.0% (**Fig. 5I**), demonstrating that fine-tuning arm lengths can effectively improve recombination outcomes.

### Iterative marker-free genetic manipulations enabled high production of sucrose in Syn2973

As described above, a sucrose-secreting strain, WTR-B3, was successfully constructed. When 150 mM NaCl was added to the medium, sucrose production was significantly enhanced in Syn2973, reaching 662.2 mg·L^-1^·OD^-1^, compared to only 125.3 mg·L^-1^·OD^-1^ under salt-free conditions (**Fig. 6B, 6C**). However, this increase in production came at the cost of inhibited cell growth. To improve sucrose production without salt stress, we applied the developed “CRISPARM” method to perform iterative marker-free genetic modifications. The key rate-limiting steps in the cyanobacterial sucrose synthesis pathway are catalyzed by sucrose-phosphate synthase (Sps) and sucrose-phosphate phosphatase (Spp). These enzymes are strongly upregulated during salt induction and are considered primary targets for enhancing sucrose biosynthesis. To bypass the requirement for NaCl, the *sps* and *spp* genes from Syn6803 were overexpressed in WTR-B3 by integrating their expression cassettes into the NSI site, generating strain WTR-BSP13. As a result, sucrose production increased significantly even in the absence of salt, while cell growth was also restored (**Fig. 6B, 6C**). The sucrose yield reached 610.5 mg·L^-1^·OD^-1^, comparable to that of WTR-B3 under NaCl induction, suggesting that overexpression of *sps* and *spp* can mimic the effect of salt stress on sucrose production. Next, to further enhance sucrose accumulation, we targeted *invA*, a gene encoding invertase responsible for hydrolyzing sucrose into glucose and fructose. The *sps* and *spp* cassette was integrated directly into the *invA* locus, generating WTR-BSPi3. This strain exhibited an additional ∼40% increase in sucrose production, albeit with a slight reduction in growth (**Fig. 6B, 6C**).

**Fig. 6.**
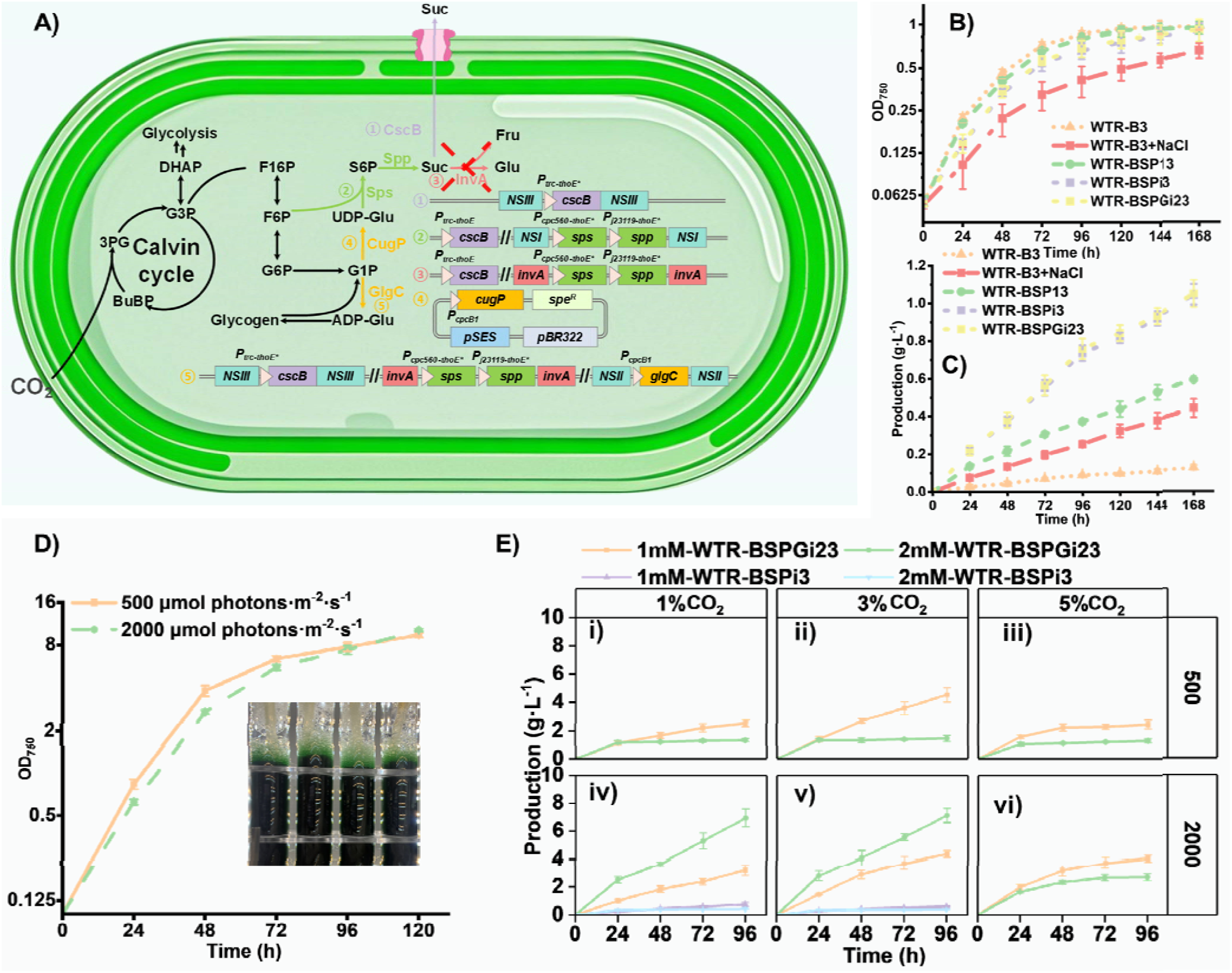
Sucrose production through iterative metabolic engineering of Syn2973. Error bars represent standard deviations from three independent replicates. **A)** Schematic diagram of the sucrose synthesis and degradation pathways in Syn2973, along with the sequential engineering steps applied in this study. **B)** Growth curves of sucrose-producing strains WTR-B3, WTR-BSP13, WTR-BSPi3, WTR-BSPGi23, and WTR-B3 cultured in the presence of 150 mM NaCl. **C)** Sucrose production of the strains listed in panel B under 150 mM NaCl stress. **D)** Growth curve of WTR-BSPGi23 cultivated in a bubble column photobioreactor with 3% CO_2_ under different light intensities. **E)** Sucrose yields of WTR-BSPi3 and WTR-BSPGi23 under varying conditions, including 1%, 3%, or 5% CO_2_ concentrations, theophylline induction at 1 mM or 2 mM, and light intensities of 500 or 2000 μmol photons·m^-2^·s^-1^.

To further boost sucrose synthesis, we explored increasing the supply of its precursors—fructose-6-phosphate (F6P) and UDP-glucose (UDP-Glc). Given F6P’s central role in carbon fixation and glycolysis, it was unlikely to be a limiting substrate. Therefore, we focused on enhancing UDP-Glc synthesis by overexpressing *cugP*, a non-GalU-type UDP-glucose pyrophosphorylase from Syn6803, in WTR-BSPi3. We attempted chromosomal integration of *cugP* at the NSII site, but no positive transformants were obtained across three independent trials. Alternatively, we used the high-efficiency free-shuttle plasmid pSES to introduce the *cugP* cassette. Although transformants were initially obtained, they failed to survive after induction in shake flask cultivation. Interestingly, despite its apparent contradiction, previous studies have reported that overexpression of *glgC*—encoding ADP-glucose pyrophosphorylase, a key enzyme in glycogen biosynthesis—can enhance sucrose production(45). It is hypothesized that increased glycogen synthesis diverts more carbon flux from glycolysis toward glucose-1-phosphate (G1P), thereby promoting UDP-Glc generation. We thus overexpressed *glgC* from Syn6803 in WTR-BSPi3 by targeting the NSII site, resulting in the strain WTR-BSPGi23. Under standard shake flask conditions, *glgC* overexpression did not significantly affect either growth or sucrose yield (**Fig. 6B, 6C**).

To further evaluate performance under more physiologically relevant conditions, we conducted bubble-column photobioreactor cultivation under various combinations of environmental factors. Specifically, we tested both WTR-BSPi3 and WTR-BSPGi23 under three CO_2_ concentrations (1%, 3%, and 5%), two light intensities (500 and 2000 μmol photons·m^-2^·s^-1^), and two concentrations of theophylline (1 mM and 2 mM) as inducers. The results showed that WTR-BSPi3 produced detectable levels of sucrose only under 1% and 3% CO_2_ at high light intensity (2000 μmol photons·m^-2^·s^-1^), but its yield was lower than that observed in shake flasks (**Fig. 6E**). In contrast, WTR-BSPGi23, overexpressing *glgC*, exhibited a significant advantage under aerated conditions. Under optimized conditions—3% CO_2_, 2000 μmol photons·m^-2^·s^-1^ light, and 2 mM theophylline—this strain reached a sucrose titer of 7.12 g·L^-1^ after 4 days, corresponding to an average productivity of 1.79 g·L^-1^·day^-1^.

## Discussion

The lack of efficient marker-free genetic manipulation tools remains a major bottleneck for performing complex synthetic biology modifications in Syn2973. In our previous work, we established a suite of regulatory elements tailored for Syn2973, including constitutive and inducible promoters, as well as gene regulation systems (33). Gene editing and gene silencing tools have also been developed in recent years (21,23). However, we find these tools are still insufficient for high-efficiency, iterative genome engineering due to three key limitations: (i) the absence of stable high-copy-number shuttle plasmids, which necessitates chromosomal integration of exogenous genes in most cases; (ii) a strong bias toward single crossover events over double crossovers, even when two homologous arms are provided, complicating the identification of correctly modified strains; and (iii) the lack of efficient marker-free editing strategies, limiting the number of sequential genetic modifications that can be performed.

Among currently available tools, RSF1010 is the only known broad-host-range replicon capable of stable maintenance in several cyanobacteria, especially Syn6803 and *Nostoc sp.* PCC 7120 (46,47). Although previous studies have employed RSF1010 for genetic manipulation in Syn2973, our findings reveal its transformation efficiency in this strain is low, requiring extensive screening to identify positive clones. We speculate that this inefficiency stems from its low plasmid copy number. Deletion of the *repF* gene partially alleviated this issue but did not fully resolve the instability, suggesting that RSF1010’s replication module does not function optimally in Syn2973. To address this, we developed two novel free-replicating plasmids, pSES and pSEL, by fusing endogenous replication elements from Syn2973 with *E. coli* replicons. These plasmids exhibited satisfactory transformation efficiency and stable maintenance. However, the presence of homologous endogenous plasmids may compete with the introduced plasmids, reducing their stability. In future work, eliminating these endogenous plasmids—if non-essential—could enhance plasmid maintenance, particularly for pSES. Furthermore, although pSEL exhibited reliable replication, its large replication module complicates plasmid construction, suggesting future efforts should focus on streamlining the replication region.

Another major challenge is Syn2973’s preference for single crossover recombination, which compromises the genetic stability of engineered strains and necessitates laborious screening to identify double crossover events. Although modifying the host genome to overcome this bias remains difficult, we developed two counter-selection marker systems that enable a two-step single crossover strategy to achieve functionally equivalent double crossovers. Despite increasing construction time, this approach is far more efficient than blind screening. Previous studies by Chen et al. introduced counter-selection systems including *SYNPCC7002_G0085* from *Synechococcus sp.* PCC 7002 and *mazF* from *E. coli* (17). Our work complements these systems by expanding the available toolkit. Through comparative analysis of recombination systems across cyanobacteria, we also identified the absence of key recombinases in Syn2973 as a possible contributor to its recombination inefficiency. While introducing exogenous recombinases moderately improved recombination outcomes, the improvement was limited. In future studies, a systems biology approach could be employed to dissect the genetic basis of recombination bias in Syn2973, allowing for rational complementation or rewiring of its recombination machinery.

To date, very few studies have reported sequential, marker-free genome editing in Syn2973. To our knowledge, only Ungerer et al. have used CRISPR/Cpf1 to simultaneously mutate three genes (23). Building on our newly developed free-replicating plasmids and negative selection systems, we established three distinct strategies for marker-free genetic manipulation: *i*) T4CROSS, which uses two plasmids and two transformation steps, enabling both point mutations and gene insertions with high precision. *ii*) TRIPLEARM, a single-step transformation approach ideal for gene insertions. However, it is not suitable for point mutations or fine-tuned sequence changes. Additionally, the use of three homologous arms increases the risk of incorrect recombination, which can be mitigated by optimizing homologous arm lengths. *iii*) CRISPRARM, the most efficient and rapid approach, but with some limitations. Its large plasmid size presents cloning challenges, and as it is an integrative system, the post-editing removal of the editing plasmid is more complex than that of free-replicating systems. Future directions include the use of smaller Cas proteins, such as IscB(48), and further optimization of free-replicating plasmids to combine efficiency with ease of use. Collectively, the systems developed in this study not only address several key challenges in Syn2973 engineering but also provide a scalable foundation for future multi-gene and pathway-level genome engineering in this promising photosynthetic chassis.

In this study, a marker-free sucrose-producing strain was constructed using the methods described above. Following fermentation optimization, a final sucrose titer of 7.12 g·L^-^¹ was achieved within 4 days, approaching the current highest reported levels of 8 g·L^-^¹ (4 days, salt-induced) (14) and 8.7 g·L^-^¹ (21 days) (49). These results demonstrate the effectiveness of the proposed strategies for constructing Syn2973-based cell factories. In the future, further investigation into the interplay between sucrose and glycogen biosynthesis could enable more precise carbon flux redistribution toward sucrose production, while culture condition optimization and increased cell density could further enhance sucrose yields.

## Data availability statement

All the data have been included in the supplementary file.

## SUPPLEMENTARY DATA

Supplementary Data are available online.

