## Supplemental Tables and figures for "A Versatile Marker-Free Genome Engineering Platform to Overcome Homologous Recombination Bias in Microbes: A Case Study in *Synechococcus elongatus* UTEX 2973"

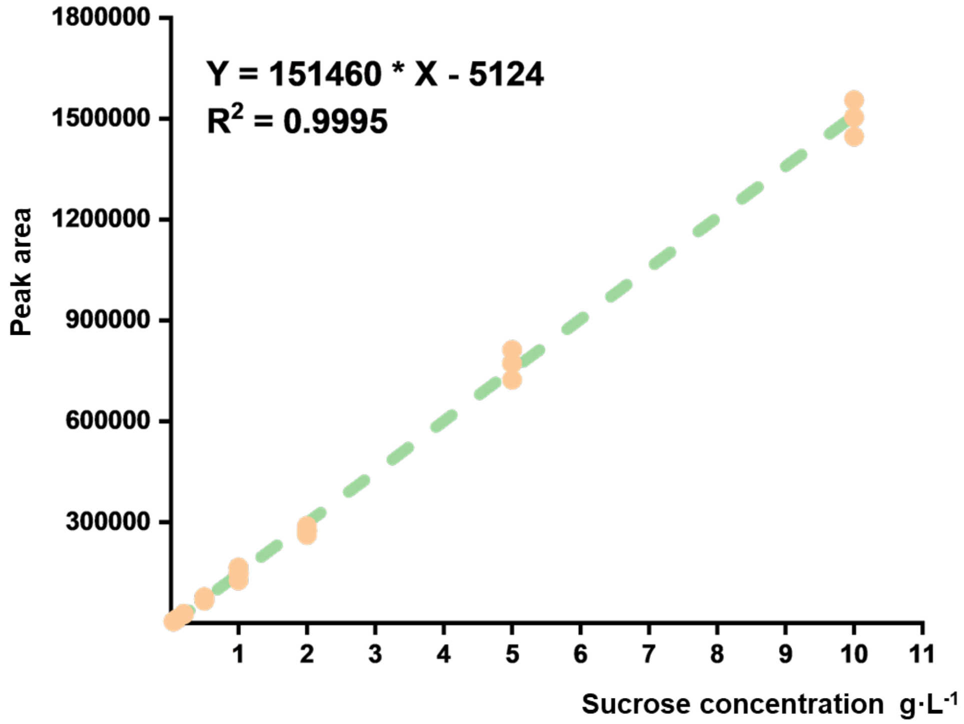

**Fig. S1.** Standard calibration curve showing the relationship between sucrose concentration and peak area as measured by HPLC with refractive index detection (RID).

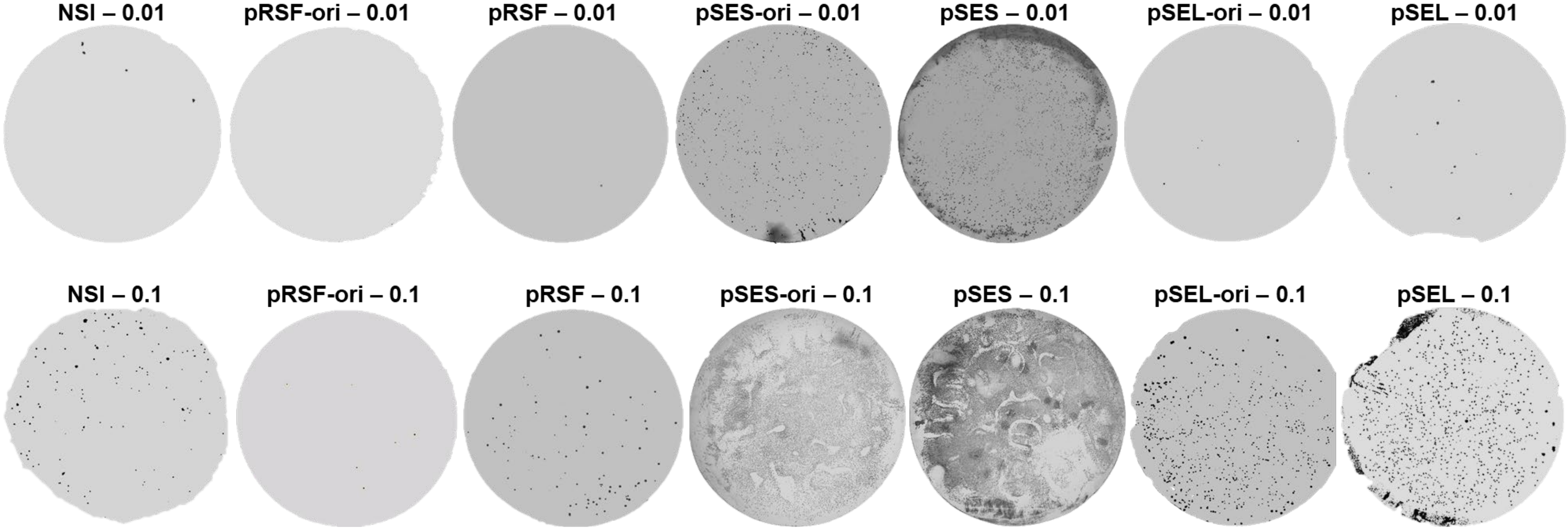

**Fig. S2.** Representative plate images showing transformant growth for NSI, pRSF-ori, pRSF, pSES-ori, pSES, pSEL-ori, and pSEL at initial DNA input amounts of 0.01 and 0.1.

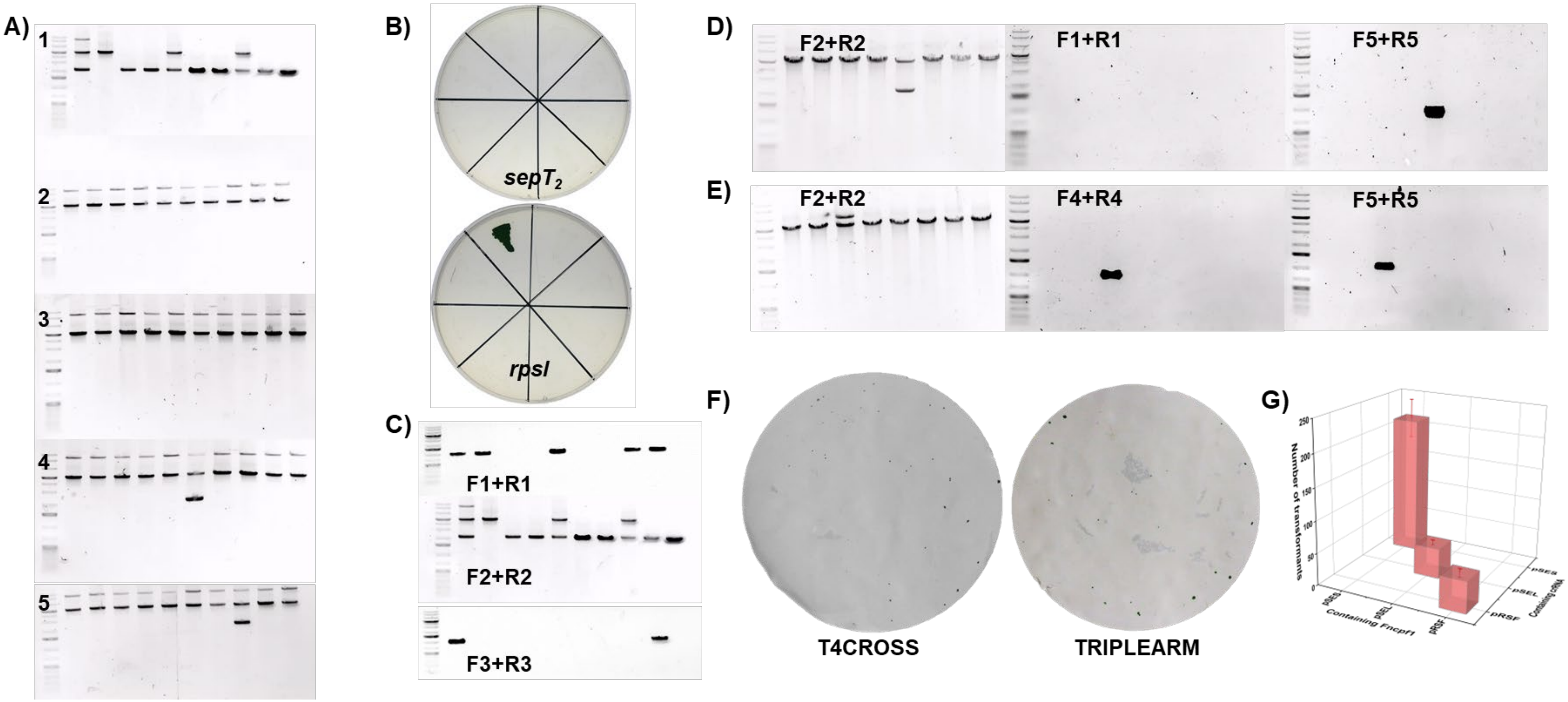

**Fig. S3. A)** PCR verification of single colonies isolated from each generation shown in Fig. 3A using primers F2 and R2 from Fig. 3. **B)** Growth of single colonies on BG11 plates supplemented with kanamycin after plasmid curing via *sepT₂* and *rpsl* selection. **C)** PCR verification of selected single colonies from Fig. 3H using primers F1-R1, F2-R2, and F3-R3 from Fig. 3. **D)** PCR verification of WTR-pilNm strains obtained using the "T4CROSS" strategy in Fig. 3L, with primers F2R2, F1R1, and F5R5 from Fig. 3. **E)** PCR verification of WTR-pilNm strains obtained using the "TRIPLEARM" strategy in Fig. 3L, with primers F2-R2, F4-R4, and F5-R5 from Fig. 3. **F)** Natural transformation results of WTR-pilNm strains obtained by the "T4CROSS" and "TRIPLEARM" strategies. **G)** Number of transformants in the first step of FnCpf1 introduction using plasmids pSES, pSEL, and pRSF, as shown in Fig. 4E.

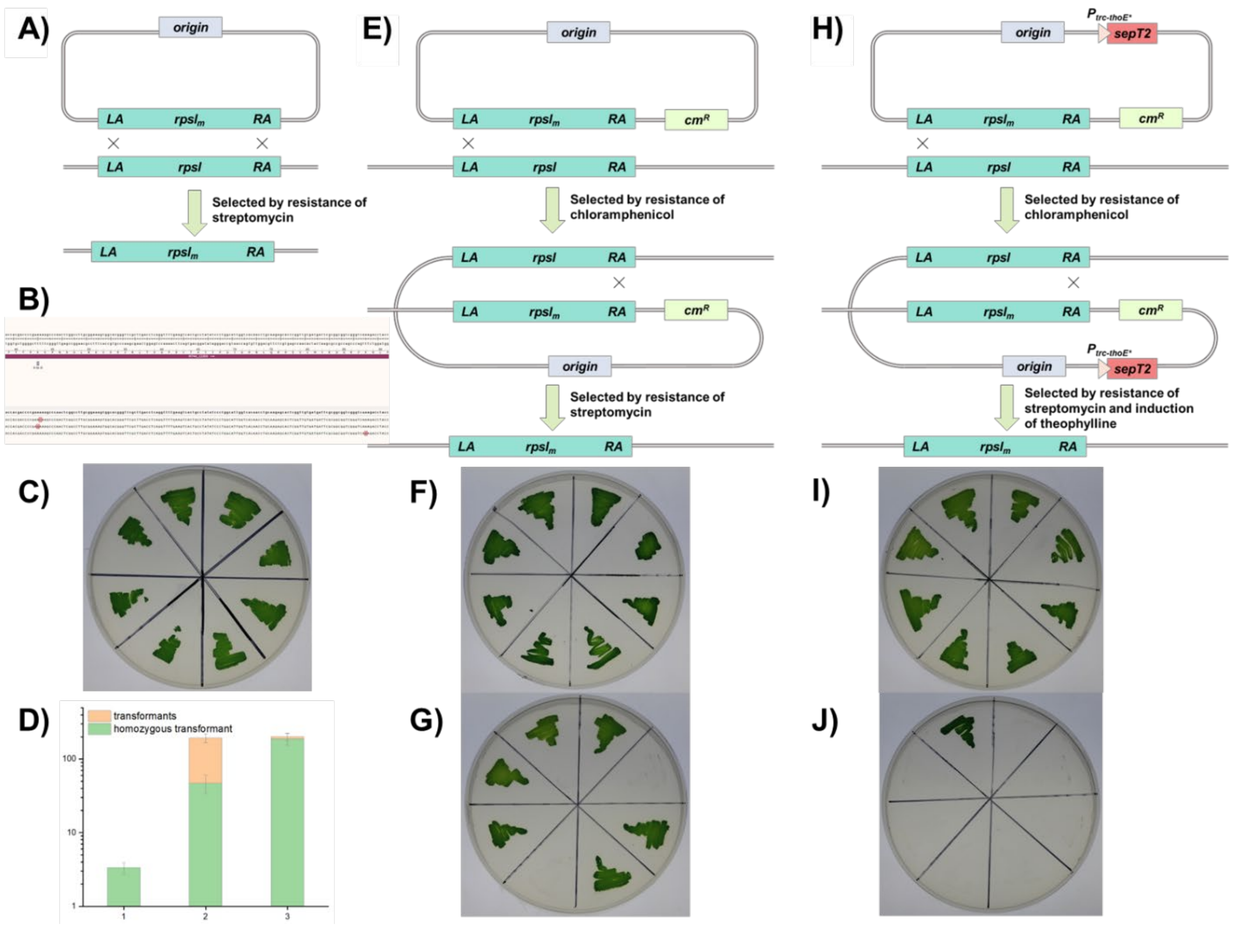

**Fig. S4. A)** Introduction of the mutation via homologous double crossover. **B)** Unexpected mutations observed during the process depicted in panel A. **C)** Growth of the resulting strains from panel A on solid medium supplemented with streptomycin. **D)** Transformation efficiency and homozygosity of rpsl12 point mutations using the strategies described in panels A, E, and H. **E)** Schematic diagram of point mutation introduction via a two-step strategy: homologous single crossover with a positive selection marker, followed by homologous double crossover using rpsl12 as a negative selection marker. **F)** Growth of strains obtained using the method in panel E on streptomycin-containing medium. **G)** Growth of strains obtained using the method in panel E on chloramphenicol-containing medium. **H)** Modified version of the strategy in panel E, incorporating SepT2 as a negative selection marker to facilitate the second crossover. **I)** Growth of strains obtained using the method in panel H on streptomycin-containing medium. J) Growth of strains obtained using the method in panel H on chloramphenicol-containing medium.

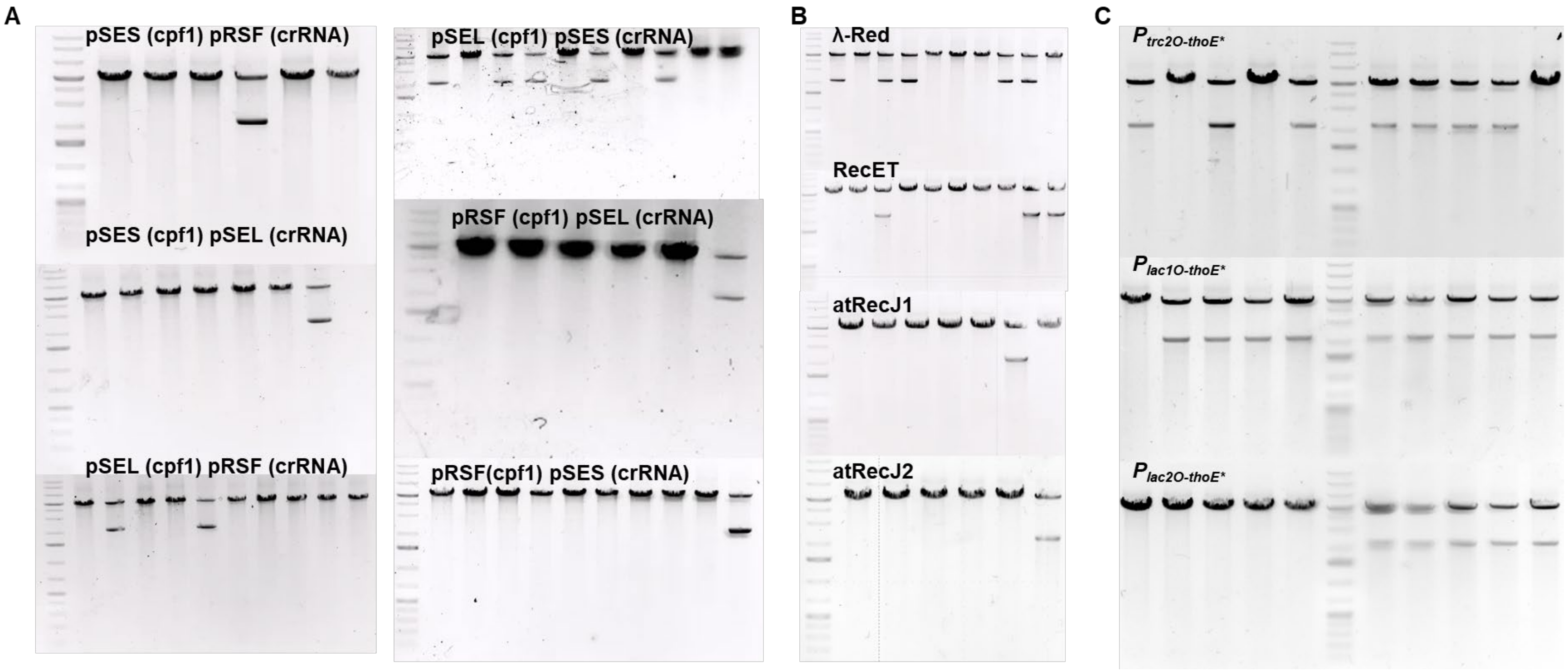

**Fig. S5 A)** Verification of survived transformants from Fig. 4E. **B)** Verification of survived transformants from Fig. 4H. **C)** Verification of surviving transformants from Fig. 4I before and after induction; lanes to the left of the marker indicate samples before induction, and lanes to the right indicate samples after induction.

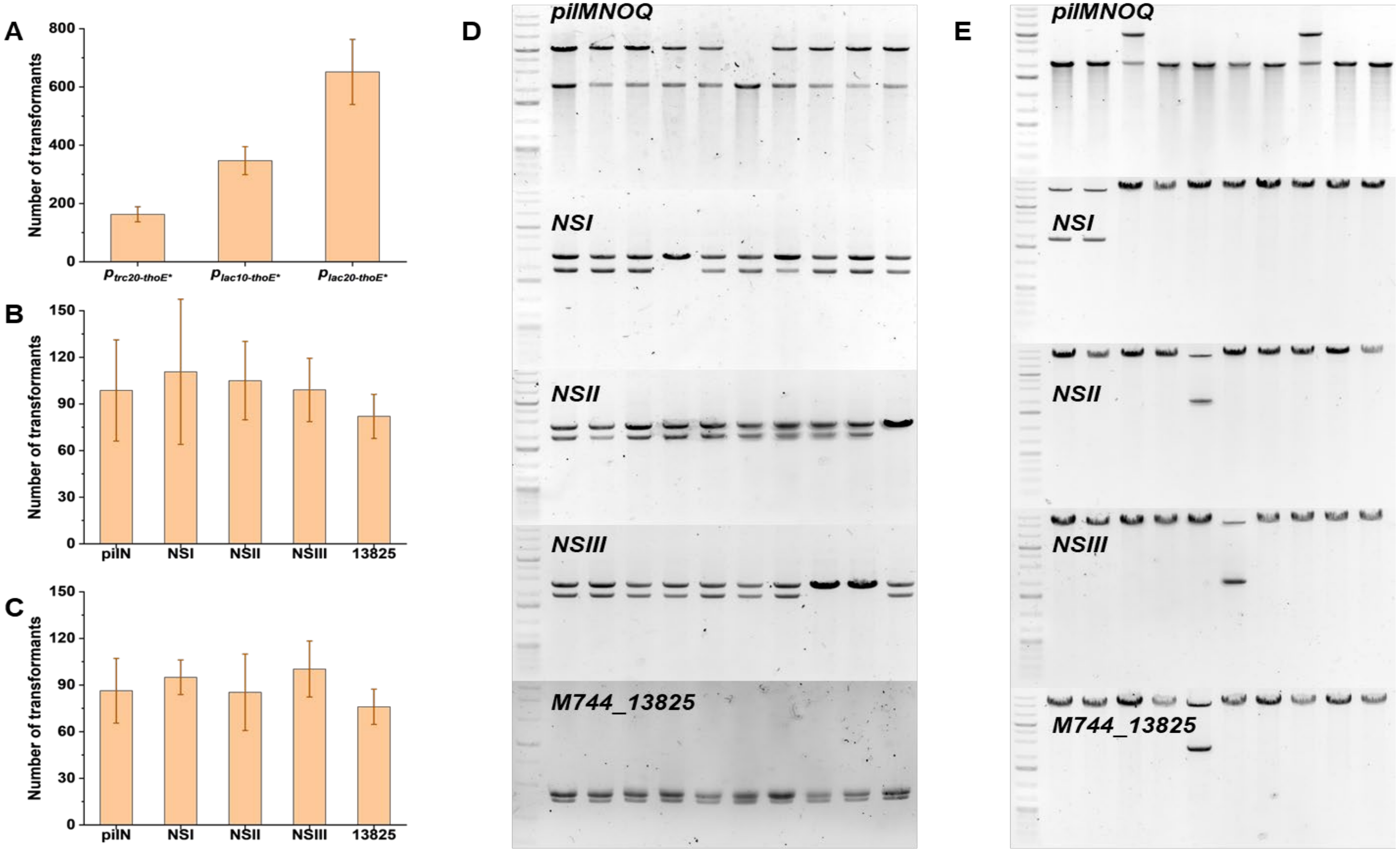

**Fig. S6. A)** Number of transformants obtained in Fig. 4I. **B)** Number of transformants obtained in Fig. 4L. **C)** Number of transformants obtained in Fig. 4M. **D)** PCR verification of surviving transformants from Fig. 4L using gene-specific F2-R2 primers as indicated in Fig. 4. **E)** PCR verification of surviving transformants from Fig. 4M using gene-specific F2-R2 primers as indicated in Fig. 4.

**Table S1.** The plasmids used in this article, the methods employed for their construction and the corresponding primers are presented. The templates marked in red are the existing ones in the laboratory, while those marked in green are the synthesized ones.

| SEQUENCE NO. | FORWARD PRIMER | SEQUNENCE | REVERSE PRIMER | SEQUNENCE | TEMPLATE | ASSEMBLY METHOD | PLASMID NAME |
|---|---|---|---|---|---|---|---|
| 1 | pbr-tb-2 | GTGACCGTCCCGGGAGCTGCATGTGTCAGA | pbr-tb-3 | TGATACCGCGcGACCCACGCTCACCGGCTC | pBR322 | DNA seamless cloning | pBR322m |
| 2 | pbr-tb-4 | GCGTGGGTCgCGCGGTATCATTGCAGCACT | pbr-tb-1 | GCAGCTCCCGGGACGGTCACAGCTTGTCTG | pBR322 | | |
| 3 | pans-spe-F | gctgtcagtgcccagctcggCGCACACCGTGGAAACGGAT | amp-spe-R | AAACAAATAGGGGTTCCGCGTTATTTGCCGACTACCTTGG | pCP3031 | DNA seamless cloning | pSES-ori |
| 4 | pans-F | ccgagctgggcactgacagc | pans-R | tcaagagcctggactgatcctcgcg | Syn2973 | | |
| 5 | pans-pbr322-F | ggatcagtccaggctcttgaCCACCTCGACCTGAATGGAAGC | amp-F | CGCGGAACCCCTATTTGTTTAT | pBR322m | | |
| 6 | pSES-F1 | ttccgtagatgtagtgctgac | pSES-R1 | aaccggcgcatatcgctggg | pSES-ori | DNA seamless clon | pSES |
| 7 | pSES-F2 | cccagcgatatgcgccggtttcagggtgttgaccaacgat | pSES-R2 | aacgggcttttaccgcgcc | pSES-ori | | |

| | | | | | | | |
|---|---|---|---|---|---|---|---|
| 8 | pSES-F3 | ggcgcggtaaaagcccgtttcgaactgcacctcgctag | pSES-R3 | tcagcactacatctacggaaaccgcagcatgacacaag | pSES-ori | ing | |
| 9 | panl-F | ccatgatcgagcgatcgg | panl-R | ccataaagcaatgcggaccg | Syn2973 | DNA seamless cloning | pSEL-ori |
| 10 | panl-spe-F | ggccgatcgctcgatcatggCGCACACCGTGGAAACGGAT | panl-pbr322-F | cggtccgcattgctttatggCCACCTCGACCTGAATGGAAGC | pSES-ori | | |
| 11 | SD-pSEL-F | atcgccctctCCACCTCGACCTGAATGGAAGC | SD-pSEL-R | GTCGAGGTGGagagggcgatcggttctcggat | pSEL-L | DNA seamless cloning | pSEL |
| 12 | rsf-spe-F | attaccgcctttgagtgagcCGCACACCGTGGAAACGGAT | rsf-spe-R | caaggcggcaggctgaccccTTATTTGCCGACTACCTTGG | pSES-ori | DNA seamless cloning | pRSF-ori |
| 13 | rsf-F | gctcactcaaaggcggtaatcaattg | rsf-R | ggggtcagcctgccgccttg | pJA-tho-PilA | | |
| 14 | repF-del-R | gcgcttggcgatcagtcaccgagaaacttg | repF-del-F | ggtgactgatcgccaagcgcgatatgccga | pRSF-ori | DNA seamless cloning | pRSF |
| 15 | spe-F | CGCACACCGTGGAAACGGAT | amp-F | CGCGGAACCCCTATTTGTTT | pSI-trc- | DNA | pSI-singl |

| | | | | | | | |
|---|---|---|---|---|---|---|---|
| | | | | | lacZ | seamless cloning | e |
| 16 | spe-in-RF | ATCCGTTTCCACGGTGTGCG | spe-in-F | GAAGGCACGAACCCAGTGGA | pSES | DNA seamless cloning | pSES-2k |
| 17 | spe-go-F | TCCACTGGGTTCGTGCCTTCttaaacaggatctgcattgc | spe-2k-R | CGCACACCGTGGAAACGGATgttgagcgactggcggatta | MG1655 | | |
| 18 | spe-in-RF | ATCCGTTTCCACGGTGTGCG | spe-in-F | GAAGGCACGAACCCAGTGGA | pSES | DNA seamless cloning | pSES-4k |
| 19 | spe-go-F | TCCACTGGGTTCGTGCCTTCttaaacaggatctgcattgc | spe-4k-R | CGCACACCGTGGAAACGGATcactaaagcgcggctggcag | MG1655 | | |
| 20 | spe-in-RF | ATCCGTTTCCACGGTGTGCG | spe-in-F | GAAGGCACGAACCCAGTGGA | pSES | DNA seamless cloning | pSES-8k |
| 21 | spe-go-F | TCCACTGGGTTCGTGCCTTCttaaacaggatctgcattgc | spe-8k-R | CGCACACCGTGGAAACGGATggctgaaggaatcgcttctt | MG1655 | | |
| 22 | spe-in-RF | ATCCGTTTCCACGGTGTGCG | spe-in-F | GAAGGCACGAACCCAGTGGA | pSES | DNA seamless cloning | pSES-12k |
| 23 | spe-go-F | TCCACTGGGTTCGTGCCTTCttaaacaggatctgcattgc | spe-12k-R | CGCACACCGTGGAAACGGATtaccgaccataccgaactgg | MG1655 | | |

| | | | | | | | |
|---|---|---|---|---|---|---|---|
| 24 | spe-in-RF | ATCCGTTTCCACGGTGTGCG | spe-in-F | GAAGGCACGAACCCAGTGGA | pSEL | DNA seamless cloning | pSEL-2k |
| 25 | spe-go-F | TCCACTGGGTTCGTGCCTTCttaaacaggatctgcattgc | spe-2k-R | CGCACACCGTGGAAACGGATgttgagcgactggcggatta | MG1655 | | |
| 26 | spe-in-RF | ATCCGTTTCCACGGTGTGCG | spe-in-F | GAAGGCACGAACCCAGTGGA | pSEL | DNA seamless cloning | pSEL-4k |
| 27 | spe-go-F | TCCACTGGGTTCGTGCCTTCttaaacaggatctgcattgc | spe-4k-R | CGCACACCGTGGAAACGGATcactaaagcgcggctggcag | MG1655 | | |
| 28 | spe-in-RF | ATCCGTTTCCACGGTGTGCG | spe-in-F | GAAGGCACGAACCCAGTGGA | pSEL | DNA seamless cloning | pSEL-8k |
| 29 | spe-go-F | TCCACTGGGTTCGTGCCTTCttaaacaggatctgcattgc | spe-8k-R | CGCACACCGTGGAAACGGATggctgaaggaatcgcttctt | MG1655 | | |
| 30 | spe-in-RF | ATCCGTTTCCACGGTGTGCG | spe-in-F | GAAGGCACGAACCCAGTGGA | pSEL | DNA seamless cloning | pSEL-12k |
| 31 | spe-go-F | TCCACTGGGTTCGTGCCTTCttaaacaggatctgcattgc | spe-12k-R | CGCACACCGTGGAAACGGATtaccgaccataccgaactgg | MG1655 | | |
| 32 | spe-in-RF | ATCCGTTTCCACGGTGTGCG | spe-in-F | GAAGGCACGAACCCAGTGGA | pRSF | DNA seamless | pRSF-2k |
| 33 | spe-go- | TCCACTGGGTTCGTGCCTTCttaaacaggatctgcattgc | spe-2k-R | CGCACACCGTGGAAACGGATgttgagcgactggcggatta | MG1655 | | |

| | | | | | | | |
|---|---|---|---|---|---|---|---|
| | F | | | | | cloning | |
| 34 | spe-in-RF | ATCCGTTTCCACGGTGTGCG | spe-in-F | GAAGGCACGAACCCAGTGGA | pRSF | DNA seamless cloning | pRSF-4k |
| 35 | spe-go-F | TCCACTGGGTTCGTGCCTTCttaaacaggatctgcattgc | spe-4k-R | CGCACACCGTGGAAACGGATcactaaagcgcggctggcag | MG1655 | | |
| 36 | spe-in-RF | ATCCGTTTCCACGGTGTGCG | spe-in-F | GAAGGCACGAACCCAGTGGA | pRSF | DNA seamless cloning | pRSF-8k |
| 37 | spe-go-F | TCCACTGGGTTCGTGCCTTCttaaacaggatctgcattgc | spe-8k-R | CGCACACCGTGGAAACGGATggctgaaggaatcgcttctt | MG1655 | | |
| 38 | spe-in-RF | ATCCGTTTCCACGGTGTGCG | spe-in-F | GAAGGCACGAACCCAGTGGA | pRSF | DNA seamless cloning | pRSF-12k |
| 39 | spe-go-F | TCCACTGGGTTCGTGCCTTCttaaacaggatctgcattgc | spe-12k-R | CGCACACCGTGGAAACGGATtaccgaccataccgaactgg | MG1655 | | |
| 40 | spe-trc-F | ATCCGTTTCCACGGTGTGCGATGAGCTGTTGACAATTAATCAT | pSES-lacZ-R | gctgtcagtgcccagctcggttacttctgacaccaaaccaac | pSI-trc-lacZ | DNA seamless cloning | pSES-lacZ |
| 41 | spe-F | CGCACACCGTGGAAACGGAT | pans-F | ccgagctgggcactgacagc | pSES | | |
| 42 | spe-trc-F | ATCCGTTTCCACGGTGTGCGATGAGCTGTTGACAATTAATCAT | pSES-lacZ | gctgtcagtgcccagctcggttacttctgacaccaaaccaac | pSI-trc-lacZ | DNA sea | pSES-ori-lacZ |

| | | | | | | | |
|---|---|---|---|---|---|---|---|
| | | | -R | | | mless cloning | |
| 43 | spe-F | CGCACACCGTGGAAACGGAT | pans-F | ccgagctgggcactgacagc | pSES-ori | | |
| 44 | spe-trc-F | ATCCGTTTCCACGGTGTGCGATGAGCTGTTGACAATTAATCAT | pSEL-lacZ-R | ggccgatcgctcgatcatggttacttctgacaccaaaccaac | pSI-trc-lacZ | DNA seamless cloning | pSEL-lacZ |
| 45 | spe-F | CGCACACCGTGGAAACGGAT | panl-F | ccatgatcgagcgatcgg | pSEL | | |
| 46 | spe-trc-F | ATCCGTTTCCACGGTGTGCGATGAGCTGTTGACAATTAATCAT | pSEL-lacZ-R | ggccgatcgctcgatcatggttacttctgacaccaaaccaac | pSI-trc-lacZ | DNA seamless cloning | pSEL-L-lacZ |
| 47 | spe-F | CGCACACCGTGGAAACGGAT | panl-F | ccatgatcgagcgatcgg | pSEL-L | | |
| 48 | spe-trc-F | ATCCGTTTCCACGGTGTGCGATGAGCTGTTGACAATTAATCAT | pRSF-lacZ-R | attaccgcctttgagtgagcttacttctgacaccaaaccaac | pSI-trc-lacZ | DNA seamless cloning | pRSF-lacZ |
| 49 | spe-F | CGCACACCGTGGAAACGGAT | rsf-F | gctcactcaaaggcggtaatcaattg | pRSF | | |
| 50 | spe-trc-F | ATCCGTTTCCACGGTGTGCGATGAGCTGTTGACAATTAATCAT | pRSF-lacZ-R | attaccgcctttgagtgagcttacttctgacaccaaaccaac | pSI-trc-lacZ | DNA seamless cloning | pRSF-ori-lacZ |
| 51 | spe-F | CGCACACCGTGGAAACGGAT | rsf-F | gctcactcaaaggcggtaatcaattg | pRSF-ori | | |
| 52 | sac | aaggatcgatcctctagctactgatccttcaactcagcaa | amp | AAACAAATAGGGGTTCCGCGTTAgaaaaactcatcgagca | km | DN | pSE |

| | | | | | | | |
|---|---|---|---|---|---|---|---|
| | B-km-F | | -km-R | | | A seamless cloning | L-sacB |
| 53 | panl-F | ccatgatcgagcgatcgg | amp-F | CGCGGAACCCCTATTTGTTTAT | pSEL | | |
| 54 | pSEL-sacB-R | ggccgatcgctcgatcatggttatttgttaactgttaatt | sacB-F | tagctagaggatcgatcctt | pCPF1b | | |
| 55 | gg-spe-F | tacggtctcgACGGATGAAGGCACGAAC | gg-spe-R | tacggtctcaTTTGCCGACTACCTTGGTG | pSI-trc-lacZ | Golden Gate assembly bsaI | pAC-lacI-sepA2 |
| 56 | gg-pac-F | tacggtctcacaaataaTTTTGAGGTGCTCCAGTGGC | gg-pac-R | tacggtctcttcaaTCCGTTAGCGAGGTGCCG | pACYC184 | | |
| 57 | gg-lacI-F | tacggtctctTTGACAGCTAGCTCAGTCCTAGGTATAATGCTAGCATCTATACTGGAAGAGAGTCAATTCAGGGTGGTGAATGTGAAACCAGTAACGTTAT | gg-lacI-R | tacggtctccTGCCCGCTTTCCAGTCGG | mg1655 | | |
| 58 | gg-sepA2-F | tacggtctccggcagtgaTCAGTAGGTTGCAGCTCC | gg-sepA2-R | tacggtctcgccgtttccacggtgtgcgCTAGTCTTCAGGCCAGTC | Syn2973 | | |
| 59 | panl-F | ccatgatcgagcgatcgg | tho-km-F | TAGGAATAACTAAGGAATTCctgatccttcaactcagcaa | pSEL-sacB | DNA seamless cloning | pSEL-sepT2 |
| 60 | tho-F | GAATTCCTTAGTTATTCCTATTCTGCAC | pSEL-sepT2-R | ggccgatcgctcgatcatggCGCAGAAAGGCCCACCCGAA | tho-sepT2 | | |
| 61 | psbA2-R | TTTGCGATGAGTCCTTAGTT | psbA2-F | GAAAAGTCTGAAAGTTCTTTA | Syn2973 | DNA seamles | pSEL-68rps1 |
| 62 | panl | ggccgatcgctcgatcatggttatttcttcgctttttcccg | psba | AACTAAGGACTCATCGCAAAatgccaactatccagcagct | Syn6 | | |

| | | | | | | | |
|---|---|---|---|---|---|---|---|
| | -rpsl-R | | 2-rpsl-F | | 803 | s cloning | |
| 63 | panl-F | ccatgatcgagcgatcgg | psab2-km-F | TAAAGAACTTTCAGACTTTTCctgatccttcaactcagcaa | pSEL | | |
| 64 | tetA-km-F | GCACTTTTCGGGGAAATGTGctgatccttcaactcagcaa | panl-F | ccatgatcgagcgatcgg | pSEL | DNA seamless cloning | pSEL-tetA |
| 65 | tetA-F | CACATTTCCCCGAAAAGTGC | panl-tetA-R | ggccgatcgctcgatcatggTCAGGTCGAGGTGGCCC | pBR322m | | |
| 66 | rpsl-F1 | AAACAAATAGGGGTTCCGCGaggaggagggccactcctcg | rpsl-R1 | ttgggctttctcggggtcgtggtgtagacgc | Syn2973 | DNA seamless cloning | pBR322-rpslm |
| 67 | rpsl-F2 | acgaccccgagaaagcccaactcggccttg | rpsl-R2 | ctagtagcggtagtgggcaaaa | Syn2973 | | |
| 68 | rpsl-pbr-F | ttgcccactaccgctactagCCACCTCGACCTGAATGGAAGC | amp-F | CGCGGAACCCCTATTTGTTTAT | pBR322m | | |
| 69 | cm-R | TTACGCCCCGCCCTGCC | rpsl-cm-F | ttgcccactaccgctactagTACCTGTGACGGAAGATCAC | pSII-cm | DNA seamless cloning | pBR322-rpslm-cm |
| 70 | rpsl-R2 | ctagtagcggtagtgggcaaaa | cm-pbr-F | GGCAGGGCGGGGCGTAACCACCTCGACCTGAATGGAA | pBR322-rpslm | | |
| 71 | rpsl-R2 | ctagtagcggtagtgggcaaaa | sepT2-cm- | TTCGGGTGGGCCTTTCTGCGTACCTGTGACGGAAGATCAC | pBR322-rpsl | DNA sea | pBR322-rpsl |

| | | | | | | | |
|---|---|---|---|---|---|---|---|
| | | | F | | m-cm | mless cloning | m-sepT2 |
| 72 | rpsl-tho-F | ttgcccactaccgctactagGAATTCCTTAGTTATTCCTATTCTGCAC | sepT2-R | CGCAGAAAGGCCCACCCGAA | pSEL-sepT2 | | |
| 73 | amp-F | CGCGGAACCCCTATTTGTTTAT | pbr-F | CCACCTCGACCTGAATGGAAGC | pBR322m | DNA seamless cloning | pBR-pil-km |
| 74 | amp-pil-up-F | AAACAAATAGGGGTTCCGCGcttagcgcttactctctcgg | pil-up-R | ttagttcagatcatccaggc | Syn2973 | | |
| 75 | pil-km-F | cgttctcagaccgccacttctttcgacctcattctattagact | pil-km-R | gcctggatgatctgaactaattagaaaaactcatcgagcatc | pSEL-sacB | | |
| 76 | pil-down-F | gaagtggcggtctgagaacg | pbr-pil-down-R | TTCCATTCAGGTCGAGGTGGagagtaatgaagccattgtc | Syn2973 | | |
| 77 | pil-d-R | agagtaatgaagccattgtc | erm-pbr-F | GTTTTTTCTTTGTGAGTCCACCACCTCGACCTGAATGGAA | pBR-pil-Km | DNA seamless cloning | pBR-pil-km-sepT2 |
| 78 | sepT2-erm-F | CGGGTGGGCCTTTCTGCGCGCACACCGTGGAAACGGAT | erm-R | TGGACTCACAAAGAAAAAACGC | Erm | | |
| 79 | pil-sepT2-F | gacaatggcttcattactctGAATTCCTTAGTTATTCCTATTC | sepT2-R | CGCAGAAAGGCCCACCCG | pSEL-sepT2 | | |
| 80 | amp-F | CGCGGAACCCCTATTTGTTTAT | pbr-F | CCACCTCGACCTGAATGGAAGC | pBR322 | DNA | pBR-pil- |

| | | | | | | | |
|---|---|---|---|---|---|---|---|
| | | | | | m | seamless cloning | KO |
| 81 | amp-pil-up-F | AAACAAATAGGGGTTCCGCGcttagcgcttactctctcgg | pil-up-R | ttagttcagatcatccaggc | Syn2973 | | |
| 82 | pil-cm-R | gcctggatgatctgaactaattacgccccgccctgccactca | psba2-cm-F | CTCGTGAGTAAAATCTAACAAGCAGATTGAGTTTTGTAAAGAACTTTCAGACTTTTCtgatcggcacgtaagaggtt | pACYC184 | | |
| 83 | psba2-rpsl-F | TAGATTTTACTCACGAGGCTATTAAGTCTCGTAAATAGTTCAACTAAGGACTCATCGCAAAatgccaactatccagcagct | pil-rpsl-R | cgttctcagaccgccacttcttatttcttcgctttttcccg | pSEL-68rpsl | | |
| 84 | pil-down-F | gaagtggcggtctgagaacg | pbr-pil-down-R | TTCCATTCAGGTCGAGGTGGagagtaatgaagccattgtc | Syn2973 | | |
| 85 | pil-R | agagtaatgaagccattgtc | erm-pbr-F | GTTTTTTCTTTGTGAGTCCACCACCTCGACCTGAATGGAA | pBR-pil-KO | DNA seamless cloning | pBR-pil-sepT2 |
| 86 | pil-sept2-F | gacaatggcttcattactctGAATTCCTTAGTTATTCCTATTC | erm-R | TGGACTCACAAAGAAAAAACGC | pBR-pil-km-sepT2 | | |
| 87 | pil-amp-F | ccgagagagtaagcgctaagaCGCGGAACCCCTATTTGTTT | pil-tho-F | gacaatggcttcattactctGAATTCCTTAGTTATTCCTA | pBR-pil-sepT2 | DNA seamless cloning | pBR-pilNm-sepT2 |
| 88 | pil-up-F | tcttagcgcttactctctcgg | pil-down-R | agagtaatgaagccattgtc | Syn2973 | | |
| 89 | amp | CGCGGAACCCCTATTTGTTTAT | pbr- | CCACCTCGACCTGAATGGAAGC | pBR | DN | pBR- |

| | | | | | | | |
|---|---|---|---|---|---|---|---|
| | -F | | F | | 322 | A seamless cloning | pil-KO-km |
| 90 | amp-pil-up-F | AAACAAATAGGGGTTCCGCGtgatcacgacttgggcgatc | pil-up-R | gagaagcgcctttttatttg | Syn2973 | | |
| 91 | pil-km-R | caaataaaaaggcgcttctcttagaaaaactcatcgagcatc | pil-rpsl-R | cgttctcagaccgccacttcttatttcttcgctttttcccg | pSEL-68rpsl | | |
| 92 | pil-down-F | gaagtggcggtctgagaacg | pbr-pil-down-R | TTCCATTCAGGTCGAGGTGGctagggcagcggagcgaccg | Syn2973 | | |
| 93 | tho-F | GAATTCCTTAGTTATTCCTA | cm-pil-R | agtggcagggcggggcgtaaagagtaatgaagccattgtc | pBR-pilNm-sepT2 | DNA seamless cloning | pBR3-pilNm-sepT2 |
| 94 | cm-R | ttacgccccgccctgccactc | tho-pil-R | TAGGAATAACTAAGGAATTCagagtaatgaagccattgtc | pBR-pil-sepT2 | | |
| 95 | gg-km-F | tacggtctcgTCAACTCAGCAAAAGTTCGATTTATTC | gg-pSEL-F | tacggtctcatagcCATGATCGAGCGATCGG | pSEL-sepT2 | Golden Gate assembly bsaI | pLLACCP-pilN-km-un |
| 96 | gg-lac-F | tacggtctcgttgaaggatcagTTTACACTTTATGCTTCCGgctcgtatgttGTGTGGTGCTAAGGAGGCAACAAGatgtcaatttatcaagaatt | gg-cpf1-R | tacggtctcaTTAGTTATTCCTATTCTGCAC | pCPF1b | | |
| 97 | gg-cr-F | tacggtctcactaaTTGACAGCTAGCTCAGTC | gg-cr-R | tacggtctctAAATGACCTTCATAAATCGC | crRNA-pilN | | |
| 98 | gg- | ggctacggtctctattttttttgccggtcttagcgcttact | gg- | ggctacggtctctgaagctgcaggcgattcc | Syn2 | | |

| | | | | | | | |
|---|---|---|---|---|---|---|---|
| | pil-F1 | | pil-R1 | | 973 | | |
| 99 | gg-km-R2 | ggctacggtctctcttcaggattttgtttagaaaaactcatcgagc | gg-km-F2 | ggctacggtctcgtggggatcgcccatttcgacctcattctattag | pSEL-sacB | | |
| 100 | gg-pil-F2 | ggctacggtctcgcccaagcaatcaactttgcgac | gg-pil-R2 | ggctacggtctccgctactaagagtaatgaagccattgtc | Syn2973 | | |
| 101 | amp-cm-F | AAACAAATAGGGGTTCCGCGttacgccccgccctgccact | km-cm-F | ccggaagcataaagtgtaaatgatcggcacgtaagaggtt | pBR-pil-KO | DNA seamless cloning | pLLACCP-pilN-km |
| 102 | amp-F | CGCGGAACCCCTATTTGTTT | lac-F | tttacactttatgcttccgg | pLLACCP-pilN-km-un | | |
| 103 | tho-cm-R | attaattgtcaacagctcattgatcggcacgtaagaggtt | tho-cpf1-F | cctgctaaggaggcaacaagatgtcaatttatcaagaatttgtta | pLLACCP-pilN-km | DNA seamless cloning | pLTHOCP-pilN-km |
| 104 | tho-R | cttgttgcctccttagcagg | tho-F | atgagctgttgacaattaat | pSI-tho-lacZ | | |
| 105 | trc-cm-F | TCCACACATTATACGAGCCGGATGattaattgtcaacagctcattgatcggcacgtaagaggtt | trc-cpf1-F | CGGCTCGTATAATGTGTGGAATTGTGAGCGGATAACAATTTCACACAatgtcaatttatcaagaatttgtta | pLLACCP-pilN-km | DNA seamless cloning | pLTRCCP-pilN-km |

| | | | | | | | |
|---|---|---|---|---|---|---|---|
| 106 | pSES-pil-R | gctgtcagtgcccagctcggagagtaatgaagccattgtc | pbr322-F | ggatcagtccaggctcttgaCCACCTCGACCTGAATGGAAGC | pLLACCP-pilN-km | DNA seamless cloning | pSLACCP-pilN-km |
| 107 | pans-F | ccgagctgggcactgacagc | pans-R | tcaagagcctggactgatcctcgcg | pSES | | |
| 108 | pSES-pil-R | gctgtcagtgcccagctcggagagtaatgaagccattgtc | pbr322-F | ggatcagtccaggctcttgaCCACCTCGACCTGAATGGAAGC | pLTRCCP-pilN-km | DNA seamless cloning | pSTRCCP-pilN-km |
| 109 | pans-F | ccgagctgggcactgacagc | pans-R | tcaagagcctggactgatcctcgcg | pSES | | |
| 110 | pSES-pil-R | gctgtcagtgcccagctcggagagtaatgaagccattgtc | pbr322-F | ggatcagtccaggctcttgaCCACCTCGACCTGAATGGAAGC | pLTHOCP-pilN-km | DNA seamless cloning | pSTHOCP-pilN-km |
| 111 | pans-F | ccgagctgggcactgacagc | pans-R | tcaagagcctggactgatcctcgcg | pSES | | |
| 112 | RSF-pil-R | attaccgcctttgagtgagcagagtaatgaagccattgt | amp-R | TTACCAATGCTTAATCAGTGAG | pLLACCP-pilN-km | DNA seamless cloning | pRFLACCP-pilN-km |
| 113 | RSF-F | gctcactcaaaggcggtaat | amp-RSF-R | CACTGATTAAGCATTGGTAAggggtcagcctgccgcctt | pRSF-ori | | |
| 114 | RSF-pil-R | attaccgcctttgagtgagcagagtaatgaagccattgt | amp-R | TTACCAATGCTTAATCAGTGAG | pLTRCCP- | DNA sea | pRFTRCCP- |

| | | | | | | | |
|---|---|---|---|---|---|---|---|
| | | | | | pilN-km | mless cloning | pilN-km |
| 115 | RSF-F | gctcactcaaaggcggtaat | amp-RSF-R | CACTGATTAAGCATTGGTAAggggtcagcctgccgcctt | pRSF-ori | | |
| 116 | RSF-pil-R | attaccgcctttgagtgagcagagtaatgaagccattgt | amp-R | TTACCAATGCTTAATCAGTGAG | pLTHOCP-pilN-km | DNA seamless cloning | pRFTHOCP-pilN-km |
| 117 | RSF-F | gctcactcaaaggcggtaat | amp-RSF-R | CACTGATTAAGCATTGGTAAggggtcagcctgccgcctt | pRSF-ori | | |
| 118 | RSF-pil-R | attaccgcctttgagtgagcagagtaatgaagccattgt | amp-R | TTACCAATGCTTAATCAGTGAG | pLLACCP-pilN-km | DNA seamless cloning | pRLACCP-pilN-km |
| 119 | RSF-F | gctcactcaaaggcggtaat | amp-RSF-R | CACTGATTAAGCATTGGTAAggggtcagcctgccgcctt | pRSF | | |
| 120 | RSF-pil-R | attaccgcctttgagtgagcagagtaatgaagccattgt | amp-R | TTACCAATGCTTAATCAGTGAG | pLTRCCP-pilN-km | DNA seamless cloning | pRTRCCP-pilN-km |
| 121 | RSF-F | gctcactcaaaggcggtaat | amp-RSF-R | CACTGATTAAGCATTGGTAAggggtcagcctgccgcctt | pRSF | | |

| | | | | | | | |
|---|---|---|---|---|---|---|---|
| 122 | RSF-pil-R | attaccgcctttgagtgagcagagtaatgaagccattgt | amp-R | TTACCAATGCTTAATCAGTGAG | pLTHOCP-pilN-km | DNA seamless cloning | pRTHOCP-pilN-km |
| 123 | RSF-F | gctcactcaaaggcggtaat | amp-RSF-R | CACTGATTAAGCATTGGTAAggggtcagcctgccgcctt | pRSF | | |
| 124 | cr-del-R | ggagcgaccgttagttattcctattctgcacgaa | cr-del-F | gaataactaacggtcgctccgctgccctag | pSTHOCP-pilN-km | DNA seamless cloning | pSTHOCP |
| 125 | cr-del-R | ggagcgaccgttagttattcctattctgcacgaa | cr-del-F | gaataactaacggtcgctccgctgccctag | pLTHOCP-pilN-km | DNA seamless cloning | pLTHOCP |
| 126 | cr-del-R | ggagcgaccgttagttattcctattctgcacgaa | cr-del-F | gaataactaacggtcgctccgctgccctag | pRTHOCP-pilN-km | DNA seamless cloning | pRTHOCP |
| 127 | amp-pil-R | AAACAAATAGGGGTTCCGCGagagtaatgaagccattgtc | spe-cr-F | CCAAGGTAGTCGGCAAATAAttgacagctagctcagtcct | pSTHOCP- | DNA sea | pSCR-pilN- |

| | | | | | | | |
|---|---|---|---|---|---|---|---|
| | | | | | pilN-km | mless cloning | km |
| 128 | amp-F | CGCGGAACCCCTATTTGTTT | spe-R | TTATTTGCCGACTACCTTGG | pSES | | |
| 129 | amp-pil-R | AAACAAATAGGGGTTCCGCGagagtaatgaagccattgtc | spe-cr-F | CCAAGGTAGTCGGCAAATAAttgacagctagctcagtcct | pSTHOCP-pilN-km | DNA seamless cloning | pLCR-pilN-km |
| 130 | amp-F | CGCGGAACCCCTATTTGTTT | spe-R | TTATTTGCCGACTACCTTGG | pSEL | | |
| 131 | spe-F | CGCACACCGTGGAAACGGAT | RSF-F | gctcactcaaaggcggtaat | pRSF | DNA seamless cloning | pRCR-pilN-km |
| 132 | spe-cr-F | ATCCGTTTCCACGGTGTGCGttgacagctagctcagtcct | RSF-pil-R | attaccgcctttgagtgagcagagtaatgaagccattgtc | pSTHOCP-pilN-km | | |
| 133 | recE-cpcB1-R | aagagtggttttgtgctcattcaaccagtctcctgttctc | spe-cpcB1-F | ATCCGTTTCCACGGTGTGCGcagtctcagctgcatgctgg | Syn2973 | DNA seamless cloning | pRSF-recET |
| 134 | RSF-F | gctcactcaaaggcggtaat | RSF-R | ggggtcagcctgccgccttg | pRSF | | |
| 135 | recE-F | atgagcacaaaaccactctt | RSF-recT-R | attaccgcctttgagtgagcttattcctctgaattatcgattac | MG1655 | | |
| 136 | erm-F | CGCACACCGTGGAAACGGAT | RSF-erm-R | caaggcggcaggctgaccccTGGACTCACAAAGAAAAACGC | pBR-pilNm-sepT | | |

| | | | | | | | |
|---|---|---|---|---|---|---|---|
| | | | | | 2 | | |
| 137 | cpcB-red-F | gagaacaggagactggttgaatggatattaatactgaaactgag | RSF-red-R | attaccgcctttgagtgagcttaactcaacagaagatgctttg | pKD46 | DNA seamless cloning | pRSF-λred |
| 138 | cpcB-R | tcaaccagtctcctgttctc | RSF-F | gctcactcaaaggcggtaat | pRSF-recET | | |
| 139 | cpcB-R | tcaaccagtctcctgttctc | RSF-F | gctcactcaaaggcggtaat | pRSF-recET | DNA seamless cloning | pRSF-atRecJ1 |
| 140 | pRSF-1886-R | attaccgcctttgagtgagcAGGAGGAATTAACCATGCAGTGGTGGTGGTGGTGGTGCttgagtctccctacaattga | cpcB-pt-1886-F | gagaacaggagactggttgaAGGAGGAATTAACCATGCAGTGGTGGTGGTGGTGGTGCcaaggatccaacgctgctcg | Syn2973 | | |
| 141 | cpcB-pt-0517-F | gagaacaggagactggttgaAGGAGGAATTAACCATGCAGTGGTGGTGGTGGTGGTGCccagcacctcggcgtagtca | pRSF-0517-R | attaccgcctttgagtgagcAGGAGGAATTAACCATGCAGTGGTGGTGGTGGTGGTGCatgcagcggacctggcgatc | Syn2973 | DNA seamless cloning | pRSF-atRecJ2 |
| 142 | cpcB-R | tcaaccagtctcctgttctc | RSF-F | gctcactcaaaggcggtaat | pRSF-recET | | |
| 143 | cm-lacI-R | acctcttacgtgccgatcatcactgcccgctttccagtc | trc-j23-F | caacgacagaccaaaatatcaattgtgagcgctcacaattttgacagctagctcagtcct | pAC-lacI-sepA2 | DNA seamless clon | pLTRC2OTCP |
| 144 | cm-F | tgatcggcacgtaagaggt | trc-cpf1 | CCTGCTAAGGAGGCAACAAGatgtcaatttatcaagaatttg | pLTHOC | | |

| | | | | | | | |
|---|---|---|---|---|---|---|---|
| | | | -F | | P | ing | |
| 145 | trc20-F | gatattttggtctgtcgttg | tho-R | CTTGTTGCCTCCTTAGCAGG | trc2o-tho | | |
| 146 | trc-j23-F | caacgacagaccaaaatatcaattgtgagcgctcacaattttgacagctagctcagtcct | trc-cpf1-F | CCTGCTAAGGAGGCAACAAGatgtcaatttatcaagaatttg | pLTRC2OTCP | DNA seamless cloning | pLLAC2OTCP |
| 147 | trc20-F | gatattttggtctgtcgttg | tho-R | CTTGTTGCCTCCTTAGCAGG | lac2o-tho | | |
| 148 | trc-1o-j23-F | caacgacagaccaaaatatcttgacagctagctcagtcct | trc-cpf1-F | CCTGCTAAGGAGGCAACAAGatgtcaatttatcaagaatttg | pLTRC2OTCP | DNA seamless cloning | pLLAC1OTCP |
| 149 | trc-R | CTTGTTGCCTCCTTAGCAGG | trc1o-F | gatattttggtctgtcgttg | lac2o-tho | | |
| 150 | gg-cm-F | ggctacggtctcgagttttacgccccgccctgcca | gg-cpf1-R | tac**ggtctcc*****ctga***gactgTTAGTTATTCCTATTCTGCAC | pLLAC2OTCP | Golden Gate assembly bsaI | pCPF1-pil |
| 151 | gg-cpcB-F | tac**ggtctc**c***TCAG***CTGCATGCTGGTTG | gg-λred-R | tac**ggtctc**g***atta***ACTCAACAGAAGATGCTTTGTG | pRSF-λred | | |
| 152 | gg-cr-F | tacggtctcgtaatTGACAGCTAGCTCAGTCCTAGGTATAATGCTAG | gg-in-R | tacggtctcaGGCAGagagtaatgaagccattgtc | pSTHOCP-pilN-km | | |
| 153 | gg-pbr-F | tac**ggtctca***tgcc*ctagCACATTTCCCCGAAAAGTG | gg-pbr-R | tac**ggtctc**g***aact***GTCAGACCAAGTTTAC | pBR322m | | |
| 154 | λred | agcatcttctgttgagttaattgacagctagctcagtcct | cr-R | aaaaaaatgaccttcataaa | crRN | DN | pCP |

| | | | | | | | |
|---|---|---|---|---|---|---|---|
| | -j23-F | | | | A-NSI-1 | A seamless cloning | F1-NSI |
| 155 | in-F | ttaaacaggatctgcattgc | in-R | ggacatgtgttcccgacgca | MG1655 | | |
| 156 | cr-nsi-up-F | **tttatgaaggtcattttttt**ggaagggcgatcgcagcacg | in-nsi-up-R | gcaatgcagatcctgtttaacgggtcaatccgactcgccc | Syn2973 | | |
| 157 | in-nsi-down-F | tgcgtcgggaacacatgtcccgatcacatcggcttgaagt | pcpf1-nsi-down-R | **GCACTTTTCGGGGAAATGTG**caatgccttctccaagggcg | Syn2973 | | |
| 158 | pcpf1-F | CACATTTCCCCGAAAAGTGC | λred-R | ttaactcaacagaagatgctttg | pCPF1-pil | | |
| 159 | λred-j23-F | agcatcttctgttgagttaattgacagctagctcagtcct | cr-R | aaaaaaatgaccttcataaa | crRNA-NSII-1 | DNA seamless cloning | pCPF1-NSII |
| 160 | in-F | ttaaacaggatctgcattgc | in-R | ggacatgtgttcccgacgca | MG1655 | | |
| 161 | cr-nsii-up-F | tttatgaaggtcattttttaattctggatgaggcgactt | in-nsii-up-R | gcaatgcagatcctgtttaagccgtcgtggtaaaaatcgc | Syn2973 | | |
| 162 | in-nsii-down-F | tgcgtcgggaacacatgtccgaagggggagctgacgatcg | pcpf1-nsii-down-R | GCACTTTTCGGGGAAATGTGtaacccctgctcgcgccgca | Syn2973 | | |
| 163 | pcpf | CACATTTCCCCGAAAAGTGC | λred | ttaactcaacagaagatgctttg | pCP | | |

| | | | | | | | |
|---|---|---|---|---|---|---|---|
| | 1-F | | -R | | F1-pil | | |
| 164 | λred-j23-F | agcatcttctgttgagttaattgacagctagctcagtcct | cr-R | aaaaaaatgaccttcataaa | crRNA-NSIII-1 | DNA seamless cloning | pCPF1-NSIII |
| 165 | in-F | ttaaacaggatctgcattgc | in-R | ggacatgtgttcccgacgca | MG1655 | | |
| 166 | cr-nsiii-up-F | tttatgaaggtcatttttttatcacagtcggcgtcacggc | in-nsiii-up-R | gcaatgcagatcctgtttaagctccttgccactgtcacgg | Syn2973 | | |
| 167 | in-nsiii-down-F | tgcgtcgggaacacatgtccatcgcgtgacgccagctaca | pcpf1-nsiii-down-R | GCACTTTTCGGGGAAATGTGcgacgacggtgctcggtgcc | Syn2973 | | |
| 168 | pcpf1-F | CACATTTCCCCGAAAAGTGC | λred-R | ttaactcaacagaagatgctttg | pCPF1-pil | | |
| 169 | λred-j23-F | agcatcttctgttgagttaattgacagctagctcagtcct | cr-R | aaaaaaatgaccttcataaa | crRNA-13825-1 | DNA seamless cloning | pCPF1-13825 |
| 170 | in-F | ttaaacaggatctgcattgc | in-R | ggacatgtgttcccgacgca | MG1655 | | |
| 171 | cr-13825-up-F | tttatgaaggtcatttttttccaacagctggtgcccgcgc | in-13825-R | gcaatgcagatcctgtttaaggccgctgccatctagcttc | Syn2973 | | |
| 172 | in- | tgcgtcgggaacacatgtccatgcgcctcggcaaagttgt | pcpf | GCACTTTTCGGGGAAATGTGggtggatttaagatggaaaa | Syn2 | | |

| | | | | | | | |
|---|---|---|---|---|---|---|---|
| | 13825-F | | 1-13825-R | | 973 | | |
| 173 | pcpf1-F | CACATTTCCCCGAAAAGTGC | λred-R | ttaactcaacagaagatgctttg | pCPF1-pil | | |
| 174 | λred-j23-F | agcatcttctgttgagttaattgacagctagctcagtcct | cr-R | aaaaaaatgaccttcataaa | crRNA-06260-1 | DNA seamless cloning | pCPF1-06260 |
| 175 | in-F | ttaaacaggatctgcattgc | in-R | tcagccaacagcgactgaca | MG1655 | | |
| 176 | cr-06260-up-F | tttatgaaggtcatttttttggcacccacaccggagctgc | in-06260-up-R | gcaatgcagatcctgtttaactgcagatagttcagaggtg | Syn2973 | | |
| 177 | in-06260-down-F | tgtcagtcgctgttggctgatgtttagccgctatctgcag | pcpf1-06260-down-R | GCACTTTTCGGGGAAATGTGtggttgtccctgttggagca | Syn2973 | | |
| 178 | pcpf1-F | CACATTTCCCCGAAAAGTGC | λred-R | ttaactcaacagaagatgctttg | pCPF1-pil | | |
| 179 | λred-j23-F | agcatcttctgttgagttaattgacagctagctcagtcct | cr-R | aaaaaaatgaccttcataaa | crRNA-anl-1 | DNA seamless cloning | pCPF1-anl |
| 180 | in-F | ttaaacaggatctgcattgc | in-R | tcagccaacagcgactgaca | MG1655 | | |
| 181 | cr- | tttatgaaggtcatttttttagactatctcaagatagatt | in- | gcaatgcagatcctgtttaaaggcgggacacattttcatg | Syn2 | | |

| | | | | | | | |
|---|---|---|---|---|---|---|---|
| | anl-up-F | | anl-up-R | | 973 | | |
| 182 | in-anl-down-F | tgtcagtcgctgttggctgattatggatgaggctgggga | pcpf1-anl-down-R | GCACTTTTCGGGGAAATGTGatcgcatcaatagtcttttg | Syn2973 | | |
| 183 | pcpf1-F | CACATTTCCCCGAAAAGTGC | λred-R | ttaactcaacagaagatgctttg | pCPF1-pil | | |
| 184 | cpcB-F | cagtctcagctgcatgctgg | cpf1-R | ttagttattcctattctgca | pCPF1-NSI | DNA seamless cloning | pCPF1-NSI-rpsl |
| 185 | cpcB-rpsl-R | ccagcatgcagctgagactgttatttcttcgctttttccc | cpf1-psba2-F | tgcagaataggaataactaaGAAAAGTCTGAAAGTTCTTTA | pSEL-68rpsl | | |
| 186 | pcpf-gj-R | ATGGAAGCCGGCGGCACCTC | in-nsi-down-F | cgcgagagtatcgctggcagcgggtcaatccgactcgccc | pCPF1-NSI-rpsl | DNA seamless cloning | pCPF3-NSI-1 |
| 187 | pcpf1-nsi-F | GAGGTGCCGCCGGCTTCCATcgggtcaatccgactcgccc | tcr-nsi-R | GCCGGGCCACCTCGACCTGAggaagggcgatcgcagcacg | Syn2973 | | |
| 188 | tcr-R | TCAGGTCGAGGTGGCCCGGC | nsi-ds-R | cgatcacatcggcttgaagt | pCPF1-NSI-rpsl | | |
| 189 | nsi-in-F | acttcaagccgatgtgatcgttaaacaggatctgcattgc | in-R | ctgccagcgatactctcgcg | MG1655 | | |
| 190 | cr- | atgaccttcataaatcgcta | cr- | gctgatttaggcaaaaacgg | crRN | DN | pCP |

| | | | | | | | |
|---|---|---|---|---|---|---|---|
| | cz-2 | | cz-3 | | A-NSI-2 | A seamless cloning | F3-NSI-2 |
| 191 | cr-cz-1 | tagcgatttatgaaggtcat | cr-cz-4 | ccgtttttgcctaaatcagc | pCPF3-NSI-1 | | |
| 192 | cpcB-F | cagtctcagctgcatgctgg | cpf1-R | ttagttattcctattctgca | pCPF1-NSII | DNA seamless cloning | pCPF1-NSII-rpsl |
| 193 | cpcB-rpsl-R | ccagcatgcagctgagactgttatttcttcgctttttccc | cpf1-psba2-F | tgcagaataggaataactaaGAAAAGTCTGAAAGTTCTTTA | pSEL-68rpsl | | |
| 194 | pcpf-gj-R | ATGGAAGCCGGCGGCACCTC | in-nsii-us-F | cgcgagagtatcgctggcaggccgtcgtggtaaaaatcgcg | pCPF1-NSII-rpsl | DNA seamless cloning | pCPF3-NSII-1 |
| 195 | pcpf1-nsii-F | GAGGTGCCGCCGGCTTCCATgccgtcgtggtaaaaatcgc | tcr-nsii-R | GCCGGGCCACCTCGACCTGAaattctggatgaggcgactt | Syn2973 | | |
| 196 | tcr-R | TCAGGTCGAGGTGGCCCGGC | nsii-ds-R | gaagggggagctgacgatcg | pCPF1-NSII-rpsl | | |
| 197 | nsii-in-F | cgatcgtcagctccccсttcttaaacaggatctgcattgc | in-R | ctgccagcgatactctcgcg | MG1655 | | |
| 198 | cr-cz-2 | atgaccttcataaatcgcta | cr-cz-3 | gctgatttaggcaaaaacgg | crRNA-NSII-2 | DNA seamless | pCPF3-NSII-2 |
| 199 | cr- | tagcgatttatgaaggtcat | cr- | ccgtttttgcctaaatcagc | pCP | | |

| | | | | | | | |
|---|---|---|---|---|---|---|---|
| | cz-1 | | cz-4 | | F3-NSII-1 | cloning | |
| 200 | cpcB-F | cagtctcagctgcatgctgg | cpf1-R | ttagttattcctattctgca | pCPF1-NSIII | DNA seamless cloning | pCPF1-NSIII-rpsl |
| 201 | cpcB-rpsl-R | ccagcatgcagctgagactgttatttcttcgctttttccc | cpf1-psba2-F | tgcagaataggaataactaaGAAAAGTCTGAAAGTTCTTTA | pSEL-68rpsl | | |
| 202 | pcpf-gj-R | ATGGAAGCCGGCGGCACCTC | in-nsiii-down-F | cgcgagagtatcgctggcagatcgcgtgacgccagctaca | pCPF1-NSIII-rpsl | DNA seamless cloning | pCPF3-NSIII-1 |
| 203 | pcpf1-nsiii-F | GAGGTGCCGCCGGCTTCCATgctccttgccactgtcacgg | tcr-nsiii-R | GCCGGGCCACCTCGACCTGAatcacagtcggcgtcacggc | Syn2973 | | |
| 204 | nsiii-R | gctccttgccactgtcacgg | tcr-R | TCAGGTCGAGGTGGCCCGGC | pCPF1-NSIII-rpsl | | |
| 205 | nsiii-in-F | gccgtgacagtggcaaggagcttaaacaggatctgcattgc | in-R | ctgccagcgatactctcgcg | MG1655 | | |
| 206 | cr-cz-2 | atgaccttcataaatcgcta | cr-cz-3 | gctgatttaggcaaaaacgg | crRNA-NSIII-2 | DNA seamless cloning | pCPF3-NSIII-2 |
| 207 | cr-cz-1 | tagcgatttatgaaggtcat | cr-cz-4 | ccgtttttgcctaaatcagc | pCPF3- | | |

| | | | | | | | |
|---|---|---|---|---|---|---|---|
| | | | | | NSIII-1 | ing | |
| 208 | cpcB-F | cagtctcagctgcatgctgg | cpf1-R | ttagttattcctattctgca | pCPF1-13825 | DNA seamless cloning | pCPF1-13825-rpsl |
| 209 | cpcB-rpsl-R | ccagcatgcagctgagactgttatttcttcgctttttccc | cpf1-psba2-F | tgcagaataggaataactaaGAAAAGTCTGAAAGTTCTTTA | pSEL-68rpsl | | |
| 210 | pcpf-gj-R | ATGGAAGCCGGCGGCACCTC | in-13825-down-F | cgcgagagtatcgctggcagggccgctgccatctagcttc | pCPF1-13825-rpsl | DNA seamless cloning | pCPF3-13825-1 |
| 211 | pcpf1-13825-F | GAGGTGCCGCCGGCTTCCATggccgctgccatctagcttc | tcr-13825-R | GCCGGGCCACCTCGACCTGAccaacagctggtgcccgcgc | Syn2973 | | |
| 212 | tcr-R | TCAGGTCGAGGTGGCCCGGC | 13825-R | atgcgcctcggcaaagttgt | pCPF1-13825-rpsl | | |
| 213 | 13825-in-F | acaactttgccgaggcgcatttaaacaggatctgcattgc | in-R | ctgccagcgatactctcgcg | MG1655 | | |
| 214 | cr-cz-2 | atgaccttcataaatcgcta | cr-cz-3 | gctgatttaggcaaaaacgg | crRNA-13825-2 | DNA seamless | pCPF3-13825-2 |
| 215 | cr- | tagcgatttatgaaggtcat | cr- | ccgtttttgcctaaatcagc | pCP | | |

| | | | | | | | |
|---|---|---|---|---|---|---|---|
| | cz-1 | | cz-4 | | F3-13825-1 | cloning | |
| 216 | cpcB-F | cagtctcagctgcatgctgg | cpf1-R | ttagttattcctattctgca | pCPF1-06260 | DNA seamless cloning | pCPF1-06260-rpsl |
| 217 | cpcB-rpsl-R | ccagcatgcagctgagactgttatttcttcgctttttccc | cpf1-psba2-F | tgcagaataggaataactaaGAAAAGTCTGAAAGTTCTTTA | pSEL-68rpsl | | |
| 218 | pcpf-gj-R | ATGGAAGCCGGCGGCACCTC | in-06260-down-F | cgcgagagtatcgctggcagggGTCACGctgcagatagtt | pCPF1-06260-rpsl | DNA seamless cloning | pCPF3-06260-1 |
| 219 | pcpf1-06260-F | GAGGTGCCGCCGGCTTCCATggGTCACGctgcagatagtt | tcr-06260-R | GCCGGGCCACCTCGACCTGAggcacccacaccggagctgc | Syn2973 | | |
| 220 | tcr-R | TCAGGTCGAGGTGGCCCGGC | 06260-R | CTTtgtttagccgctatctg | pCPF1-06260-rpsl | | |
| 221 | 06260-in-F | cagatagcggctaaacaAAGttaaacaggatctgcattgc | in-R | ctgccagcgatactctcgcg | MG1655 | | |
| 222 | cr-cz-2 | atgaccttcataaatcgcta | cr-cz-3 | gctgatttaggcaaaaacgg | crRNA-06260-2 | DNA seamles | pCPF3-06260-2 |

| | | | | | | | |
|---|---|---|---|---|---|---|---|
| 223 | cr-cz-1 | tagcgatttatgaaggtcat | cr-cz-4 | ccgtttttgcctaaatcagc | pCPF3-06260-1 | s cloning | |
| 224 | cpcB-F | cagtctcagctgcatgctgg | cpf1-R | ttagttattcctattctgca | pCPF1-anl | DNA seamless cloning | pCPF1-anl-rpsl |
| 225 | cpcB-rpsl-R | ccagcatgcagctgagactgttatttcttcgctttttccc | cpf1-psba2-F | tgcagaataggaataactaaGAAAAGTCTGAAAGTTCTTTA | pSEL-68rpsl | | |
| 226 | pcpf-gj-R | ATGGAAGCCGGCGGCACCTC | in-anl-down-F | cgcgagagtatcgctggcagaggcgggacacattttcatg | pCPF1-anl-rpsl | DNA seamless cloning | pCPF3-anl-1 |
| 227 | pcpf1-anl-F | GAGGTGCCGCCGGCTTCCATaggcgggacacattttcatg | tcr-anl-R | GCCGGGCCACCTCGACCTGAagactatctcaagatagatt | Syn2973 | | |
| 228 | tcr-R | TCAGGTCGAGGTGGCCCGGC | anl-R | ttatggatgaggctgggga | pCPF1-anl-rpsl | | |
| 229 | anl-in-F | tcccccagcctcatccataattaaacaggatctgcattgc | in-R | ctgccagcgatactctcgcg | MG1655 | | |
| 230 | cr-cz-2 | atgaccttcataaatcgcta | cr-cz-3 | gctgatttaggcaaaaacgg | crRNA-anl-2 | DNA seamless cloning | pCPF3-anl-2 |
| 231 | cr-cz-1 | tagcgatttatgaaggtcat | cr-cz-4 | ccgtttttgcctaaatcagc | pCPF3-anl-1 | | |
| 232 | pcpf | ttaactcaacagaagatgctttg | pCP | ATGGAAGCCGGCGGCACCT | pCP | DN | pCP |

| | | | | | | | |
|---|---|---|---|---|---|---|---|
| | 1-R | | F1-F | | F3-anl-2 | A seamless cloning | F3-pil |
| 233 | tcr-pil-F | GCCGGGCCACCTCGACCTGAtgccggtcttagcgcttact | cp-pil-R | AGGTGCCGCCGGCTTCCATacaaaatcctgaagctgcag | pCPF1-pil | | |
| 234 | tcr-R | TCAGGTCGAGGTGGCCCGGC | cp-pil-F | agcatcttctgttgagttaattgacagctagctcagtcct | pCPF1-pil | | |
| 235 | amp-nsiii-F1 | AAACAAATAGGGGTTCCGCGatcacagtcggcgtcacggc | km-nsiii-R1 | tgctcgatgagtttttctaagctccttgccactgtcacgg | Syn2973 | DNA seamless cloning | pBR-NSIII-sepT2 |
| 236 | amp-F | CGCGGAACCCCTATTTGTTT | nsiii-pbr-F | caccgagcaccgtcgtcgGAATTCCTTAGTTATTCCTA | pBR-pil-sepT2 | | |
| 237 | rpsl-nsiii-F2 | gggaaaaagcgaagaaataaatcgcgtgacgccagctaca | nsiii-R2 | cgacgacggtgctcggtg | Syn2973 | | |
| 238 | km-R | ttagaaaaactcatcgagca | rpsl12-R | ttatttcttcgctttttccc | pSEL-68rpsl | | |
| 239 | nsiii-F | atcgcgtgacgccagctaca | nsiii-R | gctccttgccactgtcacgg | pBR-NSIII-sepT2 | DNA seamless cloning | pBR-NSIII-cscB |
| 240 | nsiii-tho-F | ccgtgacagtggcaaggagcatgagctgttgacaattaat | tho-R | cttgttgcctccttagcagg | pSI-tho-lacZ | | |

| | | | | | | | |
|---|---|---|---|---|---|---|---|
| 241 | tho-cscB-F | cctgctaaggaggcaacaagatggcactgaatattccatt | rbcl-cscB-R | CCGACAATCCAAACACCGGTctatattgctgaaggtacag | Escherichia coli W | | |
| 242 | rbcl-F | ACCGGTGTTTGGATTGTCGG | nsiii-rbcl-R | tgtagctggcgtcacgcgatGCTGTCGAAGTTGAACATCA | Syn2973 | | |
| 243 | tho-nsiii-R | TAGGAATAACTAAGGAATTCcgacgacggtgctcggtgcc | km-R | ttagaaaaactcatcgagca | pBR-NSIII-sepT2 | DNA seamless cloning | pBR3-NSIII-cscB |
| 244 | km-nsiii-R | tgctcgatgagtttttctaacgacgacggtgctcggtgcc | tho-F | GAATTCCTTAGTTATTCCTA | pBR-NSIII-cscB-sepT2 | | |
| 245 | cr-F | ttgacagctagctcagtcct | cr-R | aaaaaaatgaccttcataaatcgc | crRNA-NSIII-1 | DNA seamless cloning | pCPF3-NSIII-cscB |
| 246 | cr-nsiii-R | tttatgaaggtcatttttttcgacgacggtgctcggtgcc | amp-R | TTACCAATGCTTAATCAGTG | pBR3-NSIII-cscB | | |
| 247 | amp-nsiii | CACTGATTAAGCATTGGTAAcgacgacggtgctcggtgcc | cpf1-nsiii | GAGGTGCCGCCGGCTTCCATatcgcgtgacgccagctaca | Syn2973 | | |

| | | | | | | | |
|---|---|---|---|---|---|---|---|
| | -R | | -F | | | | |
| 248 | pcpf-R | ATGGAAGCCGGCGGCACCTC | cr-cpf1-R | aggactgagctagctgtcaattaactcaacagaagatgctttg | pCPF3-NSI-1 | | |
| 249 | NSI-cpc560-F | ctgccagcgatactctcgcgACCTGTAGAGAAGAGTCCCT | tho-cpc-R | gcatcaagacgatgctggtatcaccggtaccaattgtgagGACTTTATGAGTTGGGATTTTCTTAAAC | pCPF3-NSI-1 | DNA seamless cloning | pCPF3-NSI-sps-spp |
| 250 | NSI-FX-R | cgcgagagtatcgctggcag | NSI-FX-F | gcaatgcagatcctgtttaa | Syn6803 | | |
| 251 | tho-sps-F | cagcatcgtcttgatgcccttggcagcaccctgctaaggaggcaacaagatgagctattcatcaaaatac | j23-sps-R | TAGCATTATACCTAGGACTGAGCTAGCTGTCAAGAGAGCGTTCACCGACAAACAACAGATAAAACGAAAGGCCCAGTCTTTCGACTGAGCCTTTCGTTTTATTTGttaaacggggtctaacaactc | Syn6803 | | |
| 252 | j23-spp-F | AGTCCTAGGTATAATGCTAGCctcacaattggtaccggtgataccagcatcgtcttgatgcccttggcagcaccctgctaaggaggcaacaagatgcgacagttattgctaat | NSI-spp-R | ttaaacaggatctgcattgcGGCTCACCTTCGGGTGGGCCTTTCTGCGtcagctcaaaaaatcgaaat | Syn6803 | | |
| 253 | amp-invaup-F | AAACAAATAGGGGTTCCGCGttattgcggagttcccggcg | inva-up-R | ggcaaagtcctgcgggagaa | Syn2973 | DNA seamless cloning | pBR-inva-sps-spp-sepT2 |
| 254 | inva-cpc-F | ttctcccgcaggactttgccACCTGTAGAGAAGAGTCCCT | inva-T-R | cgagcagtgggttttggttgGGCTCACCTTCGGGTGGGCC | pCPF3-NSI-sps-spp | | |
| 255 | amp-F | CGCGGAACCCCTATTTGTTT | tho-F | GAATTCCTTAGTTATTCCTATTCTG | pBR-NSIII- | | |

| | | | | | | | |
|---|---|---|---|---|---|---|---|
| | | | | | cscB-sepT2 | | |
| 256 | inva-down-F | caaccaaaacccactgctcg | tho-inva-R | TAGGAATAACTAAGGAATTCgataacggcaggcgcgagta | Syn2973 | | |
| 257 | cr-F | ttgacagctagctcagtccta | cr-R | aaaaaaatgaccttcataaatcgc | crRNA-invA | DNA seamless cloning | pCPF3-inva-sps-spp |
| 258 | cr-red-R | aggactgagctagctgtcaattaactcaacagaagatgct | pcpf1-F | ATGGAAGCCGGCGGCACCT | pCPF3-NSI-sps-spp | | |
| 259 | cpf1-inva-F | AGGTGCCGCCGGCTTCCATcaaccaaaacccactgctcg | amp-inva-R | CACTGATTAAGCATTGGTAAgataacggcaggcgcgagta | Syn2973 | | |
| 260 | cr-inva-R | tttatgaaggtcatttttttgataacggcaggcgcgagta | amp-F | TTACCAATGCTTAATCAGTG | pBR-inva-sps-spp-sepT2 | | |
| 261 | nsii-in-cpcB-F | ctgccagcgatactctcgcgcagtctcagctgcatgctgg | cpcB-R | tcaaccagtctcctgttctc | Syn2973 | DNA seamless cloning | pCPF3-NSII-cugP |
| 262 | cpcB-cug | gagaacaggagactggttgaatgaaagccatgattttggc | rbcl-cugP-R | CCGACAATCCAAACACCGGTttattccggctggagaagct | Syn6803 | | |

| | | | | | | | |
|---|---|---|---|---|---|---|---|
| | P-F | | | | | | |
| 263 | NSII-in-R | cgcgagagtatcgctggcag | nsii-in-F | gcaatgcagatcctgtttaa | pCPF3-NSII-1 | | |
| 264 | rbcl-F | ACCGGTGTTTGGATTGTCGG | in-rbcl-R | ttaaacaggatctgcattgcGCTGTCGAAGTTGAACATCA | Syn6803 | | |
| 265 | nsii-in-cpcB-F | ctgccagcgatactctcgcgcagtctcagctgcatgctgg | cpcB-R | tcaaccagtctcctgttctc | Syn2973 | DNA seamless cloning | pCPF3-NSII-glgC |
| 266 | cpcB-glgc-F | gagaacaggagactggttgaatgtgttgttggcaatcgag | rbcl-glgc-R | CCGACAATCCAAACACCGGTctagattaccgtgccgtcgg | Syn6803 | | |
| 267 | NSII-in-R | cgcgagagtatcgctggcag | nsii-in-F | gcaatgcagatcctgtttaa | pCPF3-NSII-1 | | |
| 268 | rbcl-F | ACCGGTGTTTGGATTGTCGG | in-rbcl-R | ttaaacaggatctgcattgcGCTGTCGAAGTTGAACATCA | Syn6803 | | |
| 269 | cpcB-RF | gagaacaggagactggttga | rbcl-FR | CCGACAATCCAAACACCGGT | ptglgC | DNA seamless cloning | pCPF3-NSII-ptglgC |
| 270 | cpcB-R | tcaaccagtctcctgttctc | red-in-F | tgcgtttgatgacgatgttg | pCPF3-NSII-glgC | | |
| 271 | rbcl-F | ACCGGTGTTTGGATTGTCGG | red-in-R | caacatcgtcatcaaacgca | pCPF3- | | |

| | | | | | | | |
|---|---|---|---|---|---|---|---|
| | | | | | NSII-glgC | | |
| 272 | spe-cpcB-F | ACCAAGGTAGTCGGCAAATAAcagtctcagctgcatgctgg | rbcl-R | GCTGTCGAAGTTGAACATCA | pCPF3-NSII-cugP | DNA seamless cloning | pSES-cugP |
| 273 | spe-R | TTATTTGCCGACTACCTTGGT | rbcl-amp-F | TGATGTTCAACTTCGACAGCCGCGGAACCCCTATTTGTTT | pSES | | |
| 274 | spe-cpcB-F | ACCAAGGTAGTCGGCAAATAAcagtctcagctgcatgctgg | rbcl-R | GCTGTCGAAGTTGAACATCA | pCPF3-NSII-glgC | DNA seamless cloning | pSES-glgC |
| 275 | spe-R | TTATTTGCCGACTACCTTGGT | rbcl-amp-F | TGATGTTCAACTTCGACAGCCGCGGAACCCCTATTTGTTT | pSES | | |
| 276 | spe-cpcB-F | ACCAAGGTAGTCGGCAAATAAcagtctcagctgcatgctgg | rbcl-R | GCTGTCGAAGTTGAACATCA | pCPF3-NSII-ptglgC | DNA seamless cloning | pSES-ptglgC |
| 277 | spe-R | TTATTTGCCGACTACCTTGGT | rbcl-amp-F | TGATGTTCAACTTCGACAGCCGCGGAACCCCTATTTGTTT | pSES | | |
| 278 | cpf1-Gj-F | ATGGAAGCCGGCGGCACCTC | T0-R | aaaaaaatgaccttcataaatcgcta | pCPF3-NSIII-cscB | DNA seamless | pCPF3-NSIII-cscB |

| | | | | | | | |
|---|---|---|---|---|---|---|---|
| 279 | t0-nsiiiU-300-F | tttatgaaggtcatttttttcgtgggtcagccgtcagcgc | rbcl-nsiii-U-R | TGATGTTCAACTTCGACAGCatcgcgtgacgccagctaca | Syn2973 | cloning | -1200-300 |
| 280 | trbcl-R | GCTGTCGAAGTTGAACATCA | nsiii-d-R | atcacagtcggcgtcacggc | pCPF3-NSIII-cscB | | |
| 281 | nsiii-d-in-R | gccgtgacgccgactgtgatcgctcgcttcgtagtcgccg | amp-nsiii-d-R | AAACAAATAGGGGTTCCGCGatgacgacttcaccatcccg | Syn2973 | | |
| 282 | cpf1-gj-nsiii-U-R | GAGGTGCCGCCGGCTTCCATatcgcgtgacgccagctaca | amp-nsiii-U-1200-F | CACTGATTAAGCATTGGTAAccggaagtcgtagacagcga | Syn2973 | | |
| 283 | amp-R | TTACCAATGCTTAATCAGTG | amp-F | CGCGGAACCCCTATTTGTTT | pCPF3-NSIII-cscB | | |
| 284 | amp-nsiii-d-800-R | AAACAAATAGGGGTTCCGCGatcacagtcggcgtcacggc | cpf1-Gj-F | ATGGAAGCCGGCGGCACCTC | pCPF3-NSIII-cscB-1200-300 | DNA seamless cloning | pCPF3-NSIII-cscB-800-300 |

| | | | | | | | |
|---|---|---|---|---|---|---|---|
| 285 | amp-F | CGCGGAACCCCTATTTGTTT | amp-R | TTACCAATGCTTAATCAGTG | pCPF3-NSIII-cscB-1200-300 | | |
| 286 | amp-nsiii-U-800-F | CACTGATTAAGCATTGGTAAcgcgacaggttgcgctcaag | cpf1-gj-nsiii-U-R | GAGGTGCCGCCGGCTTCCATatcgcgtgacgccagctaca | pCPF3-NSIII-cscB-1200-300 | | |
| 287 | cpf1-gj-nsiii-U-R | GAGGTGCCGCCGGCTTCCATatcgcgtgacgccagctaca | amp-nsiii-U-500-F | CACTGATTAAGCATTGGTAAtagatgcggtgacgttcacc | pCPF3-NSIII-cscB-1200-300 | DNA seamless cloning | pCPF3-NSIII-cscB-500-300 |
| 288 | amp-nsiii-d-500-R | AAACAAATAGGGGTTCCGCGcaaaagacgcagttcgagga | cpf1-Gj-F | ATGGAAGCCGGCGGCACCTC | pCPF3-NSIII-cscB-1200-300 | | |
| 289 | amp | TTACCAATGCTTAATCAGTG | amp | CGCGGAACCCCTATTTGTTT | pCP | | |

| | | | | | | | |
|---|---|---|---|---|---|---|---|
| | -R | | -F | | F3-NSIII-cscB-1200-300 | | |
| 290 | T0-R | aaaaaaatgaccttcataaatcgcta | rbcl-R | GCTGTCGAAGTTGAACATCA | pCPF3-NSIII-cscB-1200-300 | DNA seamless cloning | pCPF3-NSIII-cscB-500-500 |
| 291 | T0-nsiii-F | tttatgaaggtcattttttttagatgcggtgacgttcacc | rbcl-nsiii-U-R | TGATGTTCAACTTCGACAGCatcgcgtgacgccagctaca | pCPF3-NSIII-cscB-1200-300 | | |
| 292 | trc2o-F | GAATTCCTTAGTTATTCCTATTCTGC | amp-F | CGCGGAACCCCTATTTGTTT | pBR3-NSIII-cscB | DNA seamless cloning | pBR3-NSIII-cscB-1200-300 |
| 293 | amp-nsiii-F | AAACAAATAGGGGTTCCGCGatgacgacttcaccatcccg | nsiii-in-R | gccgtgacgccgactgtgatcgctcgcttcgtagtcgccg | Syn2973 | | |
| 294 | nsiii | atcacagtcggcgtcacggc | nsiii | cgtgggtcagccgtcagcgc | pBR | | |

| | | | | | | | |
|---|---|---|---|---|---|---|---|
| | -d-F1 | | -u-300-R | | 3-NSIII-cscB | | |
| 295 | nsiii-u-km-R | gcgctgacggctgacccacgttagaaaaactcatcgagca | NSIII-un-in-R | cgacgacggtgctcggtgcc | pBR3-NSIII-cscB | | |
| 296 | nsiii-u-in-F | ggcaccgagcaccgtcgtcgacacgcttcccgatgtgctc | trc-nsiii-1200-R | ATAGGAATAACTAAGGAATTCccggaagtcgtagacagcga | Syn2973 | | |
| 297 | amp-nsiii-d-800-F | AAACAAATAGGGGTTCCGCGagcgatcacagtcggcgtca | trc-nsiii-u-800-R | TAGGAATAACTAAGGAATTCcgcgacaggttgcgctcaag | pBR3-NSIII-cscB-1200-300 | DNA seamless cloning | pBR3-NSIII-cscB-800-300 |
| 298 | amp-F | CGCGGAACCCCTATTTGTTT | trc2o-F | GAATTCCTTAGTTATTCCTATTCTGCACG | pBR3-NSIII-cscB-1200-300 | | |
| 299 | amp-nsiii | AAACAAATAGGGGTTCCGCGcaaaagacgcagttcgagga | trc-nsiii-u- | TAGGAATAACTAAGGAATTCtagatgcggtgacgttcacc | pBR3-NSII | DNA sea | pBR3-NSII |

| | | | | | | | |
|---|---|---|---|---|---|---|---|
| | -d-500-F | | 500-R | | I-cscB-1200-300 | mless cloning | I-cscB-500-300 |
| 300 | amp-F | CGCGGAACCCCTATTTGTTT | trc2o-F | GAATTCCTTAGTTATTCCTATTCTGCACG | pBR3-NSIII-cscB-1200-300 | | |
| 301 | trbcl-nsiii-U-500-F | TGATGTTCAACTTCGACAGCatcgcgtgacgccagctaca | km-nsiii-U-500-R | tgctcgatgagtttttctaatagatgcggtgacgttcacc | pBR3-NSIII-cscB-500-300 | DNA seamless cloning | pBR3-NSIII-cscB-500-500 |
| 302 | rbcl-R | GCTGTCGAAGTTGAACATCA | km-R | ttagaaaaactcatcgagcatc | pBR3-NSIII-cscB-500-300 | | |

**Table S2.** Sequences of key genes.

| Name | Sequences |
|---|---|
| km | ctgatccttcaactcagcaaaagttcgatttattcaacaaagccacgttgtgtctcaaaatctctgatgttacattgcacaagataaaaatatatcatcatgaacaataaaactgtctgcttacataaacagtaatacaaggggtgttatgagccatattcaacgggaaacgtcttgctccaggccgcgattaaattccaacatggatgctgatttatatgggtataaatgggctcgcgataatgtcgggcaatcaggtgcgacaatctatcgattgtatgggaagcccgatgcgccagagttgtttctgaaacatggcaaaggtagcgttgccaatgatgttacagatgagatggtcagactaaactggctgacggaatttatgcctcttccgaccatcaagcattttatccgtactcctgatgatgcatggttactcaccactgcgatccccgggaaaacagcattccaggtattagaagaatatcctgattcaggtgaaaatattgttgatgcgctggcagtgttcctgcgccggttgcattcgattcctgtttgtaattgtccttttaacagcgatcgcgtatttcgtctggctcaggcgcaatcacgaatgaataacggtttggttgatgcgagtgattttgatgacgagcgtaatggctggcctgttgaacaagtctggaaagaaatgcataagcttttgccattctcaccggattcagtcgtcactcatggtgatttctcacttgataaccttatttttgacgaggggaaattaataggttgtattgatgttggacgagtcggaatcgcagaccgataccaggatcttgccatcctatggaactgcctcggtgagttttctccttcattacagaaacggctttttcaaaaatatggtattgataatcctgatatgaataaattgcagtttcatttgatgctcgatgagtttttctaa |
| tho-sepT2 | GAATTCCTTAGTTATTCCTATTCTGCACGAACTCAAAATACTCTTCATTTTTGATAACCAAATTGAGTTTTTTGCCCTCTTGATTATTTTTGATCCTACCTAGTAGCCTCTTCTGCTCCTGCAGAATTGTGAGCGCTCACAATTGATATTTTGGTCTGTCGTTGCGATCGCCCGTTGCAGGCCGACATGAAGGATTGACAATTAATCATCCGGCTCGTATAATGAATTGTGAGCGCTCACAATTGGTACCGGTGATACCAGCATCGTCTTGATGCCCTTGGCAGCACCCTGCTAAGGAGGCAACAAGatgcaagcggtctttgataccaacatcctcatctaccacctcaaaggctgtcttcctgaagcggggagtcaaattctgcgcagcagtctggggcgcggtgccgtctgttcagtcattacccgcctagaggttttgggttacgaccaaccttggccagaaagactgaaagcacaagctttgctccagctatttcgagaacgggcattggatgaatcgattgctgattgcacgattcagctgcgtcaacaacaacggatcaaactgcccgatgccatcgttgccgcaactgcgctgacagagaacttgccactcgtgacgcgcaacaccaaagactttaaagccatcgctggcttacaactgattaacccctttcaaccgaactag |
| Erm | CGCACACCGTGGAAACGGATGAAGGCACGAACCCAGTGGACATAAGCCTGTTCGGTTCGTAAGCTGTAATGCAAGTAGCGTATGCGCTCACGCAACTGGTCCAGAACCTTGACCGAACGCAGCGGTGGTAACGGCGCAGTGGCGGTTTTCATGGCTTGTTATGACTGTTTTTTTGGGGTACAGTCTATGCCTCGGGCATCCAAGCAGCAAGCGCGTTACGCCGTGGGTCGATGTTTGATGTTATGGAGCAGCAACGATGTTACGCAGCAGGGCAGTCGCCCTAAAACAAAGTTAAACATCatgaacgagaaaaatataaaacacagtcaaaactttattacttcaaaacataatatagataaaataatgacaaatataagattaaatgaacatgataatatctttgaaatcggctcaggaaaaggccattttacccttgaattagtaaagaggtgtaatttcgtaactgccattgaaatagaccataaattatgcaaaactacagaaaataaacttgttgatcacgataatttccaagttttaaacaaggatatattgcagtttaaatttcctaaaaaccaatcctataaaatatatggtaatatacctataacataagtacggatataatacgcaaaattgtttttgatagtatagctaatgagatttatttaatcgtggaatacgggtttgctaaaagattattaaatacaaaacgctcattggcattacttttaatggcagaagttgatatttctatattaagtatggttccaagagaatattttcatcctaaacctaaagtgaatagctcacttatcagattaagtagaaaaaaatcaagaatatcacacaaagataaacaaaagtataattatttcgttatgaaatgggttaacaaagaatacaagaaaatatttacaaaaaatcaatttaacaattccttaaaacatgcaggaattgacgatttaaacaatattagctttgaacaattcttatctcttttcaatagctataaattatttaataagtaaTTGTTCAGAACGCTCGGTCTTGCACACCGGGCGTTTTTTCTTTGTGAGTCCA |
| crRNA-pilN | ttgacagctagctcagtcctaggtataatgctagcgctgatttaggcaaaaacgggtctaagaactttaaataatttctactgttgtagatGTTTCCGTGGAGCTACCGGATGgtctaagaactttaaataatttctactgttgtagattagcgatttatgaaggtcattttttt |
| trc2o-tho | gatattttggtctgtcgttgcgatcgcccgttgcaggccgacatgaaggattgacaattaatcatccggctcgtataatgaattgtgagcgctcacaattggtaccggtgataccAGCATCGTCTTGATGCCCTTGGCAGCACCCTGCTAAGGAGGCAACAAG |
| lac2o-tho | gatattttggtctgtcgttgcgatcgcccgttgcaggccgacatgaaggatttacactttatgcttccggctcgtatgttGTGTGGaattgtgagcgctcacaattggtaccggtgataccAGCATCGTCTTGATGCCCTTGGCAGCACCCTGCTAAGGAGGCAACAAG |
| crRNA-NSI-1 | ttgacagctagctcagtcctaggtataatgctagcgctgatttaggcaaaaacgggtctaagaactttaaataatttctactgttgtagatGGTAGCGCTGCCTTCCCCTTCGgtctaagaactttaaataatttctactgttgtagattagcgatttatgaaggtcattttttt |
| crR | ttgacagctagctcagtcctaggtataatgctagcgctgatttaggcaaaaacgggtctaagaactttaaataatttctactgttgtagatCATCTTCGACGATA |

| | |
|---|---|
| NA-NSI-2 | CGGGCGGCgtctaagaactttaaataatttctactgttgtagattagcgatttatgaaggtcattttttt |
| crRNA-NSII-1 | ttgacagctagctcagtcctaggtataatgctagcgctgatttaggcaaaaacgggtctaagaactttaaataatttctactgttgtagatCATCACGAAGCGGGTCACTACTgtctaagaactttaaataatttctactgttgtagattagcgatttatgaaggtcattttttt |
| crRNA-NSII-2 | ttgacagctagctcagtcctaggtataatgctagcgctgatttaggcaaaaacgggtctaagaactttaaataatttctactgttgtagatGCCTCTTGGTGCTGTTCAGTCTgtctaagaactttaaataatttctactgttgtagattagcgatttatgaaggtcattttttt |
| crRNA-NSIII-1 | ttgacagctagctcagtcctaggtataatgctagcgctgatttaggcaaaaacgggtctaagaactttaaataatttctactgttgtagatCTACGATTCCCGCGCCCACCATgtctaagaactttaaataatttctactgttgtagattagcgatttatgaaggtcattttttt |
| crRNA-NSIII-2 | ttgacagctagctcagtcctaggtataatgctagcgctgatttaggcaaaaacgggtctaagaactttaaataatttctactgttgtagatAACTGATCCTGCAGGACTTGATgtctaagaactttaaataatttctactgttgtagattagcgatttatgaaggtcattttttt |
| crRNA-13825-1 | ttgacagctagctcagtcctaggtataatgctagcgctgatttaggcaaaaacgggtctaagaactttaaataatttctactgttgtagatggcttttcgctgaactcgaccagtctaagaactttaaataatttctactgttgtagattagcgatttatgaaggtcattttttt |
| crRNA-13825-2 | ttgacagctagctcagtcctaggtataatgctagcgctgatttaggcaaaaacgggtctaagaactttaaataatttctactgttgtagatccattggcccgattctggtgccgtctaagaactttaaataatttctactgttgtagattagcgatttatgaaggtcattttttt |
| crRNA-06260-1 | ttgacagctagctcagtcctaggtataatgctagcgctgatttaggcaaaaacgggtctaagaactttaaataatttctactgttgtagatCGACAAAATCCGGCAGCGCTCTgtctaagaactttaaataatttctactgttgtagattagcgatttatgaaggtcattttttt |
| crRNA-06260- | ttgacagctagctcagtcctaggtataatgctagcgctgatttaggcaaaaacgggtctaagaactttaaataatttctactgttgtagatTCGCAAATCGGAATGGCGGGTCgtctaagaactttaaataatttctactgttgtagattagcgatttatgaaggtcattttttt |

| | |
|---|---|
| 2 | |
| crRNA-anl-1 | ttgacagctagctcagtcctaggtataatgctagcgctgatttaggcaaaaacgggtctaagaactttaaataatttctactgttgtagatTTGAACTCATAACAATCAGGCTgtctaagaactttaaataatttctactgttgtagattagcgatttatgaaggtcattttttt |
| crRNA-anl-2 | ttgacagctagctcagtcctaggtataatgctagcgctgatttaggcaaaaacgggtctaagaactttaaataatttctactgttgtagatCGTTCAATCGATTCCTCAGTTCgtctaagaactttaaataatttctactgttgtagattagcgatttatgaaggtcattttttt |
| crRNA-invA | ttgacagctagctcagtcctaggtataatgctagcgctgatttaggcaaaaacgggtctaagaactttaaataatttctactgttgtagattctccagagccaggcactgcaagtctaagaactttaaataatttctactgttgtagattagcgatttatgaaggtcattttttt |
| ptglgC | gagaacaggagactggttgaAGGAGGAATTAACCATGCAGTGGTGGTGGTGGTGCtgcccgccagggggaccgctggtttggcgcgctgtttggttagtggatagagacgactgcctgcgcctccaccgagaatgatcgccagcacgtttttcacGCACCACCACCACCACCACTGCATGGTTAATTCCTCCTACCGGTGTTTGGATTGTCGG |

**Table S3.** Primer sequences of qRT-PCR.

| Name | Sequences |
|---|---|
| 4-NSII-qRT-R2 | CCCGCTTCGTGATGCAAAAT |
| 4-NSIII-F1 | GCTGGGCTGAGCAAATCAAC |
| 4-NSIII-R1 | CAAGCCACGCATTCAACCAA |
| 4-NSIII-qRT-F2 | GGTCAACGATGAGGGACTGG |
| 4-NSIII-qRT-R2 | TGAAGGGCATTGACCACCTC |
| 4-13825-F1 | ggagcaataacaacggatttttg |
| 4-13825-R1 | gcttcacagcttgtgtggct |
| 4-13825-qRT-F2 | ACGTCTTCAAGCGGGATGTT |
| 4-13825-qRT-R2 | CTGAAGGGTGGGCTAGGTTG |
| 5-qRT-F1 | ttgaccggaccgtttatgat |
| 5-qRT-R1 | gagaccaaccagaagttgat |
| 5-qRT-F2 | GTTTACACCACCACCCCCAA |
| 5-qRT-R2 | TCTTTAACTCCGGTGGCGTC |
| 5-qRT-F3 | CCGCTTACCGGATACCTGTC |
| 5-qRT-R3 | ATCCTGTTACCAGTGGCTGC |
| 5-qRT-F4 | GGTCAACGATGAGGGACTGG |
| 5-qRT-F4 | TGAAGGGCATTGACCACCTC |
| 6-NSI-qRT-F | AAAAGCTCAAGCGGAAGGGA |
| 6-NSI-qRT-R | AGCAAGCTAGCGATTTGGGT |
| 6-sps-qRT-F | TACGTGGGCACCAGACTTTC |
| 6-sps-qRT-R | GGGTTTCTTCCTCCGCGTTA |
| 6-invA-qRT-F | TCACAGACGAAACTGGTCGG |
| 6-invA-qRT-R | GGGCAAAACTGACGTTGTCC |
| 6-NSII-qRT-F | CTGTTCAGTCTGGATGCGGT |
| 6-NSII-qRT-R | CCCGCTTCGTGATGCAAAAT |
| 6-GlgC-qRT-F | TCTACCGCATGGATTACGCC |
| 6-GlgC-qRT-R | CTCGGGTGCCTTTCTGTCAT |

**Table S4**. Backbone sequences of plasmids.

| PLASMID NAME | SEQUNENCE FROM 5' TO 3' |
|---|---|
| pBR322m | TTCTCATGTTTGACAGCTTATCATCGATAAGCTTTAATGCGGTAGTTTATCACAGTTAAATTGCTAACGC<br>AGTCAGGCACCGTGTATGAAATCTAACAATGCGCTCATCGTCATCCTCGGCACCGTCACCCTGGATGCT<br>GTAGGCATAGGCTTGGTTATGCCGGTACTGCCGGGCCTCTTGCGGGATATCGTCCATTCCGACAGCATC<br>GCCAGTCACTATGGCGTGCTGCTAGCGCTATATGCGTTGATGCAATTTCTATGCGCACCCGTTCTCGGAG<br>CACTGTCCGACCGCTTTGGCCGCCGCCCAGTCCTGCTCGCTTCGCTACTTGGAGCCACTATCGACTACG<br>CGATCATGGCGACCACACCCGTCCTGTGGATCCTCTACGCCGGACGCATCGTGGCCGGCATCACCGGC<br>GCCACAGGTGCGGTTGCTGGCGCCTATATCGCCGACATCACCGATGGGGAAGATCGGGCTCGCCACTT<br>CGGGCTCATGAGCGCTTGTTTCGGCGTGGGTATGGTGGCAGGCCCCGTGGCCGGGGGACTGTTGGGCG<br>CCATCTCCTTGCATGCACCATTCCTTGCGGCGGCGGTGCTCAACGGCCTCAACCTACTACTGGGCTGCT<br>TCCTAATGCAGGAGTCGCATAAGGGAGAGCGTCGACCGATGCCCTTGAGAGCCTTCAACCCAGTCAGC<br>TCCTTCCGGTGGGCGCGGGGCATGACTATCGTCGCCGCACTTATGACTGTCTTCTTTATCATGCAACTCG<br>TAGGACAGGTGCCGGCAGCGCTCTGGGTCATTTTCGGCGAGGACCGCTTTCGCTGGAGCGCGACGATG<br>ATCGGCCTGTCGCTTGCGGTATTCGGAATCTTGCACGCCCTCGCTCAAGCCTTCGTCACTGGTCCCGCC<br>ACCAAACGTTTCGGCGAGAAGCAGGCCATTATCGCCGGCATGGCGGCCGACGCGCTGGGCTACGTCTT<br>GCTGGCGTTCGCGACGCGAGGCTGGATGGCCTTCCCCATTATGATTCTTCTCGCTTCCGGCGGCATCGG<br>GATGCCCGCGTTGCAGGCCATGCTGTCCAGGCAGGTAGATGACGACCATCAGGGACAGCTTCAAGGAT<br>CGCTCGCGGCTCTTACCAGCCTAACTTCGATCATTGGACCGCTGATCGTCACGGCGATTTATGCCGCCT<br>CGGCGAGCACATGGAACGGGTTGGCATGGATTGTAGGCGCCGCCCTATACCTTGTCTGCCTCCCCGCGT<br>TGCGTCGCGGTGCATGGAGCCGGGCCACCTCGACCTGAATGGAAGCCGGCGGCACCTCGCTAACGGA<br>TTCACCACTCCAAGAATTGGAGCCAATCAATTCTTGCGGAGAACTGTGAATGCGCAAACCAACCCTTG<br>GCAGAACATATCCATCGCGTCCGCCATCTCCAGCAGCCGCACGCGGCGCATCTCGGGCAGCGTTGGGT<br>CCTGGCCACGGGTGCGCATGATCGTGCTCCTGTCGTTGAGGACCCGGCTAGGCTGGCGGGGTTGCCTT<br>ACTGGTTAGCAGAATGAATCACCGATACGCGAGCGAACGTGAAGCGACTGCTGCTGCAAAACGTCTG<br>CGACCTGAGCAACAACATGAATGGTCTTCGGTTTCCGTGTTTCGTAAAGTCTGGAAACGCGGAAGTCA<br>GCGCCCTGCACCATTATGTTCCGGATCTGCATCGCAGGATGCTGCTGGCTACCCTGTGGAACACCTACA<br>TCTGTATTAACGAAGCGCTGGCATTGACCCTGAGTGATTTTTCTCTGGTCCCGCCGCATCCATACCGCC<br>AGTTGTTTACCCTCACAACGTTCCAGTAACCGGGCATGTTCATCATCAGTAACCCGTATCGTGAGCATC<br>CTCTCTCGTTTCATCGGTATCATTACCCCCATGAACAGAAATCCCCCTTACACGGAGGCATCAGTGACC<br>AAACAGGAAAAAACCGCCCTTAACATGGCCCGCTTTATCAGAAGCCAGACATTAACGCTTCTGGAGAA<br>ACTCAACGAGCTGGACGCGGATGAACAGGCAGACATCTGTGAATCGCTTCACGACCACGCTGATGAG<br>CTTTACCGCAGCTGCCTCGCGCGTTTCGGTGATGACGGTGAAAACCTCTGACACATGCAGCTCCCGGG<br>ACGGTCACAGCTTGTCTGTAAGCGGATGCCGGGAGCAGACAAGCCCGTCAGGGCGCGTCAGCGGGTG<br>TTGGCGGGTGTCGGGGCGCAGCCATGACCCAGTCACGTAGCGATAGCGGAGTGTATACTGGCTTAACT<br>ATGCGGCATCAGAGCAGATTGTACTGAGAGTGCACCATATGCGGTGTGAAATACCGCACAGATGCGTA<br>AGGAGAAAATACCGCATCAGGCGCTCTTCCGCTTCCTCGCTCACTGACTCGCTGCGCTCGGTCGTTCG<br>GCTGCGGCGAGCGGTATCAGCTCACTCAAAGGCGGTAATACGGTTATCCACAGAATCAGGGGATAACG<br>CAGGAAAGAACATGTGAGCAAAAGGCCAGCAAAAGGCCAGGAACCGTAAAAAGGCCGCGTTGCTGG<br>CGTTTTTCCATAGGCTCCGCCCCCCTGACGAGCATCACAAAAATCGACGCTCAAGTCAGAGGTGGCGA<br>AACCCGACAGGACTATAAAGATACCAGGCGTTTCCCCCTGGAAGCTCCCTCGTGCGCTCTCCTGTTCC<br>GACCCTGCCGCTTACCGGATACCTGTCCGCCTTTCTCCCTTCGGGAAGCGTGGCGCTTTCTCATAGCTC |

| | |
|---|---|
| | ACGCTGTAGGTATCTCAGTTCGGTGTAGGTCGTTCGCTCCAAGCTGGGCTGTGTGCACGAACCCCCCG<br>TTCAGCCCGACCGCTGCGCCTTATCCGGTAACTATCGTCTTGAGTCCAACCCGGTAAGACACGACTTAT<br>CGCCACTGGCAGCAGCCACTGGTAACAGGATTAGCAGAGCGAGGTATGTAGGCGGTGCTACAGAGTTC<br>TTGAAGTGGTGGCCTAACTACGGCTACACTAGAAGGACAGTATTTGGTATCTGCGCTCTGCTGAAGCC<br>AGTTACCTTCGGAAAAAGAGTTGGTAGCTCTTGATCCGGCAAACAAACCACCGCTGGTAGCGGTGGTT<br>TTTTTGTTTGCAAGCAGCAGATTACGCGCAGAAAAAAAGGATCTCAAGAAGATCCTTTGATCTTTTCTA<br>CGGGGTCTGACGCTCAGTGGAACGAAAACTCACGTTAAGGGATTTTGGTCATGAGATTATCAAAAAGG<br>ATCTTCACCTAGATCCTTTTAAATTAAAAATGAAGTTTTAAATCAATCTAAAGTATATATGAGTAAACTTG<br>GTCTGACAGTTACCAATGCTTAATCAGTGAGGCACCTATCTCAGCGATCTGTCTATTTCGTTCATCCATA<br>GTTGCCTGACTCCCCGTCGTGTAGATAACTACGATACGGGAGGGCTTACCATCTGGCCCCAGTGCTGCA<br>ATGATACCGCGcGACCCACGCTCACCGGCTCCAGATTTATCAGCAATAAACCAGCCAGCCGGAAGGGC<br>CGAGCGCAGAAGTGGTCCTGCAACTTTATCCGCCTCCATCCAGTCTATTAATTGTTGCCGGGAAGCTAG<br>AGTAAGTAGTTCGCCAGTTAATAGTTTGCGCAACGTTGTTGCCATTGCTGCAGGCATCGTGGTGTCACG<br>CTCGTCGTTTGGTATGGCTTCATTCAGCTCCGGTTCCCAACGATCAAGGCGAGTTACATGATCCCCCAT<br>GTTGTGCAAAAAAGCGGTTAGCTCCTTCGGTCCTCCGATCGTTGTCAGAAGTAAGTTGGCCGCAGTGT<br>TATCACTCATGGTTATGGCAGCACTGCATAATTCTCTTACTGTCATGCCATCCGTAAGATGCTTTTCTGTG<br>ACTGGTGAGTACTCAACCAAGTCATTCTGAGAATAGTGTATGCGGCGACCGAGTTGCTCTTGCCCGGC<br>GTCAACACGGGATAATACCGCGCCACATAGCAGAACTTTAAAAGTGCTCATCATTGGAAAACGTTCTTC<br>GGGGCGAAAACTCTCAAGGATCTTACCGCTGTTGAGATCCAGTTCGATGTAACCCACTCGTGCACCCA<br>ACTGATCTTCAGCATCTTTTACTTTCACCAGCGTTTCTGGGTGAGCAAAAACAGGAAGGCAAAATGCC<br>GCAAAAAAGGGAATAAGGGCGACACGGAAATGTTGAATACTCATACTCTTCCTTTTTCAATATTATTGA<br>AGCATTTATCAGGGTTATTGTCTCATGAGCGGATACATATTTGAATGTATTTAGAAAAATAAACAAATAG<br>GGGTTCCGCGCACATTTCCCCGAAAAGTGCCACCTGACGTCTAAGAAACCATTATTATCATGACATTAA<br>CCTATAAAAATAGGCGTATCACGAGGCCCTTTCGTCTTCAAGAA |
| pS<br>ES<br>-<br>ori | TTATTTGCCGACTACCTTGGTGATCTCGCCTTTCACGTAGTGGACAAATTCTTCCAACTGATCTGCGCGC<br>GAGGCCAAGCGATCTTCTTCTTGTCCAAGATAAGCCTGTCTAGCTTCAAGTATGACGGGCTGATACTGG<br>GCCGGCAGGCGCTCCATTGCCCAGTCGGCAGCGACATCCTTCGGCGCGATTTTGCCGGTTACTGCGCT<br>GTACCAAATGCGGGACAACGTAAGCACTACATTTCGCTCATCGCCAGCCCAGTCGGGCGGCGAGTTCC<br>ATAGCGTTAAGGTTTCATTTAGCGCCTCAAATAGATCCTGTTCAGGAACCGGATCAAAGAGTTCCTCCG<br>CCGCTGGACCTACCAAGGCAACGCTATGTTCTCTTGCTTTTGTCAGCAAGATAGCCAGATCAATGTCGA<br>TCGTGGCTGGCTCGAAGATACCTGCAAGAATGTCATTGCGCTGCCATTCTCCAAATTGCAGTTCGCGCT<br>TAGCTGGATAACGCCACGGAATGATGTCGTCGTGCACAACAATGGTGACTTCTACAGCGCGGAGAATC<br>TCGCTCTCTCCAGGGGAAGCCGAAGTTTCCAAAAGGTCGTTGATCAAAGCTCGCCGCGTTGTTTCATC<br>AAGCCTTACGGTCACCGTAACCAGCAAATCAATATCACTGTGTGGCTTCAGGCCGCCATCCACTGCGG<br>AGCCGTACAAATGTACGGCCAGCAACGTCGGTTCGAGATGGCGCTCGATGACGCCAACTACCTCTGAT<br>AGTTGAGTCGATACTTCGGCGATCACCGCTTCCCTCATGATGTTTAACTTTGTTTTAGGGCGACTGCCCT<br>GCTGCGTAACATCGTTGCTGCTCCATAACATCAAACATCGACCCACGGCGTAACGCGCTTGCTGCTTGG<br>ATGCCCGAGGCATAGACTGTACCCCAAAAAAACAGTCATAACAAGCCATGAAAACCGCCACTGCGCC<br>GTTACCACCGCTGCGTTCGGTCAAGGTTCTGGACCAGTTGCGTGAGCGCATACGCTACTTGCATTACAG<br>CTTACGAACCGAACAGGCTTATGTCCACTGGGTTCGTGCCTTCATCCGTTTCCACGGTGTGCGccgagctgg<br>gcactgacagcctcagtgagcggctctcggaagcgctcgagcattggccacatgtgcttgggtgctgcgatggccagcgttccactatcgaagctgtaggcgcggca<br>gcggttagccgcaaaccatcgcaagccttcgttctcgatcgcggcgatcgcagttgtccaaagctgatcgagttcggcttgtgtcatgctgcggtctccgtagatgtagt<br>gctgacctgacctgacgcggtgtcacgaaaatgcgcagtgattggttttgatcttctattagggatttcgtgacagctccccagtccctgactagatcgggcgttttcaagc<br>caatgagcttcgatcgcctcaagcggcagttcagcaatcgcctcattcggcacgacttggtagatccaagtggacttgcccgcgatgcgcttctgccgtgccttgagctt<br>gaagccgatcatccgcagcagcgcccgcacgatggtgctggctctggtcttgtcggtgatctgaatcttgaggatctgtgcggcactagcccggtggcttgtggcggtc<br>gtgacgatctcttgaatcagcggatcatcaacgctgaaccagtcgcggcgttgcagcaactcttccaccttcagcccatgcaggtaccgcacttggggcgcgatacag<br>ctgcccacaagatcgggtccgaacatcgggcgatcgcgccgtttgtcgagcttggcggcatcgcgagcggcagcggcatccttgccaaggctgagccagaacagc<br>agtcttagcggggcgtatagaccgtcgtcatcagcggctagtagttcacgggtgcagtctccccagcgttgctggcagcggtgccgagcaagggcagccaactgttc<br>cgcgttagcgaagtccgcagacgcgatcgcctcagcttccttcggggtcagatcaggtgaactcaggatcgcctcggctgaggcatcagccagcgcttcagcagtctc |

ccgttgcagttcgtcaacgacctcttgaagaaagggatcgatcgcgccgtcccagtcttgccaagtgaacccgttgagccgtagaagctgctccaccgtggcggcata
ggtcaacccgtcacggttgcgacgggcggccaattctgcccatgcagtcaagtaggcttgcgatgcgaccggctcgccgatcttgatcgccgcctgtgacagctgag
ccaagatcacttgttggtgccgcttcaggccgtcggcaacctcggttgggtcgatgctaccgttgccagctcgcatcaaattggacgtgtcggggcaggtgatgaatcg
ggggacgttgctccgcacccgtcctgcaatctgcgcgatcgcacctggggcgatgtgcccagctccggcgaagatcagcacggcatcaaatgggcgatcgcctacc
ccttgggtgatcgacaagccagcatcgatcacgctcgtgaagacaaggtgggggcaacagttgatgaacatctgcgggttgagcttcagcgtttccaaagcccccgc
ttattggcttcgctgtcgatgaccggaaccggctgaccgcaggcttcggagagctgttgggcgatcgtcgctgcggaatacagggacggccgcgactggctcagcttc
tggcggtcgcagaagactgctaagcgcttgcccgcctgaaccatttcgaccgccatcgccatgagttcgcgatcgctgccaacctgatagaccggagcaccgctgaa
gggttggacggtactgctgaccagatatgcggttgatcctgtcagtgcttccaacagggcgatcgacacatccgacagcatcgcgtcagcggcgataacctgccgccc
ccctgcgagtagtccagcaatttcagccatgacttcggggcgatgtttagcgatctcggtcttgccgctcagcaggtgctcagcccacagttccacctcatcgatgacga
cgatcgcccgttgccatccgtgggcgctgaagcgcagcttgctggcggggtgcatggagtctaggcaaccgccaaggccgatcatccgaaggtcggaaccgggaa
cgcattcttcagcccacggcacccctaagcgcttgcccaaggccgcgcccaaggtacggcgatgcgtgatcaacacgacaggaatgccggtaccaatgagcggctg
gagatgccccgcgatcgcctctgtcttgcccgtccccatcgcagcattcaggcagacaaggcgatgctgttccggctgcgggatgtagtcgccgttgatgtacccgcc
ggtgtgtaagcggttagggctacgggtcagccgctgccactgcttcgcgacctcgcgctgatattcccgctgctcttgtcgagtcagctttggcgggcggtagccgcctt
ctttggccagcttggccagtgtggcgatcgtgatgccgctgcccttgaagctggcccacttccgctctagcttgcggttgccgtcgaacttggacgattgcgctgaccat
gcaatccagtcatctaacagggcatcgctgactgactgcagggccatcccgacagccacccagctctcgtagtcgtctgcgtagctggcaggtacatgggccagcat
ggtgcgggcgatcgcgacctcatcatcagaagctggagcactgacgatcggcttataggatttcggcgcggtctgccggtcttgccgctccattgcatccagcaattcg
ggcggggcctctgcaacctcgatctcccaaggtgcgcagccttccacccagcgatatgcgccggtctcagggtgttgaccaacgatgactgattgcagcttgttccaac
gcagctctaacagatcaccgtctgggctgtgaagcttctgcgtggcgatcgcagcccatcgctcttggggtacctgatagagccgttggtggcgtcctggcttgccaga
tgtatatgcaactgtgggcggtaattctgtcagcgtcagcccttctggcagtgcctcatgagccgttgggccatcgaagtcaactgcgattaacccaccgcttgcggggc
cagtaagcagaccgattccggtaacgcgccgatcgctgcggatacgttcagcaacagcagcatggtcaagcgggttattttgccagccagggtcaaatggtgccttgc
gactgttaacgacgcagaagcgccagtttgggggcagccggtcaagttgatcaattaattcgcagcgggtcgggatctgcgacctttgcagatgattggcgataaatgc
catacttctgttgagttatttttcctgacctcccctgggcaaagtttgccgacctggcctgggggggggttttgctttgattgccgataactatagctggaatgaatacggcag
gtggcaaggctgcctcgaattgatcggcaagccagaggcgcgatcgcccctggctgtgatgagacttagacgccggacgctgtcgagatccgaacgatcggacggt
gatcgactacgcagcagccgaagctgtgctcagccttggcgcggtaaaagcccgtctcgaactgcacctcgctagttagctgaggcgcgggcgtcaggcgtcccag
ctccagaggtaggttgccgcgtgcaactaggtagtagtagctgccgctcagccgtgcgtcgatcgcaacctccagcgaccaagggttgaactcatcgcggctgttgac
ttgcagatctgccaccaactgacggataggcagctcactctcaggcgacgcgaggatcagtccaggctcttgaCCACCTCGACCTGAATGGAAGC
CGGCGGCACCTCGCTAACGGATTCACCACTCCAAGAATTGGAGCCAATCAATTCTTGCGGAGAACTGT
GAATGCGCAAACCAACCCTTGGCAGAACATATCCATCGCGTCCGCCATCTCCAGCAGCCGCACGCGGC
GCATCTCGGGCAGCGTTGGGTCCTGGCCACGGGTGCGCATGATCGTGCTCCTGTCGTTGAGGACCCGG
CTAGGCTGGCGGGGTTGCCTTACTGGTTAGCAGAATGAATCACCGATACGCGAGCGAACGTGAAGCGA
CTGCTGCTGCAAAACGTCTGCGACCTGAGCAACAACATGAATGGTCTTCGGTTTCCGTGTTTCGTAAA
GTCTGGAAACGCGGAAGTCAGCGCCCTGCACCATTATGTTCCGGATCTGCATCGCAGGATGCTGCTGG
CTACCCTGTGGAACACCTACATCTGTATTAACGAAGCGCTGGCATTGACCCTGAGTGATTTTTCTCTGG
TCCCGCCGCATCCATACCGCCAGTTGTTTACCCTCACAACGTTCCAGTAACCGGGCATGTTCATCATCA
GTAACCCGTATCGTGAGCATCCTCTCTCGTTTCATCGGTATCATTACCCCCATGAACAGAAATCCCCCTT
ACACGGAGGCATCAGTGACCAAACAGGAAAAAACCGCCCTTAACATGGCCCGCTTTATCAGAAGCCA
GACATTAACGCTTCTGGAGAAACTCAACGAGCTGGACGCGGATGAACAGGCAGACATCTGTGAATCG
CTTCACGACCACGCTGATGAGCTTTACCGCAGCTGCCTCGCGCGTTTCGGTGATGACGGTGAAAACCT
CTGACACATGCAGCTCCCGGGACGGTCACAGCTTGTCTGTAAGCGGATGCCGGGAGCAGACAAGCCC
GTCAGGGCGCGTCAGCGGGTGTTGGCGGGTGTCGGGGCGCAGCCATGACCCAGTCACGTAGCGATAG
CGGAGTGTATACTGGCTTAACTATGCGGCATCAGAGCAGATTGTACTGAGAGTGCACCATATGCGGTGT
GAAATACCGCACAGATGCGTAAGGAGAAAATACCGCATCAGGCGCTCTTCCGCTTCCTCGCTCACTGA
CTCGCTGCGCTCGGTCGTTCGGCTGCGGCGAGCGGTATCAGCTCACTCAAAGGCGGTAATACGGTTAT
CCACAGAATCAGGGGATAACGCAGGAAAGAACATGTGAGCAAAAGGCCAGCAAAAGGCCAGGAACC
GTAAAAAGGCCGCGTTGCTGGCGTTTTTCCATAGGCTCCGCCCCCCTGACGAGCATCACAAAAATCGA
CGCTCAAGTCAGAGGTGGCGAAACCCGACAGGACTATAAAGATACCAGGCGTTTCCCCCTGGAAGCT
CCCTCGTGCGCTCTCCTGTTCCGACCCTGCCGCTTACCGGATACCTGTCCGCCTTTCTCCCTTCGGGAA
GCGTGGCGCTTTCTCATAGCTCACGCTGTAGGTATCTCAGTTCGGTGTAGGTCGTTCGCTCCAAGCTGG
GCTGTGTGCACGAACCCCCCGTTCAGCCCGACCGCTGCGCCTTATCCGGTAACTATCGTCTTGAGTCCA

| | |
|---|---|
| | ACCCGGTAAGACACGACTTATCGCCACTGGCAGCAGCCACTGGTAACAGGATTAGCAGAGCGAGGTAT<br>GTAGGCGGTGCTACAGAGTTCTTGAAGTGGTGGCCTAACTACGGCTACACTAGAAGGACAGTATTTGG<br>TATCTGCGCTCTGCTGAAGCCAGTTACCTTCGGAAAAAGAGTTGGTAGCTCTTGATCCGGCAAACAAA<br>CCACCGCTGGTAGCGGTGGTTTTTTTGTTTGCAAGCAGCAGATTACGCGCAGAAAAAAAGGATCTCAA<br>GAAGATCCTTTGATCTTTTCTACGGGGTCTGACGCTCAGTGGAACGAAAACTCACGTTAAGGGATTTTG<br>GTCATGAGATTATCAAAAAGGATCTTCACCTAGATCCTTTTAAATTAAAAATGAAGTTTTAAATCAATCT<br>AAAGTATATATGAGTAAACTTGGTCTGACAGTTACCAATGCTTAATCAGTGAGGCACCTATCTCAGCGAT<br>CTGTCTATTTCGTTCATCCATAGTTGCCTGACTCCCCGTCGTGTAGATAACTACGATACGGGAGGGCTTA<br>CCATCTGGCCCCAGTGCTGCAATGATACCGCGcGACCCACGCTCACCGGCTCCAGATTTATCAGCAATA<br>AACCAGCCAGCCGGAAGGGCCGAGCGCAGAAGTGGTCCTGCAACTTTATCCGCCTCCATCCAGTCTAT<br>TAATTGTTGCCGGGAAGCTAGAGTAAGTAGTTCGCCAGTTAATAGTTTGCGCAACGTTGTTGCCATTGC<br>TGCAGGCATCGTGGTGTCACGCTCGTCGTTTGGTATGGCTTCATTCAGCTCCGGTTCCCAACGATCAAG<br>GCGAGTTACATGATCCCCCATGTTGTGCAAAAAAGCGGTTAGCTCCTTCGGTCCTCCGATCGTTGTCAG<br>AAGTAAGTTGGCCGCAGTGTTATCACTCATGGTTATGGCAGCACTGCATAATTCTCTTACTGTCATGCCA<br>TCCGTAAGATGCTTTTCTGTGACTGGTGAGTACTCAACCAAGTCATTCTGAGAATAGTGTATGCGGCGA<br>CCGAGTTGCTCTTGCCCGGCGTCAACACGGGATAATACCGCGCCACATAGCAGAACTTTAAAAGTGCT<br>CATCATTGGAAAACGTTCTTCGGGGCGAAAACTCTCAAGGATCTTACCGCTGTTGAGATCCAGTTCGAT<br>GTAACCCACTCGTGCACCCAACTGATCTTCAGCATCTTTTACTTTCACCAGCGTTTCTGGGTGAGCAAA<br>AACAGGAAGGCAAAATGCCGCAAAAAAGGGAATAAGGGCGACACGGAAATGTTGAATACTCATACTC<br>TTCCTTTTTCAATATTATTGAAGCATTTATCAGGGTTATTGTCTCATGAGCGGATACATATTTGAATGTATT<br>TAGAAAAATAAACAAATAGGGGTTCCGCG |
| pS<br>ES | TTATTTGCCGACTACCTTGGTGATCTCGCCTTTCACGTAGTGGACAAATTCTTCCAACTGATCTGCGCGC<br>GAGGCCAAGCGATCTTCTTCTTGTCCAAGATAAGCCTGTCTAGCTTCAAGTATGACGGGCTGATACTGG<br>GCCGGCAGGCGCTCCATTGCCCAGTCGGCAGCGACATCCTTCGGCGCGATTTTGCCGGTTACTGCGCT<br>GTACCAAATGCGGGACAACGTAAGCACTACATTTCGCTCATCGCCAGCCCAGTCGGGCGGCGAGTTCC<br>ATAGCGTTAAGGTTTCATTTAGCGCCTCAAATAGATCCTGTTCAGGAACCGGATCAAAGAGTTCCTCCG<br>CCGCTGGACCTACCAAGGCAACGCTATGTTCTCTTGCTTTTGTCAGCAAGATAGCCAGATCAATGTCGA<br>TCGTGGCTGGCTCGAAGATACCTGCAAGAATGTCATTGCGCTGCCATTCTCCAAATTGCAGTTCGCGCT<br>TAGCTGGATAACGCCACGGAATGATGTCGTCGTGCACAACAATGGTGACTTCTACAGCGCGGAGAATC<br>TCGCTCTCTCCAGGGGAAGCCGAAGTTTCCAAAAGGTCGTTGATCAAAGCTCGCCGCGTTGTTTCATC<br>AAGCCTTACGGTCACCGTAACCAGCAAATCAATATCACTGTGTGGCTTCAGGCCGCCATCCACTGCGG<br>AGCCGTACAAATGTACGGCCAGCAACGTCGGTTCGAGATGGCGCTCGATGACGCCAACTACCTCTGAT<br>AGTTGAGTCGATACTTCGGCGATCACCGCTTCCCTCATGATGTTTAACTTTGTTTTAGGGCGACTGCCCT<br>GCTGCGTAACATCGTTGCTGCTCCATAACATCAAACATCGACCCACGGCGTAACGCGCTTGCTGCTTGG<br>ATGCCCGAGGCATAGACTGTACCCCAAAAAAACAGTCATAACAAGCCATGAAAACCGCCACTGCGCC<br>GTTACCACCGCTGCGTTCGGTCAAGGTTCTGGACCAGTTGCGTGAGCGCATACGCTACTTGCATTACAG<br>CTTACGAACCGAACAGGCTTATGTCCACTGGGTTCGTGCCTTCATCCGTTTCCACGGTGTGCGccgagctgg<br>gcactgacagcctcagtgagcggctctcggaagcgctcgagcattggccacatgtgcttgggtgctgcgatggccagcgttccactatcgaagctgtaggcgcggca<br>gcggttagccgcaaaccatcgcaagccttcgttctcgatcgcggcgatcgcagttgtccaaagctgatcgagttcggcttgtgtcatgctgcggtttccgtagatgtagtg<br>ctgacctgacctgacgcggtgtcacgaaaatgcgcagtgattggttttgatcttctattagggatttcgtgacagctccccagtccctgactagatcgggcgttttcaagcc<br>aatgagcttcgatcgcctcaagcggcagttcagcaatcgcctcattcggcacgacttggtagatccaagtggacttgcccgcgatgcgcttctgccgtgccttgagcttg<br>aagccgatcatccgcagcagcgcccgcacgatggtgctggctctggtcttgtcggtgatctgaatcttgaggatctgtgcggcactagcccggtggcttgtggcggtcg<br>tgacgatctcttgaatcagcggatcatcaacgctgaaccagtcgcggcgttgcagcaactcttccaccttcagcccatgcaggtaccgcacttggggcgcgatacagct<br>gcccacaagatcgggtccgaacatcgggcgatcgcgccgtttgtcgagcttggcggcatcgcgagcggcagcggcatccttgccaaggctgagccagaacagcag<br>tcttagcggggcgtatagaccgtcgtcatcagcggctagtagttcacgggtgcagtctccccagcgttgctggcagcggtgccgagcaagggcagccaactgttccg<br>cgttagcgaagtccgcagacgcgatcgcctcagcttccttcggggtcagatcaggtgaactcaggatcgcctcggctgaggcatcagccagcgcttcagcagtctccc<br>gttgcagttcgtcaacgacctcttgaagaaagggatcgatcgcgccgtcccagtcttgccaagtgaacccgttgagccgtagaagctgctccaccgtggcggcatagg<br>tcaacccgtcacggttgcgacgggcggccaattctgcccatgcagtcaagtaggcttgcgatgcgaccggctcgccgatcttgatcgccgcctgtgacagctgagcca<br>agatcacttgttggtgccgcttcaggccgtcggcaacctcggttgggtcgatgctaccgttgccagctcgcatcaaattggacgtgtcggggcaggtgatgaatcgggg |

gacgttgctccgcacccgtcctgcaatctgcgcgatcgcacctggggcgatgtgcccagctccggcgaagatcagcacggcatcaaatgggcgatcgcctacccctt
gggtgatcgacaagccagcatcgatcacgctcgtgaagacaaggtgggggcaacagttgatgaacatctgcgggttgagcttcagcgtttccaaagccccccgcttat
tggcttcgctgtcgatgaccggaaccggctgaccgcaggcttcggagagctgttgggcgatcgtcgctgcggaatacagggacggccgcgactggctcagcttctgg
cggtcgcagaagactgctaagcgcttgcccgcctgaaccatttcgaccgccatcgccatgagttcgcgatcgctgccaacctgatagaccggagcaccgctgaaggg
ttggacggtactgctgaccagatatgcggttgatcctgtcagtgcttccaacagggcgatcgacacatccgacagcatcgcgtcagcggcgataacctgccgcccccc
tgcgagtagtccagcaatttcagccatgacttcggggcgatgtttagcgatctcggtcttgccgctcagcaggtgctcagcccacagttccacctcatcgatgacgacga
tcgcccgttgccatccgtgggcgctgaagcgcagcttgctggcggggtgcatggagtctaggcaaccgccaaggccgatcatccgaaggtcggaaccgggaacgc
attcttcagcccacggcacccctaagcgcttgcccaaggccgcgcccaaggtacggcgatgcgtgatcaacacgacaggaatgccggtaccaatgagcggctgga
gatgccccgcgatcgcctctgtcttgcccgtccccatcgcagcattcaggcagacaaggcgatgctgttccggctgcgggatgtagtcgccgttgatgtacccgccggt
gtgtaagcggttagggctacgggtcagccgctgccactgcttcgcgacctcgcgctgatattcccgctgctcttgtcgagtcagctttggcgggcggtagccgccttcttt
ggccagcttggccagtgtggcgatcgtgatgccgctgcccttgaagctggcccacttccgctctagcttgcggttgccgtcgaacttggacgattgcgctgaccatgca
atccagtcatctaacagggcatcgctgactgactgcagggccatcccgacagccacccagctctcgtagtcgtctgcgtagctggcaggtacatgggccagcatggtg
cgggcgatcgcgacctcatcatcagaagctggagcactgacgatcggcttataggatttcggcgcggtctgccggtcttgccgctccattgcatccagcaattcgggcg
gggcctctgcaacctcgatctcccaaggtgcgcagccttccacccagcgatatgcgccggtttcagggtgttgaccaacgatgactgattgcagcttgttccaacgcag
ctctaacagatcaccgtctgggctgtgaagcttctgcgtggcgatcgcagcccatcgctcttggggtacctgatagagccgttggtggcgtcctggcttgccagatgtat
atgcaactgtgggcggtaattctgtcagcgtcagcccttctggcagtgcctcatgagccgttgggccatcgaagtcaactgcgattaacccaccgcttgcggggccagt
aagcagaccgattccggtaacgcgccgatcgctgcggatacgttcagcaacagcagcatggtcaagcgggttattttgccagccagggtcaaatggtgccttgcgact
gttaacgacgcagaagcgccagtttgggggcagccggtcaagttgatcaattaattcgcagcgggtcgggatctgcgacctttgcagatgattggcgataaatgccata
cttctgttgagttatttttcctgacctcccctgggcaaagtttgccgacctggcctggggggggttttgctttgattgccgataactatagctggaatgaatacggcaggtgg
caaggctgcctcgaattgatcggcaagccagaggcgcgatcgcccctggctgtgatgagacttagacgccggacgctgtcgagatccgaacgatcggacggtgatc
gactacgcagcagccgaagctgtgctcagccttggcgcggtaaaagcccgtttcgaactgcacctcgctagttagctgaggcgcgggcgtcaggcgtcccagctcca
gaggtaggttgccgcgtgcaactaggtagtagtagctgccgctcagccgtgcgtcgatcgcaacctccagcgaccaagggttgaactcatcgcggctgttgacttgca
gatctgccaccaactgacggataggcagctcactctcaggcgacgcgaggatcagtccaggctcttgaCCACCTCGACCTGAATGGAAGCCG
GCGGCACCTCGCTAACGGATTCACCACTCCAAGAATTGGAGCCAATCAATTCTTGCGGAGAACTGTGA
ATGCGCAAACCAACCCTTGGCAGAACATATCCATCGCGTCCGCCATCTCCAGCAGCCGCACGCGGCGC
ATCTCGGGCAGCGTTGGGTCCTGGCCACGGGTGCGCATGATCGTGCTCCTGTCGTTGAGGACCCGGCT
AGGCTGGCGGGGTTGCCTTACTGGTTAGCAGAATGAATCACCGATACGCGAGCGAACGTGAAGCGACT
GCTGCTGCAAAACGTCTGCGACCTGAGCAACAACATGAATGGTCTTCGGTTTCCGTGTTTCGTAAAGT
CTGGAAACGCGGAAGTCAGCGCCCTGCACCATTATGTTCCGGATCTGCATCGCAGGATGCTGCTGGCT
ACCCTGTGGAACACCTACATCTGTATTAACGAAGCGCTGGCATTGACCCTGAGTGATTTTTCTCTGGTC
CCGCCGCATCCATACCGCCAGTTGTTTACCCTCACAACGTTCCAGTAACCGGGCATGTTCATCATCAGT
AACCCGTATCGTGAGCATCCTCTCTCGTTTCATCGGTATCATTACCCCCATGAACAGAAATCCCCCTTAC
ACGGAGGCATCAGTGACCAAACAGGAAAAAACCGCCCTTAACATGGCCCGCTTTATCAGAAGCCAGA
CATTAACGCTTCTGGAGAAACTCAACGAGCTGGACGCGGATGAACAGGCAGACATCTGTGAATCGCTT
CACGACCACGCTGATGAGCTTTACCGCAGCTGCCTCGCGCGTTTCGGTGATGACGGTGAAAACCTCTG
ACACATGCAGCTCCCGGGACGGTCACAGCTTGTCTGTAAGCGGATGCCGGGAGCAGACAAGCCCGTC
AGGGCGCGTCAGCGGGTGTTGGCGGGTGTCGGGGCGCAGCCATGACCCAGTCACGTAGCGATAGCGG
AGTGTATACTGGCTTAACTATGCGGCATCAGAGCAGATTGTACTGAGAGTGCACCATATGCGGTGTGAA
ATACCGCACAGATGCGTAAGGAGAAAATACCGCATCAGGCGCTCTTCCGCTTCCTCGCTCACTGACTC
GCTGCGCTCGGTCGTTCGGCTGCGGCGAGCGGTATCAGCTCACTCAAAGGCGGTAATACGGTTATCCA
CAGAATCAGGGGATAACGCAGGAAAGAACATGTGAGCAAAAGGCCAGCAAAAGGCCAGGAACCGTA
AAAAGGCCGCGTTGCTGGCGTTTTTCCATAGGCTCCGCCCCCCTGACGAGCATCACAAAAATCGACGC
TCAAGTCAGAGGTGGCGAAACCCGACAGGACTATAAAGATACCAGGCGTTTCCCCCTGGAAGCTCCCT
CGTGCGCTCTCCTGTTCCGACCCTGCCGCTTACCGGATACCTGTCCGCCTTTCTCCCTTCGGGAAGCGT
GGCGCTTTCTCATAGCTCACGCTGTAGGTATCTCAGTTCGGTGTAGGTCGTTCGCTCCAAGCTGGGCTG
TGTGCACGAACCCCCCGTTCAGCCCGACCGCTGCGCCTTATCCGGTAACTATCGTCTTGAGTCCAACCC
GGTAAGACACGACTTATCGCCACTGGCAGCAGCCACTGGTAACAGGATTAGCAGAGCGAGGTATGTAG
GCGGTGCTACAGAGTTCTTGAAGTGGTGGCCTAACTACGGCTACACTAGAAGGACAGTATTTGGTATCT
GCGCTCTGCTGAAGCCAGTTACCTTCGGAAAAAGAGTTGGTAGCTCTTGATCCGGCAAACAAACCACC

| | |
|---|---|
| | GCTGGTAGCGGTGGTTTTTTTGTTTGCAAGCAGCAGATTACGCGCAGAAAAAAAGGATCTCAAGAAGA<br>TCCTTTGATCTTTTCTACGGGGTCTGACGCTCAGTGGAACGAAAACTCACGTTAAGGGATTTTGGTCAT<br>GAGATTATCAAAAAGGATCTTCACCTAGATCCTTTTAAATTAAAAATGAAGTTTTAAATCAATCTAAAGT<br>ATATATGAGTAAACTTGGTCTGACAGTTACCAATGCTTAATCAGTGAGGCACCTATCTCAGCGATCTGTC<br>TATTTCGTTCATCCATAGTTGCCTGACTCCCCGTCGTGTAGATAACTACGATACGGGAGGGCTTACCATC<br>TGGCCCCAGTGCTGCAATGATACCGCGcGACCCACGCTCACCGGCTCCAGATTTATCAGCAATAAACCA<br>GCCAGCCGGAAGGGCCGAGCGCAGAAGTGGTCCTGCAACTTTATCCGCCTCCATCCAGTCTATTAATT<br>GTTGCCGGGAAGCTAGAGTAAGTAGTTCGCCAGTTAATAGTTTGCGCAACGTTGTTGCCATTGCTGCAG<br>GCATCGTGGTGTCACGCTCGTCGTTTGGTATGGCTTCATTCAGCTCCGGTTCCCAACGATCAAGGCGAG<br>TTACATGATCCCCCATGTTGTGCAAAAAAGCGGTTAGCTCCTTCGGTCCTCCGATCGTTGTCAGAAGTA<br>AGTTGGCCGCAGTGTTATCACTCATGGTTATGGCAGCACTGCATAATTCTCTTACTGTCATGCCATCCGT<br>AAGATGCTTTTCTGTGACTGGTGAGTACTCAACCAAGTCATTCTGAGAATAGTGTATGCGGCGACCGA<br>GTTGCTCTTGCCCGGCGTCAACACGGGATAATACCGCGCCACATAGCAGAACTTTAAAAGTGCTCATC<br>ATTGGAAAACGTTCTTCGGGGCGAAAACTCTCAAGGATCTTACCGCTGTTGAGATCCAGTTCGATGTA<br>ACCCACTCGTGCACCCAACTGATCTTCAGCATCTTTTACTTTCACCAGCGTTTCTGGGTGAGCAAAAAC<br>AGGAAGGCAAAATGCCGCAAAAAAGGGAATAAGGGCGACACGGAAATGTTGAATACTCATACTCTTC<br>CTTTTTCAATATTATTGAAGCATTTATCAGGGTTATTGTCTCATGAGCGGATACATATTTGAATGTATTTAG<br>AAAAATAAACAAATAGGGGTTCCGCG |
| pSEL-ori | TTATTTGCCGACTACCTTGGTGATCTCGCCTTTCACGTAGTGGACAAATTCTTCCAACTGATCTGCGCGC<br>GAGGCCAAGCGATCTTCTTCTTGTCCAAGATAAGCCTGTCTAGCTTCAAGTATGACGGGCTGATACTGG<br>GCCGGCAGGCGCTCCATTGCCCAGTCGGCAGCGACATCCTTCGGCGCGATTTTGCCGGTTACTGCGCT<br>GTACCAAATGCGGGACAACGTAAGCACTACATTTCGCTCATCGCCAGCCCAGTCGGGCGGCGAGTTCC<br>ATAGCGTTAAGGTTTCATTTAGCGCCTCAAATAGATCCTGTTCAGGAACCGGATCAAAGAGTTCCTCCG<br>CCGCTGGACCTACCAAGGCAACGCTATGTTCTCTTGCTTTTGTCAGCAAGATAGCCAGATCAATGTCGA<br>TCGTGGCTGGCTCGAAGATACCTGCAAGAATGTCATTGCGCTGCCATTCTCCAAATTGCAGTTCGCGCT<br>TAGCTGGATAACGCCACGGAATGATGTCGTCGTGCACAACAATGGTGACTTCTACAGCGCGGAGAATC<br>TCGCTCTCTCCAGGGGAAGCCGAAGTTTCCAAAAGGTCGTTGATCAAAGCTCGCCGCGTTGTTTCATC<br>AAGCCTTACGGTCACCGTAACCAGCAAATCAATATCACTGTGTGGCTTCAGGCCGCCATCCACTGCGG<br>AGCCGTACAAATGTACGGCCAGCAACGTCGGTTCGAGATGGCGCTCGATGACGCCAACTACCTCTGAT<br>AGTTGAGTCGATACTTCGGCGATCACCGCTTCCCTCATGATGTTTAACTTTGTTTTAGGGCGACTGCCCT<br>GCTGCGTAACATCGTTGCTGCTCCATAACATCAAACATCGACCCACGGCGTAACGCGCTTGCTGCTTGG<br>ATGCCCGAGGCATAGACTGTACCCCAAAAAAACAGTCATAACAAGCCATGAAAACCGCCACTGCGCC<br>GTTACCACCGCTGCGTTCGGTCAAGGTTCTGGACCAGTTGCGTGAGCGCATACGCTACTTGCATTACAG<br>CTTACGAACCGAACAGGCTTATGTCCACTGGGTTCGTGCCTTCATCCGTTTCCACGGTGTGCGccatgatcga<br>gcgatcggcctcgggggcgttgcagttcatccgcatgatctcgcccctgtcttggatgcccattgccgttatggcctttggcattggcgatctgcccgtttattttctgctag<br>cgatcgcagcggtctggccaattctactcagcaccagcagcggtactgcggctgttaaccacaagctcttacttctagctcgtagcctttgtgccactcgcagtgaaacg<br>attcggcgcatcgtgattcctgcgatcgtgccgcaaatccttgtgggtgttcgactcgcgatcggcacgatctggattgtgctcgtccccgctgagatgctgggcgttagc<br>tcaggcctcggttacttcattcttgatacccgcgatcgcattgcctataacgaacttaccgctgttttattagcgatcggcatcattggctgcgccctagattggagcctgca<br>attcctgcaaaagtactggcaacccagtcgctaaacctcttggcatccattctccaaacctctcaaaggcgtctacagggttgagcttgtggggaggggcggctatcctc<br>gcccgccaaaagcccgatcgtcatcactggcaagcccagcaacgacagatggaactgaactcagtccaagggcagctcgatgccctggtcgtcaaagcaccctacg<br>caggcagggttaggcgggtacgctggctagagcaggtgggcggacaggtcaaggtggagattgctcttcaagtctcaaacactccctaagttttgaccttgattacagt<br>ggccccatctagtttttgtatatacaatgactaggtgcgtatcacctacgatccagtcaaacgcgacaagacactgcttgagcgaggactcgacttcgagagtgcgatcg<br>aagtctttgcagggctcacgctagaagtcgaagacacccgacgcgactatggcgaaaagcgagtgctctgcgtcggcttccttgatggccgcatggtgatggttggct<br>ataccccctcgcggtcgaacacgtcacatcttttcgatgaggaaatgcaatgcccgagaaattcgccgctacacgccatacttcgagcaagcctgatgacgacttgcccg<br>agttgaccgatgaaatgctcagtcgggcggtgctcaagcagggcggtaagcgtatcggcagaccccgatcgcaatctccaaaagtagcgatcagcctgcgcttagaa<br>gctgaagtcctgcagcgctggcgtgacagtggccctggctggcaaactcgtatggctgaattgctctcgaaagaaaccccctaagtctttgatcgctcaagcccaacgg<br>caggacaagctacaccagtggcaaacccagcagcgacagatggaacttaactcagtccaagcgcagctcgatgcactggtcgtcaaagcaccttactccggcagag<br>tcaggcgggtgcgttggctagagcaagtcggtggacaggtgaaagtagagattgctcttcagatctttaacgagtgataaattcatagctttgtcattatctattatcgaga<br>gcaaataagagctatcaaggagcacctaagtgatctgctacttggtaaccttttttcagttagtctataaacttctcctttgccgccaagtgactccattagctcatatttaataa |

gctctgaaatagcacttttccaagaggcctcactacgagcattacccatatctgcaagctgcttattattagtttcaatcatcaatccacccataactggcagacaaattaatt
ctccattagaatctgatgctacctctattaacagatcttttccaagatcgcttaagcgtggtaattcaggagcaaaagatacttctaaagtattattagtgctttctaagctactg
taattttgtataattttctgagatatctgccgactaaacttatgctgaaactcctccaatgactcatatgtttcaattaagccttttttgttgacacaaactacgaaaatcacatagtg
ccttgtattgatcctgatcaacgctttctagatatactggagctctggaaaagtaaatcatcgcttgcttgccagacttaagatgttcttctatttcctcaacagttccactaata
aatttaccagtaggtgttcctaatcttgtccaaaaaacagcaacaagtagatcgcagtctttcaaaatttgcttgtttattatggattgtgctcgtccgctcatatcaggatggg
catgtgtttcccaccctactggatcaagtactattttacgatcaatagaattaatggtattccactcgtatattatgttccgaataatgttgcgttcttttgcaacatcgctcggtg
atgcaatcataatcctaagaactgtagcgttgtatggcattgctagccccttagagaaaatttcagtagcatttactatttattagctatatttacataaattttttaaactcatcag
agataatgactcatattttctatacgatcgcaagtaacttcactgccccttcaccacctccgtatcgcccgagaaagcaaagtgctccgcctgactcaccagcgactctg
gcaacggcagcgtaatccgatagcactccagaaactcacccagcactcccatcgcatgggctttgaccgaatccgagaaccgatcgccctctgataggaacactgag
atcagcttttctcccgccgaggccaagggctgattgagaccatccagcaggaaactataggttgggcgactgaagtagcttttctctagctcctctgccgatagctcatgc
agcatttcatagagagcatcagtcgcagcttccgtcccgatccgcttcagggtgtgggcaacccagcagcgcacttccatgtctggatcggccagcaggggcttgagg
cgagcgatcagctcggcgctctgggctgcttccgtgtccttcgcaaggcgcttgtagagcgcatgggcaacgctgccacggatctccggctccatctccagcagttgc
agcaggatcgacaccgattgattcatgacccgaacaccccaaggcatcaagacagccttcagcgttccctctgggtagatcaagctaacttctgcatcctagatgagta
caaataaccaactattgctatgagggagcataggccagcttttatgtctccttaaggatcaaatcaacataaatgagtctcgaaaggctgcctctgtcttctctatgggacaa
tgtaagccagaacactaatagttacttttgaatttaaagtcagtaacaattacacagtgaaaccgagaaaataaaaagggaatctaaacttgagagctcagttaggtgctct
tcgctaaactcctcttaaccttcactggcctcggccaaggttcgccaaaccgcccagatcaggcgatactatcctgcaaccccagctgtcaaggagctgcctagcgtcc
attgctctccctcccctcagtgcctcgaccgcaccccttagccttgggtgtgcactgaccacaatcgattgaactgagtccggtgtttgagctgctgtttggcgacggcag
atttcagacccctaaaaagttcgaagccccgagttggcgctcagggcttcaggccgggtgtgtcctagcctcgggggacaaaatcttactcccgatgctaccacctcgc
gccgcagaccaacgtgtctatgcaccggcaggtgtgaggaaaatatgaaacttttcgtccgaggaggccgctgatggtctcctctatccctgtatctggcagcctcgaa
acccagctccagcaagagttctgccaatccagcgcgatcgctttgagcctctttcagcgggcgatcgcccttgtgccagatcttgaaattgatccagtcactcacgaggt
gatcggtacgccgatcgcagacgccttgggctggcagttcacccgctttgggctgcaagcccgcgctaaccagaccgccgcgatttttgtacaggaatcaaacgaggt
ctggcaggccaagattttcggtggtgagtccggtaagcgaagcggctcctacttagctcccaagggcattggcaaccgcgcttatctcccgcccgttgacccagaaac
ccgttctcagcttggtgctcctgctgaaggaagtttctgggactgggttgaatcaactccttctatccccatcgtgccgaccgaaggcggcaagaagtctctctctgctctt
agtgctggctccgtagcgatcgccctctatggctgtgattgcggcaatagtcctgacctgactcgcttcttagtcccaggtcgtgaagtaatcatcgccttcgaccaagac
gctaaggccaagaccagagccagagtgaatcgcagcattgcacggctggcacggcgagcgatcgcggctgggtgctcggtccgcattgctttatggCCACCT
CGACCTGAATGGAAGCCGGCGGCACCTCGCTAACGGATTCACCACTCCAAGAATTGGAGCCAATCAAT
TCTTGCGGAGAACTGTGAATGCGCAAACCAACCCTTGGCAGAACATATCCATCGCGTCCGCCATCTCC
AGCAGCCGCACGCGGCGCATCTCGGGCAGCGTTGGGTCCTGGCCACGGGTGCGCATGATCGTGCTCCT
GTCGTTGAGGACCCGGCTAGGCTGGCGGGGTTGCCTTACTGGTTAGCAGAATGAATCACCGATACGCG
AGCGAACGTGAAGCGACTGCTGCTGCAAAACGTCTGCGACCTGAGCAACAACATGAATGGTCTTCGG
TTTCCGTGTTTCGTAAAGTCTGGAAACGCGGAAGTCAGCGCCCTGCACCATTATGTTCCGGATCTGCAT
CGCAGGATGCTGCTGGCTACCCTGTGGAACACCTACATCTGTATTAACGAAGCGCTGGCATTGACCCTG
AGTGATTTTTCTCTGGTCCCGCCGCATCCATACCGCCAGTTGTTTACCCTCACAACGTTCCAGTAACCG
GGCATGTTCATCATCAGTAACCCGTATCGTGAGCATCCTCTCTCGTTTCATCGGTATCATTACCCCCATG
AACAGAAATCCCCCTTACACGGAGGCATCAGTGACCAAACAGGAAAAAACCGCCCTTAACATGGCCC
GCTTTATCAGAAGCCAGACATTAACGCTTCTGGAGAAACTCAACGAGCTGGACGCGGATGAACAGGC
AGACATCTGTGAATCGCTTCACGACCACGCTGATGAGCTTTACCGCAGCTGCCTCGCGCGTTTCGGTG
ATGACGGTGAAAACCTCTGACACATGCAGCTCCCGGGACGGTCACAGCTTGTCTGTAAGCGGATGCCG
GGAGCAGACAAGCCCGTCAGGGCGCGTCAGCGGGTGTTGGCGGGTGTCGGGGCGCAGCCATGACCC
AGTCACGTAGCGATAGCGGAGTGTATACTGGCTTAACTATGCGGCATCAGAGCAGATTGTACTGAGAGT
GCACCATATGCGGTGTGAAATACCGCACAGATGCGTAAGGAGAAAATACCGCATCAGGCGCTCTTCCG
CTTCCTCGCTCACTGACTCGCTGCGCTCGGTCGTTCGGCTGCGGCGAGCGGTATCAGCTCACTCAAAG
GCGGTAATACGGTTATCCACAGAATCAGGGGATAACGCAGGAAAGAACATGTGAGCAAAAGGCCAGC
AAAAGGCCAGGAACCGTAAAAAGGCCGCGTTGCTGGCGTTTTTCCATAGGCTCCGCCCCCCTGACGA
GCATCACAAAAATCGACGCTCAAGTCAGAGGTGGCGAAACCCGACAGGACTATAAAGATACCAGGCG
TTTCCCCCTGGAAGCTCCCTCGTGCGCTCTCCTGTTCCGACCCTGCCGCTTACCGGATACCTGTCCGCC
TTTCTCCCTTCGGGAAGCGTGGCGCTTTCTCATAGCTCACGCTGTAGGTATCTCAGTTCGGTGTAGGTC
GTTCGCTCCAAGCTGGGCTGTGTGCACGAACCCCCCGTTCAGCCCGACCGCTGCGCCTTATCCGGTAA
CTATCGTCTTGAGTCCAACCCGGTAAGACACGACTTATCGCCACTGGCAGCAGCCACTGGTAACAGGA

| | |
|---|---|
| | TTAGCAGAGCGAGGTATGTAGGCGGTGCTACAGAGTTCTTGAAGTGGTGGCCTAACTACGGCTACACT<br>AGAAGGACAGTATTTGGTATCTGCGCTCTGCTGAAGCCAGTTACCTTCGGAAAAAGAGTTGGTAGCTC<br>TTGATCCGGCAAACAAACCACCGCTGGTAGCGGTGGTTTTTTTGTTTGCAAGCAGCAGATTACGCGCA<br>GAAAAAAAGGATCTCAAGAAGATCCTTTGATCTTTTCTACGGGGTCTGACGCTCAGTGGAACGAAAAC<br>TCACGTTAAGGGATTTTGGTCATGAGATTATCAAAAAGGATCTTCACCTAGATCCTTTTAAATTAAAAAT<br>GAAGTTTTAAATCAATCTAAAGTATATATGAGTAAACTTGGTCTGACAGTTACCAATGCTTAATCAGTGA<br>GGCACCTATCTCAGCGATCTGTCTATTTCGTTCATCCATAGTTGCCTGACTCCCCGTCGTGTAGATAACT<br>ACGATACGGGAGGGCTTACCATCTGGCCCCAGTGCTGCAATGATACCGCGcGACCCACGCTCACCGGC<br>TCCAGATTTATCAGCAATAAACCAGCCAGCCGGAAGGGCCGAGCGCAGAAGTGGTCCTGCAACTTTAT<br>CCGCCTCCATCCAGTCTATTAATTGTTGCCGGGAAGCTAGAGTAAGTAGTTCGCCAGTTAATAGTTTGC<br>GCAACGTTGTTGCCATTGCTGCAGGCATCGTGGTGTCACGCTCGTCGTTTGGTATGGCTTCATTCAGCT<br>CCGGTTCCCAACGATCAAGGCGAGTTACATGATCCCCCATGTTGTGCAAAAAAGCGGTTAGCTCCTTC<br>GGTCCTCCGATCGTTGTCAGAAGTAAGTTGGCCGCAGTGTTATCACTCATGGTTATGGCAGCACTGCAT<br>AATTCTCTTACTGTCATGCCATCCGTAAGATGCTTTTCTGTGACTGGTGAGTACTCAACCAAGTCATTCT<br>GAGAATAGTGTATGCGGCGACCGAGTTGCTCTTGCCCGGCGTCAACACGGGATAATACCGCGCCACAT<br>AGCAGAACTTTAAAAGTGCTCATCATTGGAAAACGTTCTTCGGGGCGAAAACTCTCAAGGATCTTACC<br>GCTGTTGAGATCCAGTTCGATGTAACCCACTCGTGCACCCAACTGATCTTCAGCATCTTTTACTTTCACC<br>AGCGTTTCTGGGTGAGCAAAAACAGGAAGGCAAAATGCCGCAAAAAAGGGAATAAGGGCGACACGG<br>AAATGTTGAATACTCATACTCTTCCTTTTTCAATATTATTGAAGCATTTATCAGGGTTATTGTCTCATGAG<br>CGGATACATATTTGAATGTATTTAGAAAAATAAACAAATAGGGGTTCCGCG |
| pS<br>EL | TTATTTGCCGACTACCTTGGTGATCTCGCCTTTCACGTAGTGGACAAATTCTTCCAACTGATCTGCGCGC<br>GAGGCCAAGCGATCTTCTTCTTGTCCAAGATAAGCCTGTCTAGCTTCAAGTATGACGGGCTGATACTGG<br>GCCGGCAGGCGCTCCATTGCCCAGTCGGCAGCGACATCCTTCGGCGCGATTTTGCCGGTTACTGCGCT<br>GTACCAAATGCGGGACAACGTAAGCACTACATTTCGCTCATCGCCAGCCCAGTCGGGCGGCGAGTTCC<br>ATAGCGTTAAGGTTTCATTTAGCGCCTCAAATAGATCCTGTTCAGGAACCGGATCAAAGAGTTCCTCCG<br>CCGCTGGACCTACCAAGGCAACGCTATGTTCTCTTGCTTTTGTCAGCAAGATAGCCAGATCAATGTCGA<br>TCGTGGCTGGCTCGAAGATACCTGCAAGAATGTCATTGCGCTGCCATTCTCCAAATTGCAGTTCGCGCT<br>TAGCTGGATAACGCCACGGAATGATGTCGTCGTGCACAACAATGGTGACTTCTACAGCGCGGAGAATC<br>TCGCTCTCTCCAGGGGAAGCCGAAGTTTCCAAAAGGTCGTTGATCAAAGCTCGCCGCGTTGTTTCATC<br>AAGCCTTACGGTCACCGTAACCAGCAAATCAATATCACTGTGTGGCTTCAGGCCGCCATCCACTGCGG<br>AGCCGTACAAATGTACGGCCAGCAACGTCGGTTCGAGATGGCGCTCGATGACGCCAACTACCTCTGAT<br>AGTTGAGTCGATACTTCGGCGATCACCGCTTCCCTCATGATGTTTAACTTTGTTTTAGGGCGACTGCCCT<br>GCTGCGTAACATCGTTGCTGCTCCATAACATCAAACATCGACCCACGGCGTAACGCGCTTGCTGCTTGG<br>ATGCCCGAGGCATAGACTGTACCCCAAAAAAACAGTCATAACAAGCCATGAAAACCGCCACTGCGCC<br>GTTACCACCGCTGCGTTCGGTCAAGGTTCTGGACCAGTTGCGTGAGCGCATACGCTACTTGCATTACAG<br>CTTACGAACCGAACAGGCTTATGTCCACTGGGTTCGTGCCTTCATCCGTTTCCACGGTGTGCGccatgatcga<br>gcgatcggcctcgggggcgttgcagttcatccgcatgatctcgcccctgtcttggatgcccattgccgttatggcctttggcattggcgatctgcccgtttattttctgctag<br>cgatcgcagcggtctggccaattctactcagcaccagcagcggtactgcggctgttaaccacaagctcttacttctagctcgtagcctttgtgccactcgcagtgaaacg<br>attcggcgcatcgtgattcctgcgatcgtgccgcaaatccttgtgggtgttcgactcgcgatcggcacgatctggattgtgctcgtccccgctgagatgctgggcgttagc<br>tcaggcctcggttacttcattcttgatacccgcgatcgcattgcctataacgaacttaccgctgttttattagcgatcggcatcattggctgcgccctagattggagcctgca<br>attcctgcaaaagtactggcaacccagtcgctaaacctcttggcatccattctccaaacctctcaaaggcgtctacagggttgagcttgtggggaggggcggctatcctc<br>gcccgccaaaagcccgatcgtcatcactggcaagcccagcaacgacagatggaactgaactcagtccaagggcagctcgatgccctggtcgtcaaagcaccctacg<br>caggcagggttaggcgggtacgctggctagagcaggtgggcggacaggtcaaggtggagattgctcttcaagtctcaaacactccctaagttttgaccttgattacagt<br>ggccccatctagtttttgtatatacaatgactaggtgcgtatcacctacgatccagtcaaacgcgacaagacactgcttgagcgaggactcgacttcgagagtgcgatcg<br>aagtctttgcagggctcacgctagaagtcgaagacacccgacgcgactatggcgaaaagcgagtgctctgcgtcggcttccttgatggccgcatggtgatggttggct<br>atacccctcgcggtcgaacacgtcacatcttttcgatgaggaaatgcaatgcccgagaaattcgccgctacacgccatacttcgagcaagcctgatgacgacttgcccg<br>agttgaccgatgaaatgctcagtcgggcggtgctcaagcagggcggtaagcgtatcggcagaccccgatcgcaatctccaaaagtagcgatcagcctgcgcttagaa<br>gctgaagtcctgcagcgctggcgtgacagtggccctggctggcaaactcgtatggctgaattgctctcgaaagaaacccccctaagtctttgatcgctcaagcccaacgg<br>caggacaagctacaccagtggcaaacccagcagcgacagatggaacttaactcagtccaagcgcagctcgatgcactggtcgtcaaagcaccttactccggcagag |

tcaggcgggtgcgttggctagagcaagtcggtggacaggtgaaagtagagattgctcttcagatctttaacgagtgataaattcatagctttgtcattatctattatcgaga
gcaaataagagctatcaaggagcacctaagtgatctgctacttggtaacctttttcagttagtctataaacttctcctttgccgccaagtgactccattagctcatatttaataa
gctctgaaatagcacttttccaagaggcctcactacgagcattacccatatctgcaagctgcttattattagtttcaatcatcaatccacccataactggcagacaaattaatt
ctccattagaatctgatgctacctctattaacagatcttttccaagatcgcttaagcgtggtaattcaggagcaaaagatacttctaaagtattattagtgctttctaagctactg
taattttgtataattttctgagatatctgccgactaaacttatgctgaaactcctccaatgactcatatgtttcaattaagcctttttgttgacacaaactacgaaaatcacatagtg
ccttgtattgatcctgatcaacgctttctagatatactggagctctggaaaagtaaatcatcgcttgcttgccagacttaagatgttcttctatttcctcaacagttccactaata
aatttaccagtaggtgttcctaatcttgtccaaaaaacagcaacaagtagatcgcagtctttcaaaatttgcttgtttattatggattgtgctcgtccgctcatatcaggatggg
catgtgtttcccaccctactggatcaagtactattttacgatcaatagaattaatggtattccactcgtatattatgttccgaataatgttgcgttcttttgcaacatcgctcggtg
atgcaatcataatcctaagaactgtagcgttgtatggcattgctagccccttagagaaaatttcagtagcatttactatttattagctatatttacataaatttttaaactcatcag
agataatgactcatattttctatacgatcgcaagtaacttcactgccccttcaccacctccgtatcgcccgagaaagcaaagtgctccgcctgactcaccagcgactctg
gcaacggcagcgtaatccgatagcactccagaaactcacccagcactcccatcgcatgggctttgaccgaatccgagaaccgatcgccctctCCACCTCGAC
CTGAATGGAAGCCGGCGGCACCTCGCTAACGGATTCACCACTCCAAGAATTGGAGCCAATCAATTCTT
GCGGAGAACTGTGAATGCGCAAACCAACCCTTGGCAGAACATATCCATCGCGTCCGCCATCTCCAGCA
GCCGCACGCGGCGCATCTCGGGCAGCGTTGGGTCCTGGCCACGGGTGCGCATGATCGTGCTCCTGTCG
TTGAGGACCCGGCTAGGCTGGCGGGGTTGCCTTACTGGTTAGCAGAATGAATCACCGATACGCGAGCG
AACGTGAAGCGACTGCTGCTGCAAAACGTCTGCGACCTGAGCAACAACATGAATGGTCTTCGGTTTCC
GTGTTTCGTAAAGTCTGGAAACGCGGAAGTCAGCGCCCTGCACCATTATGTTCCGGATCTGCATCGCA
GGATGCTGCTGGCTACCCTGTGGAACACCTACATCTGTATTAACGAAGCGCTGGCATTGACCCTGAGTG
ATTTTTCTCTGGTCCCGCCGCATCCATACCGCCAGTTGTTTACCCTCACAACGTTCCAGTAACCGGGCAT
GTTCATCATCAGTAACCCGTATCGTGAGCATCCTCTCTCGTTTCATCGGTATCATTACCCCCATGAACAG
AAATCCCCCTTACACGGAGGCATCAGTGACCAAACAGGAAAAAACCGCCCTTAACATGGCCCGCTTTA
TCAGAAGCCAGACATTAACGCTTCTGGAGAAACTCAACGAGCTGGACGCGGATGAACAGGCAGACAT
CTGTGAATCGCTTCACGACCACGCTGATGAGCTTTACCGCAGCTGCCTCGCGCGTTTCGGTGATGACG
GTGAAAACCTCTGACACATGCAGCTCCCGGGACGGTCACAGCTTGTCTGTAAGCGGATGCCGGGAGC
AGACAAGCCCGTCAGGGCGCGTCAGCGGGTGTTGGCGGGTGTCGGGGCGCAGCCATGACCCAGTCAC
GTAGCGATAGCGGAGTGTATACTGGCTTAACTATGCGGCATCAGAGCAGATTGTACTGAGAGTGCACCA
TATGCGGTGTGAAATACCGCACAGATGCGTAAGGAGAAAATACCGCATCAGGCGCTCTTCCGCTTCCTC
GCTCACTGACTCGCTGCGCTCGGTCGTTCGGCTGCGGCGAGCGGTATCAGCTCACTCAAAGGCGGTAA
TACGGTTATCCACAGAATCAGGGGATAACGCAGGAAAGAACATGTGAGCAAAAGGCCAGCAAAAGGC
CAGGAACCGTAAAAAGGCCGCGTTGCTGGCGTTTTTCCATAGGCTCCGCCCCCCTGACGAGCATCACA
AAAATCGACGCTCAAGTCAGAGGTGGCGAAACCCGACAGGACTATAAAGATACCAGGCGTTTCCCCCT
GGAAGCTCCCTCGTGCGCTCTCCTGTTCCGACCCTGCCGCTTACCGGATACCTGTCCGCCTTTCTCCCT
TCGGGAAGCGTGGCGCTTTCTCATAGCTCACGCTGTAGGTATCTCAGTTCGGTGTAGGTCGTTCGCTCC
AAGCTGGGCTGTGTGCACGAACCCCCCGTTCAGCCCGACCGCTGCGCCTTATCCGGTAACTATCGTCTT
GAGTCCAACCCGGTAAGACACGACTTATCGCCACTGGCAGCAGCCACTGGTAACAGGATTAGCAGAG
CGAGGTATGTAGGCGGTGCTACAGAGTTCTTGAAGTGGTGGCCTAACTACGGCTACACTAGAAGGACA
GTATTTGGTATCTGCGCTCTGCTGAAGCCAGTTACCTTCGGAAAAAGAGTTGGTAGCTCTTGATCCGGC
AAACAAACCACCGCTGGTAGCGGTGGTTTTTTTGTTTGCAAGCAGCAGATTACGCGCAGAAAAAAAG
GATCTCAAGAAGATCCTTTGATCTTTTCTACGGGGTCTGACGCTCAGTGGAACGAAAACTCACGTTAA
GGGATTTTGGTCATGAGATTATCAAAAAGGATCTTCACCTAGATCCTTTTAAATTAAAAATGAAGTTTTA
AATCAATCTAAAGTATATATGAGTAAACTTGGTCTGACAGTTACCAATGCTTAATCAGTGAGGCACCTAT
CTCAGCGATCTGTCTATTTCGTTCATCCATAGTTGCCTGACTCCCCGTCGTGTAGATAACTACGATACGG
GAGGGCTTACCATCTGGCCCCAGTGCTGCAATGATACCGCGcGACCCACGCTCACCGGCTCCAGATTTA
TCAGCAATAAACCAGCCAGCCGGAAGGGCCGAGCGCAGAAGTGGTCCTGCAACTTTATCCGCCTCCAT
CCAGTCTATTAATTGTTGCCGGGAAGCTAGAGTAAGTAGTTCGCCAGTTAATAGTTTGCGCAACGTTGT
TGCCATTGCTGCAGGCATCGTGGTGTCACGCTCGTCGTTTGGTATGGCTTCATTCAGCTCCGGTTCCCA
ACGATCAAGGCGAGTTACATGATCCCCCATGTTGTGCAAAAAAGCGGTTAGCTCCTTCGGTCCTCCGAT
CGTTGTCAGAAGTAAGTTGGCCGCAGTGTTATCACTCATGGTTATGGCAGCACTGCATAATTCTCTTACT
GTCATGCCATCCGTAAGATGCTTTTCTGTGACTGGTGAGTACTCAACCAAGTCATTCTGAGAATAGTGT

| | |
|---|---|
| | ATGCGGCGACCGAGTTGCTCTTGCCCGGCGTCAACACGGGATAATACCGCGCCACATAGCAGAACTTT<br>AAAAGTGCTCATCATTGGAAAACGTTCTTCGGGGCGAAAACTCTCAAGGATCTTACCGCTGTTGAGAT<br>CCAGTTCGATGTAACCCACTCGTGCACCCAACTGATCTTCAGCATCTTTTACTTTCACCAGCGTTTCTG<br>GGTGAGCAAAAACAGGAAGGCAAAATGCCGCAAAAAAGGGAATAAGGGCGACACGGAAATGTTGAA<br>TACTCATACTCTTCCTTTTTCAATATTATTGAAGCATTTATCAGGGTTATTGTCTCATGAGCGGATACATAT<br>TTGAATGTATTTAGAAAAATAAACAAATAGGGGTTCCGCG |
| pRSF-ori | TTATTTGCCGACTACCTTGGTGATCTCGCCTTTCACGTAGTGGACAAATTCTTCCAACTGATCTGCGCGC<br>GAGGCCAAGCGATCTTCTTCTTGTCCAAGATAAGCCTGTCTAGCTTCAAGTATGACGGGCTGATACTGG<br>GCCGGCAGGCGCTCCATTGCCCAGTCGGCAGCGACATCCTTCGGCGCGATTTTGCCGGTTACTGCGCT<br>GTACCAAATGCGGGACAACGTAAGCACTACATTTCGCTCATCGCCAGCCCAGTCGGGCGGCGAGTTCC<br>ATAGCGTTAAGGTTTCATTTAGCGCCTCAAATAGATCCTGTTCAGGAACCGGATCAAAGAGTTCCTCCG<br>CCGCTGGACCTACCAAGGCAACGCTATGTTCTCTTGCTTTTGTCAGCAAGATAGCCAGATCAATGTCGA<br>TCGTGGCTGGCTCGAAGATACCTGCAAGAATGTCATTGCGCTGCCATTCTCCAAATTGCAGTTCGCGCT<br>TAGCTGGATAACGCCACGGAATGATGTCGTCGTGCACAACAATGGTGACTTCTACAGCGCGGAGAATC<br>TCGCTCTCTCCAGGGGAAGCCGAAGTTTCCAAAAGGTCGTTGATCAAAGCTCGCCGCGTTGTTTCATC<br>AAGCCTTACGGTCACCGTAACCAGCAAATCAATATCACTGTGTGGCTTCAGGCCGCCATCCACTGCGG<br>AGCCGTACAAATGTACGGCCAGCAACGTCGGTTCGAGATGGCGCTCGATGACGCCAACTACCTCTGAT<br>AGTTGAGTCGATACTTCGGCGATCACCGCTTCCCTCATGATGTTTAACTTTGTTTTAGGGCGACTGCCCT<br>GCTGCGTAACATCGTTGCTGCTCCATAACATCAAACATCGACCCACGGCGTAACGCGCTTGCTGCTTGG<br>ATGCCCGAGGCATAGACTGTACCCCAAAAAAACAGTCATAACAAGCCATGAAAACCGCCACTGCGCC<br>GTTACCACCGCTGCGTTCGGTCAAGGTTCTGGACCAGTTGCGTGAGCGCATACGCTACTTGCATTACAG<br>CTTACGAACCGAACAGGCTTATGTCCACTGGGTTCGTGCCTTCATCCGTTTCCACGGTGTGCGgctcactcaa<br>aggcggtaatcaattgagttcttttaccctcagccgaaatgcctgccgttgctagacattgccagccagtgcccgtcactcccgtactaactgtcacgaacccctgcaata<br>actgtcacgccccccctgcaataactgtcacgaacccctgcaataactgtcacgcccccaaacctgcaaacccagcaggggcgggggctggcggggtgttggaaaaa<br>tccatccatgattatctaagaataatccactaggcgcggttatcagcgcccttgtggggcgctgctgcccttgcccaatatgcccggccagaggccggatagctggtcta<br>ttcgctgcgctaggctacacaccgccccaccgctgcgcggcaggggggaaaggcgggcaaagcccgctaaaccccacaccaaaccccgcagaaatacgctggag<br>cgcttttagccgctttagcggcctttccccctacccgaagggtgggggcgcgtgtgcagccccgcagggcctgtctcggtcgatcattcagcccggctcatccttctgg<br>cgtggcggcagaccgaacaaggcgcggtcgtggtcgcgttcaaggtacgcatccattgccgccatgagccgatcctccggccactcgctgctgttcaccttggccaa<br>aatcatggcccccaccagcaccttgcgccttgtttcgttcttgcgctcttgctgctgttcccttgcccgcacccgctgaatttcggcattgattcgcgctcgttgttcttcgag<br>cttggccagccgatccgccgccttgttgctccccttaaccatcttgacaccccattgttaatgtgctgtctcgtaggctatcatggaggcacagcggcggcaatcccgacc<br>ctactttgtaggggagggcgcacttaccggtttctcttcgagaaactggcctaacggccacccttcgggcggtgcgctctccgagggccattgcatggagccgaaaag<br>caaaagcaacagcgaggcagcatggcgatttatcaccttacggcgaaaaccggcagcaggtcgggcggccaatcggccagggccaaggccgactacatccagcg<br>cgaaggcaagtatgcccgcgacatggatgaagtcttgcacgccgaatccgggcacatgccggagttcgtcgagcggcccgccgactactgggatgctgccgacctg<br>tatgaacgcgccaatgggcggctgttcaaggaggtcgaatttgccctgccggtcgagctgaccctcgaccagcagaaggcgctggcgtccgagttcgcccagcacct<br>gaccggtgccgagcgcctgccgtatacgctggccatccatgccggtggcggcgagaacccgcactgccacctgatgatctccgagcggatcaatgacggcatcga<br>gcggcccgccgctcagtggttcaagcggtacaacggcaagaccccggagaagggcggggcacagaagaccgaagcgctcaagcccaaggcatggcttgagca<br>gacccgcgaggcatgggccgaccatgccaaccgggcattagagcgggctggccacgacgcccgcattgaccacagaacacttgaggcgcagggcatcgagcgc<br>ctgcccggtgttcacctggggccgaacgtggtggagatggaaggccggggcatccgcaccgaccgggcagacgtggccctgaacatcgacaccgccaacgccca<br>gatcatcgacttacaggaataccgggaggcaatagaccatgaacgcaatcgacagagtgaagaaatccagaggcatcaacgagttagcggagcagatcgaaccgct<br>ggcccagagcatggcgacactggccgacgaagcccggcaggtcatgagccagacccagcaggccagcgaggcgcaggcggcggagtggctgaaagcccagc<br>gccagacaggggcggcatgggtggagctggccaaagagttgcgggaggtagccgccgaggtgagcagcgccgcgcagagcgcccggagcgcgtcgcggggg<br>tggcactggaagctatggctaaccgtgatgctggcttccatgatgcctacggtggtgctgctgatcgcatcgttgctcttgctcgacctgacgccactgacaaccgagga<br>cggctcgatctggctgcgcttggtggcccgatgaagaacgacaggactttgcaggccataggccgacagctcaaggccatgggctgtgagcgcttcgatatcggcgt<br>cagggacgccaccaccggccagatgatgaaccgggaatggtcagccgccgaagtgctccagaacacgccatggctcaagcggatgaatgcccagggcaatgacg<br>tgtatatcaggcccgccgagcaggagcggcatggtctggtgctggtggacgacctcagcgagtttgacctggatgacatgaaagccgagggccgggagcctgccct<br>ggtagtggaaaccagcccgaagaactatcaggcatgggtcaaggtggccgacgccgcaggcggtgaacttcgggggcagattgcccggacgctggccagcgagt<br>acgacgccgacccggccagcgccgacagccgccactatggccgcttggcgggcttcaccaaccgcaaggacaagcacaccacccgcgccggttatcagccgtgg<br>gtgctgctgcgtgaatccaagggcaagaccgccaccgctggcccggcgctggtgcagcaggctggccagcagatcgagcaggcccagcggcagcaggagaag<br>gcccgcaggctggccagcctcgaactgcccgagcggcagcttagccgccaccggcgcacggcgctggacgagtaccgcagcgagatggccgggctggtcaagc |

| | |
|---|---|
| | gcttcggtgatgacctcagcaagtgcgactttatcgccgcgcagaagctggccagccggggccgcagtgccgaggaaatcggcaaggccatggccgaggccagc<br>ccagcgctggcagagcgcaagcccggccacgaagcggattacatcgagcgcaccgtcagcaaggtcatgggtctgcccagcgtccagcttgcgcgggccgagct<br>ggcacgggcaccggcaccccgccagcgaggcatggacaggggcgggccagatttcagcatgtagtgcttgcgttggtactcacgcctgttatactatgagtactcac<br>gcacagaagggggttttatggaatacgaaaaaagcgcttcagggtcggtctacctgatcaaaagtgacaagggctattggttgcccggtggctttggttatacgtcaaac<br>aaggccgaggctggccgcttttcagtcgctgatatggccagccttaaccttgacggctgcaccttgtccttgttccgcgaagacaagcctttcggccccggcaagtttctc<br>ggtgactgatatgaaagaccaaaaggacaagcagaccggcgacctgctggccagccctgacgctgtacgccaagcgcgatatgccgagcgcatgaaggccaaag<br>ggatgcgtcagcgcaagttctggctgaccgacgacgaatacgaggcgctgcgcgagtgcctggaagaactcagagcggcgcagggcgggggtagtgaccccgc<br>cagcgcctaaccaccaactgcctgcaaaggaggcaatcaatggctacccataagcctatcaatattctggaggcgttcgcagcagcgccgccaccgctggactacgtt<br>ttgcccaacatggtggccggtacggtcggggcgctggtgtcgcccggtggtgccggtaaatccatgctggccctgcaactggccgcacagattgcaggcgggccgg<br>atctgctggaggtgggcgaactgcccaccggcccggtgatctacctgcccgccgaagacccgcccaccgccattcatcaccgcctgcacgcccttggggcgcacct<br>cagcgccgaggaacggcaagccgtggctgacggcctgctgatccagccgctgatcggcagcctgcccaacatcatggccccggagtggttcgacggcctcaagcg<br>cgccgccgagggccgccgcctgatggtgctggacacgctgcgccggttccacatcgaggaagaaaacgccagcggccccatggcccaggtcatcggtcgcatgg<br>aggccatcgccgccgataccgggtgctctatcgtgttcctgcaccatgccagcaagggcgcggccatgatgggcgcaggcgaccagcagcaggccagccggggc<br>agctcggtactggtcgataacatccgctggcagtcctacctgtcgagcatgaccagcgccgaggccgaggaatggggtgtggacgacgaccagcgccggttcttcgt<br>ccgcttcggtgtgagcaaggccaactatggcgcaccgttcgctgatcggtggttcaggcggcatgacggcggggtgctcaagcccgccgtgctggagaggcagcg<br>caagagcaagggggtgccccgtggtgaagcctaagaacaagcacagcctcagccacgtccggcacgacccggcgcactgtctggcccccggcctgttccgtgccc<br>tcaagcggggcgagcgcaagcgcagcaagctggacgtgacgtatgactacggcgacggcaagcggatcgagttcagcggcccggagccgctgggcgctgatga<br>tctgcgcatcctgcaagggctggtggccatggctgggcctaatggcctagtgcttggcccggaacccaagaccgaaggcggacggcagctccggctgttcctggaa<br>cccaagtgggaggccgtcaccgctgatgccatggtggtcaaaggtagctatcgggcgctggcaaaggaaatcggggcagaggtcgatagtggtggggcgctcaag<br>cacatacaggactgcatcgagcgcctttggaaggtatccatcatcgcccagaatggccgcaagcggcaggggtttcggctgctgtcggagtacgccagcgacgagg<br>cggacgggcgcctgtacgtggccctgaaccccttgatcgcgcaggccgtcatgggtggcggccagcatgtgcgcatcagcatggacgaggtgcgggcgctggaca<br>gcgaaaccgcccgcctgctgcaccagcggctgtgtggctggatcgaccccggcaaaaccggcaaggcttccatagataccttgtgcggctatgtctggccgtcagag<br>gccagtggttcgaccatgcgcaagcgccgccagcgggtgcgcgaggcgttgccggagctggtcgcgctgggctggacggtaaccgagttcgcggcgggcaagta<br>cgacatcacccggcccaaggcggcaggctgacccc |
| pRSF | TTATTTGCCGACTACCTTGGTGATCTCGCCTTTCACGTAGTGGACAAATTCTTCCAACTGATCTGCGCGC<br>GAGGCCAAGCGATCTTCTTCTTGTCCAAGATAAGCCTGTCTAGCTTCAAGTATGACGGGCTGATACTGG<br>GCCGGCAGGCGCTCCATTGCCCAGTCGGCAGCGACATCCTTCGGCGCGATTTTGCCGGTTACTGCGCT<br>GTACCAAATGCGGGACAACGTAAGCACTACATTTCGCTCATCGCCAGCCCAGTCGGGCGGCGAGTTCC<br>ATAGCGTTAAGGTTTCATTTAGCGCCTCAAATAGATCCTGTTCAGGAACCGGATCAAAGAGTTCCTCCG<br>CCGCTGGACCTACCAAGGCAACGCTATGTTCTCTTGCTTTTGTCAGCAAGATAGCCAGATCAATGTCGA<br>TCGTGGCTGGCTCGAAGATACCTGCAAGAATGTCATTGCGCTGCCATTCTCCAAATTGCAGTTCGCGCT<br>TAGCTGGATAACGCCACGGAATGATGTCGTCGTGCACAACAATGGTGACTTCTACAGCGCGGAGAATC<br>TCGCTCTCTCCAGGGGAAGCCGAAGTTTCCAAAAGGTCGTTGATCAAAGCTCGCCGCGTTGTTTCATC<br>AAGCCTTACGGTCACCGTAACCAGCAAATCAATATCACTGTGTGGCTTCAGGCCGCCATCCACTGCGG<br>AGCCGTACAAATGTACGGCCAGCAACGTCGGTTCGAGATGGCGCTCGATGACGCCAACTACCTCTGAT<br>AGTTGAGTCGATACTTCGGCGATCACCGCTTCCCTCATGATGTTTAACTTTGTTTTAGGGCGACTGCCCT<br>GCTGCGTAACATCGTTGCTGCTCCATAACATCAAACATCGACCCACGGCGTAACGCGCTTGCTGCTTGG<br>ATGCCCGAGGCATAGACTGTACCCCAAAAAAACAGTCATAACAAGCCATGAAAACCGCCACTGCGCC<br>GTTACCACCGCTGCGTTCGGTCAAGGTTCTGGACCAGTTGCGTGAGCGCATACGCTACTTGCATTACAG<br>CTTACGAACCGAACAGGCTTATGTCCACTGGGTTCGTGCCTTCATCCGTTTCCACGGTGTGCGgctcactcaa<br>aggcggtaatcaattgagttcttttaccctcagccgaaatgcctgccgttgctagacattgccagccagtgcccgtcactcccgtactaactgtcacgaacccctgcaata<br>actgtcacgccccccctgcaataactgtcacgaacccctgcaataactgtcacgcccccaaacctgcaaacccagcaggggcgggggctggcggggtgttggaaaaa<br>tccatccatgattatctaagaataatccactaggcgcggttatcagcgcccttgtggggcgctgctgcccttgcccaatatgcccggccagaggccggatagctggtcta<br>ttcgctgcgctaggctacacaccgccccaccgctgcgcggcagggggaaaggcgggcaaagcccgctaaaccccacaccaaaccccgcagaaatacgctggag<br>cgcttttagccgctttagcggcctttccccctacccgaagggtgggggcgcgtgtgcagccccgcagggcctgtctcggtcgatcattcagcccggctcatccttctgg<br>cgtggcggcagaccgaacaaggcgcggtcgtggtcgcgttcaaggtacgcatccattgccgccatgagccgatcctccggccactcgctgctgttcaccttggccaa<br>aatcatggcccccaccagcaccttgcgccttgtttcgttcttgcgctcttgctgctgttcccttgcccgcacccgctgaatttcggcattgattcgcgctcgttgttcttcgag<br>cttggccagccgatccgccgccttgttgctccccttaaccatcttgacaccccattgttaatgtgctgtctcgtaggctatcatggaggcacagcggcggcaatcccgacc<br>ctactttgtaggggagggcgcacttaccggtttctcttcgagaaactggcctaacggccacccttcgggcggtgcgctctccgagggccattgcatggagccgaaaag |

| | |
|---|---|
| | caaaagcaacagcgaggcagcatggcgatttatcaccttacggcgaaaaccggcagcaggtcgggcggccaatcggccagggccaaggccgactacatccagcgcgaaggcaagtatgcccgcgacatggatgaagtcttgcacgccgaatccgggcacatgccggagttcgtcgagcggcccgccgactactgggatgctgccgacctgtatgaacgcgccaatgggcggctgttcaaggaggtcgaatttgccctgccggtcgagctgaccctcgaccagcagaaggcgctggcgtccgagttcgcccagcacctgaccggtgccgagcgcctgccgtatacgctggccatccatgccggtggcggcgagaacccgcactgccacctgatgatctccgagcggatcaatgacggcatcgagcggcccgccgctcagtggttcaagcggtacaacggcaagaccccggagaagggcggggcacagaagaccgaagcgctcaagcccaaggcatggcttgagcagacccgcgaggcatgggccgaccatgccaaccgggcattagagcgggctggccacgacgcccgcattgaccacagaacacttgaggcgcagggcatcgagcgcctgcccggtgttcacctggggccgaacgtggtggagatggaaggccggggcatccgcaccgaccgggcagacgtggccctgaacatcgacaccgccaacgcccagatcatcgacttacaggaataccgggaggcaatagaccatgaacgcaatcgacagagtgaagaaatccagaggcatcaacgagttagcggagcagatcgaaccgctggcccagagcatggcgacactggccgacgaagcccggcaggtcatgagccagacccagcaggccagcgaggcgcaggcggcggagtggctgaaagcccagcgccagacaggggcggcatgggtggagctggccaaagagttgcgggaggtagccgccgaggtgagcagcgccgcgcagagcgcccggagcgcgtcgcgggggtggcactggaagctatggctaaccgtgatgctggcttccatgatgcctacggtggtgctgctgatcgcatcgttgctcttgctcgacctgacgccactgacaaccgaggacggctcgatctggctgcgcttggtggcccgatgaagaacgacaggactttgcaggccataggccgacagctcaaggccatgggctgtgagcgcttcgatatcggcgtcagggacgccaccaccggccagatgatgaaccgggaatggtcagccgccgaagtgctccagaacacgccatggctcaagcggatgaatgcccagggcaatgacgtgtatatcaggcccgccgagcaggagcggcatggtctggtgctggtggacgacctcagcgagtttgacctggatgacatgaaagccgagggccgggagcctgccctggtagtggaaaccagcccgaagaactatcaggcatgggtcaaggtggccgacgccgcaggcggtgaacttcggggggcagattgcccggacgctggccagcgagtacgacgccgacccggccagcgccgacagccgccactatggccgcttggcgggcttcaccaaccgcaaggacaagcacaccacccgcgccggttatcagccgtgggtgctgctgcgtgaatccaagggcaagaccgccaccgctggcccggcgctggtgcagcaggctggccagcagatcgagcaggcccagcggcagcaggagaaggcccgcaggctggccagcctcgaactgcccgagcggcagcttagccgccaccggcgcacggcgctggacgagtaccgcagcgagatggccgggctggtcaagcgcttcggtgatgacctcagcaagtgcgactttatcgccgcgcagaagctggccagccgggggccgcagtgccgaggaaatcggcaaggccatggccgaggccagcccagcgctggcagagcgcaagcccggccacgaagcggattacatcgagcgcaccgtcagcaaggtcatgggtctgcccagcgtccagcttgcgcgggccgagctggcacgggcaccggcaccccgccagcgaggcatggacaggggcgggccagatttcagcatgtagtgcttgcgttggtactcacgcctgttatactatgagtactcacgcacagaaggggggttttatggaatacgaaaaaagcgcttcagggtcggtctacctgatcaaaagtgacaagggctattggttgcccggtggctttggttatacgtcaaacaaggccgaggctggccgcttttcagtcgctgatatggccagccttaaccttgacggctgcaccttgtccttgttccgcgaagacaagcctttcggccccggcaagtttctcggtgactgatcgccaagcgcgatatgccgagcgcatgaaggccaaagggatgcgtcagcgcaagttctggctgaccgacgacgaatacgaggcgctgcgcgagtgcctggaagaactcagagcggcgcagggcgggggtagtgaccccgccagcgcctaaccaccaactgcctgcaaaggaggcaatcaatggctacccataagcctatcaatattctggaggcgttcgcagcagcgccgccaccgctggactacgttttgcccaacatggtggccggtacggtcggggcgctggtgtcgcccggtggtgccggtaaatccatgctggccctgcaactggccgcacagattgcaggcgggccggatctgctggaggtgggcgaactgcccaccggcccggtgatctacctgcccgccgaagacccgcccaccgccattcatcaccgcctgcacgcccttggggcgcacctcagcgccgaggaacggcaagccgtggctgacggcctgctgatccagccgctgatcggcagcctgcccaacatcatggccccggagtggttcgacggcctcaagcgcgccgccgagggccgccgcctgatggtgctggacacgctgcgccggttccacatcgaggaagaaaacgccagcggccccatggcccaggtcatcggtcgcatggaggccatcgccgccgataccgggtgctctatcgtgttcctgcaccatgccagcaagggcgcggccatgatgggcgcaggcgaccagcagcaggccagccggggcagctcggtactggtcgataacatccgctggcagtcctacctgtcgagcatgaccagcgccgaggccgaggaatggggtgtggacgacgaccagcgccggttcttcgtccgcttcggtgtgagcaaggccaactatggcgcaccgttcgctgatcggtggttcaggcggcatgacggcggggtgctcaagcccgccgtgctggagaggcagcgcaagagcaaggggggtgccccgtggtgaagcctaagaacaagcacagcctcagccacgtccggcacgacccggcgcactgtctggcccccggcctgttccgtgccctcaagcggggcgagcgcaagcgcagcaagctggacgtgacgtatgactacggcgacggcaagcggatcgagttcagcggcccggagccgctgggcgctgatgatctgcgcatcctgcaagggctggtggccatggctgggcctaatggcctagtgcttggcccggaacccaagaccgaaggcggacggcagctccggctgttcctggaacccaagtgggaggccgtcaccgctgatgccatggtggtcaaaggtagctatcgggcgctggcaaaggaaatcggggcagaggtcgatagtggtggggcgctcaagcacatacaggactgcatcgagcgcctttggaaggtatccatcatcgcccagaatggccgcaagcggcaggggtttcggctgctgtcggagtacgccagcgacgaggcggacgggcgcctgtacgtggccctgaaccccttgatcgcgcaggccgtcatgggtggcggccagcatgtgcgcatcagcatggacgaggtgcgggcgctggacagcgaaaccgcccgcctgctgcaccagcggctgtgtggctggatcgaccccggcaaaaccggcaaggcttccatagatacctgtgcggctatgtctggccgtcagaggccagtggttcgaccatgcgcaagcgccgccagcgggtgcgcgaggcgttgccggagctggtcgcgctgggctggacggtaaccgagttcgcggcgggcaagtacgacatcacccggcccaaggcggcaggctgacccc |
| pSI-single | gatccggcagccggcggagcgctgctttcttggcaagcggtcgccagccccaacgccagggctgccagcccgaaacagcggggcaaggcagcttggaagggcgatcgcagcacgggcatggcaatgtctctctgaaggaatgcagaccttattcgtacagccagggttgaatcgtgggggtccaatcacttagctctgctgggctaaaccagagagcaatttcctgttgtgctgtttcgattgcatccgagccatggatgatgttgcggccaatattgacaccaaaatcaccacggatggtgcccggttctgccgtcagcggattggtagcgccgatcaacttgcgagcagccgccacaacgccttcgccttccaagacgatcgccacgatcggcccagaggtgatgaactcgacgaggccattgaagaaggggcgctcgcggtggacagcatagtgctgttcggccagctcgcgactgggcttcagctgctttaggcccaccagtttgaagccttttttgctcaaagcggccgatgatcgtaccgaccaaaccccgctgaacgccatcgggcttgatggcaataaatgtgcgttccacagacatctagatagtcctcaagacgaggcaagcattgagcttgccttcctatggttcgggatcactgggattcttgacaagcgatcgcggtcacatcgctatctcttaggacttcgcagcgggcgagtcggattgacccggtagggatttcgccagatcaatgcccgtggtttgtttcagcttctccagcaagctagcgatttgggtagcgctgccttccccttcgccaatcacagtgatcgactccacgtcgatatctggcacggtgcct |

gaaagcgtgacgagcagggactcgaTATAAACGCAGAAAGGCCCACCCGAAGGTGAGCCAGTGTGACTCTAGTAGA
GAGCGTTCACCGACAAACAACAGATAAAACGAAAGGCCCAGTCTTTCGACTGAGCCTTTCGTTTTATT
TGATGCCTGGTTATTTGCCGACTACCTTGGTGATCTCGCCTTTCACGTAGTGGACAAATTCTTCCAACTG
ATCTGCGCGCGAGGCCAAGCGATCTTCTTCTTGTCCAAGATAAGCCTGTCTAGCTTCAAGTATGACGGG
CTGATACTGGGCCGGCAGGCGCTCCATTGCCCAGTCGGCAGCGACATCCTTCGGCGCGATTTTGCCGG
TTACTGCGCTGTACCAAATGCGGGACAACGTAAGCACTACATTTCGCTCATCGCCAGCCCAGTCGGGC
GGCGAGTTCCATAGCGTTAAGGTTTCATTTAGCGCCTCAAATAGATCCTGTTCAGGAACCGGATCAAAG
AGTTCCTCCGCCGCTGGACCTACCAAGGCAACGCTATGTTCTCTTGCTTTTGTCAGCAAGATAGCCAGA
TCAATGTCGATCGTGGCTGGCTCGAAGATACCTGCAAGAATGTCATTGCGCTGCCATTCTCCAAATTGC
AGTTCGCGCTTAGCTGGATAACGCCACGGAATGATGTCGTCGTGCACAACAATGGTGACTTCTACAGC
GCGGAGAATCTCGCTCTCTCCAGGGGAAGCCGAAGTTTCCAAAAGGTCGTTGATCAAAGCTCGCCGC
GTTGTTTCATCAAGCCTTACGGTCACCGTAACCAGCAAATCAATATCACTGTGTGGCTTCAGGCCGCCA
TCCACTGCGGAGCCGTACAAATGTACGGCCAGCAACGTCGGTTCGAGATGGCGCTCGATGACGCCAAC
TACCTCTGATAGTTGAGTCGATACTTCGGCGATCACCGCTTCCCTCATGATGTTTAACTTTGTTTTAGGG
CGACTGCCCTGCTGCGTAACATCGTTGCTGCTCCATAACATCAAACATCGACCCACGGCGTAACGCGCT
TGCTGCTTGGATGCCCGAGGCATAGACTGTACCCCAAAAAAACAGTCATAACAAGCCATGAAAACCGC
CACTGCGCCGTTACCACCGCTGCGTTCGGTCAAGGTTCTGGACCAGTTGCGTGAGCGCATACGCTACT
TGCATTACAGCTTACGAACCGAACAGGCTTATGTCCACTGGGTTCGTGCCTTCATCCGTTTCCACGGTG
TGCGCGCGGAACCCCTATTTGTTTATTTTTCTAAATACATTCAAATATGTATCCGCTCATGAGACAATAAC
CCTGATAAATGCTTCAATAATATTGAAAAAGGAAGAGTATGAGTATTCAACATTTCCGTGTCGCCCTTAT
TCCCTTTTTTGCGGCATTTTGCCTTCCTGTTTTTGCTCACCCAGAAACGCTGGTGAAAGTAAAAGATGC
TGAAGATCAGTTGGGTGCACGAGTGGGTTACATCGAACTGGATCTCAACAGCGGTAAGATCCTTGAGA
GTTTTCGCCCCGAAGAACGTTTTCCAATGATGAGCACTTTTAAAGTTCTGCTATGTGGCGCGGTATTATC
CCGTGTTGACGCCGGGCAAGAGCAACTCGGTCGCCGCATACACTATTCTCAGAATGACTTGGTTGAGT
ACTCACCAGTCACAGAAAAGCATCTTACGGATGGCATGACAGTAAGAGAATTATGCAGTGCTGCCATA
ACCATGAGTGATAACACTGCGGCCAACTTACTTCTGACAACGATCGGAGGACCGAAGGAGCTAACCGC
TTTTTTGCACAACATGGGGGATCATGTAACTCGCCTTGATCGTTGGGAACCGGAGCTGAATGAAGCCAT
ACCAAACGACGAGCGTGACACCACGATGCCTGCAGCAATGGCAACAACGTTGCGCAAACTATTAACT
GGCGAACTACTTACTCTAGCTTCCCGGCAACAATTAATAGACTGGATGGAGGCGGATAAAGTTGCAGG
ACCACTTCTGCGCTCGGCCCTTCCGGCTGGCTGGTTTATTGCTGATAAATCTGGAGCCGGTGAGCGTGG
GTCTCGCGGTATCATTGCAGCACTGGGGCCAGATGGTAAGCCCTCCCGTATCGTAGTTATCTACACGAC
GGGGAGTCAGGCAACTATGGATGAACGAAATAGACAGATCGCTGAGATAGGTGCCTCACTGATTAAGC
ATTGGTAACTGTCAGACCAAGTTTACTCATATATACTTTAGATTGATTTAAAACTTCATTTTTAATTTAAA
AGGATCTAGGTGAAGATCCTTTTTGATAATCTCATGACCAAAATCCCTTAACGTGAGTTTTCGTTCCACT
GAGCGTCAGACCCCGTAGAAAAGATCAAAGGATCTTCTTGAGATCCTTTTTTTCTGCGCGTAATCTGCT
GCTTGCAAACAAAAAAACCACCGCTACCAGCGGTGGTTTGTTTGCCGGATCAAGAGCTACCAACTCTT
TTTCCGAAGGTAACTGGCTTCAGCAGAGCGCAGATACCAAATACTGTCCTTCTAGTGTAGCCGTAGTTA
GGCCACCACTTCAAGAACTCTGTAGCACCGCCTACATACCTCGCTCTGCTAATCCTGTTACCAGTGGCT
GCTGCCAGTGGCGATAAGTCGTGTCTTACCGGGTTGGACTCAAGACGATAGTTACCGGATAAGGCGCA
GCGGTCGGGCTGAACGGGGGGTTCGTGCACACAGCCCAGCTTGGAGCGAACGACCTACACCGAACTG
AGATACCTACAGCGTGAGCTATGAGAAAGCGCCACGCTTCCCGAAGGGAGAAAGGCGGACAGGTATC
CGGTAAGCGGCAGGGTCGGAACAGGAGAGCGCACGAGGGAGCTTCCAGGGGGAAACGCCTGGTATC
TTTATAGTCCTGTCGGGTTTCGCCACCTCTGACTTGAGCGTCGATTTTTGTGATGCTCGTCAGGGGGGC
GGAGCCTATGGAAAAACGCCAGCAACGCGGCCTTTTTACGGTTCCTGGCCTTTTGCTGGCCTTTTGCTC
ACATGTTCTTTCCTGCGTTATCCCCTGATTCTGTGGATAACCGTATTACCGCCTTTGAGTGAGCTGATAC
CGCTCGCCGCAGCCGAACGACCGAGCGCAGCGAGTCAGTGAGCGAGGAAGCGGAAGAGCGCCTGAT
GCGGTATTTTCTCCTTACGCATCTGTGCGGTATTTCACACCGCATATGGTGCACTCTCAGTACAATCTGC
TCTGATGCCGCATAGTTAAGCCAGTATACACTCCGCTATCGCTACGTGACTGGGTCATGGCTGCGCCCC
GACACCCGCCAACACCCGCTGACGCGCCCTGACGGGCTTGTCTGCTCCCGGCATCCGCTTACAGACAA

| | |
|---|---|
| | GCTGTGACCGTCTCCGGGAGCTGCATGTGTCAGAGGTTTTCACCGTCATCACCGAAACGCGCGAGGCA<br>GCTGCGGTAAAGCTCATCAGCGTGGTCGTGAAGCGATTCACAGATGTCTGCCTGTTCATCCGCGTCCA<br>GCTCGTTGAGTTTCTCCAGAAGCGTTAATGTCTGGCTTCTGATAAAGCGGGCCATGTTAAGGGCGGTTT<br>TTTCCTGTTTGGTCACTGATGCCTCCGTGTAAGGGGGATTTCTGTTCATGGGGGTAATGATACCGATGA<br>AACGAGAGAGGATGCTCACGATACGGGTTACTGATGATGAACATGCCCGGTTACTGGAACGTTGTGAG<br>GGTAAACAACTGGCGGTATGGATGCGGCGGGACCAGAGAAAAATCACTCAGGGTCAATGCCAGCGCT<br>TCGTTAATACAGATGTAGGTGTTCCACAGGGTAGCCAGCAGCATCCTGCGATGCAGATCCGGAACATAA<br>TGGTGCAGGGCGCTGACTTCCGCGTTTCCAGACTTTACGAAACACGGAAACCGAAGACCATTCATGTT<br>GTTGCTCAGGTCGCAGACGTTTTGCAGCAGCAGTCGCTTCACGTTCGCTCGCGTATCGGTGATTCATTC<br>TGCTAACCAGTAAGGCAACCCCGCCAGCCTAGCCGGGTCCTCAACGACAGGAGCACGATCATGCGCA<br>CCCGTGGCCAGGACCCAACGCTGCCCGAGATGCGCCGCGTGCGGCTGCTGGAGATGGCGGACGCGAT<br>GGATATGTTCTGCCAAGGGTTGGTTTGCGCATTCACAGTTCTCCGCAAGAATTGATTGGCTCCAATTCT<br>TGGAGTGGTGAATCCGTTAGCGAGGTGCCGCCGGCTTCCAT |
| pSEL-sepT2 | ttagaaaaactcatcgagcatcaaatgaaactgcaatttattcatatcaggattatcaataccatatttttgaaaaagccgtttctgtaatgaaggagaaaactcaccgaggc<br>agttccataggatggcaagatcctggtatcggtctgcgattccgactcgtccaacatcaatacaacctattaatttcccctcgtcaaaaataaggttatcaagtgagaaatc<br>accatgagtgacgactgaatccggtgagaatggcaaaagcttatgcatttctttccagacttgttcaacaggccagccattacgctcgtcatcaaaatcactcgcatcaac<br>caaaccgttattcattcgtgattgcgcctgagcgagacgaaatacgcgatcgctgttaaaaggacaattacaaacaggaatcgaatgcaaccggcgcaggaacactgc<br>cagcgcatcaacaatattttcacctgaatcaggatattcttctaatacctggaatgctgttttcccggggatcgcagtggtgagtaaccatgcatcatcaggagtacggata<br>aaatgcttgatggtcggaagaggcataaattccgtcagccagtttagtctgaccatctcatctgtaacatcattggcaacgctacctttgccatgtttcagaaacaactctgg<br>cgcatcgggcttcccatacaatcgatagattgtcgcacctgattgcccgacattatcgcgagcccatttatacccatataaatcagcatccatgttggaatttaatcgcggc<br>ctggagcaagacgtttcccgttgaatatggctcataacaccccttgtattactgtttatgtaagcagacagttttattgttcatgatgatatatttttatcttgtgcaatgtaacatc<br>agagattttgagacacaacgtggctttgttgaataaatcgaacttttgctgagttgaaggatcagGAATTCCTTAGTTATTCCTATTCTGCACGA<br>ACTCAAAATACTCTTCATTTTTGATAACCAAATTGAGTTTTTTGCCCTCTTGATTATTTTTGATCCTACCT<br>AGTAGCCTCTTCTGCTCCTGCAGAATTGTGAGCGCTCACAATTGATATTTTGGTCTGTCGTTGCGATCGC<br>CCGTTGCAGGCCGACATGAAGGATTGACAATTAATCATCCGGCTCGTATAATGAATTGTGAGCGCTCAC<br>AATTGGTACCGGTGATACCAGCATCGTCTTGATGCCCTTGGCAGCACCCTGCTAAGGAGGCAACAAGat<br>gcaagcggtctttgataccaacatcctcatctaccacctcaaaggctgtcttcctgaagcggggagtcaaattctgcgcagcagtctggggcgcggtgccgtctgttcag<br>tcattacccgcctagaggttttgggttacgaccaaccttggccagaaagactgaaagcacaagctttgctccagctatttcgagaacgggcattggatgaatcgattgctg<br>attgcacgattcagctgcgtcaacaacaacggatcaaactgcccgatgccatcgttgccgcaactgcgctgacagagaacttgccactcgtgacgcgcaacaccaaa<br>gactttaaagccatcgctggcttacaactgattaacccctttcaaccgaactagGGCTCACCTTCGGGTGGGCCTTTCTGCGccatgatcgagc<br>gatcggcctcgggggcgttgcagttcatccgcatgatctcgcccctgtcttggatgcccattgccgttatggcctttggcattggcgatctgcccgtttattttctgctagcg<br>atcgcagcggtctggccaattctactcagcaccagcagcggtactgcggctgttaaccacaagctcttacttctagctcgtagcctttgtgccactcgcagtgaaacgatt<br>cggcgcatcgtgattcctgcgatcgtgccgcaaatccttgtgggtgttcgactcgcgatcggcacgatctggattgtgctcgtccccgctgagatgctgggcgttagctc<br>aggcctcggttacttcattcttgatacccgcgatcgcattgcctataacgaacttaccgctgttttattagcgatcggcatcattggctgcgccctagattggagcctgcaatt<br>cctgcaaaagtactggcaacccagtcgctaaacctcttggcatccattctccaaacctctcaaaggcgtctacagggttgagcttgtggggaggggcggctatcctcgc<br>ccgccaaaagcccgatcgtcatcactggcaagcccagcaacgacagatggaactgaactcagtccaagggcagctcgatgccctggtcgtcaaagcaccctacgca<br>ggcagggttaggcgggtacgctggctagagcaggtgggcggacaggtcaaggtggagattgctcttcaagtctcaaacactccctaagttttgaccttgattacagtgg<br>ccccatctagtttttgtatatacaatgactaggtgcgtatcacctacgatccagtcaaacgcgacaagacactgcttgagcgaggactcgacttcgagagtgcgatcgaa<br>gtctttgcagggctcacgctagaagtcgaagacacccgacgcgactatggcgaaaagcgagtgctctgcgtcggcttccttgatggccgcatggtgatggttggctata<br>cccctcgcggtcgaacacgtcacatcttttcgatgaggaaatgcaatgcccgagaaattcgccgctacacgccatacttcgagcaagcctgatgacgacttgcccgagt<br>tgaccgatgaaatgctcagtcgggcggtgctcaagcagggcggtaagcgtatcggcagaccccgatcgcaatctccaaaagtagcgatcagcctgcgcttagaagct<br>gaagtcctgcagcgctggcgtgacagtggccctggctggcaaactcgtatggctgaattgctctcgaaagaaacccccctaagtctttgatcgctcaagcccaacggca<br>ggacaagctacaccagtggcaaacccagcagcgacagatggaacttaactcagtccaagcgcagctcgatgcactggtcgtcaaagcaccttactccggcagagtc<br>aggcgggtgcgttggctagagcaagtcggtggacaggtgaaagtagagattgctcttcagatctttaacgagtgataaattcatagctttgtcattatctattatcgagagc<br>aaataagagctatcaaggagcacctaagtgatctgctacttggtaacctttttcagttagtctataaacttctcctttgccgccaagtgactccattagctcatatttaataagct<br>ctgaaatagcacttttccaagaggcctcactacgagcattacccatatctgcaagctgcttattattagtttcaatcatcaatccacccataactggcagacaaattaattctc<br>cattagaatctgatgctacctctattaacagatcttttccaagatcgcttaagcgtggtaattcaggagcaaaagatacttctaaagtattattagtgctttctaagctactgtaa<br>ttttgtataattttctgagatatctgccgactaaacttatgctgaaactcctccaatgactcatatgtttcaattaagcctttttgttgacacaaactacgaaaatcacatagtgcct<br>tgtattgatcctgatcaacgctttctagatatactggagctctggaaaagtaaatcatcgcttgcttgccagacttaagatgttcttctatttcctcaacagttccactaataaatt |

taccagtaggtgttcctaatcttgtccaaaaaacagcaacaagtagatcgcagtctttcaaaatttgcttgtttattatggattgtgctcgtccgctcatatcaggatgggcatg
tgtttcccaccctactggatcaagtactattttacgatcaatagaattaatggtattccactcgtatattatgttccgaataatgttgcgttcttttgcaacatcgctcggtgatgc
aatcataatcctaagaactgtagcgttgtatggcattgctagccccttagagaaaatttcagtagcatttactatttattagctatatttacataaatttttaaactcatcagagat
aatgactcatattttctatacgatcgcaagtaacttcactgccccccttcaccacctccgtatcgcccgagaaagcaaagtgctccgcctgactcaccagcgactctggcaa
cggcagcgtaatccgatagcactccagaaactcacccagcactcccatcgcatgggctttgaccgaatccgagaaccgatcgccctctCCACCTCGACCT
GAATGGAAGCCGGCGGCACCTCGCTAACGGATTCACCACTCCAAGAATTGGAGCCAATCAATTCTTGC
GGAGAACTGTGAATGCGCAAACCAACCCTTGGCAGAACATATCCATCGCGTCCGCCATCTCCAGCAGC
CGCACGCGGCGCATCTCGGGCAGCGTTGGGTCCTGGCCACGGGTGCGCATGATCGTGCTCCTGTCGTT
GAGGACCCGGCTAGGCTGGCGGGGTTGCCTTACTGGTTAGCAGAATGAATCACCGATACGCGAGCGAA
CGTGAAGCGACTGCTGCTGCAAAACGTCTGCGACCTGAGCAACAACATGAATGGTCTTCGGTTTCCGT
GTTTCGTAAAGTCTGGAAACGCGGAAGTCAGCGCCCTGCACCATTATGTTCCGGATCTGCATCGCAGG
ATGCTGCTGGCTACCCTGTGGAACACCTACATCTGTATTAACGAAGCGCTGGCATTGACCCTGAGTGAT
TTTTCTCTGGTCCCGCCGCATCCATACCGCCAGTTGTTTACCCTCACAACGTTCCAGTAACCGGGCATG
TTCATCATCAGTAACCCGTATCGTGAGCATCCTCTCTCGTTTCATCGGTATCATTACCCCCATGAACAGA
AATCCCCCTTACACGGAGGCATCAGTGACCAAACAGGAAAAAACCGCCCTTAACATGGCCCGCTTTAT
CAGAAGCCAGACATTAACGCTTCTGGAGAAACTCAACGAGCTGGACGCGGATGAACAGGCAGACATC
TGTGAATCGCTTCACGACCACGCTGATGAGCTTTACCGCAGCTGCCTCGCGCGTTTCGGTGATGACGG
TGAAAACCTCTGACACATGCAGCTCCCGGGACGGTCACAGCTTGTCTGTAAGCGGATGCCGGGAGCA
GACAAGCCCGTCAGGGCGCGTCAGCGGGTGTTGGCGGGTGTCGGGGCGCAGCCATGACCCAGTCACG
TAGCGATAGCGGAGTGTATACTGGCTTAACTATGCGGCATCAGAGCAGATTGTACTGAGAGTGCACCAT
ATGCGGTGTGAAATACCGCACAGATGCGTAAGGAGAAAATACCGCATCAGGCGCTCTTCCGCTTCCTC
GCTCACTGACTCGCTGCGCTCGGTCGTTCGGCTGCGGCGAGCGGTATCAGCTCACTCAAAGGCGGTAA
TACGGTTATCCACAGAATCAGGGGATAACGCAGGAAAGAACATGTGAGCAAAAGGCCAGCAAAAGGC
CAGGAACCGTAAAAAGGCCGCGTTGCTGGCGTTTTTCCATAGGCTCCGCCCCCCTGACGAGCATCACA
AAAATCGACGCTCAAGTCAGAGGTGGCGAAACCCGACAGGACTATAAAGATACCAGGCGTTTCCCCCT
GGAAGCTCCCTCGTGCGCTCTCCTGTTCCGACCCTGCCGCTTACCGGATACCTGTCCGCCTTTCTCCCT
TCGGGAAGCGTGGCGCTTTCTCATAGCTCACGCTGTAGGTATCTCAGTTCGGTGTAGGTCGTTCGCTCC
AAGCTGGGCTGTGTGCACGAACCCCCCGTTCAGCCCGACCGCTGCGCCTTATCCGGTAACTATCGTCTT
GAGTCCAACCCGGTAAGACACGACTTATCGCCACTGGCAGCAGCCACTGGTAACAGGATTAGCAGAG
CGAGGTATGTAGGCGGTGCTACAGAGTTCTTGAAGTGGTGGCCTAACTACGGCTACACTAGAAGGACA
GTATTTGGTATCTGCGCTCTGCTGAAGCCAGTTACCTTCGGAAAAAGAGTTGGTAGCTCTTGATCCGGC
AAACAAACCACCGCTGGTAGCGGTGGTTTTTTTGTTTGCAAGCAGCAGATTACGCGCAGAAAAAAAG
GATCTCAAGAAGATCCTTTGATCTTTTCTACGGGGTCTGACGCTCAGTGGAACGAAAACTCACGTTAA
GGGATTTTGGTCATGAGATTATCAAAAAGGATCTTCACCTAGATCCTTTTAAATTAAAAATGAAGTTTTA
AATCAATCTAAAGTATATATGAGTAAACTTGGTCTGACAGTTACCAATGCTTAATCAGTGAGGCACCTAT
CTCAGCGATCTGTCTATTTCGTTCATCCATAGTTGCCTGACTCCCCGTCGTGTAGATAACTACGATACGG
GAGGGCTTACCATCTGGCCCCAGTGCTGCAATGATACCGCGcGACCCACGCTCACCGGCTCCAGATTTA
TCAGCAATAAACCAGCCAGCCGGAAGGGCCGAGCGCAGAAGTGGTCCTGCAACTTTATCCGCCTCCAT
CCAGTCTATTAATTGTTGCCGGGAAGCTAGAGTAAGTAGTTCGCCAGTTAATAGTTTGCGCAACGTTGT
TGCCATTGCTGCAGGCATCGTGGTGTCACGCTCGTCGTTTGGTATGGCTTCATTCAGCTCCGGTTCCCA
ACGATCAAGGCGAGTTACATGATCCCCCATGTTGTGCAAAAAAGCGGTTAGCTCCTTCGGTCCTCCGAT
CGTTGTCAGAAGTAAGTTGGCCGCAGTGTTATCACTCATGGTTATGGCAGCACTGCATAATTCTCTTACT
GTCATGCCATCCGTAAGATGCTTTTCTGTGACTGGTGAGTACTCAACCAAGTCATTCTGAGAATAGTGT
ATGCGGCGACCGAGTTGCTCTTGCCCGGCGTCAACACGGGATAATACCGCGCCACATAGCAGAACTTT
AAAAGTGCTCATCATTGGAAAACGTTCTTCGGGGCGAAAACTCTCAAGGATCTTACCGCTGTTGAGAT
CCAGTTCGATGTAACCCACTCGTGCACCCAACTGATCTTCAGCATCTTTTACTTTCACCAGCGTTTCTG
GGTGAGCAAAAACAGGAAGGCAAAATGCCGCAAAAAAGGGAATAAGGGCGACACGGAAATGTTGAA
TACTCATACTCTTCCTTTTTCAATATTATTGAAGCATTTATCAGGGTTATTGTCTCATGAGCGGATACATAT
TTGAATGTATTTAGAAAAATAAACAAATAGGGGTTCCGCG

| | |
|---|---|
| pSEL-68rpsl | ttagaaaaactcatcgagcatcaaatgaaactgcaatttattcatatcaggattatcaataccatatttttgaaaaagccgtttctgtaatgaaggagaaaactcaccgaggc<br>agttccataggatggcaagatcctggtatcggtctgcgattccgactcgtccaacatcaatacaacctattaatttcccctcgtcaaaaataaggttatcaagtgagaaatc<br>accatgagtgacgactgaatccggtgagaatggcaaaagcttatgcatttctttccagacttgttcaacaggccagccattacgctcgtcatcaaaatcactcgcatcaac<br>caaaccgttattcattcgtgattgcgcctgagcgagacgaaatacgcgatcgctgttaaaaggacaattacaaacaggaatcgaatgcaaccggcgcaggaacactgc<br>cagcgcatcaacaatattttcacctgaatcaggatattcttctaatacctggaatgctgttttcccggggatcgcagtggtgagtaaccatgcatcatcaggagtacggata<br>aaatgcttgatggtcggaagaggcataaattccgtcagccagtttagtctgaccatctcatctgtaacatcattggcaacgctacctttgccatgtttcagaaacaactctgg<br>cgcatcgggcttcccatacaatcgatagattgtcgcacctgattgcccgacattatcgcgagcccatttatacccatataaatcagcatccatgttggaatttaatcgcggc<br>ctggagcaagacgtttcccgttgaatatggctcataacaccccttgtattactgtttatgtaagcagacagttttattgttcatgatgatatatttttatcttgtgcaatgtaacatc<br>agagattttgagacacaacgtggctttgttgaataaatcgaacttttgctgagttgaaggatcagGAAAAGTCTGAAAGTTCTTTACAAAACTC<br>AATCTGCTTGTTAGATTTTACTCACGAGGCTATTAAGTCTCGTAAATAGTTCAACTAAGGACTCATCGCA<br>AAatgccaactatccagcagctaattcgtagcgaacgctcgaaggtacagaagaaaactaaatcccctgccctcaagcaatgtccccaacggcggggagtctgcac<br>tagggtttacaccaccacccccaaaaagcccaactccgccctccggaaagtggcccgggtacgcctcacctccgggtttgaagtaactgcctatatccctggcattggc<br>cacaacctgcaagaacactccgtagtactaatccggggcggtcgggtaaaagatttgcctggggttcgctaccatattgtgcggggcacgttggacgccaccggagtt<br>aaagaccgcaaacagggtcgctccaaatacggcaccaaacgggaaaaagcgaagaaataaccatgatcgagcgatcggcctcgggggcgttgcagttcatccgca<br>tgatctcgcccctgtcttggatgcccattgccgttatggcctttggcattggcgatctgcccgtttattttctgctagcgatcgcagcggtctggccaattctactcagcacca<br>gcagcggtactgcggctgttaaccacaagctcttacttctagctcgtagcctttgtgccactcgcagtgaaacgattcggcgcatcgtgattcctgcgatcgtgccgcaaa<br>tccttgtgggtgttcgactcgcgatcggcacgatctggattgtgctcgtccccgctgagatgctgggcgttagctcaggcctcggttacttcattcttgatacccgcgatcg<br>cattgcctataacgaacttaccgctgttttattagcgatcggcatcattggctgcgccctagattggagcctgcaattcctgcaaaagtactggcaacccagtcgctaaacc<br>tcttggcatccattctccaaacctctcaaaggcgtctacagggttgagcttgtggggaggggcggctatcctcgcccgccaaaagcccgatcgtcatcactggcaagcc<br>cagcaacgacagatggaactgaactcagtccaagggcagctcgatgccctggtcgtcaaagcaccctacgcaggcagggttaggcgggtacgctggctagagcag<br>gtgggcggacaggtcaaggtggagattgctcttcaagtctcaaacactccctaagttttgaccttgattacagtggccccatctagtttttgtatatacaatgactaggtgcg<br>tatcacctacgatccagtcaaacgcgacaagacactgcttgagcgaggactcgacttcgagagtgcgatcgaagtctttgcagggctcacgctagaagtcgaagacac<br>ccgacgcgactatggcgaaaagcgagtgctctgcgtcggcttccttgatggccgcatggtgatggttggctatacccctcgcggtcgaacacgtcacatcttttcgatga<br>ggaaatgcaatgcccgagaaattcgccgctacacgccatacttcgagcaagcctgatgacgacttgcccgagttgaccgatgaaatgctcagtcgggcggtgctcaa<br>gcagggcggtaagcgtatcggcagaccccgatcgcaatctccaaaagtagcgatcagcctgcgcttagaagctgaagtcctgcagcgctggcgtgacagtggccct<br>ggctggcaaactcgtatggctgaattgctctcgaaagaaacccccctaagtctttgatcgctcaagcccaacggcaggacaagctacaccagtggcaaacccagcagc<br>gacagatggaacttaactcagtccaagcgcagctcgatgcactggtcgtcaaagcaccttactccggcagagtcaggcgggtgcgttggctagagcaagtcggtgga<br>caggtgaaagtagagattgctcttcagatctttaacgagtgataaattcatagctttgtcattatctattatcgagagcaaataagagctatcaaggagcacctaagtgatct<br>gctacttggtaacctttttcagttagtctataaacttctcctttgccgccaagtgactccattagctcatatttaataagctctgaaatagcacttttccaagaggcctcactacg<br>agcattacccatatctgcaagctgcttattattagtttcaatcatcaatccacccataactggcagacaaattaattctccattagaatctgatgctacctctattaacagatcttt<br>tccaagatcgcttaagcgtggtaattcaggagcaaaagatacttctaaagtattattagtgctttctaagctactgtaattttgtataattttctgagatatctgccgactaaactt<br>atgctgaaactcctccaatgactcatatgtttcaattaagcctttttgttgacacaaactacgaaaatcacatagtgccttgtattgatcctgatcaacgctttctagatatactg<br>gagctctggaaaagtaaatcatcgcttgcttgccagacttaagatgttcttctatttcctcaacagttccactaataaatttaccagtaggtgttcctaatcttgtccaaaaaac<br>agcaacaagtagatcgcagtctttcaaaatttgcttgtttattatggattgtgctcgtccgctcatatcaggatgggcatgtgtttcccaccctactggatcaagtactattttac<br>gatcaatagaattaatggtattccactcgtatattatgttccgaataatgttgcgttcttttgcaacatcgctcggtgatgcaatcataatcctaagaactgtagcgttgtatggc<br>attgctagccccttagagaaaatttcagtagcatttactatttattagctatatttacataaatttttttaaactcatcagagataatgactcatattttctatacgatcgcaagtaact<br>tcactgccccccttcaccacctccgtatcgcccgagaaagcaaagtgctccgcctgactcaccagcgactctggcaacggcagcgtaatccgatagcactccagaaact<br>cacccagcactcccatcgcatgggctttgaccgaatccgagaaccgatcgccctctCCACCTCGACCTGAATGGAAGCCGGCGGCACC<br>TCGCTAACGGATTCACCACTCCAAGAATTGGAGCCAATCAATTCTTGCGGAGAACTGTGAATGCGCAA<br>ACCAACCCTTGGCAGAACATATCCATCGCGTCCGCCATCTCCAGCAGCCGCACGCGGCGCATCTCGGG<br>CAGCGTTGGGTCCTGGCCACGGGTGCGCATGATCGTGCTCCTGTCGTTGAGGACCCGGCTAGGCTGGC<br>GGGGTTGCCTTACTGGTTAGCAGAATGAATCACCGATACGCGAGCGAACGTGAAGCGACTGCTGCTGC<br>AAAACGTCTGCGACCTGAGCAACAACATGAATGGTCTTCGGTTTCCGTGTTTCGTAAAGTCTGGAAAC<br>GCGGAAGTCAGCGCCCTGCACCATTATGTTCCGGATCTGCATCGCAGGATGCTGCTGGCTACCCTGTGG<br>AACACCTACATCTGTATTAACGAAGCGCTGGCATTGACCCTGAGTGATTTTTCTCTGGTCCCGCCGCAT<br>CCATACCGCCAGTTGTTTACCCTCACAACGTTCCAGTAACCGGGCATGTTCATCATCAGTAACCCGTATC<br>GTGAGCATCCTCTCTCGTTTCATCGGTATCATTACCCCCATGAACAGAAATCCCCCTTACACGGAGGCAT<br>CAGTGACCAAACAGGAAAAAACCGCCCTTAACATGGCCCGCTTTATCAGAAGCCAGACATTAACGCTT<br>CTGGAGAAACTCAACGAGCTGGACGCGGATGAACAGGCAGACATCTGTGAATCGCTTCACGACCACG |

| | |
|---|---|
| | CTGATGAGCTTTACCGCAGCTGCCTCGCGCGTTTCGGTGATGACGGTGAAAACCTCTGACACATGCAG<br>CTCCCGGGACGGTCACAGCTTGTCTGTAAGCGGATGCCGGGAGCAGACAAGCCCGTCAGGGCGCGTC<br>AGCGGGTGTTGGCGGGTGTCGGGGCGCAGCCATGACCCAGTCACGTAGCGATAGCGGAGTGTATACTG<br>GCTTAACTATGCGGCATCAGAGCAGATTGTACTGAGAGTGCACCATATGCGGTGTGAAATACCGCACAG<br>ATGCGTAAGGAGAAAATACCGCATCAGGCGCTCTTCCGCTTCCTCGCTCACTGACTCGCTGCGCTCGGT<br>CGTTCGGCTGCGGCGAGCGGTATCAGCTCACTCAAAGGCGGTAATACGGTTATCCACAGAATCAGGGG<br>ATAACGCAGGAAAGAACATGTGAGCAAAAGGCCAGCAAAAGGCCAGGAACCGTAAAAAGGCCGCGT<br>TGCTGGCGTTTTTCCATAGGCTCCGCCCCCCTGACGAGCATCACAAAAATCGACGCTCAAGTCAGAGG<br>TGGCGAAACCCGACAGGACTATAAAGATACCAGGCGTTTCCCCCTGGAAGCTCCCTCGTGCGCTCTCC<br>TGTTCCGACCCTGCCGCTTACCGGATACCTGTCCGCCTTTCTCCCTTCGGGAAGCGTGGCGCTTTCTCA<br>TAGCTCACGCTGTAGGTATCTCAGTTCGGTGTAGGTCGTTCGCTCCAAGCTGGGCTGTGTGCACGAACC<br>CCCCGTTCAGCCCGACCGCTGCGCCTTATCCGGTAACTATCGTCTTGAGTCCAACCCGGTAAGACACG<br>ACTTATCGCCACTGGCAGCAGCCACTGGTAACAGGATTAGCAGAGCGAGGTATGTAGGCGGTGCTACA<br>GAGTTCTTGAAGTGGTGGCCTAACTACGGCTACACTAGAAGGACAGTATTTGGTATCTGCGCTCTGCTG<br>AAGCCAGTTACCTTCGGAAAAAGAGTTGGTAGCTCTTGATCCGGCAAACAAACCACCGCTGGTAGCG<br>GTGGTTTTTTTGTTTGCAAGCAGCAGATTACGCGCAGAAAAAAAGGATCTCAAGAAGATCCTTTGATC<br>TTTTCTACGGGGTCTGACGCTCAGTGGAACGAAAACTCACGTTAAGGGATTTTGGTCATGAGATTATCA<br>AAAAGGATCTTCACCTAGATCCTTTTAAATTAAAAATGAAGTTTTAAATCAATCTAAAGTATATATGAGT<br>AAACTTGGTCTGACAGTTACCAATGCTTAATCAGTGAGGCACCTATCTCAGCGATCTGTCTATTTCGTTC<br>ATCCATAGTTGCCTGACTCCCCGTCGTGTAGATAACTACGATACGGGAGGGCTTACCATCTGGCCCCAG<br>TGCTGCAATGATACCGCGcGACCCACGCTCACCGGCTCCAGATTTATCAGCAATAAACCAGCCAGCCGG<br>AAGGGCCGAGCGCAGAAGTGGTCCTGCAACTTTATCCGCCTCCATCCAGTCTATTAATTGTTGCCGGGA<br>AGCTAGAGTAAGTAGTTCGCCAGTTAATAGTTTGCGCAACGTTGTTGCCATTGCTGCAGGCATCGTGGT<br>GTCACGCTCGTCGTTTGGTATGGCTTCATTCAGCTCCGGTTCCCAACGATCAAGGCGAGTTACATGATC<br>CCCCATGTTGTGCAAAAAAGCGGTTAGCTCCTTCGGTCCTCCGATCGTTGTCAGAAGTAAGTTGGCCG<br>CAGTGTTATCACTCATGGTTATGGCAGCACTGCATAATTCTCTTACTGTCATGCCATCCGTAAGATGCTTT<br>TCTGTGACTGGTGAGTACTCAACCAAGTCATTCTGAGAATAGTGTATGCGGCGACCGAGTTGCTCTTGC<br>CCGGCGTCAACACGGGATAATACCGCGCCACATAGCAGAACTTTAAAAGTGCTCATCATTGGAAAACG<br>TTCTTCGGGGCGAAAACTCTCAAGGATCTTACCGCTGTTGAGATCCAGTTCGATGTAACCCACTCGTGC<br>ACCCAACTGATCTTCAGCATCTTTTACTTTCACCAGCGTTTCTGGGTGAGCAAAAACAGGAAGGCAAA<br>ATGCCGCAAAAAAGGGAATAAGGGCGACACGGAAATGTTGAATACTCATACTCTTCCTTTTTCAATATT<br>ATTGAAGCATTTATCAGGGTTATTGTCTCATGAGCGGATACATATTTGAATGTATTTAGAAAAATAAACA<br>AATAGGGGTTCCGCG |
| pBR322-rpslm | aggaggagggccactcctcgtcaaatcagggcgagtggtcgtcaatccccaagccgaaggatttagccgagcctttgagatcgaagcagctccgcgcagtgcgatc<br>ggtctgatgccggatggccgcttggttctagtggctgcccacgagcaaaaccaaggccaaggccccaccctgcctcaaatggctgcgattatgcagcagctcggcgt<br>cgttgatgccctcaactttgatggcggcagctccacttctctaatcgtcaatggtcagctcgtcaatcgggctcgaggcagtgctgcccgggttcacaacgggctcggg<br>gtcttcctggggccgactacgccggccagtctgcgatgaaggttggcgatcgcgatcaacggccttggggacagttcgaaattcttgcccaaggctctggctatcaggt<br>caagcgtttggaggtgctgccagggcagcagttgagcctacagcgtcaccagcagcggcaggaacattggctagtcgtgcaagggctggcacgggtgcaactggg<br>cgatcgccagttttctctgcgcgtagggcaatcgctcgatatcgcgatcggggaatggcatcgcctgcaaaatccgggggacgagcttttagccctgattgaggtgcaa<br>atgggggcctatctcggcgaagatgacattgagcgccgcgccgatgactacggacgctgcccaacgtcgtgataatggctgtggacccagtctgttggccgctcgttc<br>cagagaggctacctggggcgatcgccttctcttcagctgtagcttgaccaagtgcatcaaccctcggtaaagtaagagattgttgtcttttccaagagcagccgaaccgg<br>ctttctgaaatgcctacaatccagcagctgattcgcgacgaacgcgaaaaaattaccaagaagacaaaatctcccgcgctgaaaaactgcccgcagcggcgtggcgtt<br>tgcactcgcgtctacaccacgaccccgagaaagcccaactcggccttgcggaaagtggcacgggttcgcttgacctcaggttttgaagtcactgcctatatccctggca<br>ttggtcacaacctgcaagagcactcggttgtgatgattcgcggcggtcgggtcaaagacctacccggtgtgcgctatcacatcatccgcggcacccttgataccgccg<br>gtgtcaaagaccgtcgccaaagccgttccaaatacggcgcgaagcgtccgaaggcctaggtccctgggtgagccttgtctgcgttgatcgtggcaacgtcaacactgc<br>aagctgctgggcagttgaccttgctcgttccagatgtgattaactctcacccctgcgatcgcccctttttcgcttcctctttgcgttgtagcccttgcaagacttctgagattca<br>ctgatgtcccgtcgtacctctgctcaaaagcgttcggttaacccggatccaaaattcaacagccgcttggcctccatgatggtggctcgcctgatggactcgggcaaaaa<br>gtccctggctttccgcatcctctactcagccttcgatctgatccaagagcggaccggaaacgaccccttggaactcttcgaacaagccgtgcgcaatgccaccccccttg<br>gtggaagtgcgagctcgtcgggtaggtggtgcaacctaccaagtcccgatggaagttcgctcggagcgcggtaccgccatggccctgcgctggttggttcagtattcc |

| | |
|---|---|
| | cgtcagcgccctggcaagtccatggccatcaagctggcaaacgagctgatggatgcagccaacgaaaccggaagttcagtgcgtaaacgcgaagaaacccacaaa<br>atggccgaagcgaacaaggcttttgcccactaccgctactagCCACCTCGACCTGAATGGAAGCCGGCGGCACCTCGCTAACG<br>GATTCACCACTCCAAGAATTGGAGCCAATCAATTCTTGCGGAGAACTGTGAATGCGCAAACCAACCCT<br>TGGCAGAACATATCCATCGCGTCCGCCATCTCCAGCAGCCGCACGCGGCGCATCTCGGGCAGCGTTGG<br>GTCCTGGCCACGGGTGCGCATGATCGTGCTCCTGTCGTTGAGGACCCGGCTAGGCTGGCGGGGTTGCC<br>TTACTGGTTAGCAGAATGAATCACCGATACGCGAGCGAACGTGAAGCGACTGCTGCTGCAAAACGTCT<br>GCGACCTGAGCAACAACATGAATGGTCTTCGGTTTCCGTGTTTCGTAAAGTCTGGAAACGCGGAAGTC<br>AGCGCCCTGCACCATTATGTTCCGGATCTGCATCGCAGGATGCTGCTGGCTACCCTGTGGAACACCTAC<br>ATCTGTATTAACGAAGCGCTGGCATTGACCCTGAGTGATTTTTCTCTGGTCCCGCCGCATCCATACCGCC<br>AGTTGTTTACCCTCACAACGTTCCAGTAACCGGGCATGTTCATCATCAGTAACCCGTATCGTGAGCATC<br>CTCTCTCGTTTCATCGGTATCATTACCCCCATGAACAGAAATCCCCCTTACACGGAGGCATCAGTGACC<br>AAACAGGAAAAAACCGCCCTTAACATGGCCCGCTTTATCAGAAGCCAGACATTAACGCTTCTGGAGAA<br>ACTCAACGAGCTGGACGCGGATGAACAGGCAGACATCTGTGAATCGCTTCACGACCACGCTGATGAG<br>CTTTACCGCAGCTGCCTCGCGCGTTTCGGTGATGACGGTGAAAACCTCTGACACATGCAGCTCCCGGG<br>ACGGTCACAGCTTGTCTGTAAGCGGATGCCGGGAGCAGACAAGCCCGTCAGGGCGCGTCAGCGGGTG<br>TTGGCGGGTGTCGGGGCGCAGCCATGACCCAGTCACGTAGCGATAGCGGAGTGTATACTGGCTTAACT<br>ATGCGGCATCAGAGCAGATTGTACTGAGAGTGCACCATATGCGGTGTGAAATACCGCACAGATGCGTA<br>AGGAGAAAATACCGCATCAGGCGCTCTTCCGCTTCCTCGCTCACTGACTCGCTGCGCTCGGTCGTTCG<br>GCTGCGGCGAGCGGTATCAGCTCACTCAAAGGCGGTAATACGGTTATCCACAGAATCAGGGGATAACG<br>CAGGAAAGAACATGTGAGCAAAAGGCCAGCAAAAGGCCAGGAACCGTAAAAAGGCCGCGTTGCTGG<br>CGTTTTTCCATAGGCTCCGCCCCCCTGACGAGCATCACAAAAATCGACGCTCAAGTCAGAGGTGGCGA<br>AACCCGACAGGACTATAAAGATACCAGGCGTTTCCCCCTGGAAGCTCCCTCGTGCGCTCTCCTGTTCC<br>GACCCTGCCGCTTACCGGATACCTGTCCGCCTTTCTCCCTTCGGGAAGCGTGGCGCTTTCTCATAGCTC<br>ACGCTGTAGGTATCTCAGTTCGGTGTAGGTCGTTCGCTCCAAGCTGGGCTGTGTGCACGAACCCCCCG<br>TTCAGCCCGACCGCTGCGCCTTATCCGGTAACTATCGTCTTGAGTCCAACCCGGTAAGACACGACTTAT<br>CGCCACTGGCAGCAGCCACTGGTAACAGGATTAGCAGAGCGAGGTATGTAGGCGGTGCTACAGAGTTC<br>TTGAAGTGGTGGCCTAACTACGGCTACACTAGAAGGACAGTATTTGGTATCTGCGCTCTGCTGAAGCC<br>AGTTACCTTCGGAAAAAGAGTTGGTAGCTCTTGATCCGGCAAACAAACCACCGCTGGTAGCGGTGGTT<br>TTTTTGTTTGCAAGCAGCAGATTACGCGCAGAAAAAAAGGATCTCAAGAAGATCCTTTGATCTTTTCTA<br>CGGGGTCTGACGCTCAGTGGAACGAAAACTCACGTTAAGGGATTTTGGTCATGAGATTATCAAAAAGG<br>ATCTTCACCTAGATCCTTTTAAATTAAAAATGAAGTTTTAAATCAATCTAAAGTATATATGAGTAAACTTG<br>GTCTGACAGTTACCAATGCTTAATCAGTGAGGCACCTATCTCAGCGATCTGTCTATTTCGTTCATCCATA<br>GTTGCCTGACTCCCCGTCGTGTAGATAACTACGATACGGGAGGGCTTACCATCTGGCCCCAGTGCTGCA<br>ATGATACCGCGcGACCCACGCTCACCGGCTCCAGATTTATCAGCAATAAACCAGCCAGCCGGAAGGGC<br>CGAGCGCAGAAGTGGTCCTGCAACTTTATCCGCCTCCATCCAGTCTATTAATTGTTGCCGGGAAGCTAG<br>AGTAAGTAGTTCGCCAGTTAATAGTTTGCGCAACGTTGTTGCCATTGCTGCAGGCATCGTGGTGTCACG<br>CTCGTCGTTTGGTATGGCTTCATTCAGCTCCGGTTCCCAACGATCAAGGCGAGTTACATGATCCCCCAT<br>GTTGTGCAAAAAAGCGGTTAGCTCCTTCGGTCCTCCGATCGTTGTCAGAAGTAAGTTGGCCGCAGTGT<br>TATCACTCATGGTTATGGCAGCACTGCATAATTCTCTTACTGTCATGCCATCCGTAAGATGCTTTTCTGTG<br>ACTGGTGAGTACTCAACCAAGTCATTCTGAGAATAGTGTATGCGGCGACCGAGTTGCTCTTGCCCGGC<br>GTCAACACGGGATAATACCGCGCCACATAGCAGAACTTTAAAAGTGCTCATCATTGGAAAACGTTCTTC<br>GGGGCGAAAACTCTCAAGGATCTTACCGCTGTTGAGATCCAGTTCGATGTAACCCACTCGTGCACCCA<br>ACTGATCTTCAGCATCTTTTACTTTCACCAGCGTTTCTGGGTGAGCAAAAACAGGAAGGCAAAATGCC<br>GCAAAAAAGGGAATAAGGGCGACACGGAAATGTTGAATACTCATACTCTTCCTTTTTCAATATTATTGA<br>AGCATTTATCAGGGTTATTGTCTCATGAGCGGATACATATTTGAATGTATTTAGAAAAATAAACAAATAG<br>GGGTTCCGCG |
| pB<br>R-<br>pil- | TTACCAATGCTTAATCAGTGAGGCACCTATCTCAGCGATCTGTCTATTTCGTTCATCCATAGTTGCCTGA<br>CTCCCCGTCGTGTAGATAACTACGATACGGGAGGGCTTACCATCTGGCCCCAGTGCTGCAATGATACCG<br>CGcGACCCACGCTCACCGGCTCCAGATTTATCAGCAATAAACCAGCCAGCCGGAAGGGCCGAGCGCAG |

| | |
|---|---|
| km | AAGTGGTCCTGCAACTTTATCCGCCTCCATCCAGTCTATTAATTGTTGCCGGGAAGCTAGAGTAAGTAG<br>TTCGCCAGTTAATAGTTTGCGCAACGTTGTTGCCATTGCTGCAGGCATCGTGGTGTCACGCTCGTCGTT<br>TGGTATGGCTTCATTCAGCTCCGGTTCCCAACGATCAAGGCGAGTTACATGATCCCCCATGTTGTGCAA<br>AAAAGCGGTTAGCTCCTTCGGTCCTCCGATCGTTGTCAGAAGTAAGTTGGCCGCAGTGTTATCACTCAT<br>GGTTATGGCAGCACTGCATAATTCTCTTACTGTCATGCCATCCGTAAGATGCTTTTCTGTGACTGGTGAG<br>TACTCAACCAAGTCATTCTGAGAATAGTGTATGCGGCGACCGAGTTGCTCTTGCCCGGCGTCAACACG<br>GGATAATACCGCGCCACATAGCAGAACTTTAAAAGTGCTCATCATTGGAAAACGTTCTTCGGGGCGAA<br>AACTCTCAAGGATCTTACCGCTGTTGAGATCCAGTTCGATGTAACCCACTCGTGCACCCAACTGATCTT<br>CAGCATCTTTTACTTTCACCAGCGTTTCTGGGTGAGCAAAAACAGGAAGGCAAAATGCCGCAAAAAA<br>GGGAATAAGGGCGACACGGAAATGTTGAATACTCATACTCTTCCTTTTTCAATATTATTGAAGCATTTAT<br>CAGGGTTATTGTCTCATGAGCGGATACATATTTGAATGTATTTAGAAAAATAAACAAATAGGGGTTCCGC<br>Gcttagcgcttactctctcgggctttgctagcctgcagcggatcagccgtggtgaaatgctcgaggtgctgcgccaagactacatccgcaccgcccgtgccaaaggct<br>tgccggagcagcgcgtcatctacgtccacgctctacgcaatgcgatcaatcccctgattacgctcttgggctttgagttcgcgaccctgctcagcggcgcttttattgctg<br>aatatttctttaactggccggggctaggccgcttaattttgcaagccgtttttgcgcaggatctctacttggtaatggccagcttgatgatgggtgctgtgatgctgattctgg<br>gcaatctgctcgcagatctgctgctgcgctgggtcgatccccgcattcgcctggatgatctgaactaattagaaaaactcatcgagcatcaaatgaaactgcaatttattca<br>tatcaggattatcaataccatatttttgaaaaagccgtttctgtaatgaaggagaaaactcaccgaggcagttccataggatggcaagatcctggtatcggtctgcgattcc<br>gactcgtccaacatcaatacaacctattaatttcccctcgtcaaaaataaggttatcaagtgagaaatcaccatgagtgacgactgaatccggtgagaatggcaaaagctt<br>atgcatttctttccagacttgttcaacaggccagccattacgctcgtcatcaaaatcactcgcatcaaccaaaccgttattcattcgtgattgcgcctgagcgagacgaaata<br>cgcgatcgctgttaaaaggacaattacaaacaggaatcgaatgcaaccggcgcaggaacactgccagcgcatcaacaatattttcacctgaatcaggatattcttctaat<br>acctggaatgctgttttcccggggatcgcagtggtgagtaaccatgcatcatcaggagtacggataaaatgcttgatggtcggaagaggcataaattccgtcagccagtt<br>tagtctgaccatctcatctgtaacatcattggcaacgctacctttgccatgtttcagaaacaactctggcgcatcgggcttcccatacaatcgatagattgtcgcacctgatt<br>gcccgacattatcgcgagcccatttatacccatataaatcagcatccatgttggaatttaatcgcggcctggagcaagacgtttcccgttgaatatggctcataacacccct<br>tgtattactgtttatgtaagcagacagttttattgttcatgatgatatatttttatcttgtgcaatgtaacatcagagattttgagacacaacgtggctttgttgaataaatcgaactt<br>ttgctgagttgaaggatcagtagctagaggatcgatcctttttaacccatcacatatacctgccgttcactattatttagtgaaatgagatattatgatattttctgaattgtgatt<br>aaaaaggcaactttatgcccatgcaacagaaactataaaaaatacagagaatgaaaagaaacagatagattttttagttctttaggcccgtagtctgcaaatccttttatgat<br>tttctatcaaacaaaagaggaaaatagaccagttgcaatccaaacgagagtctaatagaatgaggtcgaaagaagtggcggtctgagaacgaccgtaattgataatgcg<br>ggctcagtgatccagcccccaaatgacagtgtctttaaccccagtaacttaccagcagggattaacccagcgaatcccttgttggccttgccgaatcaattgaccaataat<br>ttccttggtgctttatctgcatcaattatcaacaacaaaactaagttgctgactgaccccaccttaatcattcaggaaggagaaagaagcgaagtcaaactgactagtcagg<br>ttgttcaaaagattgaatcagagactacaaccaacggcagtggcccgccaaccgttagccgaaccattgaccttgcagatgttggtctacagttaaccatcaacattgaa<br>cgaattgatgacaatggcttcattactctCCACCTCGACCTGAATGGAAGCCGGCGGCACCTCGCTAACGGATTCACCAC<br>TCCAAGAATTGGAGCCAATCAATTCTTGCGGAGAACTGTGAATGCGCAAACCAACCCTTGGCAGAACA<br>TATCCATCGCGTCCGCCATCTCCAGCAGCCGCACGCGGCGCATCTCGGGCAGCGTTGGGTCCTGGCCA<br>CGGGTGCGCATGATCGTGCTCCTGTCGTTGAGGACCCGGCTAGGCTGGCGGGGTTGCCTTACTGGTTA<br>GCAGAATGAATCACCGATACGCGAGCGAACGTGAAGCGACTGCTGCTGCAAAACGTCTGCGACCTGA<br>GCAACAACATGAATGGTCTTCGGTTTCCGTGTTTCGTAAAGTCTGGAAACGCGGAAGTCAGCGCCCTG<br>CACCATTATGTTCCGGATCTGCATCGCAGGATGCTGCTGGCTACCCTGTGGAACACCTACATCTGTATTA<br>ACGAAGCGCTGGCATTGACCCTGAGTGATTTTTCTCTGGTCCCGCCGCATCCATACCGCCAGTTGTTTA<br>CCCTCACAACGTTCCAGTAACCGGGCATGTTCATCATCAGTAACCCGTATCGTGAGCATCCTCTCTCGT<br>TTCATCGGTATCATTACCCCCATGAACAGAAATCCCCCTTACACGGAGGCATCAGTGACCAAACAGGA<br>AAAAACCGCCCTTAACATGGCCCGCTTTATCAGAAGCCAGACATTAACGCTTCTGGAGAAACTCAACG<br>AGCTGGACGCGGATGAACAGGCAGACATCTGTGAATCGCTTCACGACCACGCTGATGAGCTTTACCGC<br>AGCTGCCTCGCGCGTTTCGGTGATGACGGTGAAAACCTCTGACACATGCAGCTCCCGGACGGTCACA<br>GCTTGTCTGTAAGCGGATGCCGGGAGCAGACAAGCCCGTCAGGGCGCGTCAGCGGGTGTTGGCGGGT<br>GTCGGGGCGCAGCCATGACCCAGTCACGTAGCGATAGCGGAGTGTATACTGGCTTAACTATGCGGCATC<br>AGAGCAGATTGTACTGAGAGTGCACCATATGCGGTGTGAAATACCGCACAGATGCGTAAGGAGAAAAT<br>ACCGCATCAGGCGCTCTTCCGCTTCCTCGCTCACTGACTCGCTGCGCTCGGTCGTTCGGCTGCGGCGA<br>GCGGTATCAGCTCACTCAAAGGCGGTAATACGGTTATCCACAGAATCAGGGGATAACGCAGGAAAGAA<br>CATGTGAGCAAAAGGCCAGCAAAAGGCCAGGAACCGTAAAAAGGCCGCGTTGCTGGCGTTTTTCCAT<br>AGGCTCCGCCCCCCTGACGAGCATCACAAAAATCGACGCTCAAGTCAGAGGTGGCGAAACCCGACAG |

| | |
|---|---|
| | GACTATAAAGATACCAGGCGTTTCCCCCTGGAAGCTCCCTCGTGCGCTCTCCTGTTCCGACCCTGCCGC TTACCGGATACCTGTCCGCCTTTCTCCCTTCGGGAAGCGTGGCGCTTTCTCATAGCTCACGCTGTAGGT ATCTCAGTTCGGTGTAGGTCGTTCGCTCCAAGCTGGGCTGTGTGCACGAACCCCCCGTTCAGCCCGAC CGCTGCGCCTTATCCGGTAACTATCGTCTTGAGTCCAACCCGGTAAGACACGACTTATCGCCACTGGCA GCAGCCACTGGTAACAGGATTAGCAGAGCGAGGTATGTAGGCGGTGCTACAGAGTTCTTGAAGTGGTG GCCTAACTACGGCTACACTAGAAGGACAGTATTTGGTATCTGCGCTCTGCTGAAGCCAGTTACCTTCGG AAAAAGAGTTGGTAGCTCTTGATCCGGCAAACAAACCACCGCTGGTAGCGGTGGTTTTTTTGTTTGCA AGCAGCAGATTACGCGCAGAAAAAAAGGATCTCAAGAAGATCCTTTGATCTTTTCTACGGGGTCTGAC GCTCAGTGGAACGAAAACTCACGTTAAGGGATTTTGGTCATGAGATTATCAAAAAGGATCTTCACCTA GATCCTTTTAAATTAAAAATGAAGTTTTAAATCAATCTAAAGTATATATGAGTAAACTTGGTCTGACAG |
| pBR-pil-km-sepT2 | TTACCAATGCTTAATCAGTGAGGCACCTATCTCAGCGATCTGTCTATTTCGTTCATCCATAGTTGCCTGA CTCCCCGTCGTGTAGATAACTACGATACGGGAGGGCTTACCATCTGGCCCCAGTGCTGCAATGATACCG CGcGACCCACGCTCACCGGCTCCAGATTTATCAGCAATAAACCAGCCAGCCGGAAGGGCCGAGCGCAG AAGTGGTCCTGCAACTTTATCCGCCTCCATCCAGTCTATTAATTGTTGCCGGGAAGCTAGAGTAAGTAG TTCGCCAGTTAATAGTTTGCGCAACGTTGTTGCCATTGCTGCAGGCATCGTGGTGTCACGCTCGTCGTT TGGTATGGCTTCATTCAGCTCCGGTTCCCAACGATCAAGGCGAGTTACATGATCCCCCATGTTGTGCAA AAAAGCGGTTAGCTCCTTCGGTCCTCCGATCGTTGTCAGAAGTAAGTTGGCCGCAGTGTTATCACTCAT GGTTATGGCAGCACTGCATAATTCTCTTACTGTCATGCCATCCGTAAGATGCTTTTCTGTGACTGGTGAG TACTCAACCAAGTCATTCTGAGAATAGTGTATGCGGCGACCGAGTTGCTCTTGCCCGGCGTCAACACG GGATAATACCGCGCCACATAGCAGAACTTTAAAAGTGCTCATCATTGGAAAACGTTCTTCGGGGCGAA AACTCTCAAGGATCTTACCGCTGTTGAGATCCAGTTCGATGTAACCCACTCGTGCACCCAACTGATCTT CAGCATCTTTTACTTTCACCAGCGTTTCTGGGTGAGCAAAAACAGGAAGGCAAAATGCCGCAAAAAA GGGAATAAGGGCGACACGGAAATGTTGAATACTCATACTCTTCCTTTTTCAATATTATTGAAGCATTTAT CAGGGTTATTGTCTCATGAGCGGATACATATTTGAATGTATTTAGAAAAATAAACAAATAGGGGTTCCGC Gcttagcgcttactctctcgggctttgctagcctgcagcggatcagccgtggtgaaatgctcgaggtgctgcgccaagactacatccgcaccgcccgtgccaaaggct tgccggagcagcgcgtcatctacgtccacgctctacgcaatgcgatcaatcccctgattacgctcttgggctttgagttcgcgaccctgctcagcggcgcttttattgctg aatatttctttaactggccggggctaggccgcttaattttgcaagccgtttttgcgcaggatctctacttggtaatggccagcttgatgatgggtgctgtgatgctgattctgg gcaatctgctcgcagatctgctgctgcgctgggtcgatccccgcattcgcctggatgatctgaactaattagaaaaactcatcgagcatcaaatgaaactgcaatttattca tatcaggattatcaataccatatttttgaaaaagccgtttctgtaatgaaggagaaaactcaccgaggcagttccataggatggcaagatcctggtatcggtctgcgattcc gactcgtccaacatcaatacaacctattaatttcccctcgtcaaaaataaggttatcaagtgagaaatcaccatgagtgacgactgaatccggtgagaatggcaaaagctt atgcatttctttccagacttgttcaacaggccagccattacgctcgtcatcaaaatcactcgcatcaaccaaaccgttattcattcgtgattgcgcctgagcgagacgaaata cgcgatcgctgttaaaaggacaattacaaacaggaatcgaatgcaaccggcgcaggaacactgccagcgcatcaacaatattttcacctgaatcaggatattcttctaat acctggaatgctgttttcccggggatcgcagtggtgagtaaccatgcatcatcaggagtacggataaaatgcttgatggtcggaagaggcataaattccgtcagccagtt tagtctgaccatctcatctgtaacatcattggcaacgctacctttgccatgtttcagaaacaactctggcgcatcgggcttcccatacaatcgatagattgtcgcacctgatt gcccgacattatcgcgagcccatttatacccatataaatcagcatccatgttggaatttaatcgcggcctggagcaagacgtttcccgttgaatatggctcataacacccct tgtattactgtttatgtaagcagacagttttattgttcatgatgatatatttttatcttgtgcaatgtaacatcagagattttgagacacaacgtggctttgttgaataaatcgaactt ttgctgagttgaaggatcagtagctagaggatcgatcctttttaacccatcacatatacctgccgttcactattatttagtgaaatgagatattatgatattttctgaattgtgatt aaaaaggcaactttatgcccatgcaacagaaactataaaaaatacagagaatgaaaagaaacagatagattttttagttctttaggcccgtagtctgcaaatccttttatgat tttctatcaaacaaaagaggaaaatagaccagttgcaatccaaacgagagtctaatagaatgaggtcgaaagaagtggcggtctgagaacgaccgtaattgataatgcg ggctcagtgatccagcccccaaatgacagtgtctttaaccccagtaacttaccagcagggattaacccagcgaatcccttgttggccttgccgaatcaattgaccaataat ttccttggtgctttatctgcatcaattatcaacaacaaaactaagttgctgactgaccccaccttaatcattcaggaaggagaaagaagcgaagtcaaactgactagtcagg ttgttcaaaagattgaatcagagactacaaccaacggcagtggcccgccaaccgttagccgaaccattgaccttgcagatgttggtctacagttaaccatcaacattgaa cgaattgatgacaatggcttcattactctGAATTCCTTAGTTATTCCTATTCTGCACGAACTCAAAATACTCTTCATTTTTG ATAACCAAATTGAGTTTTTTGCCCTCTTGATTATTTTTGATCCTACCTAGTAGCCTCTTCTGCTCCTGCAG AATTGTGAGCGCTCACAATTGATATTTTGGTCTGTCGTTGCGATCGCCCGTTGCAGGCCGACATGAAGG ATTGACAATTAATCATCCGGCTCGTATAATGAATTGTGAGCGCTCACAATTGGTACCGGTGATACCAGCA TCGTCTTGATGCCCTTGGCAGCACCCTGCTAAGGAGGCAACAAGatgcaagcggtctttgataccaacatcctcatctaccacc tcaaaggctgtcttcctgaagcggggagtcaaattctgcgcagcagtctggggcgcggtgccgtctgttcagtcattacccgcctagaggttttgggttacgaccaacct tggccagaaagactgaaagcacaagctttgctccagctatttcgagaacgggcattggatgaatcgattgctgattgcacgattcagctgcgtcaacaacaacggatca |

| | |
|---|---|
| | aactgcccgatgccatcgttgccgcaactgcgctgacagagaacttgccactcgtgacgcgcaacaccaaagactttaaagccatcgctggcttacaactgattaaccc<br>ctttcaaccgaactagGGCTCACCTTCGGGTGGGCCTTTCTGCGCGCACACCGTGGAAACGGATGAAGGCACGA<br>ACCCAGTGGACATAAGCCTGTTCGGTTCGTAAGCTGTAATGCAAGTAGCGTATGCGCTCACGCAACTG<br>GTCCAGAACCTTGACCGAACGCAGCGGTGGTAACGGCGCAGTGGCGGTTTTCATGGCTTGTTATGACT<br>GTTTTTTTGGGGTACAGTCTATGCCTCGGGCATCCAAGCAGCAAGCGCGTTACGCCGTGGGTCGATGTT<br>TGATGTTATGGAGCAGCAACGATGTTACGCAGCAGGGCAGTCGCCCTAAAACAAAGTTAAACATCatgaa<br>cgagaaaaatataaaacacagtcaaaactttattacttcaaaacataatatagataaaataatgacaaatataagattaaatgaacatgataatatctttgaaatcggctcag<br>gaaaaggccattttacccttgaattagtaaagaggtgtaatttcgtaactgccattgaaatagaccataaattatgcaaaactacagaaaataaacttgttgatcacgataatt<br>tccaagttttaaacaaggatatattgcagtttaaatttcctaaaaaccaatcctataaaatatatggtaatataccttataacataagtacggatataatacgcaaaattgtttttg<br>atagtatagctaatgagatttatttaatcgtggaatacgggtttgctaaaagattattaaatacaaaacgctcattggcattacttttaatggcagaagttgatatttctatattaa<br>gtatggttccaagagaatattttcatcctaaacctaaagtgaatagctcacttatcagattaagtagaaaaaaatcaagaatatcacacaaagataaacaaaagtataattatt<br>tcgttatgaaatgggttaacaaagaatacaagaaaatatttacaaaaaatcaatttaacaattccttaaaacatgcaggaattgacgatttaaacaatattagctttgaacaatt<br>cttatctcttttcaatagctataaattatttaataagtaaTTGTTCAGAACGCTCGGTCTTGCACACCGGGCGTTTTTTCTTTGTGA<br>GTCCACCACCTCGACCTGAATGGAAGCCGGCGGCACCTCGCTAACGGATTCACCACTCCAAGAATTGG<br>AGCCAATCAATTCTTGCGGAGAACTGTGAATGCGCAAACCAACCCTTGGCAGAACATATCCATCGCGT<br>CCGCCATCTCCAGCAGCCGCACGCGGCGCATCTCGGGCAGCGTTGGGTCCTGGCCACGGGTGCGCATG<br>ATCGTGCTCCTGTCGTTGAGGACCCGGCTAGGCTGGCGGGGTTGCCTTACTGGTTAGCAGAATGAATC<br>ACCGATACGCGAGCGAACGTGAAGCGACTGCTGCTGCAAAACGTCTGCGACCTGAGCAACAACATGA<br>ATGGTCTTCGGTTTCCGTGTTTCGTAAAGTCTGGAAACGCGGAAGTCAGCGCCCTGCACCATTATGTTC<br>CGGATCTGCATCGCAGGATGCTGCTGGCTACCCTGTGGAACACCTACATCTGTATTAACGAAGCGCTGG<br>CATTGACCCTGAGTGATTTTTCTCTGGTCCCGCCGCATCCATACCGCCAGTTGTTTACCCTCACAACGTT<br>CCAGTAACCGGGCATGTTCATCATCAGTAACCCGTATCGTGAGCATCCTCTCTCGTTTCATCGGTATCAT<br>TACCCCCATGAACAGAAATCCCCCTTACACGGAGGCATCAGTGACCAAACAGGAAAAAACCGCCCTTA<br>ACATGGCCCGCTTTATCAGAAGCCAGACATTAACGCTTCTGGAGAAACTCAACGAGCTGGACGCGGAT<br>GAACAGGCAGACATCTGTGAATCGCTTCACGACCACGCTGATGAGCTTTACCGCAGCTGCCTCGCGCG<br>TTTCGGTGATGACGGTGAAAACCTCTGACACATGCAGCTCCCGGACGGTCACAGCTTGTCTGTAAGC<br>GGATGCCGGGAGCAGACAAGCCCGTCAGGGCGCGTCAGCGGGTGTTGGCGGGTGTCGGGGCGCAGC<br>CATGACCCAGTCACGTAGCGATAGCGGAGTGTATACTGGCTTAACTATGCGGCATCAGAGCAGATTGTA<br>CTGAGAGTGCACCATATGCGGTGTGAAATACCGCACAGATGCGTAAGGAGAAAATACCGCATCAGGCG<br>CTCTTCCGCTTCCTCGCTCACTGACTCGCTGCGCTCGGTCGTTCGGCTGCGGCGAGCGGTATCAGCTCA<br>CTCAAAGGCGGTAATACGGTTATCCACAGAATCAGGGGATAACGCAGGAAAGAACATGTGAGCAAAA<br>GGCCAGCAAAAGGCCAGGAACCGTAAAAAGGCCGCGTTGCTGGCGTTTTTCCATAGGCTCCGCCCCC<br>CTGACGAGCATCACAAAAATCGACGCTCAAGTCAGAGGTGGCGAAACCCGACAGGACTATAAAGATA<br>CCAGGCGTTTCCCCCTGGAAGCTCCCTCGTGCGCTCTCCTGTTCCGACCCTGCCGCTTACCGGATACCT<br>GTCCGCCTTTCTCCCTTCGGGAAGCGTGGCGCTTTCTCATAGCTCACGCTGTAGGTATCTCAGTTCGGT<br>GTAGGTCGTTCGCTCCAAGCTGGGCTGTGTGCACGAACCCCCCGTTCAGCCCGACCGCTGCGCCTTAT<br>CCGGTAACTATCGTCTTGAGTCCAACCCGGTAAGACACGACTTATCGCCACTGGCAGCAGCCACTGGT<br>AACAGGATTAGCAGAGCGAGGTATGTAGGCGGTGCTACAGAGTTCTTGAAGTGGTGGCCTAACTACGG<br>CTACACTAGAAGGACAGTATTTGGTATCTGCGCTCTGCTGAAGCCAGTTACCTTCGGAAAAAGAGTTG<br>GTAGCTCTTGATCCGGCAAACAAACCACCGCTGGTAGCGGTGGTTTTTTTGTTTGCAAGCAGCAGATT<br>ACGCGCAGAAAAAAAGGATCTCAAGAAGATCCTTTGATCTTTTCTACGGGGTCTGACGCTCAGTGGAA<br>CGAAAACTCACGTTAAGGGATTTTGGTCATGAGATTATCAAAAAGGATCTTCACCTAGATCCTTTTAAA<br>TTAAAAATGAAGTTTTAAATCAATCTAAAGTATATATGAGTAAACTTGGTCTGACAG |
| pBR-pil-KO | TTACCAATGCTTAATCAGTGAGGCACCTATCTCAGCGATCTGTCTATTTCGTTCATCCATAGTTGCCTGA<br>CTCCCCGTCGTGTAGATAACTACGATACGGGAGGGCTTACCATCTGGCCCCAGTGCTGCAATGATACCG<br>CGcGACCCACGCTCACCGGCTCCAGATTTATCAGCAATAAACCAGCCAGCCGGAAGGGCCGAGCGCAG<br>AAGTGGTCCTGCAACTTTATCCGCCTCCATCCAGTCTATTAATTGTTGCCGGGAAGCTAGAGTAAGTAG<br>TTCGCCAGTTAATAGTTTGCGCAACGTTGTTGCCATTGCTGCAGGCATCGTGGTGTCACGCTCGTCGTT<br>TGGTATGGCTTCATTCAGCTCCGGTTCCCAACGATCAAGGCGAGTTACATGATCCCCCATGTTGTGCAA |

AAAAGCGGTTAGCTCCTTCGGTCCTCCGATCGTTGTCAGAAGTAAGTTGGCCGCAGTGTTATCACTCAT
GGTTATGGCAGCACTGCATAATTCTCTTACTGTCATGCCATCCGTAAGATGCTTTTCTGTGACTGGTGAG
TACTCAACCAAGTCATTCTGAGAATAGTGTATGCGGCGACCGAGTTGCTCTTGCCCGGCGTCAACACG
GGATAATACCGCGCCACATAGCAGAACTTTAAAAGTGCTCATCATTGGAAAACGTTCTTCGGGGCGAA
AACTCTCAAGGATCTTACCGCTGTTGAGATCCAGTTCGATGTAACCCACTCGTGCACCCAACTGATCTT
CAGCATCTTTTACTTTCACCAGCGTTTCTGGGTGAGCAAAAACAGGAAGGCAAAATGCCGCAAAAAA
GGGAATAAGGGCGACACGGAAATGTTGAATACTCATACTCTTCCTTTTTCAATATTATTGAAGCATTTAT
CAGGGTTATTGTCTCATGAGCGGATACATATTTGAATGTATTTAGAAAAATAAACAAATAGGGGTTCCGC
Gcttagcgcttactctctcgggctttgctagcctgcagcggatcagccgtggtgaaatgctcgaggtgctgcgccaagactacatccgcaccgcccgtgccaaaggct
tgccggagcagcgcgtcatctacgtccacgctctacgcaatgcgatcaatcccctgattacgctcttgggctttgagttcgcgaccctgctcagcggcgcttttattgctg
aatatttctttaactggccggggctaggccgcttaattttgcaagccgtttttgcgcaggatctctacttggtaatggccagcttgatgatgggtgctgtgatgctgattctgg
gcaatctgctcgcagatctgctgctgcgctgggtcgatccccgcattcgcctggatgatctgaactaattacgccccgccctgccactcatcgcagtactgttgtaattcat
taagcattctgccgacatggaagccatcacagacggcatgatgaacctgaatcgccagcggcatcagcaccttgtcgccttgcgtataatatttgcccatggtgaaaac
gggggcgaagaagttgtccatattggccacgtttaaatcaaaactggtgaaactcacccagggattggctgagacgaaaaacatattctcaataaaccctttagggaaat
aggccaggttttcaccgtaacacgccacatcttgcgaatatatgtgtagaaactgccggaaatcgtcgtggtattcactccagagcgatgaaaacgtttcagtttgctcatg
gaaaacggtgtaacaagggtgaacactatcccatatcaccagctcaccgtctttcattgccatacggaattccggatgagcattcatcaggcgggcaagaatgtgaataa
aggccggataaaacttgtgcttatttttctttacggtctttaaaaaggccgtaatatccagctgaacggtctggttataggtacattgagcaactgactgaaatgcctcaaaat
gttctttacgatgccattgggatatatcaacggtggtatatccagtgatttttttctccattttagcttccttagctcctgaaaatctcgataactcaaaaaatacgcccggtagt
gatcttatttcattatggtgaaagttggaacctcttacgtgccgatcaGAAAAGTCTGAAAGTTCTTTACAAAACTCAATCTGCTTGT
TAGATTTTACTCACGAGGCTATTAAGTCTCGTAAATAGTTCAACTAAGGACTCATCGCAAAatgccaactatcca
gcagctaattcgtagcgaacgctcgaaggtacagaagaaaactaaatcccctgccctcaagcaatgtccccaacggcggggagtctgcactagggtttacaccacca
cccccaaaaagcccaactccgccctccggaaagtggcccgggtacgcctcacctccgggtttgaagtaactgcctatatccctggcattggccacaacctgcaagaac
actccgtagtactaatccggggcggtcgggtaaaagatttgcctggggttcgctaccatattgtgcggggcacgttggacgccaccggagttaaagaccgcaaacagg
gtcgctccaaatacggcaccaaacgggaaaaagcgaagaaataagaagtggcggtctgagaacgaccgtaattgataatgcgggctcagtgatccagcccccaaat
gacagtgtctttaaccccagtaacttaccagcagggattaacccagcgaatcccttgttggccttgccgaatcaattgaccaataatttccttggtgctttatctgcatcaatt
atcaacaacaaaactaagttgctgactgaccccaccttaatcattcaggaaggagaaagaagcgaagtcaaactgactagtcaggttgttcaaaagattgaatcagaga
ctacaaccaacggcagtggcccgccaaccgttagccgaaccattgaccttgcagatgttggtctacagttaaccatcaacattgaacgaattgatgacaatggcttcatta
ctctCCACCTCGACCTGAATGGAAGCCGGCGGCACCTCGCTAACGGATTCACCACTCCAAGAATTGGAG
CCAATCAATTCTTGCGGAGAACTGTGAATGCGCAAACCAACCCTTGGCAGAACATATCCATCGCGTCC
GCCATCTCCAGCAGCCGCACGCGGCGCATCTCGGGCAGCGTTGGGTCCTGGCCACGGGTGCGCATGAT
CGTGCTCCTGTCGTTGAGGACCCGGCTAGGCTGGCGGGGTTGCCTTACTGGTTAGCAGAATGAATCAC
CGATACGCGAGCGAACGTGAAGCGACTGCTGCTGCAAAACGTCTGCGACCTGAGCAACAACATGAAT
GGTCTTCGGTTTCCGTGTTTCGTAAAGTCTGGAAACGCGGAAGTCAGCGCCCTGCACCATTATGTTCCG
GATCTGCATCGCAGGATGCTGCTGGCTACCCTGTGGAACACCTACATCTGTATTAACGAAGCGCTGGCA
TTGACCCTGAGTGATTTTTCTCTGGTCCCGCCGCATCCATACCGCCAGTTGTTTACCCTCACAACGTTCC
AGTAACCGGGCATGTTCATCATCAGTAACCCGTATCGTGAGCATCCTCTCTCGTTTCATCGGTATCATTA
CCCCCATGAACAGAAATCCCCCTTACACGGAGGCATCAGTGACCAAACAGGAAAAAACCGCCCTTAA
CATGGCCCGCTTTATCAGAAGCCAGACATTAACGCTTCTGGAGAAACTCAACGAGCTGGACGCGGATG
AACAGGCAGACATCTGTGAATCGCTTCACGACCACGCTGATGAGCTTTACCGCAGCTGCCTCGCGCGT
TTCGGTGATGACGGTGAAAACCTCTGACACATGCAGCTCCCGGGACGGTCACAGCTTGTCTGTAAGCG
GATGCCGGGAGCAGACAAGCCCGTCAGGGCGCGTCAGCGGGTGTTGGCGGGTGTCGGGGCGCAGCC
ATGACCCAGTCACGTAGCGATAGCGGAGTGTATACTGGCTTAACTATGCGGCATCAGAGCAGATTGTAC
TGAGAGTGCACCATATGCGGTGTGAAATACCGCACAGATGCGTAAGGAGAAAATACCGCATCAGGCGC
TCTTCCGCTTCCTCGCTCACTGACTCGCTGCGCTCGGTCGTTCGGCTGCGGCGAGCGGTATCAGCTCAC
TCAAAGGCGGTAATACGGTTATCCACAGAATCAGGGGATAACGCAGGAAAGAACATGTGAGCAAAAG
GCCAGCAAAAGGCCAGGAACCGTAAAAAGGCCGCGTTGCTGGCGTTTTTCCATAGGCTCCGCCCCCCT
GACGAGCATCACAAAAATCGACGCTCAAGTCAGAGGTGGCGAAACCCGACAGGACTATAAAGATACC
AGGCGTTTCCCCCTGGAAGCTCCCTCGTGCGCTCTCCTGTTCCGACCCTGCCGCTTACCGGATACCTGT
CCGCCTTTCTCCCTTCGGGAAGCGTGGCGCTTTCTCATAGCTCACGCTGTAGGTATCTCAGTTCGGTGT

| | |
|---|---|
| | AGGTCGTTCGCTCCAAGCTGGGCTGTGTGCACGAACCCCCCGTTCAGCCCGACCGCTGCGCCTTATCC<br>GGTAACTATCGTCTTGAGTCCAACCCGGTAAGACACGACTTATCGCCACTGGCAGCAGCCACTGGTAA<br>CAGGATTAGCAGAGCGAGGTATGTAGGCGGTGCTACAGAGTTCTTGAAGTGGTGGCCTAACTACGGCT<br>ACACTAGAAGGACAGTATTTGGTATCTGCGCTCTGCTGAAGCCAGTTACCTTCGGAAAAAGAGTTGGT<br>AGCTCTTGATCCGGCAAACAAACCACCGCTGGTAGCGGTGGTTTTTTTGTTTGCAAGCAGCAGATTAC<br>GCGCAGAAAAAAAGGATCTCAAGAAGATCCTTTGATCTTTTCTACGGGGTCTGACGCTCAGTGGAACG<br>AAAACTCACGTTAAGGGATTTTGGTCATGAGATTATCAAAAAGGATCTTCACCTAGATCCTTTTAAATTA<br>AAAATGAAGTTTTAAATCAATCTAAAGTATATATGAGTAAACTTGGTCTGACAG |
| pBR3-pilNm-sepT2 | TTACCAATGCTTAATCAGTGAGGCACCTATCTCAGCGATCTGTCTATTTCGTTCATCCATAGTTGCCTGA<br>CTCCCCGTCGTGTAGATAACTACGATACGGGAGGGCTTACCATCTGGCCCCAGTGCTGCAATGATACCG<br>CGcGACCCACGCTCACCGGCTCCAGATTTATCAGCAATAAACCAGCCAGCCGGAAGGGCCGAGCGCAG<br>AAGTGGTCCTGCAACTTTATCCGCCTCCATCCAGTCTATTAATTGTTGCCGGGAAGCTAGAGTAAGTAG<br>TTCGCCAGTTAATAGTTTGCGCAACGTTGTTGCCATTGCTGCAGGCATCGTGGTGTCACGCTCGTCGTT<br>TGGTATGGCTTCATTCAGCTCCGGTTCCCAACGATCAAGGCGAGTTACATGATCCCCCATGTTGTGCAA<br>AAAAGCGGTTAGCTCCTTCGGTCCTCCGATCGTTGTCAGAAGTAAGTTGGCCGCAGTGTTATCACTCAT<br>GGTTATGGCAGCACTGCATAATTCTCTTACTGTCATGCCATCCGTAAGATGCTTTTCTGTGACTGGTGAG<br>TACTCAACCAAGTCATTCTGAGAATAGTGTATGCGGCGACCGAGTTGCTCTTGCCCGGCGTCAACACG<br>GGATAATACCGCGCCACATAGCAGAACTTTAAAAGTGCTCATCATTGGAAAACGTTCTTCGGGGCGAA<br>AACTCTCAAGGATCTTACCGCTGTTGAGATCCAGTTCGATGTAACCCACTCGTGCACCCAACTGATCTT<br>CAGCATCTTTTACTTTCACCAGCGTTTCTGGGTGAGCAAAAACAGGAAGGCAAAATGCCGCAAAAAA<br>GGGAATAAGGGCGACACGGAAATGTTGAATACTCATACTCTTCCTTTTTCAATATTATTGAAGCATTTAT<br>CAGGGTTATTGTCTCATGAGCGGATACATATTTGAATGTATTTAGAAAAATAAACAAATAGGGGTTCCGC<br>Gtcttagcgcttactctctcgggctttgctagcctgcagcggatcagccgtggtgaaatgctcgaggtgctgcgccaagactacatccgcaccgcccgtgccaaaggc<br>ttgccggagcagcgcgtcatctacgtccacgctctacgcaatgcgatcaatcccctgattacgctcttgggctttgagttcgcgaccctgctcagcggcgcttttattgctg<br>aatatttctttaactggccggggctaggccgcttaattttgcaagccgtttttgcgcaggatctctacttggtaatggccagcttgatgatgggtgctgtgatgctgattctgg<br>gcaatctgctcgcagatctgctgctgcgctgggtcgatccccgcattcgcctggatgatctgaactaaattgttcacgcttctggcttgagtcctgtagtgcgctcaggca<br>gtcttgggctgcgatcgtgcagacgcctagtttttttggcaaaccattggctaacattcaggtgctttgactctctgtgcaaataaaaaggcgcttctcgtggtcggcatttttt<br>ctaagtttcttggtggcgatcaatcgcgccttggggtggaaattaccccagagcggatcaatcttgcccacattggccgcaaggggaatcgcctgcagcttcaggatttt<br>gtttccgtggagctaccggatgggttgtttcaagacgggcgcgtgctagatagcttcagcctgtccgatctgctccgcaccgccatcaacgacaaccgcatcagaacca<br>aaagagtctccactgcggtgccaggacgagaggtgatcacccggctgctgcccatgcccgcggagctcgatgatgccgaattgcgcgagacactgatcaatcaaga<br>agctgctctgttcctgcccttcccgcgtgaagaggccgacatcgactatctgaaactggactattttctggacgcggatgaggtcgaaaagttgcacgtcctgctcgtgg<br>cgactcgtcggcaggtgaccgatgtctaccttgatctctttgcccaagctggcctgacgcctgacattgtggaagttgggagctttgcgctaatccgaacgattcgcgaa<br>gcgctgcgccagtacggattgcaagaagcggtcattctcctcaacgtggaatatgacctcacagaaatcgtgattatcgttgggggcgtgcctcggtttagccgcacga<br>tcgccatgggtactcagcagtggcagcgggctgcagcccaagctggtatcgacaatacaggacggggtattgatttcctgcaggggcgatcgttcacattgctcgaac<br>ccccggctgatctcctctctcaagctctggcacgatcggtcaccgagctctgtgatgaagtacggcgatcggctgacttctacctcagccaagagaccgatattgaagta<br>gccaagctctacgttgcaggtccaggctcagccttgccaacactacccgagtttctggggcaacgcctagggctccctgtagaactcgtcgatcctgtcgaggggttgg<br>ggctcgaggtggcggagaacctttatttgccaaatcgcgctgatgtgggcgtggtgttgggcttagccacgcgaggagtatagcggtgtacagcttagacattaactttc<br>tcaaagaacggacaaccacagcgccgccagttaccaccttcgggactgcagccgttgccactaatccctcggagcggctcccgctctggattggggtgggtatcggc<br>ttgcttttgcctgggctggcactcgtggttacggccttggctaacaatcgggtttcgaccctgactgctgacaaggccgaactcgaacaacaaattcagctggggcagcc<br>agcagaagcccgcctcaagagcattcaggctgagattgagcaaatcaataccgacaccacgtccttaattcaggtcttcccgcaggtcaaatcccagtctgctatcctca<br>ccgacctgagtcgacggaccccggtggcagttcagatcagcaagattgagcaggcgggcaagaaggtcaccctacagggcagttcttctaactacgactcggtcaat<br>gacctgctgctgaccttgcagcagtcgcgtttctttacagctagcagcctgcggattgaagaggcgaaattaggcgctacccctactcaaaatacaggggagaacagtc<br>ctaagctgtcgaatgtgaactatcgcattgtcggtgagctcagcgacgttcccacgacggatctcttgcaaaatctccgtcagctcgatgctcaaggtcttgctgctcgcct<br>gcagggattagtgaaccaaggggtactgaagccatgactgtctcgctgagtaaattcccggatgatgaacccgaggttctcgcatccaattacccaacggtttttgggct<br>gacaattacacccctcgtgggtggaattctagcggcgatcgcagggttggcaggcgctggctggatttttctcaatcttgttcagccaactcaagagcaggctaatcagc<br>tgacccaggaagttaatgacctccgcgctcgggctgaacagcaagccgctagcctgcgccaaattggcgctcagcagcaacagttagccaatgctcgagcccaaca<br>gcgtgtggtgcaaagcttctttgcatcgcccgccaccttagatacgctgttgctcgacatcaatggccaagtaaggcgggtggctggcttgcagctgcagtcctacttgc<br>cgaaggagtcggcgattgtgaccgatgggtcgctcggtgaaggcgtcaacaacaaactgaagcaaacccccgtgtttgtcgaggtgagtggcagtttcaaccaaact<br>gtgtccctactgcggggactcgaacagttgcaaccgctgttattggtgcgagacattgtcgtgacggcagagccggtgccggtcacggtcacgccgagggggcaaat |

| | |
|---|---|
| | cttggcgcggccagcccccaagctgacgacacagttcaatattgtggcgttgagtcccttgacgcccgaagaacagaagcaggttgaagaggctgctgctgccgctg<br>cgaaaaataatccaggaggaggtgcgaacccaggaggaggaacgggcaacgcaggtcagtcgggtcagtaacgaggcaagtggtaggaatcgcccttgaagcg<br>atcgcgagtgagtgaatttacggtgtgggtgtgaggtttcgaacaatgcaagcttggaaaattttggggagtggattggttcttggaactgccgcgatcgcggggcagc<br>catcaccagtctgggcccaagccgcatcgaccatcgcggcaaatccgtcggcaacggaagtcaatcgggttttggtgacgccgacaacgaatggcattagcctgtctt<br>tgagtacaacgggtactcaggcaccacaagccttttcaactaatctcgataaggtctgggtgacggatctgatcggcgctcgcctcaatctgccggagggtaagagcttt<br>actcagccgaacccggtacctgggatcgagtcgatcgcagtttcccaagtgagtgaaaacggcgttcggatcgttgtgaccggtagcgatcgcccgccgattgtgact<br>aacagcagtcggaatggtgggagcttgcagtttgaactcagtactcagactgcgatcgcagccgcacctagccccagtgctccactagtcggtgctgctccccgaccc<br>aacctgcgtccccctgcggccccggccccgcagcggactgcccaagcgacgcccccgacgccaattccagtcagcccagcaccacagccgccggtccaacccct<br>agaccctcggcaagtactgcaaccgacgccatcccctgcgccaggacaactccctcccctccagcctcgggccgttgcaccgcctttgggtgatattgcgatctccaa<br>cacagtgccccaaggcactgctgtcaccttcccgaatgcgcgtcccgtccctcgcattgccttgcgtaatgcttccgcgcgggaagtactcagcatcattgcaacgtctg<br>ctgggttaaaccttgtctatcttgatagcaacggccagccaactgctgcggccccagtggttagtaatgagccgaaagtgagtgttgatctgcaaaatattgatgctcagt<br>cagcctttaactacgttcttcagattgctgggttgcaggctagccgtcgcgataatgtcattctggtcgggagaaacctgccgccctctgctggtgggttagtttcccggac<br>tttccgcctcaaccaagcagtcgctcaaaacgtagcgggctatctcacgactttaggggctaactcagtcgtttcctttactaaaaaagtctgtgtctcgactggtgtgcag<br>gatacaactgcaacaggtggcggcgcgggcactggcgcaggacaaggtcagattcaaaccctcagtcagacttcggctcaatgttttgatgagccggaagtcaagga<br>gctcaaagttcctggcactccagccggcccattacccctcaaaaacatttcaatcaccgcagatattcgaacgaacagcctctcaatcgtgggcgatccccaagcaatc<br>aactttgcgacaggtttaattcaacagcttgaccttcgaaaaagacaagcttcagtcaatgttaaggttattgaccttgacttgaatgacctcaatgcgatcgcgactagtttc<br>tcctttggaattggcaacaccctattctcgggaagtggcggtctgagaacgaccgtaattgataatgcgggctcagtgatccagcccccaaatgacagtgtctttaaccc<br>cagtaacttaccagcagggattaacccagcgaatcccttgttggccttgccgaatcaattgaccaataatttccttggtgctttatctgcatcaattatcaacaacaaaactaa<br>gttgctgactgaccccaccttaatcattcaggaaggagaaagaagcgaagtcaaactgactagtcaggttgttcaaaagattgaatcagagactacaaccaacggcagt<br>ggcccgccaaccgttagccgaaccattgaccttgcagatgttggtctacagttaaccatcaacattgaacgaattgatgacaatggcttcattactctttacgccccgccct<br>gccactcatcgcagtactgttgtaattcattaagcattctgccgacatggaagccatcacagacggcatgatgaacctgaatcgccagcggcatcagcaccttgtcgcct<br>tgcgtataatatttgcccatggtgaaaacgggggcgaagaagttgtccatattggccacgtttaaatcaaaactggtgaaactcacccagggattggctgagacgaaaa<br>acatattctcaataaaccctttagggaaataggccaggttttcaccgtaacacgccacatcttgcgaatatatgtgtagaaactgccggaaatcgtcgtggtattcactcca<br>gagcgatgaaaacgtttcagtttgctcatggaaaacggtgtaacaagggtgaacactatcccatatcaccagctcaccgtctttcattgccatacggaattccggatgagc<br>attcatcaggcgggcaagaatgtgaataaaggccggataaaacttgtgcttatttttctttacggtctttaaaaaggccgtaatatccagctgaacggtctggttataggtac<br>attgagcaactgactgaaatgcctcaaaatgttctttacgatgccattgggatatatcaacggtggtatatccagtgatttttttctccattttagcttccttagctcctgaaaatc<br>tcgataactcaaaaaatacgcccggtagtgatcttatttcattatggtgaaagttggaacctcttacgtgccgatcaGAAAAGTCTGAAAGTTCTTTAC<br>AAAACTCAATCTGCTTGTTAGATTTACTCACGAGGCTATTAAGTCTCGTAAATAGTTCAACTAAGGACT<br>CATCGCAAAatgccaactatccagcagctaattcgtagcgaacgctcgaaggtacagaagaaaactaaatcccctgccctcaagcaatgtccccaacggcgg<br>ggagtctgcactagggtttacaccaccacccccaaaaagcccaactccgccctccggaaagtggcccgggtacgcctcacctccgggtttgaagtaactgcctatatc<br>cctggcattggccacaacctgcaagaacactccgtagtactaatccggggcggtcgggtaaaagatttgcctggggttcgctaccatattgtgcggggcacgttggac<br>gccaccggagttaaagaccgcaaacagggtcgctccaaatacggcaccaaacgggaaaaagcgaagaaataagaagtggcggtctgagaacgaccgtaattgata<br>atgcgggctcagtgatccagcccccaaatgacagtgtctttaaccccagtaacttaccagcagggattaacccagcgaatcccttgttggccttgccgaatcaattgacc<br>aataatttccttggtgctttatctgcatcaattatcaacaacaaaactaagttgctgactgaccccaccttaatcattcaggaaggagaaagaagcgaagtcaaactgacta<br>gtcaggttgttcaaaagattgaatcagagactacaaccaacggcagtggcccgccaaccgttagccgaaccattgaccttgcagatgttggtctacagttaaccatcaac<br>attgaacgaattgatgacaatggcttcattactctGAATTCCTTAGTTATTCCTATTCTGCACGAACTCAAAATACTCTTCATTT<br>TTGATAACCAAATTGAGTTTTTTGCCCTCTTGATTATTTTTGATCCTACCTAGTAGCCTCTTCTGCTCCTG<br>CAGAATTGTGAGCGCTCACAATTGATATTTTGGTCTGTCGTTGCGATCGCCCGTTGCAGGCCGACATGA<br>AGGATTGACAATTAATCATCCGGCTCGTATAATGAATTGTGAGCGCTCACAATTGGTACCGGTGATACCA<br>GCATCGTCTTGATGCCCTTGGCAGCACCCTGCTAAGGAGGCAACAAGatgcaagcggtctttgataccaacatcctcatcta<br>ccacctcaaaggctgtcttcctgaagcggggagtcaaattctgcgcagcagtctggggcgcggtgccgtctgttcagtcattacccgcctagaggttttgggttacgacc<br>aaccttggccagaaagactgaaagcacaagctttgctccagctatttcgagaacgggcattggatgaatcgattgctgattgcacgattcagctgcgtcaacaacaacg<br>gatcaaactgcccgatgccatcgttgccgcaactgcgctgacagagaacttgccactcgtgacgcgcaacaccaaagactttaaagccatcgctggcttacaactgatt<br>aacccctttcaaccgaactagGGCTCACCTTCGGGTGGGCCTTTCTGCGCGCACACCGTGGAAACGGATGAAGGC<br>ACGAACCCAGTGGACATAAGCCTGTTCGGTTCGTAAGCTGTAATGCAAGTAGCGTATGCGCTCACGCA<br>ACTGGTCCAGAACCTTGACCGAACGCAGCGGTGGTAACGGCGCAGTGGCGGTTTTCATGGCTTGTTAT<br>GACTGTTTTTTTGGGGTACAGTCTATGCCTCGGGCATCCAAGCAGCAAGCGCGTTACGCCGTGGGTCG<br>ATGTTTGATGTTATGGAGCAGCAACGATGTTACGCAGCAGGGCAGTCGCCCTAAAACAAAGTTAAACA<br>TCatgaacgagaaaaatataaaacacagtcaaaactttattacttcaaaacataatatagataaaataatgacaaatataagattaaatgaacatgataatatctttgaaatc |

| | |
|---|---|
| | ggctcaggaaaaggccattttacccttgaattagtaaagaggtgtaatttcgtaactgccattgaaatagaccataaattatgcaaaactacagaaaataaacttgttgatca<br>cgataatttccaagttttaaacaaggatatattgcagtttaaatttcctaaaaaccaatcctataaaatatatggtaatataccttataacataagtacggatataatacgcaaaa<br>ttgtttttgatagtatagctaatgagatttatttaatcgtggaatacgggtttgctaaaagattattaaatacaaaacgctcattggcattacttttaatggcagaagttgatatttct<br>atattaagtatggttccaagagaatattttcatcctaaacctaaagtgaatagctcacttatcagattaagtagaaaaaaatcaagaatatcacacaaagataaacaaaagta<br>taattatttcgttatgaaatgggttaacaaagaatacaagaaaatatttacaaaaaatcaatttaacaattccttaaaacatgcaggaattgacgatttaaacaatattagctttg<br>aacaattcttatctcttttcaatagctataaattatttaataagtaaTTGTTCAGAACGCTCGGTCTTGCACACCGGGCGTTTTTTCTTT<br>GTGAGTCCACCACCTCGACCTGAATGGAAGCCGGCGGCACCTCGCTAACGGATTCACCACTCCAAGA<br>ATTGGAGCCAATCAATTCTTGCGGAGAACTGTGAATGCGCAAACCAACCCTTGGCAGAACATATCCAT<br>CGCGTCCGCCATCTCCAGCAGCCGCACGCGGCGCATCTCGGGCAGCGTTGGGTCCTGGCCACGGGTGC<br>GCATGATCGTGCTCCTGTCGTTGAGGACCCGGCTAGGCTGGCGGGGTTGCCTTACTGGTTAGCAGAAT<br>GAATCACCGATACGCGAGCGAACGTGAAGCGACTGCTGCTGCAAAACGTCTGCGACCTGAGCAACAA<br>CATGAATGGTCTTCGGTTTCCGTGTTTCGTAAAGTCTGGAAACGCGGAAGTCAGCGCCCTGCACCATTA<br>TGTTCCGGATCTGCATCGCAGGATGCTGCTGGCTACCCTGTGGAACACCTACATCTGTATTAACGAAGC<br>GCTGGCATTGACCCTGAGTGATTTTTCTCTGGTCCCGCCGCATCCATACCGCCAGTTGTTTACCCTCACA<br>ACGTTCCAGTAACCGGGCATGTTCATCATCAGTAACCCGTATCGTGAGCATCCTCTCTCGTTTCATCGGT<br>ATCATTACCCCCATGAACAGAAATCCCCCTTACACGGAGGCATCAGTGACCAAACAGGAAAAAACCGC<br>CCTTAACATGGCCCGCTTTATCAGAAGCCAGACATTAACGCTTCTGGAGAAACTCAACGAGCTGGACG<br>CGGATGAACAGGCAGACATCTGTGAATCGCTTCACGACCACGCTGATGAGCTTTACCGCAGCTGCCTC<br>GCGCGTTTCGGTGATGACGGTGAAAACCTCTGACACATGCAGCTCCCGGGACGGTCACAGCTTGTCTG<br>TAAGCGGATGCCGGGAGCAGACAAGCCCGTCAGGGCGCGTCAGCGGGTGTTGGCGGGTGTCGGGGC<br>GCAGCCATGACCCAGTCACGTAGCGATAGCGGAGTGTATACTGGCTTAACTATGCGGCATCAGAGCAG<br>ATTGTACTGAGAGTGCACCATATGCGGTGTGAAATACCGCACAGATGCGTAAGGAGAAAATACCGCAT<br>CAGGCGCTCTTCCGCTTCCTCGCTCACTGACTCGCTGCGCTCGGTCGTTCGGCTGCGGCGAGCGGTAT<br>CAGCTCACTCAAAGGCGGTAATACGGTTATCCACAGAATCAGGGGATAACGCAGGAAAGAACATGTGA<br>GCAAAAGGCCAGCAAAAGGCCAGGAACCGTAAAAAGGCCGCGTTGCTGGCGTTTTTCCATAGGCTCC<br>GCCCCCCTGACGAGCATCACAAAAATCGACGCTCAAGTCAGAGGTGGCGAAACCCGACAGGACTATA<br>AAGATACCAGGCGTTTCCCCCTGGAAGCTCCCTCGTGCGCTCTCCTGTTCCGACCCTGCCGCTTACCGG<br>ATACCTGTCCGCCTTTCTCCCTTCGGGAAGCGTGGCGCTTTCTCATAGCTCACGCTGTAGGTATCTCAGT<br>TCGGTGTAGGTCGTTCGCTCCAAGCTGGGCTGTGTGCACGAACCCCCCGTTCAGCCCGACCGCTGCGC<br>CTTATCCGGTAACTATCGTCTTGAGTCCAACCCGGTAAGACACGACTTATCGCCACTGGCAGCAGCCAC<br>TGGTAACAGGATTAGCAGAGCGAGGTATGTAGGCGGTGCTACAGAGTTCTTGAAGTGGTGGCCTAACT<br>ACGGCTACACTAGAAGGACAGTATTTGGTATCTGCGCTCTGCTGAAGCCAGTTACCTTCGGAAAAAGA<br>GTTGGTAGCTCTTGATCCGGCAAACAAACCACCGCTGGTAGCGGTGGTTTTTTTGTTTGCAAGCAGCA<br>GATTACGCGCAGAAAAAAAGGATCTCAAGAAGATCCTTTGATCTTTTCTACGGGGTCTGACGCTCAGT<br>GGAACGAAAACTCACGTTAAGGGATTTTGGTCATGAGATTATCAAAAAGGATCTTCACCTAGATCCTTT<br>TAAATTAAAAATGAAGTTTTAAATCAATCTAAAGTATATATGAGTAAACTTGGTCTGACAG |
| pLLACCP-pilN-km | ACTCTTCCTTTTTCAATATTATTGAAGCATTTATCAGGGTTATTGTCTCATGAGCGGATACATATTTGAAT<br>GTATTTAGAAAAATAAACAAATAGGGGTTCCGCGttacgccccgccctgccactcatcgcagtactgttgtaattcattaagcattctgccg<br>acatggaagccatcacagacggcatgatgaacctgaatcgccagcggcatcagcaccttgtcgccttgcgtataatatttgcccatggtgaaaacgggggcgaagaa<br>gttgtccatattggccacgtttaaatcaaaactggtgaaactcacccagggattggctgagacgaaaaacatattctcaataaaccctttagggaaataggccaggttttca<br>ccgtaacacgccacatcttgcgaatatatgtgtagaaactgccggaaatcgtcgtggtattcactccagagcgatgaaaacgtttcagtttgctcatggaaaacggtgtaa<br>caagggtgaacactatcccatatcaccagctcaccgtctttcattgccatacggaattccggatgagcattcatcaggcgggcaagaatgtgaataaaggccggataaa<br>acttgtgcttatttttctttacggtctttaaaaaggccgtaatatccagctgaacggtctggttataggtacattgagcaactgactgaaatgcctcaaaatgttctttacgatgc<br>cattgggatatatcaacggtggtatatccagtgatttttttctccattttagcttccttagctcctgaaaatctcgataactcaaaaaatacgcccggtagtgatcttatttcattat<br>ggtgaaagttggaacctcttacgtgccgatcatttacactttatgcttccggctcgtatgttGTGTGGTGCTAAGGAGGCAACAAGatgtcaatttatc<br>aagaatttgttaataaatatagtttaagtaaaactctaagatttgagttaatcccacagggtaaaacacttgaaaacataaaagcaagaggtttgattttagatgatgagaaaa<br>gagctaaagactacaaaaaggctaaacaaataattgataaatatcatcagtttttatagaggagatattaagttcggtttgtattagcgaagatttattacaaaactattctgat<br>gtttattttaaacttaaaaagagtgatgatgataatctacaaaaagattttaaaagtgcaaaagatacgataaagaaacaaatatctgaatatataaaggactcagagaaattt<br>aagaatttgtttaatcaaaaccttatcgatgctaaaaaagggcaagagtcagatttaattctatggctaaagcaatctaaggataatggtatagaactatttaaagccaatagt |

gatatcacagatatagatgaggcgttagaaataatcaaatcttttaaaggttggacaacttattttaagggttttcatgaaaatagaaaaaatgtttatagtagcaatgatattc
ctacatctattatttataggatagtagatgataatttgcctaaatttctagaaaataaagctaagtatgagagtttaaaagacaaagctccagaagctataaactatgaacaaa
ttaaaaaagatttggcagaagagctaacctttgatattgactacaaaacatctgaagttaatcaaagagttttttcacttgatgaagtttttgagatagcaaactttaataattat
ctaaatcaaagtggtattactaaatttaatactattattggtggtaaatttgtaaatggtgaaaatacaaagagaaaaggtataaatgaatatataaatctatactcacagcaaa
taaatgataaaacactcaaaaaatataaaatgagtgttttatttaagcaaattttaagtgatacagaatctaaatcttttgtaattgataagttagaagatgatagtgatgtagtta
caacgatgcaaagtttttatgagcaaatagcagcttttaaaacagtagaagaaaaatctattaaagaaacactatctttattatttgatgatttaaaagctcaaaaacttgatttg
agtaaaatttattttaaaaatgataaatctcttactgatctatcacaacaagtttttgatgattatagtgttattggtacagcggtactagaatatataactcaacaaatagcacct
aaaaatcttgataaccctagtaagaaagagcaagaattaatagccaaaaaaactgaaaaagcaaaatacttatctctagaaactataaagcttgccttagaagaatttaata
agcatagagatatagataaacagtgtaggtttgaagaaatacttgcaaactttgcggctattccgatgatatttgatgaaatagctcaaaacaaagacaatttggcacagat
atctatcaaatatcaaaatcaaggtaaaaaagacctacttcaagctagtgcggaagatgatgttaaagctatcaaggatcttttagatcaaactaataatctcttacataaact
aaaaatatttcatattagtcagtcagaagataaggcaaatattttagacaaggatgagcatttttatctagtatttgaggagtgctactttgagctagcgaatatagtgcctcttt
ataacaaaattagaaactatataactcaaaagccatatagtgatgagaaatttaagctcaattttgagaactcgactttggctaatggttgggataaaaataaagagcctga
caatacggcaattttatttatcaaagatgataaatattatctgggtgtgatgaataagaaaaataacaaaatatttgatgataaagctatcaaagaaaataaaggcgagggtt
ataaaaaaattgtttataaacttttacctggcgcaaataaaatgttacctaaggttttcttttctgctaaatctataaaattttataatcctagtgaagatatacttagaataagaaat
cattccacacatacaaaaaatggtagtcctcaaaaaggatatgaaaaatttgagtttaatattgaagattgccgaaaatttatagatttttataaacagtctataagtaagcatc
cggagtggaaagattttggatttagattttctgatactcaaagatataattctatagatgaattttatagagaagttgaaaatcaaggctacaaactaacttttgaaaatatatca
gagagctatattgatagcgtagttaatcagggtaaattgtacctattccaaatctataataaagatttttcagcttatagcaaagggcgaccaaatctacatactttatattgga
aagcgctgtttgatgagagaaatcttcaagatgtggtttataagctaaatggtgaggcagagctttttttatcgtaaacaatcaatacctaaaaaaatcactcacccagctaaa
gaggcaatagctaataaaaacaaagataatcctaaaaaagagagtgtttttgaatatgatttaatcaaagataaacgctttactgaagataagtttttctttcactgtcctatta
caatcaattttaaatctagtggagctaataagtttaatgatgaaatcaatttattgctaaaagaaaaagcaaatgatgttcatatattaagtatagatagaggtgaaagacattt
agcttactatactttggtagatggtaaaggcaatatcatcaaacaagatactttcaacatcattggtaatgatagaatgaaaacaaactaccatgataagcttgctgcaatag
agaaagatagggattcagctaggaaagactggaaaaagataaataacatcaaagagatgaaagagggctatctatctcaggtagttcatgaaatagctaagctagttat
agagtataatgctattgtggtttttgaggatttaaattttggatttaaaagagggcgtttcaaggtagagaagcaggtctatcaaaagttagaaaaaatgctaattgagaaact
aaactatctagttttcaaagataatgagtttgataaaactgggggagtgcttagagcttatcagctaacagcaccttttgagacttttaaaaagatgggtaaacaaacaggta
ttatctactatgtaccagctggttttacttcaaaaatttgtcctgtaactggttttgtaaatcagttatatcctaagtatgaaagtgtcagcaaatctcaagagttctttagtaagttt
gacaagatttgttataaccttgataagggctattttgagtttagttttgattataaaaactttggtgacaaggctgccaaaggcaagtggactatagctagctttgggagtaga
ttgattaactttagaaattcagataaaaatcataattgggatactcgagaagtttatccaactaaagagttggagaaattgctaaaagattattctatcgaatatgggcatggc
gaatgtatcaaagcagctatttgcggtgagagcgacaaaaagtttttttgctaagctaactagtgtcctaaatactatcttacaaatgcgtaactcaaaaacaggtactgagtt
agattatctaatttcaccagtagcagatgtaaatggcaatttctttgattcgcgacaggcgccaaaaaaatatgcctcaagatgctgatgccaatggtgcttatcatattgggc
taaaaggtctgatgctactaggtaggatcaaaaataatcaagagggcaaaaaactcaatttggttatcaaaaatgaagagtattttgagttcgtgcagaataggaataacta
attgacagctagctcagtcctaggtataatgctagcgctgatttaggcaaaaacgggtctaagaactttaaataatttctactgttgtagatGTTTCCGTGGAGC
TACCGGATGgtctaagaactttaaataatttctactgttgtagattagcgatttatgaaggtcatttttttttgccggtcttagcgcttactctctcgggctttgctagcctg
cagcggatcagccgtggtgaaatgctcgaggtgctgcgccaagactacatccgcaccgcccgtgccaaaggcttgccggagcagcgcgtcatctacgtccacgctct
acgcaatgcgatcaatcccctgattacgctcttgggctttgagttcgcgaccctgctcagcggcgcttttattgctgaatatttctttaactggccggggctaggccgcttaa
ttttgcaagccgtttttgcgcaggatctctacttggtaatggccagcttgatgatgggtgctgtgatgctgattctgggcaatctgctcgcagatctgctgctgcgctgggtc
gatccccgcattcgcctggatgatctgaactaaattgttcacgcttctggcttgagtcctgtagtgcgctcaggcagtcttgggctgcgatcgtgcagacgcctagtttttttg
gcaaaccattggctaacattcaggtgctttgactctctgtgcaaataaaaaggcgcttctcgtggtcggcatttttttctaagtttcttggtggcgatcaatcgcgccttggggt
ggaaattaccccagagcggatcaatcttgcccacattggccgcaaggggaatcgcctgcagcttcaggattttgtttagaaaaactcatcgagcatcaaatgaaactgca
atttattcatatcaggattatcaataccatatttttgaaaaagccgtttctgtaatgaaggagaaaactcaccgaggcagttccataggatggcaagatcctggtatcggtct
gcgattccgactcgtccaacatcaatacaacctattaatttcccctcgtcaaaaataaggttatcaagtgagaaatcaccatgagtgacgactgaatccggtgagaatggc
aaaagcttatgcatttctttccagacttgttcaacaggccagccattacgctcgtcatcaaaatcactcgcatcaaccaaaccgttattcattcgtgattgcgcctgagcgag
acgaaatacgcgatcgctgttaaaaggacaattacaaacaggaatcgaatgcaaccggcgcaggaacactgccagcgcatcaacaatattttcacctgaatcaggata
ttcttctaatacctggaatgctgttttcccggggatcgcagtggtgagtaaccatgcatcatcaggagtacggataaaatgcttgatggtcggaagaggcataaattccgtc
agccagtttagtctgaccatctcatctgtaacatcattggcaacgctacctttgccatgtttcagaaacaactctggcgcatcgggcttcccatacaatcgatagattgtcgc
acctgattgcccgacattatcgcgagcccatttatacccatataaatcagcatccatgttggaatttaatcgcggcctggagcaagacgtttcccgttgaatatggctcataa
cacccccttgtattactgtttatgtaagcagacagttttattgttcatgatgatatatttttatcttgtgcaatgtaacatcagagattttgagacacaacgtggctttgttgaataaat
cgaacttttgctgagttgaaggatcagtagctagaggatcgatcctttttaacccatcacatatacctgccgttcactattatttagtgaaatgagatattatgatattttctgaat
tgtgattaaaaaggcaactttatgcccatgcaacagaaactataaaaaatacagagaatgaaaagaaacagatagatttttttagttctttaggcccgtagtctgcaaatcctt
ttatgattttctatcaaacaaaagaggaaaatagaccagttgcaatccaaacgagagtctaatagaatgaggtcgaaatgggcgatccccaagcaatcaactttgcgaca

ggtttaattcaacagcttgaccttcgaaaaagacaagcttcagtcaatgttaaggttattgaccttgacttgaatgacctcaatgcgatcgcgactagtttctcctttggaattg
gcaacaccctattctcgggaagtggcggtctgagaacgaccgtaattgataatgcgggctcagtgatccagcccccaaatgacagtgtctttaaccccagtaacttacca
gcagggattaacccagcgaatcccttgttggccttgccgaatcaattgaccaataatttccttggtgctttatctgcatcaattatcaacaacaaaactaagttgctgactgac
cccaccttaatcattcaggaaggagaaagaagcgaagtcaaactgactagtcaggttgttcaaaagattgaatcagagactacaaccaacggcagtggcccgccaac
cgttagccgaaccattgaccttgcagatgttggtctacagttaaccatcaacattgaacgaattgatgacaatggcttcattactcttagtagccatgatcgagcgatcggc
ctcgggggcgttgcagttcatccgcatgatctcgcccctgtcttggatgcccattgccgttatggcctttggcattggcgatctgcccgtttattttctgctagcgatcgcag
cggtctggccaattctactcagcaccagcagcggtactgcggctgttaaccacaagctcttacttctagctcgtagcctttgtgccactcgcagtgaaacgattcggcgca
tcgtgattcctgcgatcgtgccgcaaatccttgtgggtgttcgactcgcgatcggcacgatctggattgtgctcgtccccgctgagatgctgggcgttagctcaggcctcg
gttacttcattcttgatacccgcgatcgcattgcctataacgaacttaccgctgttttattagcgatcggcatcattggctgcgccctagattggagcctgcaattcctgcaaa
agtactggcaacccagtcgctaaacctcttggcatccattctccaaacctctcaaaggcgtctacagggttgagcttgtggggaggggcggctatcctcgcccgccaaa
agcccgatcgtcatcactggcaagcccagcaacgacagatggaactgaactcagtccaagggcagctcgatgccctggtcgtcaaagcaccctacgcaggcagggt
taggcgggtacgctggctagagcaggtgggcggacaggtcaaggtggagattgctcttcaagtctcaaacactccctaagttttgaccttgattacagtggccccatcta
gtttttgtatatacaatgactaggtgcgtatcacctacgatccagtcaaacgcgacaagacactgcttgagcgaggactcgacttcgagagtgcgatcgaagtctttgcag
ggctcacgctagaagtcgaagacacccgacgcgactatggcgaaaagcgagtgctctgcgtcggcttccttgatggccgcatggtgatggttggctataccccctcgcg
gtcgaacacgtcacatcttttcgatgaggaaatgcaatgcccgagaaattcgccgctacacgccatacttcgagcaagcctgatgacgacttgcccgagttgaccgatg
aaatgctcagtcgggcggtgctcaagcagggcggtaagcgtatcggcagaccccgatcgcaatctccaaaagtagcgatcagcctgcgcttagaagctgaagtcctg
cagcgctggcgtgacagtggccctggctggcaaactcgtatggctgaattgctctcgaaagaaacccccctaagtctttgatcgctcaagcccaacggcaggacaagct
acaccagtggcaaacccagcagcgacagatggaacttaactcagtccaagcgcagctcgatgcactggtcgtcaaagcaccttactccggcagagtcaggcgggtg
cgttggctagagcaagtcggtggacaggtgaaagtagagattgctcttcagatctttaacgagtgataaattcatagctttgtcattatctattatcgagagcaaataagagc
tatcaaggagcacctaagtgatctgctacttggtaaccttttttcagttagtctataaacttctcctttgccgccaagtgactccattagctcatatttaataagctctgaaatagc
acttttccaagaggcctcactacgagcattacccatatctgcaagctgcttattattagtttcaatcatcaatccacccataactggcagacaaattaattctccattagaatct
gatgctacctctattaacagatcttttccaagatcgcttaagcgtggtaattcaggagcaaaagatacttctaaagtattattagtgctttctaagctactgtaattttgtataattt
tctgagatatctgccgactaaacttatgctgaaactcctccaatgactcatatgtttcaattaagccttttttgttgacacaaactacgaaaatcacatagtgccttgtattgatcc
tgatcaacgctttctagatatactggagctctggaaaagtaaatcatcgcttgcttgccagacttaagatgttcttctatttcctcaacagttccactaataaatttaccagtagg
tgttcctaatcttgtccaaaaaacagcaacaagtagatcgcagtctttcaaaatttgcttgtttattatggattgtgctcgtccgctcatatcaggatgggcatgtgtttcccacc
ctactggatcaagtactattttacgatcaatagaattaatggtattccactcgtatattatgttccgaataatgttgcgttcttttgcaacatcgctcggtgatgcaatcataatcc
taagaactgtagcgttgtatggcattgctagccccttagagaaaatttcagtagcatttactatttattagctatatttacataaatttttttaaactcatcagagataatgactcata
ttttctatacgatcgcaagtaacttcactgccccccttcaccacctccgtatcgcccgagaaagcaaagtgctccgcctgactcaccagcgactctggcaacggcagcgta
atccgatagcactccagaaactcacccagcactcccatcgcatgggctttgaccgaatccgagaaccgatcgccctctCCACCTCGACCTGAATGGA
AGCCGGCGGCACCTCGCTAACGGATTCACCACTCCAAGAATTGGAGCCAATCAATTCTTGCGGAGAAC
TGTGAATGCGCAAACCAACCCTTGGCAGAACATATCCATCGCGTCCGCCATCTCCAGCAGCCGCACGC
GGCGCATCTCGGGCAGCGTTGGGTCCTGGCCACGGGTGCGCATGATCGTGCTCCTGTCGTTGAGGACC
CGGCTAGGCTGGCGGGGTTGCCTTACTGGTTAGCAGAATGAATCACCGATACGCGAGCGAACGTGAAG
CGACTGCTGCTGCAAAACGTCTGCGACCTGAGCAACAACATGAATGGTCTTCGGTTTCCGTGTTTCGT
AAAGTCTGGAAACGCGGAAGTCAGCGCCCTGCACCATTATGTTCCGGATCTGCATCGCAGGATGCTGC
TGGCTACCCTGTGGAACACCTACATCTGTATTAACGAAGCGCTGGCATTGACCCTGAGTGATTTTTCTC
TGGTCCCGCCGCATCCATACCGCCAGTTGTTTACCCTCACAACGTTCCAGTAACCGGGCATGTTCATCA
TCAGTAACCCGTATCGTGAGCATCCTCTCTCGTTTCATCGGTATCATTACCCCCATGAACAGAAATCCCC
CTTACACGGAGGCATCAGTGACCAAACAGGAAAAAACCGCCCTTAACATGGCCCGCTTTATCAGAAGC
CAGACATTAACGCTTCTGGAGAAACTCAACGAGCTGGACGCGGATGAACAGGCAGACATCTGTGAAT
CGCTTCACGACCACGCTGATGAGCTTTACCGCAGCTGCCTCGCGCGTTTCGGTGATGACGGTGAAAAC
CTCTGACACATGCAGCTCCCGGACGGTCACAGCTTGTCTGTAAGCGGATGCCGGGAGCAGACAAGC
CCGTCAGGGCGCGTCAGCGGGTGTTGGCGGGTGTCGGGGCGCAGCCATGACCCAGTCACGTAGCGAT
AGCGGAGTGTATACTGGCTTAACTATGCGGCATCAGAGCAGATTGTACTGAGAGTGCACCATATGCGGT
GTGAAATACCGCACAGATGCGTAAGGAGAAAATACCGCATCAGGCGCTCTTCCGCTTCCTCGCTCACT
GACTCGCTGCGCTCGGTCGTTCGGCTGCGGCGAGCGGTATCAGCTCACTCAAAGGCGGTAATACGGTT
ATCCACAGAATCAGGGGATAACGCAGGAAAGAACATGTGAGCAAAAGGCCAGCAAAAGGCCAGGAA
CCGTAAAAAGGCCGCGTTGCTGGCGTTTTTCCATAGGCTCCGCCCCCCTGACGAGCATCACAAAAATC
GACGCTCAAGTCAGAGGTGGCGAAACCCGACAGGACTATAAAGATACCAGGCGTTTCCCCCTGGAAG

| | |
|---|---|
| | CTCCCTCGTGCGCTCTCCTGTTCCGACCCTGCCGCTTACCGGATACCTGTCCGCCTTTCTCCCTTCGGG<br>AAGCGTGGCGCTTTCTCATAGCTCACGCTGTAGGTATCTCAGTTCGGTGTAGGTCGTTCGCTCCAAGCT<br>GGGCTGTGTGCACGAACCCCCCGTTCAGCCCGACCGCTGCGCCTTATCCGGTAACTATCGTCTTGAGTC<br>CAACCCGGTAAGACACGACTTATCGCCACTGGCAGCAGCCACTGGTAACAGGATTAGCAGAGCGAGG<br>TATGTAGGCGGTGCTACAGAGTTCTTGAAGTGGTGGCCTAACTACGGCTACACTAGAAGGACAGTATTT<br>GGTATCTGCGCTCTGCTGAAGCCAGTTACCTTCGGAAAAAGAGTTGGTAGCTCTTGATCCGGCAAACA<br>AACCACCGCTGGTAGCGGTGGTTTTTTTGTTTGCAAGCAGCAGATTACGCGCAGAAAAAAAGGATCTC<br>AAGAAGATCCTTTGATCTTTTCTACGGGGTCTGACGCTCAGTGGAACGAAAACTCACGTTAAGGGATT<br>TTGGTCATGAGATTATCAAAAAGGATCTTCACCTAGATCCTTTTAAATTAAAAATGAAGTTTTAAATCAA<br>TCTAAAGTATATATGAGTAAACTTGGTCTGACAGTTACCAATGCTTAATCAGTGAGGCACCTATCTCAGC<br>GATCTGTCTATTTCGTTCATCCATAGTTGCCTGACTCCCCGTCGTGTAGATAACTACGATACGGGAGGGC<br>TTACCATCTGGCCCCAGTGCTGCAATGATACCGCGcGACCCACGCTCACCGGCTCCAGATTTATCAGCA<br>ATAAACCAGCCAGCCGGAAGGGCCGAGCGCAGAAGTGGTCCTGCAACTTTATCCGCCTCCATCCAGTC<br>TATTAATTGTTGCCGGGAAGCTAGAGTAAGTAGTTCGCCAGTTAATAGTTTGCGCAACGTTGTTGCCATT<br>GCTGCAGGCATCGTGGTGTCACGCTCGTCGTTTGGTATGGCTTCATTCAGCTCCGGTTCCCAACGATCA<br>AGGCGAGTTACATGATCCCCCATGTTGTGCAAAAAAGCGGTTAGCTCCTTCGGTCCTCCGATCGTTGTC<br>AGAAGTAAGTTGGCCGCAGTGTTATCACTCATGGTTATGGCAGCACTGCATAATTCTCTTACTGTCATGC<br>CATCCGTAAGATGCTTTTCTGTGACTGGTGAGTACTCAACCAAGTCATTCTGAGAATAGTGTATGCGGC<br>GACCGAGTTGCTCTTGCCCGGCGTCAACACGGGATAATACCGCGCCACATAGCAGAACTTTAAAAGTG<br>CTCATCATTGGAAAACGTTCTTCGGGGCGAAAACTCTCAAGGATCTTACCGCTGTTGAGATCCAGTTCG<br>ATGTAACCCACTCGTGCACCCAACTGATCTTCAGCATCTTTTACTTTCACCAGCGTTTCTGGGTGAGCA<br>AAAACAGGAAGGCAAAATGCCGCAAAAAAGGGAATAAGGGCGACACGGAAATGTTGAATACTCAT |
| pCPF1-pil | ttacgccccgccctgccactcatcgcagtactgttgtaattcattaagcattctgccgacatggaagccatcacagacggcatgatgaacctgaatcgccagcggcatca<br>gcaccttgtcgccttgcgtataatatttgcccatggtgaaaacgggggcgaagaagttgtccatattggccacgtttaaatcaaaactggtgaaactcacccagggattgg<br>ctgagacgaaaaacatattctcaataaaccctttagggaaataggccaggttttcaccgtaacacgccacatcttgcgaatatatgtgtagaaactgccggaaatcgtcgt<br>ggtattcactccagagcgatgaaaacgtttcagtttgctcatggaaaacggtgtaacaagggtgaacactatcccatatcaccagctcaccgtctttcattgccatacgga<br>attccggatgagcattcatcaggcgggcaagaatgtgaataaaggccggataaaacttgtgcttatttttctttacggtctttaaaaaggccgtaatatccagctgaacggt<br>ctggttataggtacattgagcaactgactgaaatgcctcaaaatgttctttacgatgccattgggatatatcaacggtggtatatccagtgatttttttctccattttagcttcctt<br>agctcctgaaaatctcgataactcaaaaaatacgcccggtagtgatcttatttcattatggtgaaagttggaacctcttacgtgccgatcatcactgcccgctttccagtcgg<br>gaaacctgtcgtgccagctgcattaatgaatcggccaacgcgcggggagaggcggtttgcgtattgggcgccagggtggtttttcttttcaccagtgagacgggcaac<br>agctgattgcccttcaccgcctggccctgagagagttgcagcaagcggtccacgctggtttgccccagcaggcgaaaatcctgtttgatggtggttaacggcgggatat<br>aacatgagctgtcttcggtatcgtcgtatcccactaccgagatatccgcaccaacgcgcagcccggactcggtaatggcgcgcattgcgcccagcgccatctgatcgtt<br>ggcaaccagcatcgcagtgggaacgatgccctcattcagcatttgcatggtttgttgaaaaccggacatggcactaaagtcgccttcccgttccgctatcggctgaatttg<br>attgcgagtgagatatttatgccagccagccagacgcagacgcgccgagacagaacttaatgggcccgctaacagcgcgatttgctggtgacccaatgcgaccagat<br>gctccacgcccagtcgcgtaccgtcttcatgggagaaaataatactgttgatgggtgtctggtcagagacatcaagaaataacgccggaacattagtgcaggcagcttc<br>cacagcaatggcatcctggtcatccagcggatagttaatgatcagcccactgacgcgttgcgcgagaagattgtgcaccgccgctttacaggcttcgacgccgcttcgt<br>tctaccatcgacaccaccacgctggcacccagttgatcggcgcgagatttaatcgccgcgacaatttgcgacggcgcgtgcagggccagactggaggtggcaacgc<br>caatcagcaacgactgtttgcccgccagttgttgtgccacgcggttgggaatgtaattcagctccgccatcgccgcttccactttttcccgcgttttcgcagaaacgtggct<br>ggcctggttcaccacgcgggaaacggtctgataagagacaccggcatactctgcgacatcgtataacgttactggtttcacattcaccaccctgaattgactctcttccag<br>tatagatgctagcattatacctaggactgagctagctgtcaaaattgtgagcgctcacaattgatattttggtctgtcgttgcgatcgcccgttgcaggccgacatgaagga<br>tttacactttatgcttccggctcgtatgttGTGTGGaattgtgagcgctcacaattggtaccggtgataccAGCATCGTCTTGATGCCCTTGGCA<br>GCACCCTGCTAAGGAGGCAACAAGatgtcaatttatcaagaatttgttaataaatatagtttaagtaaaactctaagatttgagttaatcccacagggt<br>aaaacacttgaaaacataaaagcaagaggtttgattttagatgatgagaaaagagctaaagactacaaaaaggctaaacaaataattgataaatatcatcagtttttatag<br>aggagatattaagttcggtttgtattagcgaagatttattacaaaactattctgatgtttattttaaacttaaaaagagtgatgatgataatctacaaaaagattttaaaagtgcaa<br>aagatacgataaagaaacaaatatctgaatatataaaggactcagagaaatttaagaatttgtttaatcaaaaccttatcgatgctaaaaaagggcaagagtcagatttaatt<br>ctatggctaaagcaatctaaggataatggtatagaactatttaaagccaatagtgatatcacagatatagatgaggcgttagaaataatcaaatcttttaaaggttggacaac<br>ttattttaagggttttcatgaaaatagaaaaaatgtttatagtagcaatgatattcctacatctattatttataggatagtagatgataatttgcctaaatttctagaaaataaagct<br>aagtatgagagtttaaaagacaaagctccagaagctataaactatgaacaaattaaaaaagatttggcagaagagctaacctttgatattgactacaaaacatctgaagtt<br>aatcaaagagtttttcacttgatgaagtttttgagatagcaaactttaataattatctaaatcaaagtggtattactaaatttaatactattattggtggtaaatttgtaaatggtga |

aaatacaaagagaaaaggtataaatgaatatataaatctatactcacagcaaataaatgataaaacactcaaaaaatataaaatgagtgttttatttaagcaaattttaagtga
tacagaatctaaatcttttgtaattgataagttagaagatgatagtgatgtagttacaacgatgcaaagtttttatgagcaaatagcagcttttaaaacagtagaagaaaaatct
attaaagaaacactatctttattatttgatgatttaaaagctcaaaaacttgatttgagtaaaatttattttaaaaatgataaatctcttactgatctatcacaacaagtttttgatgat
tatagtgttattggtacagcggtactagaatatataactcaacaaatagcacctaaaaatcttgataaccctagtaagaaagagcaagaattaatagccaaaaaaactgaa
aaagcaaaatacttatctctagaaactataaagcttgccttagaagaatttaataagcatagagatatagataaacagtgtaggtttgaagaaatacttgcaaactttgcggc
tattccgatgatatttgatgaaatagctcaaaacaaagacaatttggcacagatatctatcaaatatcaaaatcaaggtaaaaaagacctacttcaagctagtgcggaagat
gatgttaaagctatcaaggatcttttagatcaaactaataatctcttacataaactaaaaatatttcatattagtcagtcagaagataaggcaaatattttagacaaggatgagc
atttttatctagtatttgaggagtgctactttgagctagcgaatatagtgcctctttataacaaaattagaaactatataactcaaaagccatatagtgatgagaaatttaagctc
aattttgagaactcgactttggctaatggttgggataaaaataaagagcctgacaatacggcaattttatttatcaaagatgataaatattatctgggtgtgatgaataagaaa
aataacaaaatatttgatgataaagctatcaaagaaaataaaggcgagggttataaaaaaattgtttataaacttttacctggcgcaaataaaatgttacctaaggttttctttt
ctgctaaatctataaaattttataatcctagtgaagatatacttagaataagaaatcattccacacatacaaaaaatggtagtcctcaaaaaggatatgaaaaatttgagtttaa
tattgaagattgccgaaaatttatagatttttataaacagtctataagtaagcatccggagtggaaagattttggatttagattttctgatactcaaagatataattctatagatga
attttatagagaagttgaaaatcaaggctacaaactaacttttgaaaatatatcagagagctatattgatagcgtagttaatcagggtaaattgtacctattccaaatctataat
aaagatttttcagcttatagcaaagggcgaccaaatctacatactttatattggaaagcgctgtttgatgagagaaatcttcaagatgtggtttataagctaaatggtgaggc
agagctttttatcgtaaacaatcaatacctaaaaaaatcactcacccagctaaagaggcaatagctaataaaaacaaagataatcctaaaaaagagagtgtttttgaatat
gatttaatcaaagataaacgctttactgaagataagtttttctttcactgtcctattacaatcaattttaaatctagtggagctaataagtttaatgatgaaatcaatttattgctaaa
agaaaaagcaaatgatgttcatatattaagtatagatagaggtgaaagacatttagcttactatactttggtagatggtaaaggcaatatcatcaaacaagatactttcaacat
cattggtaatgatagaatgaaaacaaactaccatgataagcttgctgcaatagagaaagatagggattcagctaggaaagactggaaaaagataaataacatcaaagag
atgaaagagggctatctatctcaggtagttcatgaaatagctaagctagttatagagtataatgctattgtggtttttgaggatttaaattttggatttaaaagagggcgtttcaa
ggtagagaagcaggtctatcaaaagttagaaaaaatgctaattgagaaactaaactatctagttttcaaagataatgagtttgataaaactgggggagtgcttagagcttat
cagctaacagcaccttttgagacttttaaaaagatgggtaaacaaacaggtattatctactatgtaccagctggttttacttcaaaaatttgtcctgtaactggttttgtaaatca
gttatatcctaagtatgaaagtgtcagcaaatctcaagagttctttagtaagtttgacaagatttgttataaccttgataagggctattttgagtttagttttgattataaaaactttg
gtgacaaggctgccaaaggcaagtggactatagctagctttgggagtagattgattaactttagaaattcagataaaaatcataattgggatactcgagaagtttatccaac
taaagagttggagaaattgctaaaagattattctatcgaatatgggcatggcgaatgtatcaaagcagctatttgcggtgagagcgacaaaaagtttttgctaagctaact
agtgtcctaaatactatcttacaaatgcgtaactcaaaaacaggtactgagttagattatctaatttcaccagtagcagatgtaaatggcaatttctttgattcgcgacaggcg
ccaaaaaatatgcctcaagatgctgatgccaatggtgcttatcatattgggctaaaaggtctgatgctactaggtaggatcaaaaataatcaagagggcaaaaaactcaat
ttggttatcaaaaatgaagagtattttgagttcgtgcagaataggaataactaacagtctcagctgcatgctggttgcttgccctgctgccggtcccatgagccagcgatcg
ctcagttcaagaatgtaaatgaatattacaaaccttcagctcgtaacgatttcagatccaaaagatgtagctgttgggtcagccataagtttttcttcaacttctcaagcgatc
gtcgaccatcaacttaaagcatctttacaaagctagagattgaccctagtcgtggacgacttacagattcaggattgatacggccaatcccgatcgcgatcgctctaaatc
cccgtcagtcagagcttcacaatttttagcgaatcttgtggccgcgatcgttgtataagaatgccaaggcaactggataaggttcactaatcgttgctaagcgacagtgaa
ctgcgccaattgcctgacaggcccctctcgtttaacaaacgatttaatgtaaatcattgttaagagtctctcacaatcgagagttttcttgaagaatgatggggacggttcag
gtgcagggtttccctgctagagaatgcgaaaaaaccgcgttctcgttttaggaatcgagagtcaataaaagtcgagaacaggagactggttgaatggatattaatactga
aactgagatcaagcaaaagcattcactaaccccctttcctgttttcctaatcagcccggcatttcgcgggcgatattttcacagctatttcaggagttcagccatgaacgctt
attacattcaggatcgtcttgaggctcagagctgggcgcgtcactaccagcagctcgcccgtgaagagaaagaggcagaactggcagacgacatggaaaaaggcct
gccccagcacctgtttgaatcgctatgcatcgatcatttgcaacgccacggggccagcaaaaaatccattacccgtgcgtttgatgacgatgttgagtttcaggagcgca
tggcagaacacatccggtacatggttgaaaccattgctcaccaccaggttgatattgattcagaggtataaaacgaatgagtactgcactcgcaacgctggctgggaag
ctggctgaacgtgtcggcatggattctgtcgacccacaggaactgatcaccactcttcgccagacggcatttaaaggtgatgccagcgatgcgcagttcatcgcattact
gatcgttgccaaccagtacggccttaatccgtggacgaaagaaatttacgcctttcctgataagcagaatggcatcgttccggtggtgggcgttgatggctggtcccgca
tcatcaatgaaaaccagcagtttgatggcatggactttgagcaggacaatgaatcctgtacatgccggatttaccgcaaggaccgtaatcatccgatctgcgttaccgaat
ggatggatgaatgccgccgcgaaccattcaaaactcgcgaaggcagagaaatcacggggccgtggcagtcgcatcccaaacggatgttacgtcataaagccatgatt
cagtgtgcccgtctggccttcggatttgctggtatctatgacaaggatgaagccgagcgcattgtcgaaaatactgcatacactgcagaacgtcagccggaacgcgaca
tcactccggttaacgatgaaaccatgcaggagattaacactctgctgatcgccctggataaaacatgggatgacgacttattgccgctctgttcccagatatttcgccgcg
acattcgtgcatcgtcagaactgacacaggccgaagcagtaaaagctcttggattcctgaaacagaaagccgcagagcagaaggtggcagcatgacaccggacatta
tcctgcagcgtaccgggatcgatgtgagagctgtcgaacaggggggatgatgcgtggcacaaattacggctcggcgtcatcaccgcttcagaagttcacaacgtgatag
caaaaccccgctccggaaagaagtggcctgacatgaaaatgtcctacttccacaccctgcttgctgaggtttgcaccggtgtggctccggaagttaacgctaaagcact
ggcctggggaaaacagtacgagaacgacgccagaaccctgtttgaattcacttccggcgtgaatgttactgaatccccgatcatctatcgcgacgaaagtatgcgtacc
gcctgctctcccgatggtttatgcagtgacggcaacggccttgaactgaaatgcccgtttacctcccgggatttcatgaagttccggctcggtggtttcgaggccataaag
tcagcttacatggcccaggtgcagtacagcatgtgggtgacgcgaaaaaatgcctggtactttgccaactatgacccgcgtatgaagcgtgaaggcctgcattatgtcgt
gattgagcgggatgaaaagtacatggcgagttttgacgagatcgtgccggagttcatcgaaaaaatggacgaggcactggctgaaattggttttgtatttggggagcaat

ggcgatgacgcatcctcacgataatatccgggtaggcgcaatcactttcgtctactccgttacaaagcgaggctgggtatttcccggcctttctgttatccgaaatccactg
aaagcacagcggctggctgaggagataaataataaacgaggggctgtatgcacaaagcatcttctgttgagttaattgacagctagctcagtcctaggtataatgctagc
gctgatttaggcaaaaacgggtctaagaactttaaataatttctactgttgtagatGTTTCCGTGGAGCTACCGGATGgtctaagaactttaaataatttct
actgttgtagattagcgatttatgaaggtcatttttttgccggtcttagcgcttactctctcgggctttgctagcctgcagcggatcagccgtggtgaaatgctcgaggtgct
gcgccaagactacatccgcaccgcccgtgccaaaggcttgccggagcagcgcgtcatctacgtccacgctctacgcaatgcgatcaatcccctgattacgctcttggg
ctttgagttcgcgaccctgctcagcggcgcttttattgctgaatatttctttaactggccggggctaggccgcttaattttgcaagccgtttttgcgcaggatctctacttggta
atggccagcttgatgatgggtgctgtgatgctgattctgggcaatctgctcgcagatctgctgctgcgctgggtcgatccccgcattcgcctggatgatctgaactaaatt
gttcacgcttctggcttgagtcctgtagtgcgctcaggcagtcttgggctgcgatcgtgcagacgcctagtttttggcaaaccattggctaacattcaggtgctttgactct
ctgtgcaaataaaaaggcgcttctcgtggtcggcatttttctaagtttcttggtggcgatcaatcgcgccttggggtggaaattaccccagagcggatcaatcttgcccac
attggccgcaaggggaatcgcctgcagcttcaggattttgtttagaaaaactcatcgagcatcaaatgaaactgcaatttattcatatcaggattatcaataccatatttttga
aaaagccgtttctgtaatgaaggagaaaactcaccgaggcagttccataggatggcaagatcctggtatcggtctgcgattccgactcgtccaacatcaatacaacctat
taatttcccctcgtcaaaaataaggttatcaagtgagaaatcaccatgagtgacgactgaatccggtgagaatggcaaaagcttatgcatttctttccagacttgttcaacag
gccagccattacgctcgtcatcaaaatcactcgcatcaaccaaaccgttattcattcgtgattgcgcctgagcgagacgaaatacgcgatcgctgttaaaaggacaatta
caaacaggaatcgaatgcaaccggcgcaggaacactgccagcgcatcaacaatattttcacctgaatcaggatattcttctaatacctggaatgctgttttcccggggatc
gcagtggtgagtaaccatgcatcatcaggagtacggataaaatgcttgatggtcggaagaggcataaattccgtcagccagtttagtctgaccatctcatctgtaacatca
ttggcaacgctacctttgccatgtttcagaaacaactctggcgcatcgggcttcccatacaatcgatagattgtcgcacctgattgcccgacattatcgcgagcccatttata
cccatataaatcagcatccatgttggaatttaatcgcggcctggagcaagacgtttcccgttgaatatggctcataacaccccttgtattactgtttatgtaagcagacagttt
tattgttcatgatgatatatttttatcttgtgcaatgtaacatcagagattttgagacacaacgtggctttgttgaataaatcgaacttttgctgagttgaaggatcagtagctaga
ggatcgatcctttttaacccatcacatatacctgccgttcactattatttagtgaaatgagatattatgatattttctgaattgtgattaaaaaggcaactttatgcccatgcaaca
gaaactataaaaaatacagagaatgaaaagaaacagatagattttttagttctttaggcccgtagtctgcaaatccttttatgattttctatcaaacaaaagaggaaaataga
ccagttgcaatccaaacgagagtctaatagaatgaggtcgaaatgggcgatccccaagcaatcaactttgcgacaggtttaattcaacagcttgaccttcgaaaaagac
aagcttcagtcaatgttaaggttattgaccttgacttgaatgacctcaatgcgatcgcgactagtttctcctttggaattggcaacaccctattctcgggaagtggcggtctg
agaacgaccgtaattgataatgcgggctcagtgatccagcccccaaatgacagtgtctttaaccccagtaacttaccagcagggattaacccagcgaatcccttgttggc
cttgccgaatcaattgaccaataatttccttggtgctttatctgcatcaattatcaacaacaaaactaagttgctgactgaccccaccttaatcattcaggaaggagaaagaa
gcgaagtcaaactgactagtcaggttgttcaaaagattgaatcagagactacaaccaacggcagtggcccgccaaccgttagccgaaccattgaccttgcagatgttgg
tctacagttaaccatcaacattgaacgaattgatgacaatggcttcattactctctgccctagCACATTTCCCCGAAAAGTGCCACCTGACGTC
TAAGAAACCATTATTATCATGACATTAACCTATAAAAATAGGCGTATCACGAGGCCCTTTCGTCTTCAAG
AATTCTCATGTTTGACAGCTTATCATCGATAAGCTTTAATGCGGTAGTTTATCACAGTTAAATTGCTAACG
CAGTCAGGCACCGTGTATGAAATCTAACAATGCGCTCATCGTCATCCTCGGCACCGTCACCCTGGATGC
TGTAGGCATAGGCTTGGTTATGCCGGTACTGCCGGGCCTCTTGCGGGATATCGTCCATTCCGACAGCAT
CGCCAGTCACTATGGCGTGCTGCTAGCGCTATATGCGTTGATGCAATTTCTATGCGCACCCGTTCTCGGA
GCACTGTCCGACCGCTTTGGCCGCCGCCCAGTCCTGCTCGCTTCGCTACTTGGAGCCACTATCGACTAC
GCGATCATGGCGACCACACCCGTCCTGTGGATCCTCTACGCCGGACGCATCGTGGCCGGCATCACCGG
CGCCACAGGTGCGGTTGCTGGCGCCTATATCGCCGACATCACCGATGGGGAAGATCGGGCTCGCCACT
TCGGGCTCATGAGCGCTTGTTTCGGCGTGGGTATGGTGGCAGGCCCCGTGGCCGGGGGACTGTTGGGC
GCCATCTCCTTGCATGCACCATTCCTTGCGGCGGCGGTGCTCAACGGCCTCAACCTACTACTGGGCTGC
TTCCTAATGCAGGAGTCGCATAAGGGAGAGCGTCGACCGATGCCCTTGAGAGCCTTCAACCCAGTCAG
CTCCTTCCGGTGGGCGCGGGGCATGACTATCGTCGCCGCACTTATGACTGTCTTCTTTATCATGCAACTC
GTAGGACAGGTGCCGGCAGCGCTCTGGGTCATTTTCGGCGAGGACCGCTTTCGCTGGAGCGCGACGAT
GATCGGCCTGTCGCTTGCGGTATTCGGAATCTTGCACGCCCTCGCTCAAGCCTTCGTCACTGGTCCCGC
CACCAAACGTTTCGGCGAGAAGCAGGCCATTATCGCCGGCATGGCGGCCGACGCGCTGGGCTACGTCT
TGCTGGCGTTCGCGACGCGAGGCTGGATGGCCTTCCCCATTATGATTCTTCTCGCTTCCGGCGGCATCG
GGATGCCCGCGTTGCAGGCCATGCTGTCCAGGCAGGTAGATGACGACCATCAGGGACAGCTTCAAGG
ATCGCTCGCGGCTCTTACCAGCCTAACTTCGATCATTGGACCGCTGATCGTCACGGCGATTTATGCCGC
CTCGGCGAGCACATGGAACGGGTTGGCATGGATTGTAGGCGCCGCCCTATACCTTGTCTGCCTCCCCGC
GTTGCGTCGCGGTGCATGGAGCCGGGCCACCTCGACCTGAATGGAAGCCGGCGGCACCTCGCTAACG
GATTCACCACTCCAAGAATTGGAGCCAATCAATTCTTGCGGAGAACTGTGAATGCGCAAACCAACCCT
TGGCAGAACATATCCATCGCGTCCGCCATCTCCAGCAGCCGCACGCGGCGCATCTCGGGCAGCGTTGG
GTCCTGGCCACGGGTGCGCATGATCGTGCTCCTGTCGTTGAGGACCCGGCTAGGCTGGCGGGGTTGCC

| | |
|---|---|
| | TTACTGGTTAGCAGAATGAATCACCGATACGCGAGCGAACGTGAAGCGACTGCTGCTGCAAAACGTCT<br>GCGACCTGAGCAACAACATGAATGGTCTTCGGTTTCCGTGTTTCGTAAAGTCTGGAAACGCGGAAGTC<br>AGCGCCCTGCACCATTATGTTCCGGATCTGCATCGCAGGATGCTGCTGGCTACCCTGTGGAACACCTAC<br>ATCTGTATTAACGAAGCGCTGGCATTGACCCTGAGTGATTTTTCTCTGGTCCCGCCGCATCCATACCGCC<br>AGTTGTTTACCCTCACAACGTTCCAGTAACCGGGCATGTTCATCATCAGTAACCCGTATCGTGAGCATC<br>CTCTCTCGTTTCATCGGTATCATTACCCCCATGAACAGAAATCCCCCTTACACGGAGGCATCAGTGACC<br>AAACAGGAAAAAACCGCCCTTAACATGGCCCGCTTTATCAGAAGCCAGACATTAACGCTTCTGGAGAA<br>ACTCAACGAGCTGGACGCGGATGAACAGGCAGACATCTGTGAATCGCTTCACGACCACGCTGATGAG<br>CTTTACCGCAGCTGCCTCGCGCGTTTCGGTGATGACGGTGAAAACCTCTGACACATGCAGCTCCCGGG<br>ACGGTCACAGCTTGTCTGTAAGCGGATGCCGGGAGCAGACAAGCCCGTCAGGGCGCGTCAGCGGGTG<br>TTGGCGGGTGTCGGGGCGCAGCCATGACCCAGTCACGTAGCGATAGCGGAGTGTATACTGGCTTAACT<br>ATGCGGCATCAGAGCAGATTGTACTGAGAGTGCACCATATGCGGTGTGAAATACCGCACAGATGCGTA<br>AGGAGAAAATACCGCATCAGGCGCTCTTCCGCTTCCTCGCTCACTGACTCGCTGCGCTCGGTCGTTCG<br>GCTGCGGCGAGCGGTATCAGCTCACTCAAAGGCGGTAATACGGTTATCCACAGAATCAGGGGATAACG<br>CAGGAAAGAACATGTGAGCAAAAGGCCAGCAAAAGGCCAGGAACCGTAAAAAGGCCGCGTTGCTGG<br>CGTTTTTCCATAGGCTCCGCCCCCCTGACGAGCATCACAAAAATCGACGCTCAAGTCAGAGGTGGCGA<br>AACCCGACAGGACTATAAAGATACCAGGCGTTTCCCCCTGGAAGCTCCCTCGTGCGCTCTCCTGTTCC<br>GACCCTGCCGCTTACCGGATACCTGTCCGCCTTTCTCCCTTCGGGAAGCGTGGCGCTTTCTCATAGCTC<br>ACGCTGTAGGTATCTCAGTTCGGTGTAGGTCGTTCGCTCCAAGCTGGGCTGTGTGCACGAACCCCCCG<br>TTCAGCCCGACCGCTGCGCCTTATCCGGTAACTATCGTCTTGAGTCCAACCCGGTAAGACACGACTTAT<br>CGCCACTGGCAGCAGCCACTGGTAACAGGATTAGCAGAGCGAGGTATGTAGGCGGTGCTACAGAGTTC<br>TTGAAGTGGTGGCCTAACTACGGCTACACTAGAAGGACAGTATTTGGTATCTGCGCTCTGCTGAAGCC<br>AGTTACCTTCGGAAAAAGAGTTGGTAGCTCTTGATCCGGCAAACAAACCACCGCTGGTAGCGGTGGTT<br>TTTTTGTTTGCAAGCAGCAGATTACGCGCAGAAAAAAAGGATCTCAAGAAGATCCTTTGATCTTTTCTA<br>CGGGGTCTGACGCTCAGTGGAACGAAAACTCACGTTAAGGGATTTTGGTCATGAGATTATCAAAAAGG<br>ATCTTCACCTAGATCCTTTTAAATTAAAAATGAAGTTTTAAATCAATCTAAAGTATATATGAGTAAACTTG<br>GTCTGACAG |
| pCPF3-pil | ttacgccccgccctgccactcatcgcagtactgttgtaattcattaagcattctgccgacatggaagccatcacagacggcatgatgaacctgaatcgccagcggcatca<br>gcaccttgtcgccttgcgtataatatttgcccatggtgaaaacgggggcgaagaagttgtccatattggccacgtttaaatcaaaactggtgaaactcacccagggattgg<br>ctgagacgaaaaacatattctcaataaaccctttagggaaataggccaggttttcaccgtaacacgccacatcttgcgaatatatgtgtagaaactgccggaaatcgtcgt<br>ggtattcactccagagcgatgaaaacgtttcagtttgctcatggaaaacggtgtaacaagggtgaacactatcccatatcaccagctcaccgtctttcattgccatacgga<br>attccggatgagcattcatcaggcgggcaagaatgtgaataaaggccggataaaacttgtgcttatttttctttacggtctttaaaaaggccgtaatatccagctgaacggt<br>ctggttataggtacattgagcaactgactgaaatgcctcaaaatgttctttacgatgccattgggatatatcaacggtggtatatccagtgatttttttctccattttagcttcctt<br>agctcctgaaaatctcgataactcaaaaaatacgcccggtagtgatcttatttcattatggtgaaagttggaacctcttacgtgccgatcatcactgcccgctttccagtcgg<br>gaaacctgtcgtgccagctgcattaatgaatcggccaacgcgcggggagaggcggtttgcgtattgggcgccagggtggtttttcttttcaccagtgagacgggcaac<br>agctgattgcccttcaccgcctggccctgagagagttgcagcaagcggtccacgctggtttgccccagcaggcgaaaatcctgtttgatggtggttaacggcgggatat<br>aacatgagctgtcttcggtatcgtcgtatcccactaccgagatatccgcaccaacgcgcagcccggactcggtaatggcgcgcattgcgcccagcgccatctgatcgtt<br>ggcaaccagcatcgcagtgggaacgatgccctcattcagcatttgcatggtttgttgaaaaccggacatggcactaaagtcgccttcccgttccgctatcggctgaatttg<br>attgcgagtgagatatttatgccagccagccagacgcagacgcgccgagacagaacttaatgggcccgctaacagcgcgatttgctggtgacccaatgcgaccagat<br>gctccacgcccagtcgcgtaccgtcttcatgggagaaaataatactgttgatgggtgtctggtcagagacatcaagaaataacgccggaacattagtgcaggcagcttc<br>cacagcaatggcatcctggtcatccagcggatagttaatgatcagcccactgacgcgttgcgcgagaagattgtgcaccgccgctttacaggcttcgacgccgcttcgt<br>tctaccatcgacaccaccacgctggcacccagttgatcggcgcgagatttaatcgccgcgacaatttgcgacggcgcgtgcagggccagactggaggtggcaacgc<br>caatcagcaacgactgtttgcccgccagttgttgtgccacgcggttgggaatgtaattcagctccgccatcgccgcttccactttttcccgcgttttcgcagaaacgtggct<br>ggcctggttcaccacgcgggaaacggtctgataagagacaccggcatactctgcgacatcgtataacgttactggtttcacattcaccaccctgaattgactctcttccag<br>tatagatgctagcattatacctaggactgagctagctgtcaaaattgtgagcgctcacaattgatattttggtctgtcgttgcgatcgcccgttgcaggccgacatgaagga<br>tttacactttatgcttccggctcgtatgttGTGTGGaattgtgagcgctcacaattggtaccggtgataccAGCATCGTCTTGATGCCCTTGGCA<br>GCACCCTGCTAAGGAGGCAACAAGatgtcaatttatcaagaatttgttaataaatatagtttaagtaaaactctaagatttgagttaatcccacagggt<br>aaaacacttgaaaacataaaagcaagaggtttgattttagatgatgagaaaagagctaaagactacaaaaaggctaaacaaataattgataaatatcatcagtttttatag<br>aggagatattaagttcggtttgtattagcgaagatttattacaaaactattctgatgtttattttaaacttaaaaagagtgatgatgataatctacaaaaagattttaaaagtgcaa |

aagatacgataaagaaacaaatatctgaatatataaaggactcagagaaatttaagaatttgtttaatcaaaaccttatcgatgctaaaaaagggcaagagtcagatttaatt
ctatggctaaagcaatctaaggataatggtatagaactatttaaagccaatagtgatatcacagatatagatgaggcgttagaaataatcaaatcttttaaaggttggacaac
ttattttaagggttttcatgaaaatagaaaaaatgtttatagtagcaatgatattcctacatctattatttataggatagtagatgataatttgcctaaatttctagaaaataaagct
aagtatgagagtttaaaagacaaagctccagaagctataaactatgaacaaattaaaaaagatttggcagaagagctaacctttgatattgactacaaaacatctgaagtt
aatcaaagagtttttcacttgatgaagtttttgagatagcaaactttaataattatctaaatcaaagtggtattactaaatttaatactattattggtggtaaatttgtaaatggtga
aaatacaaagagaaaaggtataaatgaatatataaatctatactcacagcaaataaatgataaaacactcaaaaaatataaaatgagtgttttatttaagcaaattttaagtga
tacagaatctaaatcttttgtaattgataagttagaagatgatagtgatgtagttacaacgatgcaaagtttttatgagcaaatagcagcttttaaaacagtagaagaaaaatct
attaaagaaacactatctttattatttgatgatttaaaagctcaaaaacttgatttgagtaaaatttattttaaaaatgataaatctcttactgatctatcacaacaagtttttgatgat
tatagtgttattggtacagcggtactagaatatataactcaacaaatagcacctaaaaatcttgataaccctagtaagaaagagcaagaattaatagccaaaaaaactgaa
aaagcaaaatacttatctctagaaactataaagcttgccttagaagaatttaataagcatagagatatagataaacagtgtaggtttgaagaaatacttgcaaactttgcggc
tattccgatgatatttgatgaaatagctcaaaacaaagacaatttggcacagatatctatcaaatatcaaaatcaaggtaaaaaagacctacttcaagctagtgcggaagat
gatgttaaagctatcaaggatcttttagatcaaactaataatctcttacataaactaaaaatatttcatattagtcagtcagaagataaggcaaatattttagacaaggatgagc
atttttatctagtatttgaggagtgctactttgagctagcgaatatagtgcctctttataacaaaattagaaactatataactcaaaagccatatagtgatgagaaatttaagctc
aattttgagaactcgactttggctaatggttgggataaaaataaagagcctgacaatacggcaattttatttatcaaagatgataaatattatctgggtgtgatgaataagaaa
aataacaaaatatttgatgataaagctatcaaagaaaataaaggcgagggttataaaaaaattgtttataaacttttacctggcgcaaataaaatgttacctaaggttttctttt
ctgctaaatctataaaattttataatcctagtgaagatatacttagaataagaaatcattccacacatacaaaaaatggtagtcctcaaaaaggatatgaaaaatttgagtttaa
tattgaagattgccgaaaatttatagatttttataaacagtctataagtaagcatccggagtggaaagattttggatttagattttctgatactcaaagatataattctatagatga
attttatagagaagttgaaaatcaaggctacaaactaacttttgaaaatatatcagagagctatattgatagcgtagttaatcagggtaaattgtacctattccaaatctataat
aaagatttttcagcttatagcaaagggcgaccaaatctacatactttatattggaaagcgctgtttgatgagagaaatcttcaagatgtggtttataagctaaatggtgaggc
agagctttttatcgtaaacaatcaatacctaaaaaaatcactcacccagctaaagaggcaatagctaataaaaacaaagataatcctaaaaaagagagtgtttttgaatat
gatttaatcaaagataaacgctttactgaagataagtttttctttcactgtcctattacaatcaattttaaatctagtggagctaataagtttaatgatgaaatcaatttattgctaaa
agaaaaagcaaatgatgttcatatattaagtatagatagaggtgaaagacatttagcttactatactttggtagatggtaaaggcaatatcatcaaacaagatactttcaacat
cattggtaatgatagaatgaaaacaaactaccatgataagcttgctgcaatagagaaagatagggattcagctaggaaagactggaaaaagataaataacatcaaagag
atgaaagagggctatctatctcaggtagttcatgaaatagctaagctagttatagagtataatgctattgtggtttttgaggatttaaattttggatttaaaagagggcgtttcaa
ggtagagaagcaggtctatcaaaagttagaaaaaatgctaattgagaaactaaactatctagttttcaaagataatgagtttgataaaactgggggagtgcttagagcttat
cagctaacagcaccttttgagacttttaaaaagatgggtaaacaaacaggtattatctactatgtaccagctggttttacttcaaaaatttgtcctgtaactggttttgtaaatca
gttatatcctaagtatgaaagtgtcagcaaatctcaagagttctttagtaagtttgacaagatttgttataaccttgataagggctattttgagtttagttttgattataaaaactttg
gtgacaaggctgccaaaggcaagtggactatagctagctttgggagtagattgattaactttagaaattcagataaaaatcataattgggatactcgagaagtttatccaac
taaagagttggagaaattgctaaaagattattctatcgaatatgggcatggcgaatgtatcaaagcagctatttgcggtgagagcgacaaaaagtttttgctaagctaact
agtgtcctaaatactatcttacaaatgcgtaactcaaaaacaggtactgagttagattatctaatttcaccagtagcagatgtaaatggcaatttctttgattcgcgacaggcg
ccaaaaaaatatgcctcaagatgctgatgccaatggtgcttatcatattgggctaaaaggtctgatgctactaggtaggatcaaaaataatcaagagggcaaaaaactcaat
ttggttatcaaaaatgaagagtattttgagttcgtgcagaataggaataactaaGAAAAGTCTGAAAGTTCTTTACAAAACTCAATCTGC
TTGTTAGATTTACTCACGAGGCTATTAAGTCTCGTAAATAGTTCAACTAAGGACTCATCGCAAAatgccaa
ctatccagcagctaattcgtagcgaacgctcgaaggtacagaagaaaactaaatcccctgccctcaagcaatgtccccaacggcggggagtctgcactagggtttaca
ccaccacccccaaaaagcccaactccgccctccggaaagtggcccgggtacgcctcacctccgggtttgaagtaactgcctatatccctggcattggccacaacctgc
aagaacactccgtagtactaatccggggcggtcgggtaaaagatttgcctggggttcgctaccatattgtgcggggcacgttggacgccaccggagttaaagaccgca
aacagggtcgctccaaatacggcaccaaacgggaaaaagcgaagaaataacagtctcagctgcatgctggttgcttgccctgctgccggtcccatgagccagcgatc
gctcagttcaagaatgtaaatgaatattacaaaccttcagctcgtaacgatttcagatccaaaagatgtagctgttgggtcagccataagtttttcttcaacttctcaagcgat
cgtcgaccatcaacttaaagcatctttacaaagctagagattgaccctagtcgtggacgacttacagattcaggattgatacggccaatcccgatcgcgatcgctctaaat
ccccgtcagtcagagcttcacaatttttagcgaatcttgtggccgcgatcgttgtataagaatgccaaggcaactggataaggttcactaatcgttgctaagcgacagtga
actgcgccaattgcctgacaggcccctctcgtttaacaaacgatttaatgtaaatcattgttaagagtctctcacaatcgagagttttcttgaagaatgatggggacggttca
ggtgcagggtttccctgctagagaatgcgaaaaaaccgcgttctcgttttaggaatcgagagtcaataaaagtcgagaacaggagactggttgaatggatattaatactg
aaactgagatcaagcaaaagcattcactaacccccctttcctgttttcctaatcagcccggcatttcgcgggcgatattttcacagctatttcaggagttcagccatgaacgct
tattacattcaggatcgtcttgaggctcagagctgggcgcgtcactaccagcagctcgcccgtgaagagaaagaggcagaactggcagacgacatggaaaaaggcc
tgccccagcacctgtttgaatcgctatgcatcgatcatttgcaacgccacggggccagcaaaaaatccattacccgtgcgtttgatgacgatgttgagtttcaggagcgc
atggcagaacacatccggtacatggttgaaaccattgctcaccaccaggttgatattgattcagaggtataaaacgaatgagtactgcactcgcaacgctggctgggaa
gctggctgaacgtgtcggcatggattctgtcgacccacaggaactgatcaccactcttcgccagacggcatttaaaggtgatgccagcgatgcgcagttcatcgcatta
ctgatcgttgccaaccagtacggccttaatccgtggacgaaagaaatttacgcctttcctgataagcagaatggcatcgttccggtggtgggcgttgatggctggtcccg
catcatcaatgaaaaccagcagtttgatggcatggactttgagcaggacaatgaatcctgtacatgccggatttaccgcaaggaccgtaatcatccgatctgcgttaccg

aatggatggatgaatgccgccgcgaaccattcaaaactcgcgaaggcagagaaatcacggggccgtggcagtcgcatcccaaacggatgttacgtcataaagccat
gattcagtgtgcccgtctggccttcggatttgctggtatctatgacaaggatgaagccgagcgcattgtcgaaaatactgcatacactgcagaacgtcagccggaacgc
gacatcactccggttaacgatgaaaccatgcaggagattaacactctgctgatcgccctggataaaacatgggatgacgacttattgccgctctgttcccagatatttcgc
cgcgacattcgtgcatcgtcagaactgacacaggccgaagcagtaaaagctcttggattcctgaaacagaaagccgcagagcagaaggtggcagcatgacaccgga
cattatcctgcagcgtaccgggatcgatgtgagagctgtcgaacaggggggatgatgcgtggcacaaattacggctcggcgtcatcaccgcttcagaagttcacaacgt
gatagcaaaaccccgctccggaaagaagtggcctgacatgaaaatgtcctacttccacaccctgcttgctgaggtttgcaccggtgtggctccggaagttaacgctaaa
gcactggcctggggaaaacagtacgagaacgacgccagaaccctgtttgaattcacttccggcgtgaatgttactgaatccccgatcatctatcgcgacgaaagtatgc
gtaccgcctgctctcccgatggtttatgcagtgacggcaacggccttgaactgaaatgcccgtttacctcccgggatttcatgaagttccggctcggtggtttcgaggcca
taaagtcagcttacatggcccaggtgcagtacagcatgtgggtgacgcgaaaaaatgcctggtactttgccaactatgacccgcgtatgaagcgtgaaggcctgcatta
tgtcgtgattgagcgggatgaaaagtacatggcgagttttgacgagatcgtgccggagttcatcgaaaaaatggacgaggcactggctgaaattggttttgtatttgggg
agcaatggcgatgacgcatcctcacgataatatccgggtaggcgcaatcactttcgtctactccgttacaaagcgaggctgggtatttcccggcctttctgttatccgaaat
ccactgaaagcacagcggctggctgaggagataaataataaacgaggggctgtatgcacaaagcatcttctgttgagttaattgacagctagctcagtcctaggtataat
gctagcgctgatttaggcaaaaacgggtctaagaactttaaataatttctactgttgtagatGTTTCCGTGGAGCTACCGGATGgtctaagaactttaaa
taatttctactgttgtagattagcgatttatgaaggtcatttttttgccggtcttagcgcttactctctcgggctttgctagcctgcagcggatcagccgtggtgaaatgctcga
ggtgctgcgccaagactacatccgcaccgcccgtgccaaaggcttgccggagcagcgcgtcatctacgtccacgctctacgcaatgcgatcaatcccctgattacgct
cttgggctttgagttcgcgaccctgctcagcggcgcttttattgctgaatatttctttaactggccggggctaggccgcttaattttgcaagccgtttttgcgcaggatctctac
ttggtaatggccagcttgatgatgggtgctgtgatgctgattctgggcaatctgctcgcagatctgctgctgcgctgggtcgatccccgcattcgcctggatgatctgaact
aaattgttcacgcttctggcttgagtcctgtagtgcgctcaggcagtcttgggctgcgatcgtgcagacgcctagtttttttggcaaaccattggctaacattcaggtgctttg
actctctgtgcaaataaaaaggcgcttctcgtggtcggcatttttttctaagtttcttggtggcgatcaatcgcgccttggggtggaaattaccccagagcggatcaatcttgc
ccacattggccgcaaggggaatcgcctgcagcttcaggattttgtttagaaaaactcatcgagcatcaaatgaaactgcaatttattcatatcaggattatcaataccatattt
ttgaaaaagccgtttctgtaatgaaggagaaaactcaccgaggcagttccataggatggcaagatcctggtatcggtctgcgattccgactcgtccaacatcaatacaac
ctattaatttcccctcgtcaaaaataaggttatcaagtgagaaatcaccatgagtgacgactgaatccggtgagaatggcaaaagcttatgcatttctttccagacttgttca
acaggccagccattacgctcgtcatcaaaatcactcgcatcaaccaaaccgttattcattcgtgattgcgcctgagcgagacgaaatacgcgatcgctgttaaaaggaca
attacaaacaggaatcgaatgcaaccggcgcaggaacactgccagcgcatcaacaatattttcacctgaatcaggatattcttctaatacctggaatgctgttttcccggg
gatcgcagtggtgagtaaccatgcatcatcaggagtacggataaaatgcttgatggtcggaagaggcataaattccgtcagccagtttagtctgaccatctcatctgtaac
atcattggcaacgctacctttgccatgtttcagaaacaactctggcgcatcgggcttcccatacaatcgatagattgtcgcacctgattgcccgacattatcgcgagcccat
ttatacccatataaatcagcatccatgttggaatttaatcgcggcctggagcaagacgtttcccgttgaatatggctcataacaccccttgtattactgtttatgtaagcagac
agttttattgttcatgatgatatatttttatcttgtgcaatgtaacatcagagattttgagacacaacgtggctttgttgaataaatcgaacttttgctgagttgaaggatcagtag
ctagaggatcgatcctttttaacccatcacatatacctgccgttcactattatttagtgaaatgagatattatgatattttctgaattgtgattaaaaaggcaactttatgcccatg
caacagaaactataaaaaatacagagaatgaaaagaaacagatagattttttagttctttaggcccgtagtctgcaaatccttttatgattttctatcaaacaaaagaggaaa
atagaccagttgcaatccaaacgagagtctaatagaatgaggtcgaaatgggcgatccccaagcaatcaactttgcgacaggtttaattcaacagcttgaccttcgaaaa
agacaagcttcagtcaatgttaaggttattgaccttgacttgaatgacctcaatgcgatcgcgactagtttctcctttggaattggcaacaccctattctcgggaagtggcgg
tctgagaacgaccgtaattgataatgcgggctcagtgatccagcccccaaatgacagtgtctttaaccccagtaacttaccagcagggattaacccagcgaatcccttgtt
ggccttgccgaatcaattgaccaataatttccttggtgctttatctgcatcaattatcaacaacaaaactaagttgctgactgaccccaccttaatcattcaggaaggagaaa
gaagcgaagtcaaactgactagtcaggttgttcaaaagattgaatcagagactacaaccaacggcagtggcccgccaaccgttagccgaaccattgaccttgcagatg
ttggtctacagttaaccatcaacattgaacgaattgatgacaatggcttcattactctctgccctagCACATTTCCCCGAAAAGTGCCACCTGAC
GTCTAAGAAACCATTATTATCATGACATTAACCTATAAAAATAGGCGTATCACGAGGCCCTTTCGTCTTC
AAGAATTCTCATGTTTGACAGCTTATCATCGATAAGCTTTAATGCGGTAGTTTATCACAGTTAAATTGCT
AACGCAGTCAGGCACCGTGTATGAAATCTAACAATGCGCTCATCGTCATCCTCGGCACCGTCACCCTGG
ATGCTGTAGGCATAGGCTTGGTTATGCCGGTACTGCCGGGCCTCTTGCGGGATATCGTCCATTCCGACA
GCATCGCCAGTCACTATGGCGTGCTGCTAGCGCTATATGCGTTGATGCAATTTCTATGCGCACCCGTTCT
CGGAGCACTGTCCGACCGCTTTGGCCGCCGCCCAGTCCTGCTCGCTTCGCTACTTGGAGCCACTATCG
ACTACGCGATCATGGCGACCACACCCGTCCTGTGGATCCTCTACGCCGGACGCATCGTGGCCGGCATC
ACCGGCGCCACAGGTGCGGTTGCTGGCGCCTATATCGCCGACATCACCGATGGGGAAGATCGGGCTCG
CCACTTCGGGCTCATGAGCGCTTGTTTCGGCGTGGGTATGGTGGCAGGCCCCGTGGCCGGGGGACTGT
TGGGCGCCATCTCCTTGCATGCACCATTCCTTGCGGCGGCGGTGCTCAACGGCCTCAACCTACTACTGG
GCTGCTTCCTAATGCAGGAGTCGCATAAGGGAGAGCGTCGACCGATGCCCTTGAGAGCCTTCAACCCA
GTCAGCTCCTTCCGGTGGGCGCGGGGCATGACTATCGTCGCCGCACTTATGACTGTCTTCTTTATCATGC
AACTCGTAGGACAGGTGCCGGCAGCGCTCTGGGTCATTTTCGGCGAGGACCGCTTTCGCTGGAGCGC

| | |
|---|---|
| | GACGATGATCGGCCTGTCGCTTGCGGTATTCGGAATCTTGCACGCCCTCGCTCAAGCCTTCGTCACTGG<br>TCCCGCCACCAAACGTTTCGGCGAGAAGCAGGCCATTATCGCCGGCATGGCGGCCGACGCGCTGGGCT<br>ACGTCTTGCTGGCGTTCGCGACGCGAGGCTGGATGGCCTTCCCCATTATGATTCTTCTCGCTTCCGGCG<br>GCATCGGGATGCCCGCGTTGCAGGCCATGCTGTCCAGGCAGGTAGATGACGACCATCAGGGACAGCTT<br>CAAGGATCGCTCGCGGCTCTTACCAGCCTAACTTCGATCATTGGACCGCTGATCGTCACGGCGATTTAT<br>GCCGCCTCGGCGAGCACATGGAACGGGTTGGCATGGATTGTAGGCGCCGCCCTATACCTTGTCTGCCTC<br>CCCGCGTTGCGTCGCGGTGCATGGAGCCGGGCCACCTCGACCTGAtgccggtcttagcgcttactctctcgggctttgctagc<br>ctgcagcggatcagccgtggtgaaatgctcgaggtgctgcgccaagactacatccgcaccgcccgtgccaaaggcttgccggagcagcgcgtcatctacgtccacg<br>ctctacgcaatgcgatcaatcccctgattacgctcttgggctttgagttcgcgaccctgctcagcggcgcttttattgctgaatatttctttaactggccggggctaggccgc<br>ttaattttgcaagccgtttttgcgcaggatctctacttggtaatggccagcttgatgatgggtgctgtgatgctgattctgggcaatctgctcgcagatctgctgctgcgctgg<br>gtcgatccccgcattcgcctggatgatctgaactaaattgttcacgcttctggcttgagtcctgtagtgcgctcaggcagtcttgggctgcgatcgtgcagacgcctagtttt<br>ttggcaaaccattggctaacattcaggtgctttgactctctgtgcaaataaaaaggcgcttctcgtggtcggcatttttttctaagtttcttggtggcgatcaatcgcgccttgg<br>ggtggaaattaccccagagcggatcaatcttgcccacattggccgcaaggggaatcgcctgcagcttcaggattttgtATGGAAGCCGGCGGCACCT<br>CGCTAACGGATTCACCACTCCAAGAATTGGAGCCAATCAATTCTTGCGGAGAACTGTGAATGCGCAAA<br>CCAACCCTTGGCAGAACATATCCATCGCGTCCGCCATCTCCAGCAGCCGCACGCGGCGCATCTCGGGC<br>AGCGTTGGGTCCTGGCCACGGGTGCGCATGATCGTGCTCCTGTCGTTGAGGACCCGGCTAGGCTGGCG<br>GGGTTGCCTTACTGGTTAGCAGAATGAATCACCGATACGCGAGCGAACGTGAAGCGACTGCTGCTGCA<br>AAACGTCTGCGACCTGAGCAACAACATGAATGGTCTTCGGTTTCCGTGTTTCGTAAAGTCTGGAAACG<br>CGGAAGTCAGCGCCCTGCACCATTATGTTCCGGATCTGCATCGCAGGATGCTGCTGGCTACCCTGTGGA<br>ACACCTACATCTGTATTAACGAAGCGCTGGCATTGACCCTGAGTGATTTTTCTCTGGTCCCGCCGCATC<br>CATACCGCCAGTTGTTTACCCTCACAACGTTCCAGTAACCGGGCATGTTCATCATCAGTAACCCGTATC<br>GTGAGCATCCTCTCTCGTTTCATCGGTATCATTACCCCCATGAACAGAAATCCCCCTTACACGGAGGCAT<br>CAGTGACCAAACAGGAAAAAACCGCCCTTAACATGGCCCGCTTTATCAGAAGCCAGACATTAACGCTT<br>CTGGAGAAACTCAACGAGCTGGACGCGGATGAACAGGCAGACATCTGTGAATCGCTTCACGACCACG<br>CTGATGAGCTTTACCGCAGCTGCCTCGCGCGTTTCGGTGATGACGGTGAAAACCTCTGACACATGCAG<br>CTCCCGGGACGGTCACAGCTTGTCTGTAAGCGGATGCCGGGAGCAGACAAGCCCGTCAGGGCGCGTC<br>AGCGGGTGTTGGCGGGTGTCGGGGCGCAGCCATGACCCAGTCACGTAGCGATAGCGGAGTGTATACTG<br>GCTTAACTATGCGGCATCAGAGCAGATTGTACTGAGAGTGCACCATATGCGGTGTGAAATACCGCACAG<br>ATGCGTAAGGAGAAAATACCGCATCAGGCGCTCTTCCGCTTCCTCGCTCACTGACTCGCTGCGCTCGGT<br>CGTTCGGCTGCGGCGAGCGGTATCAGCTCACTCAAAGGCGGTAATACGGTTATCCACAGAATCAGGGG<br>ATAACGCAGGAAAGAACATGTGAGCAAAAGGCCAGCAAAAGGCCAGGAACCGTAAAAAGGCCGCGT<br>TGCTGGCGTTTTTCCATAGGCTCCGCCCCCCTGACGAGCATCACAAAAATCGACGCTCAAGTCAGAGG<br>TGGCGAAACCCGACAGGACTATAAAGATACCAGGCGTTTCCCCCTGGAAGCTCCCTCGTGCGCTCTCC<br>TGTTCCGACCCTGCCGCTTACCGGATACCTGTCCGCCTTTCTCCCTTCGGGAAGCGTGGCGCTTTCTCA<br>TAGCTCACGCTGTAGGTATCTCAGTTCGGTGTAGGTCGTTCGCTCCAAGCTGGGCTGTGTGCACGAACC<br>CCCCGTTCAGCCCGACCGCTGCGCCTTATCCGGTAACTATCGTCTTGAGTCCAACCCGGTAAGACACG<br>ACTTATCGCCACTGGCAGCAGCCACTGGTAACAGGATTAGCAGAGCGAGGTATGTAGGCGGTGCTACA<br>GAGTTCTTGAAGTGGTGGCCTAACTACGGCTACACTAGAAGGACAGTATTTGGTATCTGCGCTCTGCTG<br>AAGCCAGTTACCTTCGGAAAAAGAGTTGGTAGCTCTTGATCCGGCAAACAAACCACCGCTGGTAGCG<br>GTGGTTTTTTTGTTTGCAAGCAGCAGATTACGCGCAGAAAAAAAGGATCTCAAGAAGATCCTTTGATC<br>TTTTCTACGGGGTCTGACGCTCAGTGGAACGAAAACTCACGTTAAGGGATTTTGGTCATGAGATTATCA<br>AAAAGGATCTTCACCTAGATCCTTTTAAATTAAAAATGAAGTTTTAAATCAATCTAAAGTATATATGAGT<br>AAACTTGGTCTGACAG |

**Table S5**. Descriptions of strains used or constructed in this study.

| Name | Description |
|---|---|
| WT | wild type *Synechococcus elongatus* UTEX 2973 |
| WT-pSES-ori | pSES-ori; *spe*$^R$ in WT |
| WT-pSES | pSES; *spe*$^R$ in WT |
| WT-pSEL-ori | pSEL-ori; *spe*$^R$ in WT |
| WT-pSEL | pSEL; *spe*$^R$ in WT |
| WT-pRSF-ori | pRSF-ori; *spe*$^R$ in WT |
| WT-pRSF | pRSF; *spe*$^R$ in WT |
| WT-NSI-single | NSI::pSI-single; *spe*$^R$ in WT |
| WT-pSES-2k | pSES-2k; *spe*$^R$ in WT |
| WT-pSES-4k | pSES-4k; *spe*$^R$ in WT |
| WT-pSES-8k | pSES-8k; *spe*$^R$ in WT |
| WT-pSES-12k | pSES-12k; *spe*$^R$ in WT |
| WT-pSEL-2k | pSEL-2k; *spe*$^R$ in WT |
| WT-pSEL-4k | pSEL-4k; *spe*$^R$ in WT |
| WT-pSEL-8k | pSEL-8k; *spe*$^R$ in WT |
| WT-pSEL-12k | pSEL-12k; *spe*$^R$ in WT |
| WT-pRSF-2k | pRSF-2k; *spe*$^R$ in WT |
| WT-pRSF-4k | pRSF-4k; *spe*$^R$ in WT |
| WT-pRSF-8k | pRSF-8k; *spe*$^R$ in WT |
| WT-pRSF-12k | pRSF-12k; *spe*$^R$ in WT |
| WT-pSES-lacZ | pSES-lacZ; *spe*$^R$ in WT |
| WT-pSES-ori-lacZ | pSES-ori-lacZ; *spe*$^R$ in WT |
| WT-pSEL-lacZ | pSEL-lacZ; *spe*$^R$ in WT |
| WT-pSEL-L-lacZ | pSEL-L-lacZ; *spe*$^R$ in WT |
| WT-pRSF-lacZ | pRSF-lacZ; *spe*$^R$ in WT |
| WT-pRSF-ori-lacZ | pRSF-ori-lacZ; *spe*$^R$ in WT |
| WT-pSEL-SacB | pSEL-sacB; *km*$^R$ *i*n WT |
| WT-pSEL-SepT$_2$ | pSEL-sepT2; *km*$^R$ in WT |
| WTR-pSEL-68Rpsl | pSEL-68rpsl; *km*$^R$ in WT-RPSLm |
| WT-pSEL-TetA | pSEL-tetA; *km*$^R$ in WT |
| WT-RPSLm | M744_12300 K43R; pBR322-rpslm; *str*$^R$ in WT |
| WT-RPSLm | M744_12300 K43R; pBR322-rpslm-cm; *str*$^R$ in WT |
| WT-RPSLm | M744_12300 K43R; pBR322-rpslm-sepT2; *str*$^R$ in WT |
| WT-PIL-KM | *pilMNOQ* :: *km*; pBR-pil-km; *km*$^R$ in WT |
| WT-PIL-KM | *pilMNOQ* :: *km*; pBR-pil-km-sepT2; *km*$^R$ in WT |
| WTR-PIL-68rpsl | *pilMNOQ* :: $P_{psba2}$-68rpsl; pBR-pil-sepT2; *cm*$^R$ in WT-RPSLm |
| WTR-PIL-KM | M744_12300 K43R; pBR322-rpslm-sepT2; *str*$^R$ in WT-PIL-KM |
| WTR-pilNm | *pilN* :: *pilNm*; pBR-pilNm-sepT2; in WTR-PIL-68rpsl |
| WTR-pilNm | *pilN :: pilNm*; pBR3-pilNm-sepT2; in WTR-PIL-KM |
| WT-LLACCP-pilN-km | *pilMNOQ* :: *km*; WpLLACCP-pilN-km; *cm*$^R$ in WT |
| WT-LTHOCP-pilN-km | *pilMNOQ* :: *km*; WpLTHOCP-pilN-km; *cm*$^R$ in WT |
| WT-LTRCCP-pilN-km | *pilMNOQ* :: *km*; WpLTRCCP-pilN-km; *cm*$^R$ in WT |

| | |
|---|---|
| WT-SLACCP-pilN-km | *pilMNOQ* :: *km*; WpSLACCP-pilN-km; $cm^R$ in WT |
| WT-STRCCP-pilN-km | *pilMNOQ* :: *km*; WpSTRCCP-pilN-km; $cm^R$ in WT |
| WT-STHOCP-pilN-km | *pilMNOQ* :: *km*; WpSTHOCP-pilN-km; $cm^R$ in WT |
| WT-RFLACCP-pilN-km | *pilMNOQ* :: *km*; WpRFLACCP-pilN-km; $cm^R$ in WT |
| WT-RFTRCCP-pilN-km | *pilMNOQ* :: *km*; WpRFTRCCP-pilN-km; $cm^R$ in WT |
| WT-RFTHOCP-pilN-km | *pilMNOQ* :: *km*; WpRFTHOCP-pilN-km; $cm^R$ in WT |
| WT-RLACCP-pilN-km | *pilMNOQ* :: *km*; WpRLACCP-pilN-km; $cm^R$ in WT |
| WT-RTRCCP-pilN-km | *pilMNOQ* :: *km*; WpRTRCCP-pilN-km; $cm^R$ in WT |
| WT-RTHOCP-pilN-km | *pilMNOQ* :: *km*; WpRTHOCP-pilN-km; $cm^R$ in WT |
| WT-STHOCP | pSTHOCP; $cm^R$ in WT |
| WT-LTHOCP | pLTHOCP; $cm^R$ in WT |
| WT-RTHOCP | pRTHOCP; $cm^R$ in WT |
| WT-LTHOCP-pSCR-pilN-km | *pilMNOQ :: km*; pSCR-pilN-km; $spe^R$ in WT-LTHOCP |
| WT-RTHOCP-pSCR-pilN-km | *pilMNOQ :: km*; pSCR-pilN-km; $spe^R$ in WT-RTHOCP |
| WT-RTHOCP-pLCR-pilN-km | *pilMNOQ :: km*; pLCR-pilN-km; $spe^R$ in WT-RTHOCP |
| WT-STHOCP-pLCR-pilN-km | *pilMNOQ :: km*; pLCR-pilN-km; $spe^R$ in WT-STHOCP |
| WT-LTHOCP-pRCR-pilN-km | *pilMNOQ :: km*; pLCR-pilN-km; $spe^R$ in WT-RTHOCP |
| WT-STHOCP-pRCR-pilN-km | *pilMNOQ :: km*; pSCR-pilN-km; $spe^R$ in WT-STHOCP |
| WT-LRTHOCPET | pRSF-recET; $erm^R$ in WT-LTHOCP |
| WT-LRTHOCPET-pSCR-pilN-km | *pilMNOQ* :: *km*; pSCR-pilN-km; $spe^R$ in WT-LRTHOCPET |
| WT-LRTHOCPRED | pRSF-λred; $erm^R$ in WT-pLTHOCP |
| WT-LRTHOCPRED-pSCR-pilN-km | *pilMNOQ* :: *km*; pSCR-pilN-km; $spe^R$ in WT-LRTHOCPRED |
| WT-LRTHOCPJ1 | pRSF-atRecJ1; ermR in WT-pLTHOCP |
| WT-LRTHOCPJ1-pSCR-pilN-km | *pilMNOQ* :: *km*; pSCR-pilN-km; $spe^R$ in WT-LRTHOCPJ1 |
| WT-LRTHOCPJ2 | pRSF-atRecJ2; ermR in WT-LTHOCP |
| WT-LRTHOCPJ2-pSCR-pilN-km | *pilMNOQ* :: *km*; pSCR-pilN-km; $spe^R$ in WT-LRTHOCPJ2 |
| WT-LTRC2OTCP | pLTRC2OTCP; $cm^R$ in WT |
| WT-LRTRC2OTCPRED | pRSF-λred; $erm^R$ in WT-LTRC2OTCP |
| WT-LRTRC2OTCPRED-pSCR-pilN-km | *pilMNOQ* :: *km*; pSCR-pilN-km; $spe^R$ in WT-LRTRC2OTCPRED |
| WT-LLAC2OTCP | pLLAC2OTCP; $cm^R$ in WT |
| WT-LRLAC2OTCPRED | pRSF-λred; $erm^R$ in WT-LLAC2OTCP |
| WT-LRLAC2OTCPRED-pSCR-pilN-km | *pilMNOQ* :: *km*; pSCR-pilN-km; $spe^R$ in WT-LRLAC2OTCPRED |
| WT-LLAC1OTCP | pLLAC1OTCP; $cm^R$ in WT |
| WT-LRLAC1OTCPRED | pRSF-λred; $erm^R$ in WT-LLAC1OTCP |
| WT-LRLAC2OTCPRED-pSCR-pilN-km | *pilMNOQ* :: *km*; pSCR-pilN-km; $spe^R$ in WT-LRLAC1OTCPRED |
| WT-PIL-KM | *pilMNOQ* :: *km*; pCPF1-pil in WT |
| WT-NSI-1K | *NSI* :: *1K*; pCPF1-NSI in WT |
| WT-NSII-1K | *NSII* :: *1K*; pCPF1-NSII in WT |
| WT-NSIII-1K | *NSIII* :: *1K*; pCPF1-NSIII in WT |
| WT-13825-1K | *M744_13825*:: *1K*; pCPF1-13825 in WT |
| WTR-NSI-8K | *NSI* :: *8K*; pCPF3-NSI-1 in WT-RPSLm |
| WTR-NSI-8K | *NSI* :: *8K*; pCPF3-NSI-2 in WT-RPSLm |
| WTR-NSII-8K | *NSII* :: *8K*; pCPF3-NSII-1 in WT-RPSLm |
| WTR-NSII-8K | *NSII* :: *8K*; pCPF3-NSII-2 in WT-RPSLm |
| WTR-NSIII-8K | *NSIII* :: *8K*; pCPF3-NSIII-1 in WT-RPSLm |

| | |
|---|---|
| WTR-NSIII-8K | *NSIII* :: *8K*; pCPF3-NSIII-2 in WT-RPSLm |
| WTR-13825-8K | *M744_13825*:: *8K*; pCPF3-13825-1 in WT-RPSLm |
| WTR-13825-8K | *M744_13825*:: *8K*; pCPF3-13825-2 in WT-RPSLm |
| WTR-06260-8K | *M744_06260*:: *7K*; pCPF3-06260-1 in WT-RPSLm |
| WTR-06260-8K | *M744_06260*:: *1K*; pCPF3-06260-2 in WT-RPSLm |
| WTR-ANL-8K | *pANL* :: *8K*; pCPF3-anl-1 in WT-RPSLm |
| WTR-ANL-8K | *pANL* :: *8K*; pCPF3-anl-1 in WT-RPSLm |
| WTR-PIL-KM | *pilMNOQ* :: *km*; pCPF3-pil in WT-RPSLm |
| WTR-NSIII-68rpsl | *NSIII* :: *68rpsl*; pBR-NSIII-sepT2; $km^R$ in WT-RPSLm |
| WTR-B3 | *NSIII* :: *cscB*; pBR-NSIII-cscB in WTR-NSIII-68rpsl |
| WTR-B3 | *NSIII* :: *cscB*; pBR3-NSIII-cscB in WT-RPSLm |
| WTR-B3 | *NSIII* :: *cscB*; pCPF3-NSIII-cscB in WT-RPSLm |
| WTR-BSP13 | *NSI* :: *sps-spp*; pCPF3-NSI-sps-spp in WTR-B3 |
| WTR-BSPi3 | *inva* :: *sps-spp*; pCPF3-inva-sps-spp in WTR-B3 |
| WTR-BSPi3-2cugP | *NSII* :: *cugP*; pCPF3-NSII-cugP in WTR-BSPi3 |
| WTR-BSPGi23 | *NSII* :: *glgC*; pCPF3-NSII-glgC in WTR-BSPi3 |
| WTR-BSPi3-2ptglgC | *NSII* :: *ptglgC*; pCPF3-NSII-ptglgC in WTR-BSPi3 |
| WTR-BSPi3-ScugP | pSES-cugP; $spe^R$ in WTR-BSPi3 |
| WTR-BSPi3-SglgC | pSES-glgC; $spe^R$ in WTR-BSPi3 |
| WTR-BSPi3-SptglgC | pSES-ptglgC; $spe^R$ in WTR-BSPi3 |
| WTR-B3 | *NSIII* :: *cscB*; pCPF3-NSIII-cscB-1200-300 in WT-RPSLm |
| WTR-B3 | *NSIII* :: *cscB*; pCPF3-NSIII-cscB-800-300 in WT-RPSLm |
| WTR-B3 | *NSIII* :: *cscB*; pCPF3-NSIII-cscB-500-300 in WT-RPSLm |
| WTR-B3 | *NSIII* :: *cscB*; pCPF3-NSIII-cscB-500-500 in WT-RPSLm |
| WTR-B3 | *NSIII* :: *cscB*; pBR3-NSIII-cscB-1200-300 in WT-RPSLm |
| WTR-B3 | *NSIII* :: *cscB*; pBR3-NSIII-cscB-800-300 in WT-RPSLm |
| WTR-B3 | *NSIII* :: *cscB*; pBR3-NSIII-cscB-500-300 in WT-RPSLm |
| WTR-B3 | *NSIII* :: *cscB*; pBR3-NSIII-cscB-500-500 in WT-RPSLm |
